# A cell-based degrader assessment platform facilitates discovery of functional NUDT5 PROTACs

**DOI:** 10.1101/2025.06.13.659494

**Authors:** Seher Alam, Maria J. Pires, Gabriel Tidestav, Ann-Sofie Jemth, Julia Eklund, Fredrik Klingegård, Rémi Caraballo, Massimiliano Gaetani, Dante Rotilli, Siebe van den Elzen, Eri van Berkum, Olov Wallner, Thomas Gustafsson, Jonas Malmström, Per I. Arvidsson, Nicholas C.K. Valerie, Mikael Altun

## Abstract

Targeted protein degradation (TPD) via PROTACs and molecular glues holds significant therapeutic promise but demands detailed mechanistic evaluation in live cells to fully understand compound behavior and optimize efficacy. Here, we present an integrated, cell-first platform that combines a modular degradation assay with E3 ligase target engagement readouts for comprehensive assessment of TPD molecules in cells and use it to evaluate PROTACs towards NUDT5. To mimic endogenous degradation conditions and TPD amenability, we established a fusion protein expression system consisting of a lysine-free FKBP12 F36V PROTAC handle (FKBPV_K0_) and used a HiBiT/akaLuc dual luciferase reporter to accurately measure degradation dynamics. This set-up identified a VHL-dependent NUDT5 PROTAC, DDD2, that induced robust NUDT5 degradation, despite impaired NUDT5 binding *in vitro* and *in cellulo*, but no CRBN-dependent degraders. NUDT5 lysine availability mapping with DDD2 and FKBP12 F36V-directed PROTACs suggested that the CRL4^CRBN^ complex is more sensitive to target lysine accessibility than CRL2^VHL^, which may have implications for E3 ligase choice and therapeutic resistance. CeTEAM drug biosensors were also established towards CRBN and VHL to quantitatively monitor degrader engagement in living cells and confirmed that the tested CRBN-directed NUDT5 PROTACs poorly engaged the E3. All together, this platform provides a versatile and scalable framework for TPD molecule discovery in a cellular context.

## INTRODUCTION

Targeted protein degradation (TPD) is a therapeutic approach that harnesses the cell’s ubiquitin-proteasome system to eliminate disease-causing proteins, including those considered undruggable ^1–3^. Small molecules such as PROTACs (proteolysis-targeting chimeras) and molecular glue degraders promote degradation by recruiting target proteins to E3 ubiquitin ligases. PROTACs are bifunctional molecules that form a ternary complex between a target and an E3 ligase, while molecular glues stabilize interactions between a ligase and a neo-substrate. Among E3 ligases, Cereblon (CRBN) and von Hippel–Lindau (VHL) are the most commonly exploited in PROTAC design ^4–6^. CRBN was originally identified as the target of immunomodulatory drugs (IMiDs) like thalidomide, which reprogram CRBN to degrade neo-substrates, a mechanism later adapted for PROTACs. Although both ligases are broadly expressed, they differ in substrate specificity and context-dependent activity. Both PROTACs and molecular glues act catalytically, enabling potent, substoichiometric knockdown of proteins. Beyond therapeutic potential, they offer reversible, non-genetic tools for probing protein functions. However, their degradation efficiency depends on multiple factors, including cellular uptake, ligand affinity, ternary complex stability, and proteasomal processing ^5,7–11^, highlighting the complexity of degrader design.

Among the key challenges is optimizing PROTACs for drug-like properties, as their relatively large molecular size often limits intracellular availability, necessitating strategies such as physicochemical optimization or prodrug design ^10,12–15^. Moreover, linker engineering is frequently a rate-limiting step, as even minor modifications can significantly affect ternary complex formation and degradation outcomes ^15,16^. Biophysical assays are widely used for initial screening and mechanistic characterization of PROTACs in vitro. While valuable, these assays do not capture key aspects of the cellular context, including subcellular localization and crowding, post-translational modifications, proteasomal capacity, or feedback regulation, all of which are critical for effective degradation in cells. Thus, in-cell assays are essential to validate and fully characterize PROTAC efficacy and mechanism of action ^8^.

Current in-cell evaluations often focus on degradation endpoints using techniques such as immunoblotting or fluorescent tagging. However, western blotting has limited throughput and is semi-quantitative, while bulky fluorescent protein tags may interfere with degradation or limited dynamic range ^17,18^. Notably, commercial direct-to-biology platforms based on HiBiT/NanoLuc degradation kinetics, such as those developed by Promega, have become a widely adopted benchmark for quantitative, high-throughput profiling of degrader activity in cells ^8^. To overcome these limitations, we developed a novel cellular platform for the real-time and high-throughput evaluation of PROTAC-mediated degradation, E3 ligase engagement, and compound uptake in a biologically relevant setting ^8^. Our approach integrates scalable degradation assays and CeTEAM (Cellular Target Engagement by Accumulation of Mutants) technology ^19,20^, a live cell assay that quantifies drug-target binding *via* ligand-induced stabilization of POI reporter constructs. We employ a fragment complementation reporter system based on the NanoLuciferase-derived NanoBiT platform ^21^, which provides high sensitivity and a broad dynamic range, for quantifying the levels of the POI and assessing E3 ligase engagement.

Distinctively, we incorporated the FKBP12 F36V (FKBPV) tag into our degradation assay system, which has cognate AP1867-derived PROTACs (dTAG-13 and FKBP degrader 3) ^22^, as an internal control for target protein degradability. To maximize the screening specificity and likeness to endogenous protein conditions, we engineered a lysine-free variant, FKBPV_K0_, that is compatible with FKBPV PROTACs to ensure target protein-specific lysine usage for degradation.

To demonstrate the utility of our platform, we synthesized PROTACs targeting NUDT5, a nucleotide-metabolizing enzyme that promotes ADP-ribosylation and supports cancer cell survival, particularly in breast cancer ^23,24^. This approach enabled the discovery of a suite of VHL-dependent NUDT5 PROTACs and highlighted that synthesized CRBN-directed counterparts failed to engage CRBN in cells and degrade NUDT5. Furthermore, we demonstrate inherent accessibility limitations of the CRL2^VHL^ and CRL4^CRBN^ complexes by NUDT5 lysine mapping, which has implications for degrader efficacy and resistance. Thus, our platform enables robust, high-throughput screening and detailed mechanistic characterization of such next-generation degraders in a physiologically relevant context.

## RESULTS

### Development of a modular and scalable cell-based degradation assay

To identify functional PROTACs for NUDT5, we first set out to develop a robust, reporter-based degradation assay that is scalable, flexible and sensitive. To this end, we engineered a fusion construct we named pENTR2x, in which the POI is linked to a reporter protein and a bicistronic reference protein to enable real-time quantification of POI levels in living cells (Fig. 1A). Our goal was predicated on flexibility so that interchangeable tagging components allow the assay to be adapted to different targets and purposes while maintaining the native-like behavior of the POI, such as replacement with fluorescent reporters (Fig. 1B, Supplemental Fig. 7).

**Figure 1.**
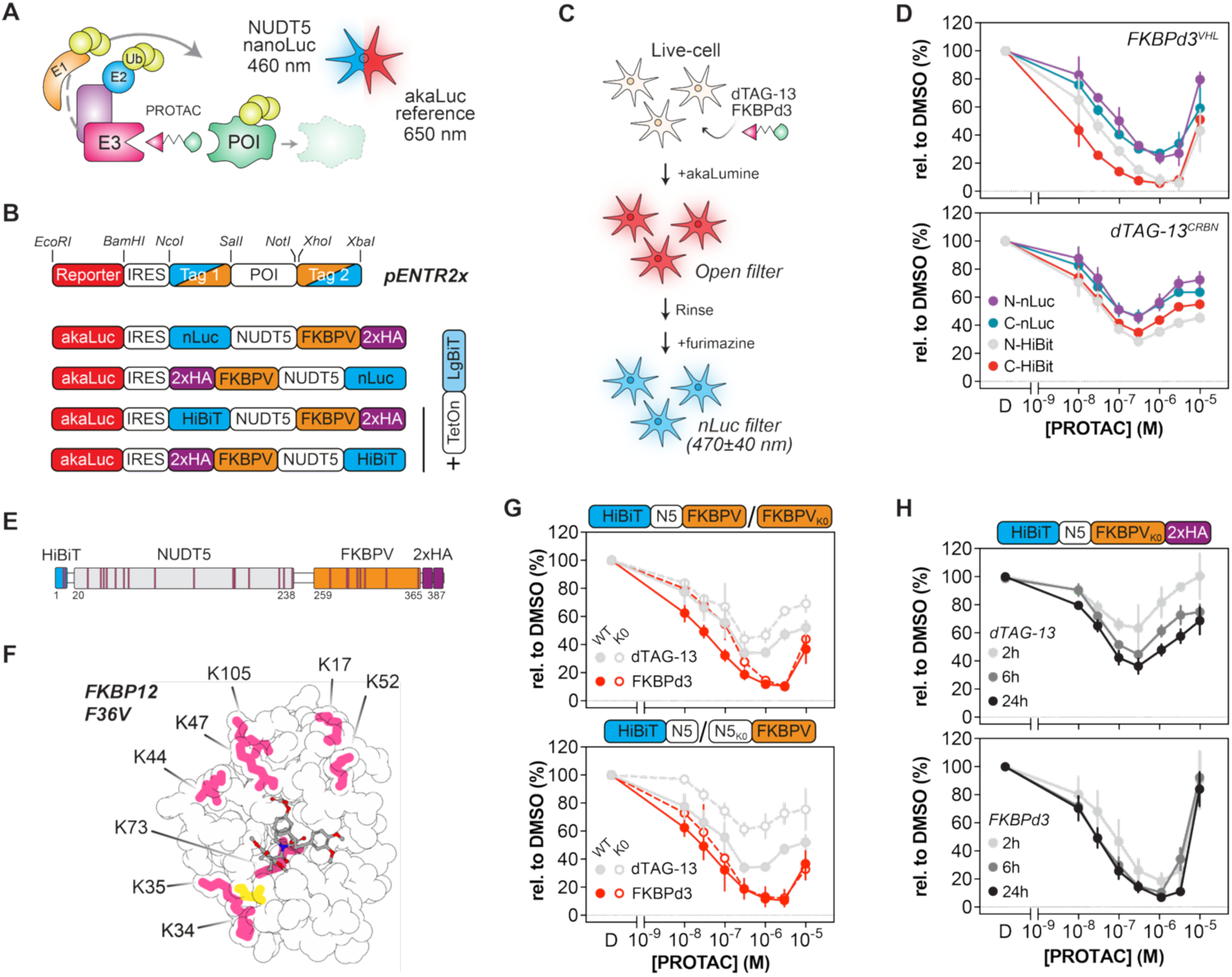
Establishment of dual reporter assay for robust screening of PROTAC-induced degradation. **A.** Illustration of the dual-luminescent reporter system for monitoring PROTAC induced degradation of a protein of interest (POI). AkaLuc provides a reference signal at 650 nm, while POI abundance is measured via NanoLuc (nLuc) emission at 460 nm. **B**. Schematic of a multimodal bicistronic expression vector (pENTR2X) with annotated restriction sites for modular assembly of degradation reporter assays. Four NUDT5 reporter variants are shown fused to nanoluciferase (nLuc) or HiBiT and FKBP12 F36V (FKBPV) with a 2x hemagglutinin (HA) peptide tag. Conditional LgBiT expression was from a separate vector via the Tet-On system for HiBiT-tagged POI detection. **C.** Illustration of single assay read out of dual-luminescence reporter by sequential measurement of signal of akaLuc with open filter and subsequent nLuc signal at 470±40 nM. **D.** Dose–response curves of normalized nLuc signal in U-2 OS cells expressing NUDT5 degradation reporter tagged with nLuc or HiBiT at the N- or C-terminus with akaLuc for internal normalization. Cells were treated for 24 h with FKBPd3 and dTAG-13 concentration gradients and luminescence was measured as described in C. **E.** Schematic of the finalized NUDT5 fusion protein indicating orientation of tags. Internal lysines are highlighted in pink. **F.** Structure of the FKBPV-F36V protein complexed with the FKBP ligand AP1867 made with Protein Imager^62^ (PDBID: 1BL4). The F36V point mutation is highlighted in yellow. Lysine residues (pink) were substituted to arginine in the lysine-free variant (FKBPV_K0_). **G**. Dose-response of dTAG-13 and FKBPd3 treatment for 24 hours in U-2 OS cells expressing FKBPV- and FKBPV_K0_-tagged degradation reporters. **H.** Dose-response of finalized degradation reporter after dTAG-13 or FKBPd3 incubation for 2, 6 and 24h. For panels D, G, and H, LgBiT was induced by doxycycline (Dox) (0.75 µg/mL) prior to 24 h compound treatment. nLuc signal was normalized to akaLuc and expressed as percentage relative to the DMSO control. Data are presented as mean ± SD (n = 3), each experiment performed in triplicate.

To enable sensitive, scalable monitoring of protein degradation in live cells, we modified a dual-luminescent reporter system reported previously that combines a NanoLuciferase (NanoLuc) readout with a red-shifted akaluciferase (akaLuc) reference ^19,25^. Here, the all-in-one format co-expresses a POI tagged with NanoLuc and as well as akaLuc separated by an IRES (Fig. 1B). Alternatively, to generate a smaller fusion protein, NanoLuc was replaced with the HiBiT fragment, while expression of the high affinity LgBiT fragment was controlled by a separate, doxycycyline (DOX)-regulable vector. In addition, we employed an FKBP12 F36V tag, which has cognate PROTACs using either CRL2^VHL^ (FKBP degrader-3 [FKBPd3]) or CRL4^CRBN^ (dTAG-13) ^22,26^, on the opposite terminus. Rather than relying on genetic loss-of-function approaches, we used FKBPV-mediated degradation as an internal, chemically inducible control to establish whether a given POI fusion was amenable to targeted degradation. To set a baseline, we treated cells expressing the NUDT5 fusion proteins in each orientation with either FKBPV PROTAC and sequentially detected akaLuc and NanoLuc for ratiometric quantification of POI levels (Fig. 1C and D). Both yielded dose-dependent fusion protein degradation with a characteristic Hook effect – saturation of both target and E3 binding sites resulting in loss of ternary complex formation. For both PROTACs, HiBiT tagging consistently yielded higher degradation efficiencies than full-length NanoLuc, supporting the use of split-luciferase designs for improved assay sensitivity. For NUDT5, the C-terminal configuration (HiBiT-POI–FKBPV) was further developed. This configuration was chosen based on comparisons across multiple POIs tested with N- and C-terminally tagged FKBPVK0 fusions, which generally showed improved degradation performance with C-terminal tagging (Supplemental Fig. 2B); for NUDT5 specifically, no substantial difference between orientations was observed, and the C-terminal configuration was retained for consistency across the panel.

Self-complementing split reporters maintain high binding affinities *in cellulo* with minimal loss of signal. The small fragment of split fluorescent proteins can be used in tandem to linearly increase signal to noise, which can be limiting in fluorescence applications. To see if this also applied for HiBiT/LgBiT, we compared signal intensities for single vs. tandem (3×) HiBiT tags. While 3×HiBiT increased luminescence, it reduced degradation efficiency, possibly due to LgBiT-mediated POI stabilization (Supplemental Fig. 1A-B). As expected, removal of FKBPV made the fusion protein completely resistant to FKBPV PROTACs. LgBiT complementation appeared to stabilize single HiBiT fusion protein abundance by western blot (Supplemental Fig. 1C), suggesting it could affect degradation dynamics. We then turned to the HiBiT/LgBiT interaction for possible improvements, as multiple HiBiT variants with varying affinities for LgBiT have been reported ^21, 27^, including a lysine-less version (HiBiT_K0_). NanoLuc signal was not appreciably affected until a reported affinity difference of 1000-fold (NP), which translated to minimal impact on degradation efficiency, unless binding was severely impaired (Supplemental Fig. 1D-F, peptide NP). As other HiBiT variants showed no distinct advantage, we continued with the parental HiBiT sequence (peptide 86). This includes the lysine-free HiBiT-K0 variant, which showed no meaningful difference in degradation efficiency relative to the parental HiBiT sequence (peptide 86) for either VHL- or CRBN-mediated degradation, indicating that lysines within the HiBiT tag do not appreciably contribute to the degradation kinetics observed here, we saw no clear benefit to switching to HiBiT-K0 or other variants for our reporter system.

The addition of HiBiT and FKBPV tags also brings associated lysine residues that are not present in endogenous proteins (2x and 8x lysine residues, respectively; Fig. 1E). To reduce potential artifacts arising from targeting available lysine residues in FKBPV, we designed a lysine-free variant (FKBPV_K0_) and again tested FKBPV PROTACS (Fig. 1F). FKBPV_K0_ degradation dynamics were similar to the original FKBPV (Fig. 1G). Comparatively, removal of lysine residues from NUDT5 (NUDT5_K0_) worsened dTAG-13-mediated degradation, and the combination blocked nearly all degradation of the fusion protein by either PROTAC (Supplemental Fig. 2A). As reported previously ^27^, while HiBiT_K0_ was functional (Supplemental Fig. 1E, F), its apparent contribution to degradation was minimal, so it was abandoned to avoid potential NanoLuc detection issues. We then confirmed the amenability of this system for multiple proteins, including KDM4C, LSD1 and KRAS^G12D^, demonstrating broad applicability and revealing target-specific differences in degradation efficiency and tag placement sensitivity (Supplemental Fig. 2B). This version of the assay displayed robust degradation kinetics for NUDT5 fusions in living cells, suggesting it was a suitable proxy for screening new degraders (Fig. 1H).

Building on the flexible reporter platform developed for assessing PROTAC-mediated degradation of POI targets, we customized this system to directly evaluate E3 ligase-recruiting compounds in a POI-independent manner, enabling functional characterization of E3 recruiters in the absence of target ligands within TPD applications. To achieve this, we implemented the HaloTag/HaloPROTAC system ^28–30^, which enables covalent recruitment of E3 ligases via chloroalkane-functionalized ligands. We replaced FKBPV with HaloTag7 in our reporter system and employed a suite of fluorescent and luminescent reporters (Supplemental Fig. 3A, B). As before, degradation efficiency was highest for split reporters (Supplemental Fig. 3C-F). The system could reliably discriminate between active and inactive HaloPROTACs, including the lack of a Hook effect due to covalent binding (Supplemental Fig. 3 G–H), confirming its usefulness for E3 ligase degradation profiling. Together, this optimized dual-luminescent system provides a sensitive, modular, and scalable platform for evaluating degrader potency and dynamics across diverse targets and E3 ligases.

### Direct-to-cell screening for functional NUDT5 PROTACs

We used our optimized dual-luminescence assay to evaluate a series of novel PROTACs designed to degrade NUDT5. A targeted library of 16 PROTACs was synthesized by conjugating NUDT5-binding warheads to either VHL (VH032) or CRBN (pomalidomide) E3 ligase recruiters. For NUDT5, we used TH5427 and related equipotent analogs, including TH10184, which has improved solubility ^24,31^ (Fig. 2A). Each compound was initially tested for degradation of NUDT5 in 3×HiBiT-FKBPV reporter cells (Supplemental Fig. 4A), which yielded a promising hit (DDD2). Like before, degradation by FKBPV PROTACs was robust but worse than with 1×HiBiT platforms (Supplemental Fig. 4B). Removal of FKBPV from the fusion protein also had no effect on degradation dynamics by DDD2, suggesting that the added bulk *per se* did not contribute to degradation. We then sought to confirm that DDD2 was a functional PROTAC by western blot, which permitted us to follow both endogenous and exogenous forms of NUDT5 (Fig. 2B). Indeed, dTAG-13 selectively degraded the fusion protein, while DDD2 degraded both forms of NUDT5 at 2 µM. For a more quantitative comparison, these were re-run in a seven-point concentration gradient in the optimized HiBiT-FKBPV_K0_ cells (Fig. 2C). Again, the FKBPV PROTACs were robust degraders (with FKBPd3 having a superior D_max_), and only DDD2 effectively degraded the NUDT5 reporter. DDD2 was hence identified as our lead compound consisting of the warhead TH10184 connected to the VHL recruiting ligand, VH032, through a saturated aliphatic eight carbon linker (Fig 2D).

**Figure 2.**
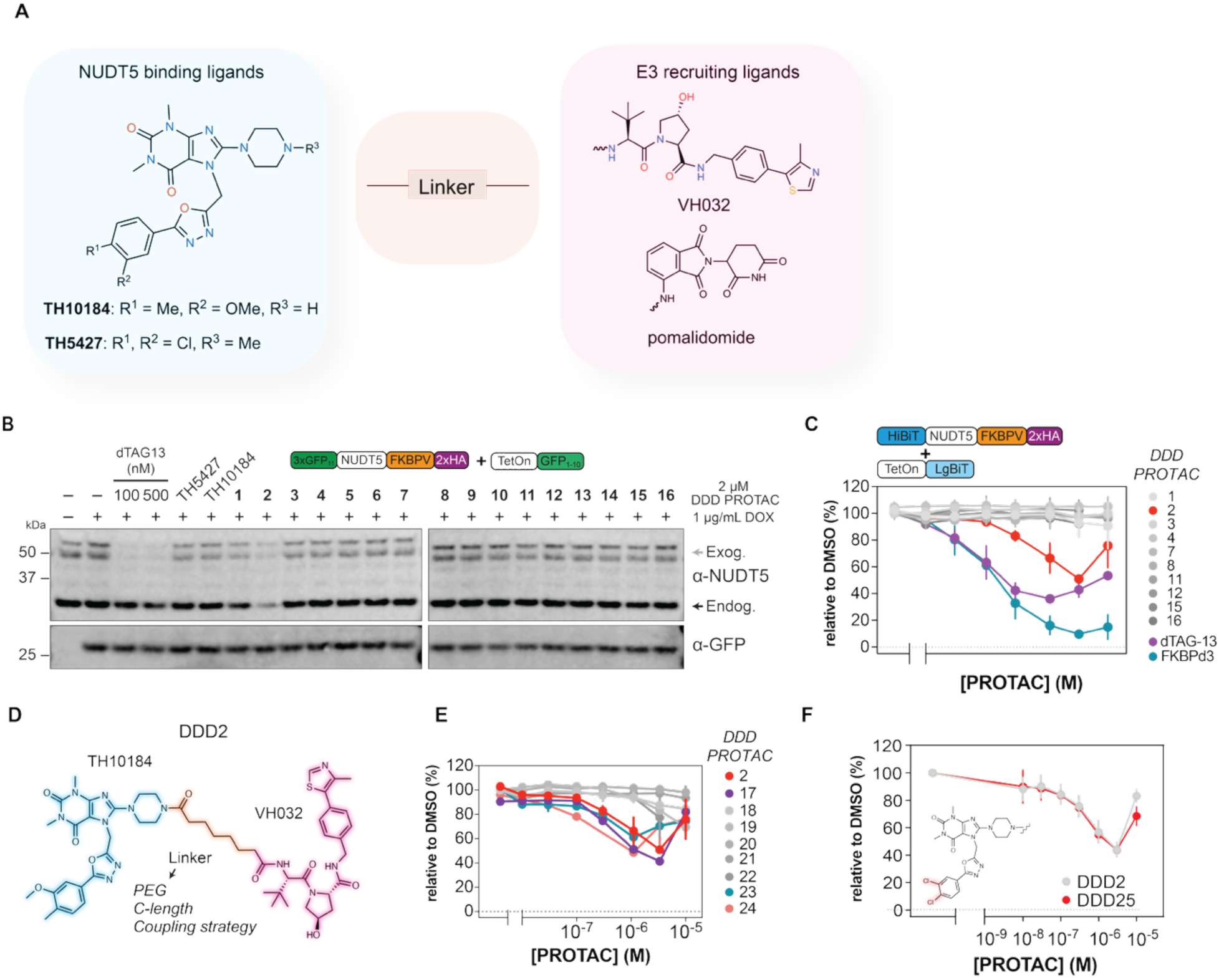
Screening and optimization of NUDT5-specific PROTACs. **A.** Simplified overview of ligands used to generate NUDT5 PROTACs. **B.** Representative western blot (n = 2) of 3×GFP11 tagged NUDT5 degradation reporter in U-2 OS cells treated with dTAG-13 (100 or 500 nM), NUDT5 inhibitors TH5427, TH10184, or DDD PROTACs 1–16 (2 µM) for 24 h (after induction of Dox (1 µg/mL)) to induce GFP1-10. Grey arrow – exogenous NUDT5; black arrow - endogenous NUDT5. **C.** Degradation effect of DDD1-16 compounds and control PROTACs, (dTAG-13 or FKBPd3) at concentrations ranging from 0.01µM and 10 µM). **D.** Chemical structure of DDD2, featuring the NUDT5 binding moiety TH10184, the VHL recruiting ligand VH032 and linker variations included in the SAR study **E.** Dose-response of DDD2 SAR derivatives incubated for 24 hours. **F.** Comparison of the effect of degradation between DDD2 and its analogue DDD25, with altered NUDT5 warhead (TH5427). In panels **C, E,** and **F,** U-2 OS cells expressing dual luminescence degradation reporter were used. LgBiT was induced by doxycycline (0.75 µg/mL) prior to 24 h compound treatment. nLuc signal was normalized to akaLuc and expressed as percentage of DMSO control. Data represent mean ±SD (n = 2), each experiment performed in triplicate.

With DDD2 as a promising lead, we sought to improve the degradation efficiency by varying the linker length, composition, and attachment chemistry for a structure–activity relationship (SAR) study. Intriguingly, degradation efficiency proved to be highly sensitive to the nature of the linker. Simply shorting the linker with one carbon atom (DDD17) or increasing up to two carbons (DDD23-24) preserved degradation activity, while replacing two carbons with oxygen (2 x PEG) abolished degradation efficiency. Replacing the piperazine in TH10184 with a piperidine analog (DDD18 and DDD19) also led to a drastic decrease in degradation efficiency, with a slight difference depending on the amide orientation. Likewise, triazole linkers (DDD20 and DDD21) were poorly tolerated. Again, a small change in activity was seen based on where the triazole ring was placed (Fig. 2E, Supplemental Table 1). To investigate alternative warheads with this geometry (DDD2), we replaced TH10184 with the cognate NUDT5 inhibitor, TH5427, and generated an analog called DDD25 (Fig. 2F). As expected, DDD25 induced degradation of NUDT5 comparable to DDD2, consistent with the high structural similarity and common binding mode in the NUDT5 active site ^24^. However, DDD25 exhibited significant solubility and formulation issues, including frequent precipitation and poor stability in DMSO (Table 1), which likely limited its cellular availability and performance. Although DDD2 and DDD25 showed similar degradation potential, DDD2 was selected as the lead compound due to its more favorable solubility and stability profile. In parallel, we tried to create a CRBN-based degrader to have both E3 ligases as recruiting units towards NUDT5, but we could not identify a functional degrader despite employing a variety of linkers (n=25 (DDD26-DDD50); Supplemental Fig. 4C, Supplemental Table 1).

**Table 1.**
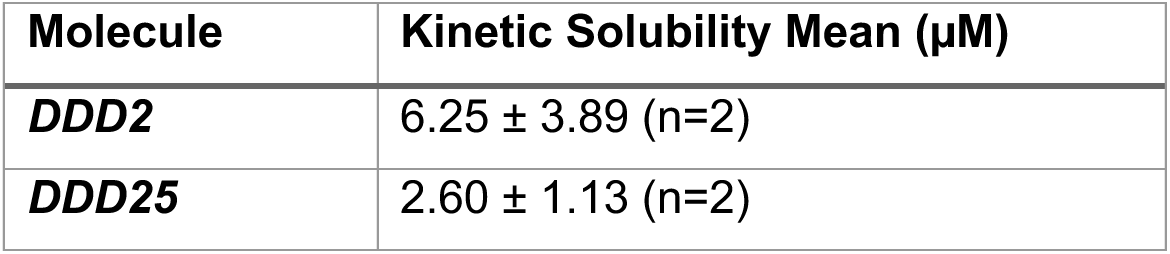
DDD2 Vs. DDD25 Kinetic solubility mean (µM)

### Validation and characterization of DDD2 towards endogenous NUDT5

After identifying DDD2 as our lead compound, we validated its activity on endogenous NUDT5. Treatment with DDD2 resulted in a concentration-dependent reduction in NUDT5 protein levels, with degradation detectable as early as 4 hours and a peak steady-state maximal effect achieved 24 hours after addition (DC_50(24hrs)_: 12 nM, D_max(24hrs)_: 91%; Fig. 3A, B). Remarkably, NUDT5 remained suppressed for at least 96 hours after treatment, indicating sustained intracellular activity (Fig. 3A, B).

**Figure 3.**
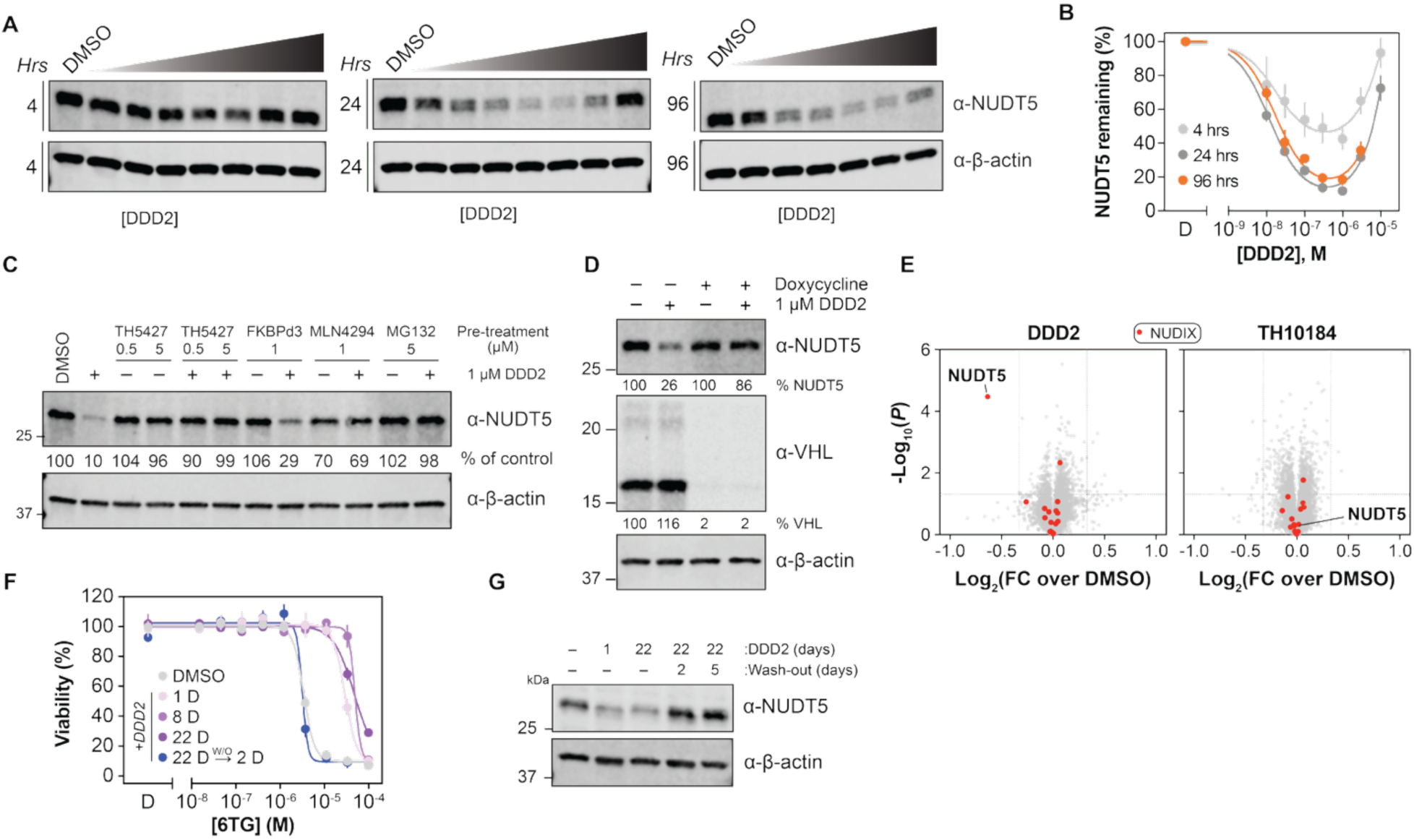
Validation of DDD2 as a selective degrader of endogenous NUDT5. **A.** Representative western blot showing concentration- and time-dependent degradation of endogenous NUDT5 in CCRF-CEM cells treated with 0, 0.01, 0.03, 0.1, 0.3, 1, 3 or 10 µM concentrations of DDD2 for 4, 24, and 96 hours. B. Quantification of NUDT5 protein levels from experiments shown in panel A (mean ± SD, n=2). **C.** Chemical rescue of NUDT5 degradation with DDD2 by 1h pre-treatment with TH5427, FKBPd3, MLN4294, or MG132 for an additional 24h (mean ± SD, n=2). **D.** Genetic rescue of NUDT5 degradation in U-2 OS cells expressing VHL shRNA. Knockdown was induced by DOX for 48hrs prior to DDD2 addition. **E.** Volcano plots showing global proteomic profiling of HEK293T cells treated with DDD2 or TH10184 after 24h. The X-axis indicates log₂ fold change over DMSO, and the Y-axis shows –log₁₀(p-value). A fold-change cut-off of 1.25 and a p-value threshold of 0.05 were applied. Members of the NUDIX family are highlighted in red; all others in grey. **F.** Viability curves from 1-, 8-, and 22-day dose–response treatments with 6-TG on cells with depleted NUDT5, including DDD2 washout on day 20, show that cells become resistant to 6-TG toxicity when NUDT5 is depleted and regain sensitivity to 6-TG when NUDT5 levels are restored. **G.** Western blot analysis of NUDT5 levels from cells treated with DDD2 for 1 or 22 days, including DDD2 washout after 17 and 20 days to show depletion and rebound of the protein. β-actin was used as a loading control.

For further insights on why DDD2 may be effective, we turned to assays for characterizing compound binding and activity with purified NUDT5. To this end, we performed differential scanning fluorimetry (DSF) and enzyme-coupled NUDT5 inhibition assays ^24^. TH5427 and TH10184 stabilized NUDT5 to thermal denaturation (up to ΔTm ≈ 10°C), whereas all PROTACs, including DDD2, were significantly worse – although NUDT5 warheads with dichlorophenyl moieties were consistently better binders (Supplemental Fig. 5A, B). Similarly, both DDD2 and DDD25 were significantly less potent NUDT5 inhibitors compared to parental molecules (∼50-200-fold activity loss; Supplemental Fig. 5C), although all compounds showed good selectivity for NUDT5 over other members of the NUDIX family (Supplemental Fig. 5D). Notably, TH10184 had reduced activity towards MTH1 compared to TH5427, suggesting a better selectivity profile within the NUDIX family. Cellular thermal shift assays (CETSA) confirmed the concentration-dependent stabilization of intracellular NUDT5 by TH5427 and TH10184, but not by DDD2 (Supplemental Fig. 5E, F). Interestingly, although TH10184 showed comparable biochemical inhibition and thermal shift to TH5427, it presented a weaker binding in the cellular context, revealing important differences between the binding profiles *in vitro* and in living cells.

Despite significantly weaker binding to NUDT5 *in vitro* and *in cellulo*, DDD2 still maintained robust cellular degradation activity. To confirm that degradation was *via* NUDT5 binding and CRL2^VHL^, we performed a series of chemical and genetic rescue experiments. Pre-treatment with TH5427 rescued NUDT5 levels, confirming that binding to the active site is required, whereas partial rescue was also observed by FKBPd3 competition. DDD2-mediated degradation was completely blocked by the neddylation inhibitor, MLN4294, and the proteasome inhibitor, MG132, consistent with CRL2^VHL^-mediated proteasomal degradation (Fig. 3C). This dependence was further confirmed by shRNA-mediated silencing of VHL, which abolished DDD2-induced degradation (Fig. 3D). These results demonstrate that in vitro binding data alone cannot predict the efficiency of degradation and emphasize the value of cellular assays to reveal functional PROTAC activity.

To assess specificity of DDD2, we performed quantitative proteomics analyses in HEK293T cells treated with DDD2 or TH10184. Over 11087 proteins were identified with 9877 confidently quantified (using at least two unique peptides) across all tested conditions and replicates. NUDT5 was the only protein that was significantly reduced by DDD2 treatment, while no changes were observed in other members of the NUDIX family (Fig. 3E). In contrast, TH10184 alone had no effect on protein levels, confirming that degradation requires both target binding and recruitment of the ubiquitin-proteasome machinery. Taken together, these results confirm that DDD2 is a selective and potent degrader of NUDT5 in cells.

To connect NUDT5 degradation to a defined cellular phenotype, we leveraged the recently reported role of NUDT5 in 6-thioguanine (6-TG)-mediated toxicity, in which loss of NUDT5 confers resistance to 6-TG through a nonenzymatic mechanism ^32^. When we treated cells with DDD2 and measured 6-TG dose–response viability over time, we found that they were resistant to 6-TG on days 1, 8, and 22, paralleling the sustained NUDT5 depletion achieved by DDD2 (Fig. 3F). To test whether this resistance was a direct and reversible consequence of NUDT5 depletion, we withdrew DDD2 after long-term treatment and rechallenged cells with 6-TG following a 2- or 5-day washout. Resistance was completely lost after 2 days and was largely restored to DMSO-like sensitivity (Fig. 3F), mirroring the recovery of NUDT5 protein levels over the same washout periods (Fig. 3G). These results show that DDD2-induced NUDT5 degradation confers reversible, phenotypically relevant resistance to 6-TG, directly linking target engagement to a biological outcome.

### NUDT5 lysine mapping reveals that CRL4^CRBN^ is more sensitive to lysine availability than CRL2^VHL^

The efficiency of PROTAC degradation is not only influenced by the E3 ligase, but also by factors such as the availability of lysine residues for ubiquitination, the plasticity of the E3 complex, and the dynamics of ternary complex formation. To investigate the contribution of specific lysine residues to NUDT5 degradation, we applied a systematic lysine mapping strategy using our luminescence-based degradation assay. NUDT5 contains 15 lysine residues, mostly in solvent-exposed regions (Fig. 4A), 11 of these 15 lysine sites have previously been reported to be ubiquitinated (Fig. 4B, Supplemental Table 2). We first generated four hybrid NUDT5 proteins (M1–M4) in which either most of the N-terminal or most of the C-terminal region was blocked from ubiquitylation by mutating lysine (K) residues to arginine (R) residues (Fig. 4C), to evaluate whether either terminus of NUDT5 is important for PROTAC-mediated degradation. M1–M4 were expressed using the FKBPV_K0_ construct to eliminate any lysine contribution from the tag. Cells were treated with 0.5 µM dTAG-13^CRBN^, 1 µM FKBPd3^VHL^ (both binding FKBPV), or 1 µM DDD2^VHL^ (binding NUDT5), and degradation was quantified by luminescence. Both VHL-based PROTACs retained nearly full degradation capacity toward all hybrids, whereas the CRBN-based PROTAC lost the ability to efficiently degrade hybrids in which the C-terminal lysines were blocked from ubiquitylation (Fig. 4D, Supplementary Table 3). To further identify which lysines are preferred for degradation, we generated a series of single-lysine NUDT5 mutants by replacing all but one lysine residue with arginine, yielding 15 unique constructs. These were expressed using the FKBPV_K0_ construct to eliminate any lysine contribution from the tag. Cells were treated with 0.5 µM dTAG-13^CRBN^, 1 µM FKBPd3^VHL^ (binding FKBPV), or 1 µM DDD2^VHL^ (binding NUDT5), and degradation was measured by luminescence (Fig. 4E, Supplementary Table 3). As before, wild-type NUDT5 (NUDT5^WT^) showed robust degradation: ∼65% with dTAG-13, ∼90% with FKBPd3, and ∼50% with DDD2, whereas NUDT5^K0^ showed no degradation, validating the assay and confirming lysine dependence.

**Figure 4.**
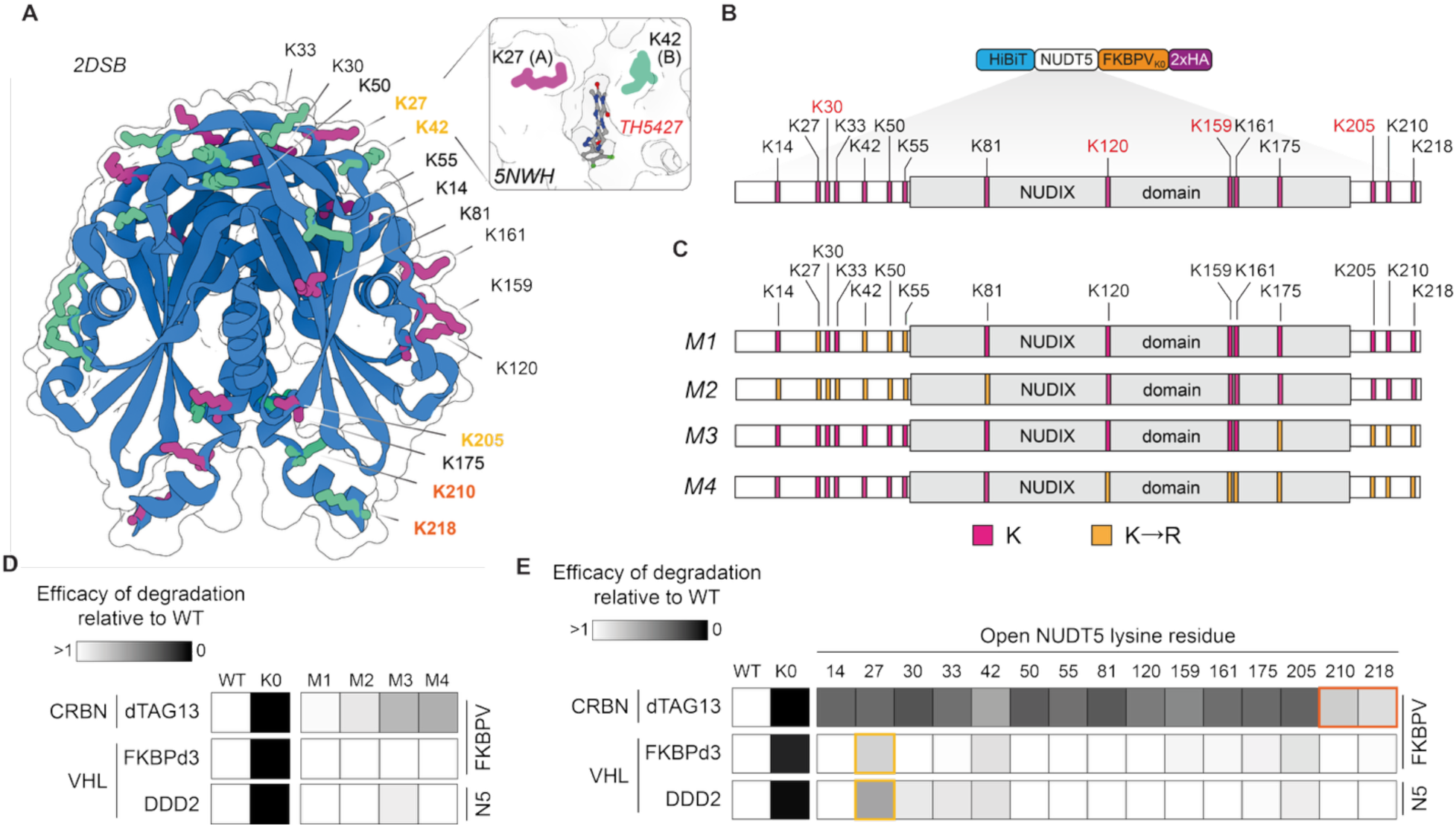
E3 ligase-dependent sensitivity to lysine positioning in NUDT5 degradation. **A.** 3D structure illustration of the NUDT5 homodimer highlighting all 15 lysine residues (pink and green denoting chain A/B specificity) and zoom-in showing the binding site of NUDT5 warhead TH5427 in proximity to K27 and K42 (made with Protein Imager; PDBID: 2DSB and 5NWH). **B.** Linear schematic of the NUDT5 protein sequence and domains, with lysine (K) residues highlighted in pink, K residues previously identified as ubiquitinated (11 of 15) in black, and non-ubiquitinated K residues in red (see Supplementary Table 2). **C.** Linear schematic of different mutant versions of the NUDT5 protein (M1–M4), with lysine residues highlighted in pink and K- to-arginine (R) mutations in orange. **D.** Heat map of residual NanoLuc (nLuc) signal in U-2 OS cells expressing different NUDT5 mutants (M1–M4), wild type (WT), or lysine-null (K0) reporter constructs after 24 h of treatment with 0.5 µM dTAG-13, 1 µM FKBPd3, or 1 µM DDD2. LgBiT expression was induced with doxycycline (0.75 µg/mL) 24 h before compound treatment. Data are normalized to WT degradation within each PROTAC (Supplementary Table 3, see Methods section for further details). **E.** Heat map of residual nLuc signal in U-2 OS cells expressing single-lysine NUDT5 mutants, wild-type (WT), or lysine-null (K0) reporter constructs after 24 h treatment with 0.5 µM dTAG-13, 1 µM FKBPd3, or 1 µM DDD2 LgBiT expression was induced with doxycycline (0.75 µg/mL) 24 h prior to compound treatment. Data is normalized to the degradation of WT within each PROTAC (Supplementary Table 3, see Methods section for further details). Residues K210 and K218 (highlighted in orange) supported robust degradation comparable to WT across all PROTACs, whereas residues K27 (highlighted in yellow) and to some degree K42, and K205 indicated that these positions are not preferred for degradation. These results suggest that VHL-based degraders can use most lysines for ubiquitination if needed, whereas the CRBN-based degrader is less flexible and strongly prefers the two C-terminal lysines. nLuc signals were normalized to akaLuc, expressed as a percentage relative to the DMSO control, and shown relative to WT degradation. Data represent mean ± SD (n=3).

Of the single lysine mutants, K210 and K218 (near the C-terminus) supported degradation comparable to wild type for all PROTACs (Fig. 4E). These residues are located in the accessible C-terminal tail region and are known ubiquitination sites *^33^* thus explaining their permissiveness for degradation. VHL- and CRBN-dependent FKBPV PROTACs exhibited different sensitivity profiles, reflecting deviations in their respective E3 complexes and recruitment strategies. The CRBN-recruiting dTAG-13 was particularly sensitive to lysine position, with most mutants showing degradation of only ∼20–40%. In contrast, the VHL-recruiting FKBPd3 maintained robust degradation (∼75-90%) in most mutants, except for K27, K42 and K205. Surprisingly, DDD2 showed a nearly identical profile to FKBPd3, despite targeting a different region of the fusion protein. K27 and K42 are directly adjacent to the NUDT5 active site (Fig. 4A), implying that this region is intrinsically more difficult to use by the CRL2^VHL^ complex. Collectively, these results suggest that CRL2^VHL^ is more tolerant to lysine positioning and therefore more resistant to potential resistance mutations affecting ubiquitination. Conversely, CRL4^CRBN^ appears to be more restrictive, such that CRBN-recruiting PROTACs may be vulnerable to lysine-based resistance. This has important implications for the development of therapies, particularly in disease contexts where accessibility to the target lysine is limited or altered *^34^*.

### CeTEAM E3 biosensors demonstrate poor intracellular availability of CRBN-based NUDT5 PROTACs

Initial screens of PROTACs targeting NUDT5 showed that CRBN-recruiting ligands were consistently ineffective degraders compared to VHL-based counterparts (Supplemental Fig. 4C). To investigate whether this discrepancy was due to insufficient binding to the target/poor intracellular accumulation, we employed the recently described cellular target engagement by accumulation of mutant (CeTEAM) assay to the VHL and CRBN E3 ligases (Fig. 5A) ^19,20^. In this setup, ligand binding protects the mutant E3 from rapid degradation, permitting reporter accumulation to be a measure of intracellular binding. Such an approach can highlight if non-functional PROTACs fail due to poor intracellular availability or related to ternary complex formation and ubiquitination. Unlike NanoBRET-based target engagement assays, which require a fluorescent tracer that competitively occupies the site of interest, or CETSA, which reports on ligand-induced thermal stabilization, CeTEAM does not depend on a validated tracer molecule and can, in principle, detect engagement by allosteric or otherwise cryptic binders in physiological conditions, broadening its applicability to E3 ligases and ligands for which no tracer currently exists ^35,36^.We first validated the CeTEAM reporters using a Western blot–based format in which VHL- or CRBN-fused constructs were labeled with V5 and co-expressed with GFP as a normalization control (Fig. 5B–C). After treatment with VHL-binding ligands, including FKBPd3 and VH-298 ^37^, the VHL-CeTEAM reporter showed dose-dependent stabilization with a ∼3.5-fold increase with FKBPd3 at 10 µM and a ∼3.5-fold increase with VH-298 at 50 µM. Similarly, CRBN ligands such as dTAG-13 and iberdomide caused a ∼2-fold stabilization of the CRBN reporter at 10 µM.

**Figure 5.**
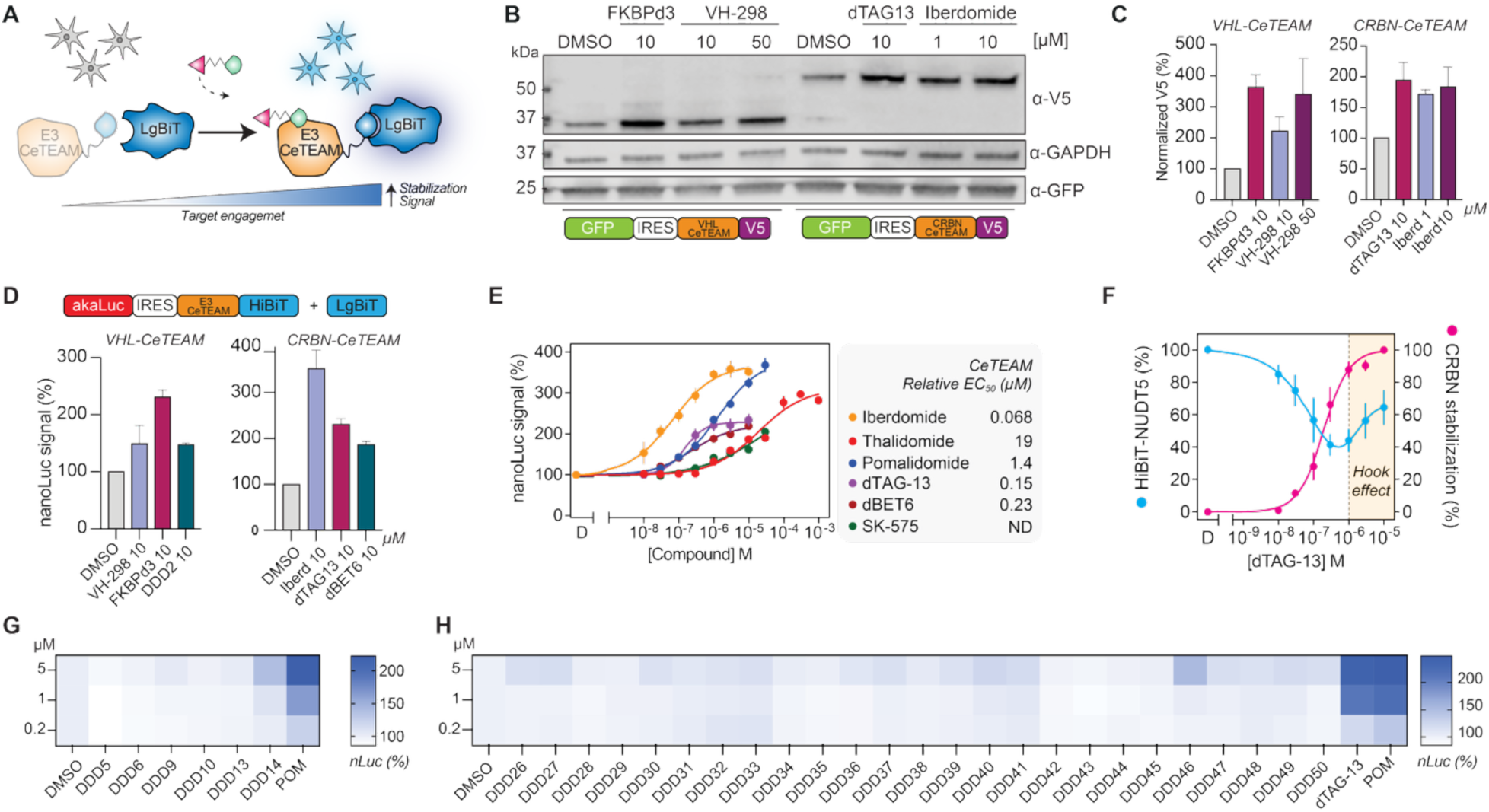
CRBN-CeTEAM assay reveals poor intracellular engagement of CRBN-based NUDT5 degraders. **A.** Schematic of the CeTEAM assay principle. A destabilized E3 ligase (VHL or CRBN) is fused to a reporter protein. Ligand binding protects the E3 fusion from degradation, allowing reporter accumulation as a readout of intracellular engagement. **B.** Western blot–based validation of the E3-CeTEAM biosensors. V5-tagged VHL or CRBN fusion constructs were expressed in U-2 OS cells alongside GFP as a loading control and treated with the indicated ligands and concentrations for 24 hrs. **C.** Quantification of Western blot signals in panel B showing stabilization of VHL and CRBN reporters upon treatment with their respective ligands (n = 2). **D.** Validation of dual-luminescence–based VHL-CeTEAM and CRBN-CeTEAM assays in U-2 OS cells. Data are presented as mean ±SD (n = 3), each performed in triplicate. **E.** Dose–response curves of CRBN-CeTEAM signal in U-2 OS cells expressing HiBiT-CRBN and akaLuc. Cells were treated for 24 h with indicated concentrations of CRBN binders; iberdomide, pomalidomide, thalidomide, dBET6 and SK575 (solubility prevented increased concentrations of SK575 to be included). Data are presented as mean ±SD (n = 3), each performed in triplicate. **F.** Dose response after 24 h treatment with dTAG-13 of degradation reporter signal superimposed on CRBN-CeTEAM stabilization signal, showing saturation and highlighting concentrations where Hook effect is observed (orange). Data are presented as mean ±SD (n = 3 [CRBN CeTEAM], n = 7 [HiBiT-NUDT5-FKBPV_K0_]), each performed in triplicate. Degradation data are fitted using a bell-shaped curve fitting. **G.** Testing of CRBN PROTACs from initial 16 NUDT5 PROTACs and pomalidomide (POM) as control for 24 h at 0.2, 1 or 5 µM using CRBN-CeTEAM expressing U-2 OS cells. Data are presented as mean ±SD (n = 2), each performed in triplicate. **H.** Screening of additional synthesized CRBN-based NUDT5 PROTACs. U-2 OS cells expressing CRBN-CeTEAM were treated with 0.2 µM, 1 µM, and 5 µM of each PROTAC for 24 h. Data are presented as mean ±SD (n = 3), each performed in triplicate. For panels **E-H**, LgBiT expression was induced with doxycycline (0.75 µg/mL) 24 h prior to compound treatment. nLuc signals were normalized to akaLuc and expressed as a percentage relative to the DMSO control.

To improve throughput and sensitivity, we next adapted E3 CeTEAM to a dual-luminescence-based format as before – using HiBiT-labeled CeTEAM E3 mutants and akaLuc as an internal normalization control (Fig. 5A). The luminescence signals correlated well with those observed in the Western blot–based system, confirming the validity and reproducibility of both formats (Fig. 5D). We further benchmarked the CRBN CeTEAM system with known CRBN-binding molecules in dose–response experiments. Molecular glues such as iberdomide, pomalidomide, and thalidomide produced sigmoidal binding curves with EC₅₀ values consistent with reported binding affinities *in vitro* (0.068, 1.4, and 19 µM, respectively; Fig. 5E) ^4,6,38–40^. PROTACs such as dTAG-13, dBET6 (BRD4), and SK575 (PARP1) also stabilized the CRBN CeTEAM biosensor with stability curve trajectories aligning with their respective CRBN ligands – albeit with less robust saturation plateaus (Fig. 5E). Engagement was detectable within 2 hours and increased over time (Supplemental Fig. 6B), highlighting the assay’s ability to capture both early kinetics and steady-state binding. Furthermore, when combined with degradation reporter data for dTAG-13, saturation of the CRBN CeTEAM reporter, a proxy of true CRBN occupancy, coincides with the observed “hook effect” (Fig. 5F), suggesting that the combined insights could be useful to discern the balance of ternary and binary complexes at scale.

After establishing the CRBN CeTEAM dual luciferase system, we then tested all the synthesized CRBN-based PROTACs, including from the initial set of 16 (Fig. 5G) and follow-up set (Fig. 5H) at 0.2 µM, 1 µM, and 5 µM for 24 hours. Despite detecting CRBN CeTEAM stabilization within two hours, we prolonged the assay time to ensure equilibrium for the NUDT5 PROTAC molecules. As before, pomalidomide and dTAG-13 controls dose-dependently stabilized the CRBN binding reporter. Comparatively, the test PROTACs resulted in minimal signal accumulation, indicating poor E3 binding. Conceivably, the poor binding data represent decreased cell permeability or high efflux rate of these analogs, which naturally would coincide with poor degradation efficiency. These results demonstrate the utility of CeTEAM as a sensitive platform to assess both the binding to targeted E3 ligases and the availability of degraders in living cells.

Taken together, our results demonstrate that a dual-luminescent reporter degradation system combined with E3 CeTEAM assays provides a sensitive, modular, and high-throughput platform for functionally assessing key mechanistic steps of PROTAC efficacy, including intracellular availability, E3 binding efficiency, and target degradation (Fig. 6). Ternary complex formation and ubiquitination capabilities can also be inferred from these readouts to facilitate the identification of failure points. Using these tools, we identified DDD2 as a selective and potent VHL-mediated degrader of NUDT5 and explored the E3 binding potency of inactive CRBN-based PROTACs, while mapping key lysine residues on NUDT5 utilized by the CRL2^VHL^ and CRL4^CRBN^ complexes. These findings underscore the importance of prioritizing integrated, cell-based assays in capturing degrader performance beyond what biochemical binding data alone can predict.

**Figure 6.**
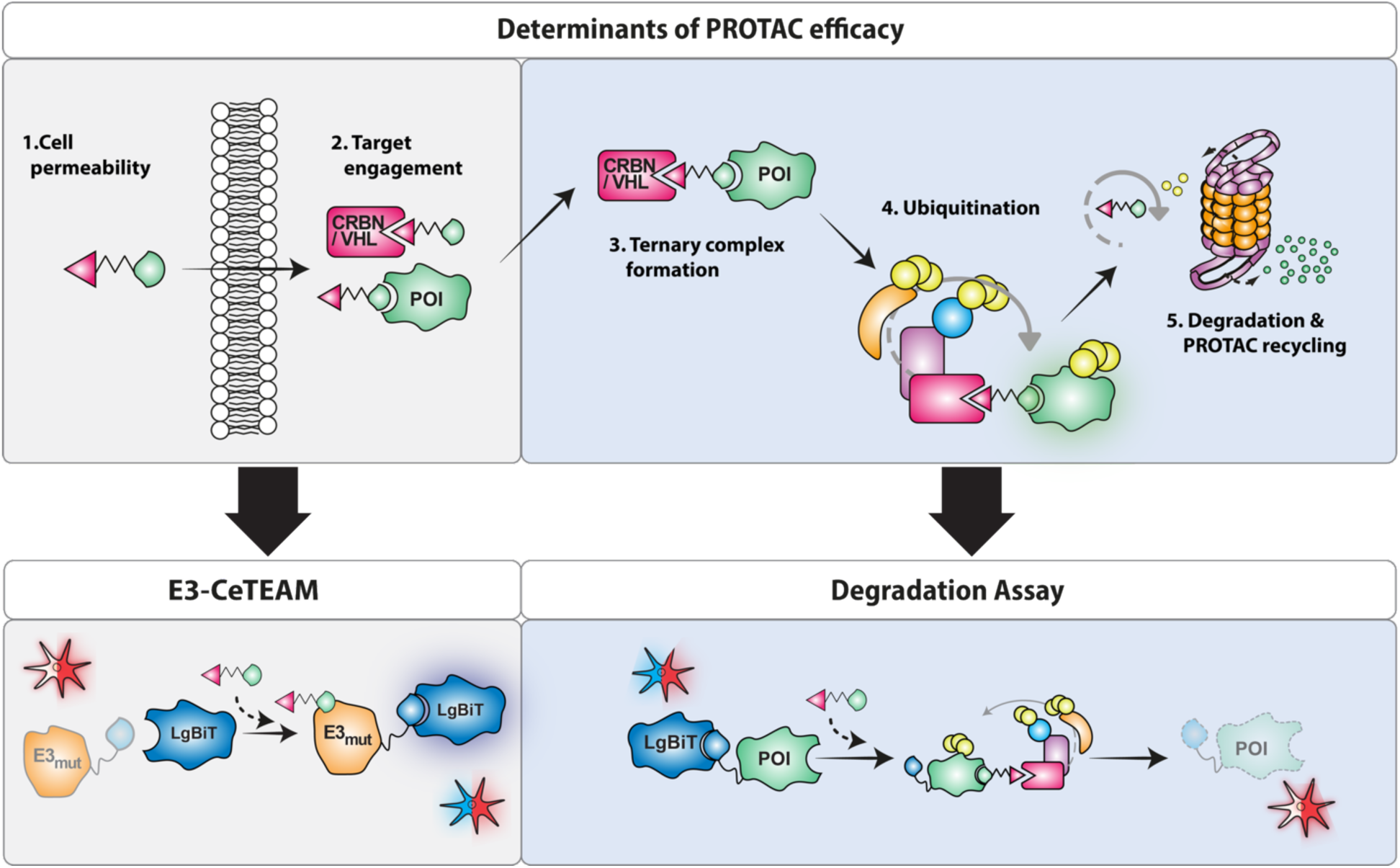
Illustration of key cellular determinants of PROTAC efficacy and the complementary assays used to dissect them mechanistically. The schematic outlines five critical steps: (1) cellular uptake, (2) target engagement, (3) ternary complex formation, (4) ubiquitination, and (5) target degradation with PROTAC recycling. These steps are addressed using our modular dual-luminescent reporter system, which comprises the E3-CeTEAM and degradation assays, providing a sensitive platform to identify mechanistic bottlenecks and guide degrader optimization in cells.

## DISCUSSION

Targeted protein degradation has emerged as a transformative modality in drug discovery, enabling pharmacologic modulation of proteins previously considered “undruggable” by conventional small-molecule inhibitors ^1–3^. By leveraging the ubiquitin-proteasome system, bifunctional degraders such as PROTACs and monovalent molecular glues co-opt cellular E3 ligases to catalyze the ubiquitination and degradation of selected proteins. These strategies offer several theoretical advantages, including sub-stoichiometric dosing, event-driven pharmacology, and potential for targeting non-enzymatic functions of scaffolding or regulatory proteins ^9,41–43^. However, the molecular and cellular determinants that govern efficient degradation are incompletely understood, and successful translation remains hampered by a lack of tools that can resolve degrader activity in live, biologically relevant contexts ^8^.

A key challenge in the field is the mechanistic complexity of degrader function. Unlike classical inhibitors, degraders must engage in a temporally and spatially coordinated cascade of events: cell entry, binary target engagement, effective ternary complex formation, E3 ligase activation, ubiquitin chain elongation, and proteasomal processing ^8^. Each of these steps is sensitive to cellular context, including the expression, localization, and post-translational modification of the target protein, E3 ligase, and relevant UPS machinery ^5,8,44^. Thus, degrader activity observed *in vitro* may not translate to cellular efficacy, and current screening paradigms often conflate ternary complex formation with productive degradation ^45^. This has led to growing calls for mechanistically informative, live-cell assays that can resolve individual steps in the degradation pathway ^8^.

To address this need, we have developed a modular, ratiometric luminescence assay platform that captures time-resolved target degradation and E3 ligase engagement directly in live cells. Using NanoLuc- or HiBiT-tagged proteins of interest (POIs) co-expressed with an internal akaLuc control, we achieve robust and scalable quantification of degradation dynamics with high temporal resolution. The inclusion of a lysine-free FKBPV_K0_ tag enables assessment of target protein degradability, tunable degradation handle configurations, and comparison across ligase-recruiting chemotypes. Importantly, we coupled this platform with an E3 ligase CeTEAM sensor system that detects the biophysical stabilization of CRBN and VHL by binding ligands as a sensitive readout of abundance, enabling holistic analysis of degrader action from intracellular binding to UPS-mediated clearance.

Our findings affirm and extend recent literature showing that the architecture of degrader vector systems, including the identity and polarity of tags, and the presence of lysine residues near the recruitment interface, profoundly influences degradation efficiency ^34,46,47^. Our modular plasmid system facilitated rapid testing of different tag configurations; for example, we observed that exchanging HiBiT for NanoLuc or altering tag orientation led to marked differences in degradation kinetics, even when degrader and target protein concentrations remained constant. These effects likely reflect differences in degron presentation and steric hindrance, which modulate ubiquitination efficiency and proteasomal recognition. While these findings underscore the importance of degrader system/reporter architecture another aspect that also influences is the presence of lysines in tags. Previous reports have shown that that tagging can introduce artifacts, particularly when lysines within the tag itself contribute to ubiquitination, thereby distorting the true degradation profile of the POI ^48,49^. This issue was notably demonstrated in the identification of a KRAS degrader that showed effective degradation when KRAS was GFP-tagged, yet failed to induce degradation of endogenous KRAS ^48^. To overcome this potential confounding factor in our degradation assay system with the FKBPV-tag, we engineered a lysine-free FKBPV_K0_. To our surprise the degradation efficiency of FKBPV-targeting PROTACS was not compromised despite possible structural perturbations and the loss of localized lysine residues, hence, we concluded that the FKBPV_K0_ handle provided a clean scaffold for POI-specific ubiquitination. Of note, while apparent stabilization of HiBiT fusions was observed following complementation with LgBiT, HiBiT versions with at least 200-fold worse reported LgBiT binding affinities showed no marked difference in degradation efficiency, suggesting that it did not contribute significantly in this context. Thus HiBiT-NUDT5-FKBPV_K0_ was established as the optimal degradation reporter. We then applied this reporter across diverse POIs, spanning different sizes, subcellular localizations, and functions. The degradation efficiency varied depending on the POI and the tagged orientation, highlighting the need for protein-specific optimization. In practice, we recommend testing both N- and C-terminally tagged FKBPV_K0_ fusions for each new POI, as no single configuration was uniformly optimal across the panel tested here (Supplemental Fig. 2B). Overall, our modular degradation system with FKBPV_K0_ demonstrated broad applicability and offers a more accurate and generalizable platform for evaluating targeted degradation of a given target protein.

Beyond profiling POI degradation, we extended this platform to enable POI-independent characterization of E3 ligase-recruiting compounds using a HaloTag7/HaloPROTAC configuration. This module allows an E3 ligand or exit vector to be functionally validated for degradation-competent engagement before it is incorporated into a full bifunctional degrader, and can reveal binding-modality differences, such as the absence of a Hook effect with covalent HaloPROTACs, that are diagnostic of the underlying recruitment chemistry. We anticipate this POI-independent module will be particularly useful for triaging new E3 ligands and exit vectors early in degrader discovery campaigns, before committing synthetic effort to a specific POI-targeting design.

We further uncovered E3 ligase–specific differences in lysine preferences for ubiquitination and susceptibility to resistance mutations using lysine availability scanning on the target protein. We could reveal that CRBN-based degraders were highly dependent on the presence of specific lysine residues proximal to the degrader binding site, whereas VHL-based compounds tolerated much broader lysine restrictions. These findings echo structural studies showing that CRBN complexes impose tighter spatial constraints on substrate lysines due to the limited flexibility of the DDB1-CRBN interface ^4,50–52^, whereas the CRL2^VHL^ complex allows for more adaptable ubiquitin transfer geometries ^5^, suggesting CRBN-recruiting degraders may be more vulnerable to therapeutic resistance via loss of key lysines. Thus, lysine availability and positioning should be carefully considered early in CRBN-based PROTAC development to mitigate potential resistance mechanisms and ensure long-term efficacy, an approach supported by the mechanistic resolution offered by our degradation reporter platform.

We applied our optimized NUDT5 degradation reporter to a series of novel PROTACs targeting NUDT5, a nucleotide hydrolase implicated in cellular metabolism and cancer cell proliferation ^23,24^. VHL-based degraders, particularly DDD2, induced robust and sustained degradation, aligning with prior observations that VHL-based PROTACs often demonstrate superior cellular potency ^5,43,53^. Proteomic profiling further confirmed DDD2 as a potent and selective tool for studying NUDT5 biology, with NUDT5 the only protein significantly altered among the >9,800 proteins confidently quantified in our unbiased proteomic profiling, and it represents a promising starting point for therapeutic degrader development. Importantly, our findings highlight a disconnect between traditional biochemical assays used for assessing small molecule interactions with the target and cellular degradation outcomes. Despite relatively weak stabilization of NUDT5 by differential scanning fluorimetry (DSF) and significantly reduced inhibition in enzymatic assays, DDD2 consistently mediated efficient degradation of NUDT5 in live-cell assays. This underscores that binary interactions are not predictive of PROTAC efficacy but, rather, the ability to form productive ternary complexes, enable efficient ubiquitination, and engage the proteasome. Nonetheless, the relative contributions of these factors to degrader efficacy appears to be highly context-dependent ^5,8,10,11,44,54^, including factors affecting compound permeability, protein conformation, and subcellular localization, which further reinforces the value of integrating direct-to-cell assays early in PROTAC development.

Our structure–activity relationship analysis further showed that NUDT5 degradation was highly sensitive to linker composition and length, with piperidine-for-piperazine substitutions and triazole-containing linkers both markedly reducing degradation efficiency despite comparable target engagement. Because piperazine-based exit vectors on VH032, such as that used in DDD2, have been reported to lose VHL-binding potency upon alkylation, exploring alternative exit vectors represents a promising avenue for further potency optimization in future work. Beyond confirming DDD2 as a selective NUDT5 degrader, our long-term treatment and washout experiments provide functional evidence linking NUDT5 abundance to sensitivity to 6-thioguanine (6-TG), a purine analog used clinically as a chemotherapeutic and immunosuppressive agent. Sustained NUDT5 depletion with DDD2 over 22 days conferred resistance to 6-TG toxicity, whereas washout of the compound restored NUDT5 protein levels and re-sensitized cells to 6-TG (Fig. 3E, 3F). This finding is consistent with a recent report describing an unexpected nonenzymatic role for NUDT5 in 6-TG–mediated toxicity, identified using an independent PROTAC-based degradation approach^32^. Together, these observations suggest that NUDT5 contributes to thiopurine drug action through mechanisms that may extend beyond its canonical nucleotide hydrolase activity, and that this contribution can be probed pharmacologically rather than only through genetic depletion. Notably, the reversibility of DDD2-induced degradation, evidenced by NUDT5 rebound following compound washout, allowed us to establish a temporal causal relationship between target protein levels and 6-TG sensitivity within the same cell population, an experimental design not readily achievable with constitutive knockout or knockdown models. This illustrates a broader advantage of degrader-based tools: their pharmacological reversibility enables dynamic interrogation of target function and supports their use in dissecting the biological consequences of protein loss over time.

In contrast to the VHL-recruiting PROTACS towards NUDT5, all tested compounds recruiting CRBN failed to elicit degradation. To interrogate this further, we employed a ligand-responsive CeTEAM reporter to assess CRBN binding directly in cells. As noted in previous reports on CeTEAM, benchmarked CRBN molecular glues and PROTACs had apparent stabilization EC_50_ values that mirrored binding affinity measurements with purified CRBN ^38,40^ further suggesting the approach is a robust biophysical proxy for live cell drug-target engagement. However, none of the CRBN-directed NUDT5 PROTACs synthesized showed appreciable CRBN engagement even at high concentrations, indicating that, whether by permeability or efflux, there is low intracellular availability. However, a CRBN-co-opting PROTAC towards NUDT5 using the TH5427 warhead was recently reported with high degradation efficacy *in cellulo* ^32^, which suggests that further linker diversification can overcome limitations present in our analogs. Interestingly, comparison of E3 CeTEAM binding and target degradation by dTAG-13 revealed that CRBN reporter binding saturation coincided with loss of degradation efficacy, i.e. the hook effect, suggesting it may be a suitable proxy readout of PROTAC intracellular binary and ternary complex dynamics. Further work is needed to evaluate the generality of this approach in understanding PROTAC intracellular ternary complex dynamics at scale.

This study has several limitations. First, while CeTEAM indicated poor intracellular availability of CRBN-directed NUDT5 PROTACs, we could not directly distinguish whether this reflects impaired cellular permeability or active efflux, and resolving this would require dedicated permeability or efflux-transporter assays. Second, our structure-activity exploration around the VH032 exit vector was not exhaustive, and alternative linker chemistries may yet yield more potent VHL-based degraders. Finally, all data presented here were generated in cultured cell lines; extending these findings to *in vivo* pharmacokinetic and pharmacodynamic contexts will be necessary to fully evaluate the therapeutic potential of NUDT5 degraders.

In sum, this work introduces a mechanistically resolved, scalable live-cell assay platform for evaluating targeted protein degraders. By combining sensitive quantification of substrate degradation with conformational biosensors for E3 ligase ligand engagement, the system enables dissection of degrader capability at multiple levels of the PROTAC mechanism-of-action cascade. Importantly, all measurements are performed in biologically relevant cell-based models rather than reconstituted in vitro systems, preserving the influence of endogenous regulation, subcellular trafficking, and proteostasis machineries. The platform is adaptable across diverse POIs, E3 ligases, and degrader modalities, including both bifunctional PROTACs and molecular glue degraders, and can resolve subtle differences in cellular outcomes between closely related compounds. As both PROTACs and molecular glues continue to advance as clinically relevant modalities with applications across oncology, neurodegeneration, and immunology, mechanistically informative tools such as this are critical. By deconstructing degrader function into modular, quantifiable steps in live cells, this framework supports rational design, SAR optimization, and deeper insight into degrader biology, thereby accelerating translation from molecular mechanism to therapeutic impact.

## MATERIAL AND METHODS

### Cell lines and culturing conditions

U-2 OS and HEK293T cells were cultured in DMEM high glucose GlutaMAX media supplemented with 1% Penicillin-Streptomycin and 10% fetal bovine serum (FBS). CCRF-CEM were cultured in RPMI media supplemented with 1% Penicillin-Streptomycin and 10% FBS. For *in vitro* luminescence and fluorescence read-outs, phenol-free DMEM, with high glucose media was used supplemented with 1x GlutaMAX, 1% Penicillin-Streptomycin and 10% FBS. All culture media and supplements were purchased from Thermo Fisher Scientific. All cells were obtained from the ATCC. Cell cultures were maintained at 37°C with 5% CO_2_ in a humidified incubator.

### Antibodies and chemical reagents

anti-GAPDH (polyclonal rabbit, cat. No. ab9485) and anti-NUDT5 (monoclonal rabbit, AB129163) were purchased from Abcam, anti-V5 (monoclonal mouse, cat. No. R960-25) was obtained from Thermo Fisher. anti-GFP (monoclonal mouse, cat. No. sc-9996) and anti-PARP1 (mouse monoclonal, cat. No. sc-8007) were purchased from Santa Cruz. anti-β-actin (monoclonal mouse, A5441) was purchased from Sigma, anti-HaloTag (mouse monoclonal, 28A8) was obtained from ChromoTek and anti-VHL (polyclonal rabbit, 68547) was obtained from Cell Signaling.

AkaLumine HCl was dissolved in MilliQ water (40 mM), aliquoted, and stored at -80°C. Furimazine was reconstituted to 1.25 mM in 15% glycerol, 85% ethanol, and 255 mM ATT (6-aza-2-thiothymine), and stored at -20°C. Doxycycline hydrochloride was dissolved in MilliQ water (2 mg/mL) and used at 0.75μg/mL. dTAG-13, FKBPd3, MLN4294, MG132, dBET6, SK575 and iberdomide were diluted in DMSO to 10 mM. Pomalidomide, thalidomide, VH-298, SK-575 were diluted in DMSO to achieve a stock concentration of 50 mM. All reagents were purchased from MedChemExpress except Doxycycline hydrochloride (DOX) and ATT obtained from Sigma-Aldrich, HaloPROTAC3 from AOBIOUS, Furimazine from Medchemtronica and AkaLumine-HCl (TokeOni) from Tocris.

### Cloning

#### General Subcloning Procedures

Standard molecular biology techniques were used for subcloning. Restriction digests were performed with FastDigest enzymes and FastAP phosphatase, T5 DNA ligase was used for ligations (Thermo Fisher). PCR for subcloning was performed with Phusion High-Fidelity Master Mix and validation colony PCRs was done with DreamTaq Master Mix (Thermo Fisher). Subcloning was done into ENTR vectors using the Gateway recombination cloning system, and propagated in DH5α bacteria with subsequent validation by Sanger sequencing. Constructs were subsequently transferred to the expression vectors using LR Clonase II in Stbl3 bacteria and positive clones were confirmed by colony PCR.

A comprehensive list of gene fragments and vectors generated is presented in Supplemental Table 4. Briefly, a bicistronic vector (pENTR2x) was constructed by subcloning a customized multiple cloning site (MCS_p2x) into pENTR1A (Addgene plasmid # 17398), detailed description can be found in Extended Methods section in Supplemental Information) and was then used for creation of the degradation reporters and CeTEAM E3 biosensors. All gene fragments encoding POI and tags were synthesized or PCR-amplified with flanking restriction sites matching positions desired in the pENTR2x vector, as illustrated in Supplemental Figure 7 and transferred to pLenti-CMV-Blast (Addgene plasmid # 17451). LgBiT, mNG2_1-10 and GFP_1-10 were subcloned as gene fragments into pENTR1A with flanking BamHI/EcoRI restriction sites prior to transfer to pCW57.1 (Addgene plasmid # 41393). To validate CRL2^VHL^ recruitment by DDD2, VHL was silenced through expression of VHL shRNA, generated by oligo-annealing of shRNA primers into an inducible shRNA plasmid ^55^.

#### Plasmids, Primers, and Synthetic DNA

Primers were obtained from Eurofins Genomics, and custom vector fragments were sourced from IDT. Plasmids not generated in the lab were acquired from Addgene or previously published (inducible shRNA ^55^ or gene ^56^). Sequences and sources are listed in Supplemental Table 4.

### Site directed mutagenesis (SDM)

A series of NUDT5 single-lysine variants were generated by reintroducing single lysine residues into the lysine-free NUDT5 backbone (see Supplemental Table 4) using site-directed mutagenesis. Each construct contained a single lysine at one of the 15 native positions in NUDT5. In addition, VHL-R167Q and CRBN-L190F were generated to create CeTEAM E3 biosensors (Supplemental Table 4). Mutagenesis was performed according to the protocol described by Zheng et al. ^57^, and successful incorporation of the mutations was confirmed by DNA sequencing. The primer sequences used for mutagenesis are listed in Table 1 in the supplement. All site-directed mutagenesis was performed on the representative ENTR plasmid and the final constructs for cellular work were generated by LR cloning into Ef1a-Tta3G-P2A-Blast ^56^ for NUDT5 single-lysine variants and pLenti-CMV-Blast (Addgene plasmid # 17451) for CeTEAM-E3 biosensors (Supplemental Table 4).

### Lentivirus Production and Transduction

For stable transduction of expression constructs in cells, lentiviral particles were produced by transfecting subconfluent HEK293T cells with third-generation packaging plasmids (Gag-Pol, Rev, and VSV-G envelope (Addgene plasmids # 12251, 12253, 12259) alongside either destination expression constructs or inducible shRNA construct, using the calcium phosphate precipitation method. Viral supernatants were collected at 48- and 72-hours post-transfection. For transduction, target cells were incubated with a 1:1 mixture of lentiviral supernatant and fresh complete growth medium supplemented with 8 μg/mL polybrene. Forty-eight hours after transduction, cells were re-seeded at low density and subjected to antibiotic selection with either blasticidin (5 μg/mL; Sigma-Aldrich) for four days or puromycin (1 μg/mL; Sigma-Aldrich) for three days, with media replenished every three days.

### In Vitro Luciferase Assays

Degradation efficiency of PROTACs and target engagement of E3 and ligands were measured with in vitro luciferase assay. 4,000 cells per well were seeded in white 96-well plates (Greiner) containing complete medium. Cells were either left untreated or exposed to 0.75 μg/mL doxycycline (DOX). The following day, cells were treated with PROTACs or ligands in different doses and for either 2, 6, or 24 hours, as specified in the figure legends. All compounds were prepared in DMSO and added to the cells at a 1:1000 dilution from the stock solutions in complete media. Sequential luminescence read-outs were performed as described by Valerie et al ^19^. In brief, luciferase substrates were diluted in phenol-red-free DMEM. First readout was performed using 200 μM AkaLumine-HCl, followed by a PBS wash. The second readout was then done with the addition of 200 µM furimazine. Luminescence signals were measured immediately after each substrate addition using the spiral averaging feature. An open filter setting for was used for akaLuc detection, and NanoLuc specifc filter (470± 40 nm) was used to measure NanoLuc signal. Measurements were carried out using CLARIOStar microplate reader (BMG Labtech). To show the percentage of WT degradation and compare the different PROTACs, with WT defined as 100% and K0 as 0% in all lysine mutant assays, the following formula was used: % = ((K0 − X) / (K0 − WT)) × 100.

### Western Blotting

To assess POI abundance after addition of degraders and ligands, Western Blot was performed. A total of 180,000 cells were initially seeded in the absence or presence of 0.75 μg/mL DOX and subsequently treated with PROTACs, naked ligands, or inhibitors the following day. 24h-post-treatment, cells were harvested and lysed using RIPA buffer and 1x Laemmli buffer. The lysates were then heated at 95°C for 5 minutes. Protein samples were either immediately used for electrophoresis or stored at -20°C for later analysis. Proteins were separated on 4-20% gradient Mini-PROTEAN or Midi-PROTEAN gels (Bio-Rad) and transferred onto 0.2 μm nitrocellulose membranes using the Trans-Blot Turbo Transfer System (Bio-Rad). Membranes were blocked with Intercept (TBS) Blocking Buffer (LI-COR) for 1 hour at room temperature. Primary antibodies were diluted in 1x ROTI®Stock TBS-T (Carl Roth) and incubated with the membranes on a shaker at 4°C overnight. The primary antibodies used included mouse monoclonal anti-GFP (1:500), mouse polyclonal anti-V5 (1:2,000), rabbit anti-NUDT5 (1:1,000), rabbit anti-GAPDH (1:2,500), mouse anti-β-actin (1:5,000), and mouse monoclonal anti-PARP1 (1:500). LI-COR secondary antibodies, diluted in 1x ROTI®Stock TBST (1:10,000), were applied and incubated at room temperature for one hour. Blots were visualized using a LI-COR Odyssey Fc system and analyzed with Image Studio Software (LI-COR).

### 6TG Viability Assay following DDD2 Wash-out

HEK293T cells were treated as described in 96-well back assay plates with clear bottom (BD Falcon or Corning) and viability was measured by resazurin reduction viability assay. DDD2 addition and/or removal was initiated prior to plating at 4000 cells per well and incubation with 6TG for an additional 96 hours before viability readings. Resazurin stocks were diluted in cell culture medium and added to cells after removal of treatment media for a final concentration of 0.01 mg/mL. Following incubation in a humidified incubator for 4-6 hours, resorufin fluorescence was measured on a CLARIOStar microplate reader (BMG LABTECH) with 600±40 nm emission filter. Background (medium with resazurin alone) was subtracted from readings prior to normalization to the max viability plateau or DMSO control to yield percent viability. Data were fit to using the [Agonist] vs. response – variable slope (four parameters) curve fitting function in GraphPad Prism (version 10) to calculate EC_50_ values and viability minima (bottom plateau).

### Proteomics

Proteomic analysis was performed to assess compound-induced changes in protein abundance using tandem mass tag (TMTpro)-based quantitative mass spectrometry as previously described ^58,59^. Briefly, HEK293T cells were treated with either vehicle (DMSO) or compounds for 24 hours, lysed in RIPA buffer with protease inhibitors, and subjected to freeze-thaw cycles followed by sonication. After protein extraction and quantification, samples were reduced, alkylated, acetone-precipitated, and digested with LysC and trypsin. TMT-labeled peptides were fractionated by high-pH reversed-phase chromatography and analyzed by high-resolution LC–MS/MS on an Orbitrap Exploris 480 instrument. Protein identification and quantification were carried out using Proteome Discoverer software, and differential protein abundances were calculated and visualized using volcano plots. Full experimental details are provided in the Extended Methods in Supplemental Information. The mass spectrometry proteomics data have been deposited to the ProteomeXchange Consortium via the PRIDE ^60^ partner repository with the dataset identifier PXD065527.

### Chemical Synthesis

#### NUDT5 warheads and PROTACs

Commercially purchased chemicals were used as obtained without further purification. Solvents as well as bulk acids and bases were obtained from Carlo Erba, anhydrous solvents from Alfa Aesar and deuterated solvents from Cambridge Isotope Laboratories, unless otherwise stated. TFA used as de-bocing agent and in HPLC mobile phases was obtained from Chemtronica. The compounds were named using software from Biovia. In addition, the commercial names or trivial names were used for the commercial starting materials and reagents. Analytical HPLC-MS was performed using an Agilent 1100 series Liquid Chromatograph/Mass Selective Detector (MSD) (Single Quadrupole) equipped with an electrospray interface and a UV diode array detector. Analyses were performed by two methods using either an ACE 3 C8 (3.0 x 50 mm) column with a gradient of acetonitrile in 0.1% aqueous TFA over 3 min and a flow of 1 mL/min, or an Xbridge C18 (3.0 x 50 mm) column with a gradient of acetonitrile in 50 mM ammonium bicarbonate over 3 min and a flow of 1 mL/min. Preparative HPLC was performed on a Gilson system equipped with a UV detector using an XBridge Prep C-18 5 µm OBD, 19 x 50 mm column. Flash column chromatography was performed using a Combi Flash RF+ Lumen system. 1H NMR spectra of major intermediates and most final products were recorded on a Bruker 400 MHz instrument or a Bruker Avance Neo 600 MHz instrument at 25 °C, in some cases along with 13C on the latter. The chemical shifts for the NMR signals were referenced relative to TMS via the residual DMSO signal (δH 2.500 ppm, δC 39.52 ppm). Peaks were labelled as follows: s – singlet, d – doublet, t triplet, p – pentet, m – multiplet. The following abbreviations were used for 2D techniques: COSY correlation spectroscopy, TOCSY – total correlation spectroscopy, HSQC – heteronuclear single-quantum coherence, HMBC – heteronuclear multiple-bond correlation, ROESY – rotating frame nuclear Overhauser enhancement spectroscopy. Processing of spectra as well as numbering of atoms was carried out in MestReNova 14.0.1. HRMS spectra of final products were acquired on a Waters Q-ToF Premier system and the experimental mass calculated from an average of duplicates, assuming M(H)=1.0078 and M(Na)=22.9898. PROTACs used in initial cellular screening were synthesized and used at a minimum purity of >80% by HPLC, judged sufficient given the substoichiometric, catalytic mechanism of PROTAC-mediated degradation; compounds selected for extended characterization, including the lead compound DDD2, were synthesized to higher purity (Supplemental Information). For further details on synthesis, see extended methods section in Supplemental Information.

#### HaloPROTACs

HaloPROTAC E was prepared according to Buckley et al ^29^. HPLC, 1H and 13C NMR were in agreement with that reported. HaloPROTAC 4b was synthesized according to Ody et al ^28^. HPLC and 1H NMR agreed with that reported.

### Differential Scanning Fluorimetry (DSF)

Differential scanning fluorimetry was performed as previously described ^24,61^. In brief, reactions were assembled in 96-well PCR plates (Bio-Rad) with a final volume of 20 µL, consisting of 5 µM purified NUDT5 protein, either 10 or 50 µM of the test compound (2.5% v/v DMSO final), and 5x Sypro Orange dye (Thermo Fisher) diluted in assay buffer (100 mM Tris-Acetate, pH 8.0, 40 mM NaCl, 10 mM MgAc, 0.005% Tween-20). Samples were subjected to a temperature gradient ranging from 21°C to 95°C, increasing at a rate of 1°C per minute, using a CFX96 Real-Time PCR machine (Bio-Rad). Fluorescence was measured every minute on the FRET channel, and melting temperatures were determined using the negative first derivative of the fluorescence curve via CFX Maestro software (Bio-Rad).

### NUDT5 Inhibition Assay

To assess NUDT5 inhibition, an enzyme-coupled malachite green assay was employed ^24,61^. Each reaction contained 6 nM purified NUDT5, 10 U/mL bovine alkaline phosphatase (Sigma-Aldrich), and various concentrations of test compounds in assay buffer (0.1 M Tris-Acetate, pH 8.0, 40 mM NaCl, 10 mM MgAc, 1 mM DTT, 0.005% Tween-20, 5% DMSO, and 10% glycerol). Compounds were dispensed using an Echo 550 acoustic dispenser (Labcyte). After addition of 50 µM ADP-ribose as substrate, reactions proceeded at 22°C for 20 minutes before quenching with malachite green reagent. The release of inorganic phosphate was quantified by absorbance at 620 nm using a Hidex Sense plate reader. Control reactions without enzyme or inhibitor were included to establish background and maximal enzyme activity.

### NUDIX Selectivity Assay

Selectivity of test compounds was evaluated at 10 µM concentration against related NUDIX enzymes including MTH1, NUDT15, NUDT18, and NUDT9. As with the NUDT5 assay, the enzyme-coupled malachite green assay was used ^24,61^. Each enzyme was assayed under optimized conditions: MTH1 (4.75 nM) with 100 µM dGTP, NUDT15 (8 nM) with 100 µM dGTP, NUDT18 (200 nM) with 50 µM 8-oxo-dGTP, and NUDT9 (20 nM) with 50 µM ADP-ribose. Pyrophosphatase (Sigma-Aldrich) was substituted for alkaline phosphatase in assays involving substrates that release pyrophosphate. The absorbance at 620 nm was measured after addition of malachite green reagent, and inhibitory effects were compared across NUDIX enzymes.

### Cellular Thermal Shift Assay (CETSA)

CETSA was used to evaluate target engagement of NUDT5 inhibitors in living cells, following protocols from Molina et al. and Page et al. ^24,36^. HEK293T cells (1 × 10^6^ per condition) were treated with test compounds or DMSO (0.1% v/v final concentration) for 2 hours at 37°C in a humidified incubator with 5% CO₂. Cells were harvested by trypsinization, washed with PBS, and resuspended in ice-cold TBS containing protease inhibitor cocktail (Roche) at a ratio of 60 µL per 10^6^ cells. Aliquots were transferred into PCR tubes and heated for 3 minutes at specific temperatures in a Veriti Thermal Cycler (Applied Biosystems), followed by 3 minutes at room temperature. Cells were lysed by three freeze-thaw cycles using a dry ice/ethanol bath and a 37°C water bath. Lysates were clarified by centrifugation at 20,000 × g for 20 minutes at 4°C, and supernatants were collected for immunoblotting. Proteins were detected using a rabbit anti-NUDT5 antibody ^24^ and mouse anti-SOD1 (Santa Cruz Biotechnology, sc-17767) at 1:1000 dilution in 1:1 Intercept Blocking Buffer (LI-COR)/TBS-T. Membranes were visualized with an Odyssey Fc Imager and analyzed using Image Studio Software (LI-COR). Band intensities for NUDT5 were normalized to SOD1, and the fraction of soluble protein was plotted to derive EC₅₀ values using sigmoidal dose-response fitting in GraphPad Prism.

### Flow Cytometry

For fluorescence HaloTag7 degradation experiments, total of 300,000 cells was seeded for each condition in complete growth medium. The following day, cells were treated with HaloPROTAC-E ^30^ at doses indicated in the Supplemental Fig. 3C for 24 hours. Cells were then harvested by trypsinization, washed twice with 10% FBS/PBS, and resuspended in 5% FBS/PBS for subsequent analysis by flow cytometry using a BD Fortessa (BD Biosciences). Data acquisition was performed using BD FACSDiva software, and final gating of cell populations and analysis was conducted with FlowJo (BD Biosciences, Supplemental Fig. 8).

### Statistical Analysis

All statistical analyses and graph generation were performed using GraphPad Prism version 10. Nonlinear regression was applied for sigmoidal dose-response and saturation curve analyses using the four-parameter logistic model ([agonist] vs. response – variable slope) and, in some case, PROTAC degradation curves as a bell-shaped curve fit. Details of specific statistical tests, sample sizes, and significance values are described within individual figure legends.

## Acknowledgments

We would like to express our gratitude to the SciLifeLab Compound Center, part of the Chemical Biology Consortium Sweden (CBCS), for assistance with compound management and spotting. Chemical Proteomics at the Karolinska Institutet (Chemistry I Division, MBB Department), also Unit of SciLifeLab and node of the Swedish National Infrastructure for Biological Mass Spectrometry (BioMS), provided full support in the experimental design, performance, and data analysis of the proteomic studies.

## Author contributions

Conceptualization (PIA, NCKV, MA), Methodology (SA, MJP, NCKV, TG, A-SJ, FK, JE, MG, SvdE, EvB, OW, MA), Validation (SA, MJP, MA, NCKV), Formal analysis (SA, NCKV MJP, A-SJ, FK, JE, MG, SvdE, EvB, OW, RE), Investigation (SA, MJP, NCKV), Resources (NCKV, MA, PIA, JM), Writing – original draft (SA, MJP), Writing – review & editing (SA, MA, NCKV), Visualization (SA, MJP, NCKV, MA), Supervision (NCKV, MA, PIA, JM, RC), Project administration (NCKV, MA), Funding acquisition (NCKV, MA, PIA).

## Funding and additional information

This research was supported by the Swedish Childhood Cancer Society (TJ2019-0036 – NCKV), Cancer Research KI (Karolinska Institutet) Blue Sky Grant (NCKV), Felix Mindus Contribution to Leukemia Research (2019-01992 – NCKV), Loo and Hans Osterman Foundation (2020-01208 – NCKV), Karolinska Institutet Research Foundation (2020-01685, 2022-01749 – NCKV), Swedish Cancer Society (21 0352 PT – NV), Hållsten Foundation (MA), SciLifeLab Technology Development Project Grant (MA, PIA), Novo Nordisk Pioneer Innovator Grant 1 (NNF22OC0076798 – MA, NV, PIA), and MJP was supported by the European Union’s Horizon 2020 research and innovation programme under the Marie Skłodowska-Curie grant agreement No 859860. The views and opinions expressed are those of the authors only and do not necessarily reflect those of the European Union or the European Commission. Neither the European Union nor the European Commission can be held responsible for them.

## Declaration of generative AI and AI-assisted technologies in the writing process

During the preparation of this work the authors used ChatGTP4 and Instatext to improve language clarity. After using these tools, the authors reviewed and edited the content as needed and takes full responsibility for the content of the publication.

