## Supplemental Information for "A cell-based degrader assessment platform facilitates discovery of functional NUDT5 PROTACs"

### Extended Methods

#### Cloning

*Creation of pENTR2x vector*

To create a versatile vector for Gateway cloning [1], we modified the pENTR1a vector (Addgene plasmid #17398) by incorporating a synthetic multiple cloning site (MCS) and an internal ribosome entry site (IRES), pENTR2x. This vector enabled bicistronic expression of a reference protein, protein of interest (POI) and reporter tags.

*Modification of pENTR1a Backbone:*

The native MCS region of pENTR1a was replaced with a redesigned MCS (MCS_p2x) containing multiple unique restriction sites to expand cloning (Supplemental Fig. 7 A, B and Supplemental Table 2). The new MCS was generated by annealing complementary oligonucleotides flanked with SalI and NotI restriction sites and inserted into pENTR1A by restriction digestion. Recombinant clones were selected in E. coli DH5α. Successful insertion was confirmed by sequencing.

*Insertion of IRES Element:*

An internal ribosome entry site (IRES) sequence was inserted in the modified pENTR1a vector to allow for translation of bicistronic expression from the same transcript, with BamHI/EcoRI sites upstream of IRES for insertion of reference gene and MCS downstream of IRES for N-terminal tag (NcoI/SalI) for POI (Sal//NotI) and C-terminal tag (XhoI/XbaI) (see Supplemental Fig. 7 A, B). The IRES sequence was synthesized with overhangs for EcoRI/NcoI and inserted in the modified pENTR1A backbone using EcoRI/NcoI digestion and ligation generating the final pENTR2x vector used for this study. Correct insertion was validated by sequencing.

#### Proteomics

*Sample processing*

Cells were treated by adding to in their culture media the same volume amounts of DMSO (control vehicle alone for the untreated control) or of each tested compound. For each treatment type, a minimum of triplicate cell cultures was provided. Cells were then collected, washed twice in PBS and resuspended in RIPA buffer with protease inhibitors. Cells were lysed by subjecting them to freeze-thaw cycles in liquid nitrogen for a total of 5 times, followed by probe sonication for 1 min (03 pulse, 03 pause, 10 times -30% amplitude) on ice.

After centrifugation for debris removal at 14,000 at 4 °C for 30 min, the supernatant was collected. The total protein concentration of extracted samples was measured using micro-BCA (bicinchoninic acid) assay and 50 µM of protein was processed by 8 mM dithiothreitol (Sigma) reduction, 25 mM iodoacetamide (Sigma) alkylation. Then samples were precipitated using cold acetone at -20 °C overnight. Briefly, as previously published [2],[3]. 50 µg of each sample was reduced, alkylated and precipitated using cold acetone. Samples were resuspended in EPPS (4-(2-Hydroxyethyl)-1-piperazinepropanesulfonic acid) buffer, 8 M urea, pH 8.0, diluted down to 4 M urea and digested by LysC, then diluted down to 1 M urea and digested with trypsin. Each digest was labeled using TMTpro technology (Thermo Fischer) for deep protein identification and quantification by LC-MS data dependent acquisition. The final multiplex sample was first cleaned by Sep-Pack C18 column (Waters) and then separated into 48 fractions by capillary reversed phase chromatography at pH 10. Each fraction was analyzed by high-resolution nLC–ESI-MS/MS (nanoscale liquid chromatography-electrospray ionization-tandem mass spectrometry) using an Orbitrap Exploris 480 instrument (Thermo Scientific).

*NanoLC-MS/MS analysis*

NanoLC-MS/MS analyses were performed on an Orbitrap Exploris 480 mass spectrometer (Thermo Fisher Scientific). The instrument was equipped with an EASY ElectroSpray source and connected online to an Ultimate 3000 nanoflow UPLC system. The samples were pre-concentrated and desalted online using a PepMap C18 nano-trap column (length - 2 cm; inner diameter - 75 µm; particle size - 3 µm; pore size - 100 Å; Thermo Fisher Scientific) with a flow rate of 3 µL/min for 5 min. Peptide separation was performed on an EASY-Spray C18 reversed-phase nano-LC column (Acclaim PepMap RSLC; length - 50 cm; inner diameter - 2 µm; particle size - 2 µm; pore size – 100 Å; Thermo Scientific) at 55 °C and a flow rate of 300 nL/min. Peptides were separated using a binary solvent system consisting of 0.1% (v/v) FA, 2% (v/v) ACN (solvent A) and 98% ACN (v/v), 0.1% (v/v) FA (solvent B). They were eluted with a gradient of 3–26% B in 97 min, and 26–95% B in 9 min. Subsequently, the analytical column was washed with 95% B for 5 min before re-equilibration with 3% B. The mass spectrometer was operated in a data-dependent acquisition mode. A survey mass spectrum (from m/z 375 to 1500) was acquired in the Orbitrap analyzer at a nominal resolution of 120,000. The automatic gain control (AGC) target was set as 100% standard, with the maximum injection time of 50 ms. The most abundant ions in charge states 2+ to 7+ were isolated in a 3 s cycle, fragmented using HCD MS/MS with 33% normalized collision energy, and detected in the Orbitrap analyzer at a nominal mass resolution of 50,000. The AGC target for MS/MS was set as 250% standard with a maximum injection time of 100 ms, whereas dynamic exclusion was set to 45 s with a 10-ppm mass window.

*Protein Identification, Quantification and Data Analysis*

Proteome Discoverer 2.5 software (Thermo Scientific) was utilized for the database search and quantification against the Uniprot Homo sapiens (Human) protein database UP000005640. Cysteine carbamidomethylation was set as a fixed modification, along with TMT-related modifications, methionine oxidation, deamidation of arginine and asparagine as variable modifications. Enzyme specificity was defined as trypsin with a maximum of two missed cleavages. A 1% false discovery rate was employed as a filter at both the protein and peptide levels. Contaminants and reversed-hit peptides were removed, and only proteins with at least two unique peptides were included in the quantitative analysis. Proteins with missing values were eliminated. The quantified abundance of each protein in each sample (labeled with a different TMT) was normalized to the total intensity of all proteins in that sample. For each protein in each compound replicate, the normalized protein abundance was divided by the average abundance of that protein in the vehicle-treated replicates. The average ratio across replicates of each compound compared to the vehicle control was calculated, and the $Log_2$ values of these ratios were determined. Two-tailed Student's t-test was employed to calculate the p-value, assuming a non-zero ratio. For each compound, the results were visualized using a volcano plot, which plotted the $Log_2$ ratio on the x-axis and the corresponding $-Log_{10}$ p-value on the y-axis.

### Chemical Synthesis

#### *Synthesis of compounds in Figure 2*

Preparation of **DDD1**

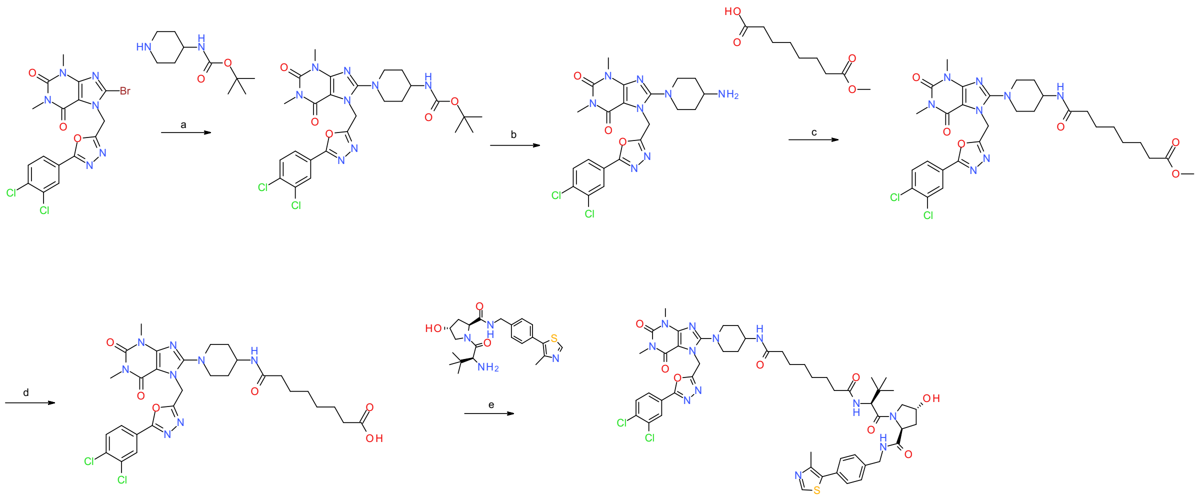

**Scheme S1.** Reagents and conditions: a) $K_2CO_3$, DMF, 80 °C, overnight; b) TFA, DCM, rt, 2.5 h; c) HATU, DIEA, DMF, rt, overnight; d) LiOH·$H_2O$, water:THF (1:1), rt, overnight; e) HATU, DIEA, DMF, rt, overnight.

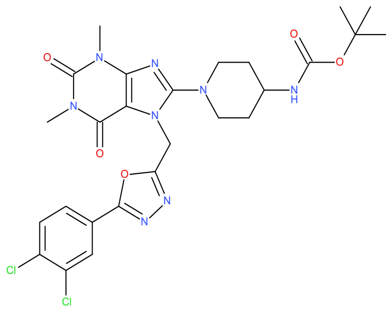

**_tert_-Butyl _N_-[1-[7-[[5-(3,4-dichlorophenyl)-1,3,4-oxadiazol-2-yl]methyl]-1,3-dimethyl-2,6-dioxo-purin-8-yl]-4-piperidyl]carbamate (8a)**

Compound **1a**[4] (0.600 g, 1.11 mmol), *tert*-butyl *N*-(4- piperidyl)carbamate (0.278 g, 1.33 mmol) and $K_2CO_3$ (0.307 g, 2.20 mmol) were dissolved in DMF (6 mL) under a nitrogen atmosphere. The mixture was heated to 80 °C and left overnight. The mixture was added to a larger volume of EtOAc and washed with brine, but a precipitate was formed. The precipitate was then extracted into DCM and the organic phase was dried over $MgSO_4$, filtered and dried under vacuum to afford the title compound (0.563 g, 0.932 mmol, yield: 84%). MS (ESI$^+$) *m/z* 605 [M+H]$^+$ .

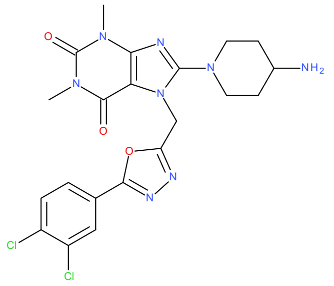

**8-(4-Amino-1-piperidyl)-7-[[5-(3,4-dichlorophenyl)-1,3,4-oxadiazol-2-yl]methyl]-1,3-dimethyl-purine-2,6-dione (9a)**

TFA (1.20 mL, 16.1 mmol) was dissolved in DCM (4.8 mL) to generate a 20% solution. This solution was added – first slowly, then rapidly – to compound **8a** (0.564 g, 0.932 mmol) and the mixture stirred in a sealed vial for 2.5 h. The mixture was neutralized with 2 M potassium carbonate solution and the product extracted to a larger volume of DCM, washed with brine and dried over magnesium sulphate. Removal of the solvent on a rotary evaporator yielded the title compound (0.533 g, 1.02 mmol, yield: >99%), which was used in the next step without further purification. MS (ESI$^+$) *m/z* 505 [M+H]$^+$. $^1$H NMR (400 MHz, DMSO-$d_6$): δ 8.16 (d, J=1.9 Hz, 1H), 7.89-7.94 (m, 2H), 5.67 (s, 2H), 3.56 (d, J=12 Hz, 2H), 3.41 (s, 3H), 3.02 (t, J=12 Hz, 2H), 3.16 (s, 3H), 2.83-2.89 (m, 1H), 1.78-1.86 (m, 2H), 1.37-1.48 (m, 2H).

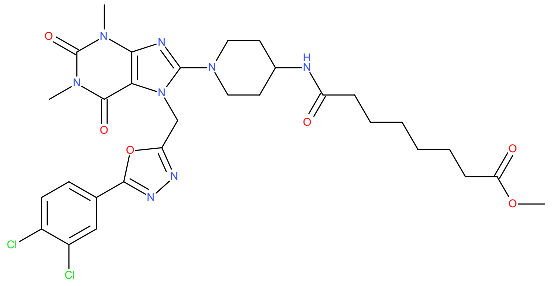

**Methyl 8-[[1-[7-[[5-(3,4-dichlorophenyl)-1,3,4-oxadiazol-2-yl]methyl]-1,3-dimethyl-2,6-dioxo-purin-8-yl]-4-piperidyl]amino]-8-oxo-octanoate (10a)**

Suberic acid monomethyl ester (0.0383 g, 0.200 mmol), DIEA (0.0697 mL, 0.403 mmol) and HATU (0.0929 g, 0.244 mmol) were dissolved in DMF (2 mL) and conditioned for 10 minutes at rt. Compound **9a** (0.100 g, 0.198 mmol) was added and the reaction mixture left overnight. The mixture was diluted with water and organics extracted with DCM (3x). The combined organic extracts were washed with brine, after which the product was dried over magnesium sulphate and the solvent was removed. This yielded the title compound (95.0%, 0.108 g, 0.152 mmol, yield: 77%), which was used in the next step without further purification. MS (ESI$^+$) *m/z* 675 [M+H]$^+$.

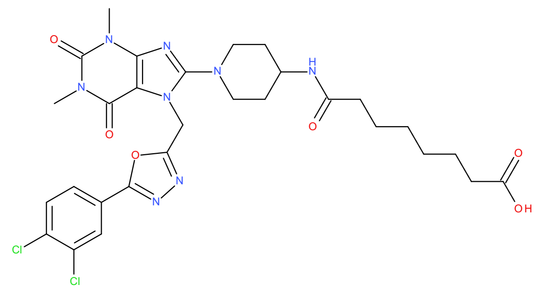

**8-[[1-[7-[[5-(3,4-Dichlorophenyl)-1,3,4-oxadiazol-2-yl]methyl]-1,3-dimethyl-2,6-dioxo-purin-8- yl]-4-piperidyl]amino]-8-oxo-octanoic acid (11a)**

Compound **10a** (0.100 g, 0.141 mmol) and LiOH·$H_2O$ (0.0181 g, 0.422 mmol) were mixed in water:THF (4 mL, 1:1) and stirred in a closed vessel at rt overnight. Monitoring of the reaction indicated incomplete formation of the product, despite a long reaction time. THF was removed in vacuo, and the crude mixture was acidification until pH=2 using 1 M HCl. The aqueous phase was extracted with DCM (4 × 25 mL), drying over $MgSO_4$ and concentrated to afford the title compound (36.0%, 0.0460 g, 0.0250 mmol, yield: 18%), which was used in the next step without further purification. MS (ESI$^+$) *m/z* 660 $[M+H]^+$.

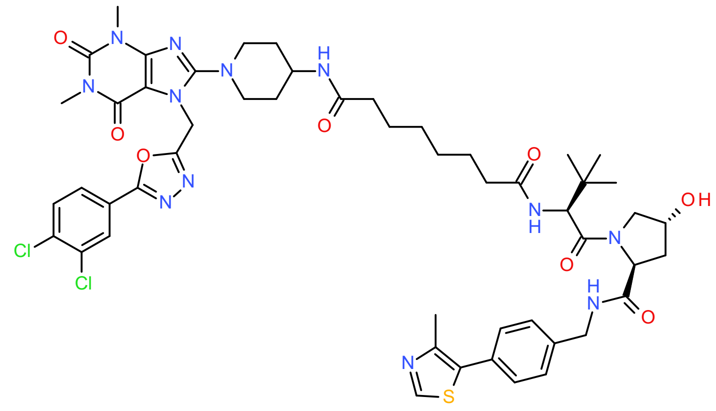

***N*-[1-[7-[[5-(3,4-Dichlorophenyl)-1,3,4-oxadiazol-2-yl]methyl]-1,3-dimethyl-2,6-dioxo-purin-8- yl]-4-piperidyl]-*N'*-[(1S)-1-[(2S,4R)-4-hydroxy-2-[[4-(4-methylthiazol-5-yl)phenyl] methylcarbamoyl]pyrrolidine-1-carbonyl]-2,2-dimethyl-propyl]octanediamide (12a)**

Compound **11a** (0.0208 g, 0.0113 mmol), DIEA (0.0165 mL, 0.0965 mmol) and HATU (0.0147 g, 0.0385 mmol) were dissolved in DMF (1.5 mL) and left to stir for 10 minutes at rt. Compound **2a** (0.0138 g, 0.0321 mmol) was added and the reaction was left overnight. The mixture was diluted with EtOAc and washed with brine, concentrated in vacuo and finally purified by preparatory HPLC (gradient of 5% to 40% acetonitrile in 50 mM ammonium bicarbonate). Lyophilizing the relevant fractions afforded the title compound (>99%, 2.50 mg, 2.33 µmol, yield: 21%). MS (ESI$^+$) *m/z* 538 $[M+2H]^{2+}$.

Preparation of **DDD2** (Large scale synthesis)

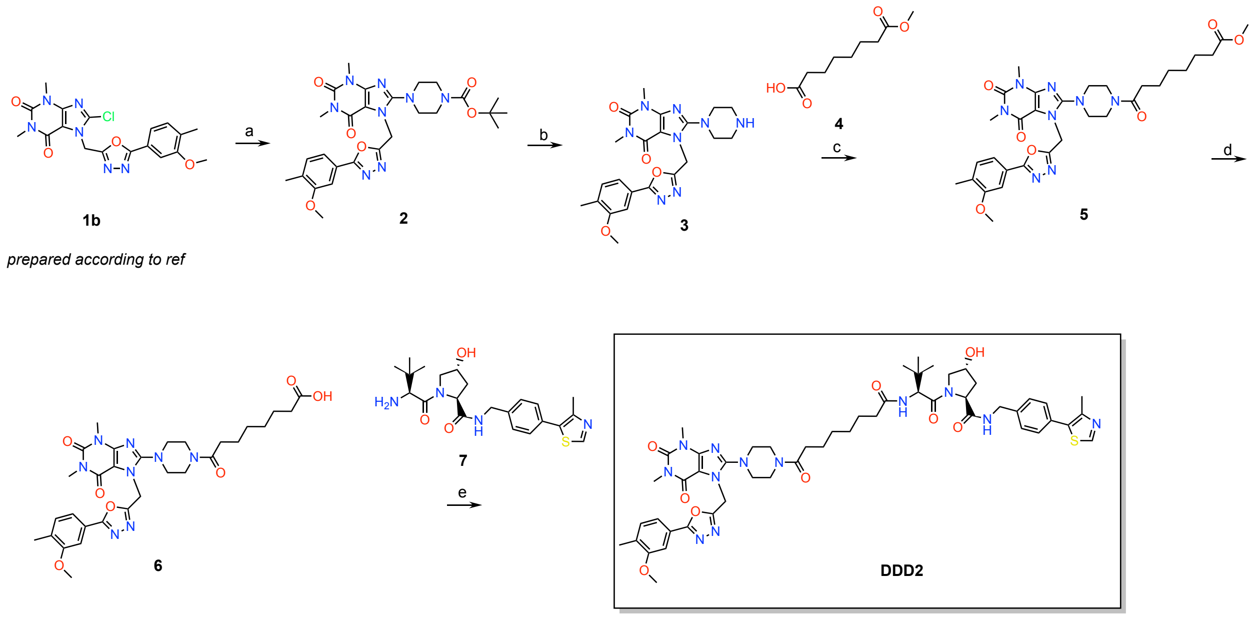

**Scheme S2.** Reagents and conditions: a) *tert*-butyl piperazine-1-carboxylate, $K_2CO_3$, DMF, 70 ℃, overnight.; b) TFA, DCM, rt, 3 h; c) HATU, DIPEA, DMF, rt, overnight; d) LiOH-$H_2O$, THF, $H_2O$, rt, 5 h; e) HATU, DIPEA, DMF, rt, overnight.

#### *tert*-Butyl 4-[7-[[5-(3-methoxy-4-methyl-phenyl)-1,3,4-oxadiazol-2-yl]methyl]-1,3-dimethyl-2,6-dioxo-purin-8-yl]piperazine-1-carboxylate (2)

8-Bromo-7-[[5-(3-methoxy-4-methyl-phenyl)-1,3,4-oxadiazol-2-yl]methyl]-1,3-dimethyl-purine-2,6-dione (**1b**)[4] (5.21 g, 6.88 mmol), *tert*-butyl piperazine-1-carboxylate (1.60 g, 8.61 mmol) and $K_2CO_3$ (1.90 g, 13.8 mmol) were mixed in DMF (50 mL) and heated at 70 °C overnight. LC-MS analysis indicated unreacted ligand was still present, so another 600 mg of BOC-piperazine was added. After another 3 h, an additional 650 mg of $K_2CO_3$ was added. The mixture was left over the weekend. The mixture was diluted with ethyl acetate (100 mL) and water:brine (1:1, 100 mL). The organic phase was separated and washed with brine (2x100 mL), dried over $MgSO_4$ and the solvent evaporated in vacuo. The crude product was purified with flash column chromatography on silica gel, using MeOH in DCM (0-10%) as eluent to afford the title compound (95.0%, 5.55 g, 9.11 mmol, yield: >99%). MS (ESI$^+$) *m/z* 567 $[M+H]^+$.

#### 7-[[5-(3-Methoxy-4-methyl-phenyl)-1,3,4-oxadiazol-2-yl]methyl]-1,3-dimethyl-8-piperazin-1-yl-purine-2,6-dione (3)

*tert*-Butyl 4-[7-[[5-(3-methoxy-4-methyl-phenyl)-1,3,4-oxadiazol-2-yl]methyl]-1,3-dimethyl-2,6-dioxo-purin-8-yl]piperazine-1-carboxylate (**2**) (5.55 g, 9.11 mmol) was treated with TFA (20.0%, 60.0 mL, 0.162 mol) in DCM and stirred for 3 h at rt. The mixture was neutralized with 2 M aqueous potassium carbonate solution and the product extracted to a larger volume of DCM, washed with brine and dried over magnesium sulphate. Removal of the solvent on a rotary evaporator yielded the title compound (93.0%, 4.67 g, 9.32 mmol, yield: >99%). MS ($ESI^+$) *m/z* 467 $[M+H]^+$.

**Methyl 8-[4-[7-[[5-(3-methoxy-4-methyl-phenyl)-1,3,4-oxadiazol-2-yl]methyl]-1,3-dimethyl-2,6-dioxo-purin-8-yl]piperazin-1-yl]-8-oxo-octanoate (5)**

Suberic acid monomethyl ester (**4**) (0.807 g, 4.29 mmol), DIEA (1.50 mL, 8.76 mmol) and HATU (1.96 g, 5.14 mmol) were conditioned in DMF (30 mL) for 30 minutes at rt. 7-[[5-(3-Methoxy-4-methyl-phenyl)-1,3,4-oxadiazol-2-yl]methyl]-1,3-dimethyl-8-piperazin-1-yl-purine-2,6-dione (**3**) (2.15 g, 4.29 mmol) was added and the reaction mixture was left overnight. The mixture was diluted with 1 M HCl (25 mL) and organics extracted with EtOAc (50 mL). The organic phase was then washed with brine (50 mL) and dried over $MgSO_4$. This extraction process was repeated from the DMF/HCl mixture. Pooling of the organic extracts and removal of the solvent on a rotary evaporator yielded the title compound (80.0%, 4.19 g, 5.27 mmol, yield: >99 %) as a viscous liquid - indicating DMF was still present. The material was used in the next step without further purification. MS ($ESI^+$) *m/z* 637 $[M+H]^+$.

**8-[4-[7-[[5-(3-Methoxy-4-methyl-phenyl)-1,3,4-oxadiazol-2-yl]methyl]-1,3-dimethyl-2,6-dioxo-purin-8-yl]piperazin-1-yl]-8-oxo-octanoic acid (6)**

To a water:THF mixture (1:1) (60 mL) were added methyl 8-[4-[7-[[5-(3-methoxy-4-methyl-phenyl)-1,3,4-oxadiazol-2-yl]methyl]-1,3-dimethyl-2,6-dioxo-purin-8-yl]piperazin-1-yl]-8-oxo-octanoate (**5**) (4.19 g, 5.27 mmol) and $LiOH \cdot H_2O$ (0.332 g, 7.90 mmol) and the mixture was stirred at rt for 5 h. THF was removed in vacuo and water (150 mL) and DCM (50 mL) were added. As no separation of phases was observed, volatile solvents were removed in vacuo overnight. The pH was then adjusted to 2 with 1 M HCl and organics extracted with DCM (3x50 mL). Each organic extract was passed over brine (50 mL), after which they were pooled and dried over $MgSO_4$. Removal of the solvent in vacuo yielded the crude product which was purified by flash column chromatography on silica gel, using MeOH in DCM (0-5%) as eluent. Pooling and drying of the relevant fractions yielded the title compound (93.0%, 1.45 g, 2.16 mmol, yield: 41%). MS ($ESI^-$) *m/z* 621 $[M-H]^-$.

***(2S,4R)*-4-Hydroxy-1-[(*2S*)-2-[[8-[4-[7-[[5-(3-methoxy-4-methyl-phenyl)-1,3,4-oxadiazol-2-yl]methyl]-1,3-dimethyl-2,6-dioxo-purin-8-yl]piperazin-1-yl]-8-oxo-octanoyl]amino]-3,3-dimethyl-butanoyl]-*N*-[[4-(4-methylthiazol-5-yl)phenyl] methyl] pyrrolidine-2-carboxamide (DDD02)**

8-[4-[7-[[5-(3-Methoxy-4-methyl-phenyl)-1,3,4-oxadiazol-2-yl]methyl]-1,3-dimethyl-2,6-dioxo-purin-8-yl] piperazin-1-yl]-8-oxo-octanoic acid (**6**) (0.500 g, 0.747 mmol), (*2S*,*4R*)-1-[(*2S*)-2-amino-3,3-dimethyl-butanoyl]-4-hydroxy-*N*-[[4-(4-methylthiazol-5-yl)phenyl]methyl]pyrrolidine-2-carboxamide;hydrochloride (**7**) (0.349 g, 0.747 mmol) and DIEA (0.384 mL, 2.24 mmol) were dissolved in DMF (9 mL) and conditioned at rt for 10 min. HATU (0.341 g, 0.896 mmol) was added and the solution stirred overnight. The mixture was diluted with saturated $NaHCO_3$ (40 mL) and organics extracted with EtOAc (50 mL). The organic phase was washed with brine (2x50 mL) and dried over $MgSO_4$. After the solvent was removed in vacuo, the product was purified by preparatory HPLC (gradient of 20% to 30% acetonitrile in 50 mM ammonium bicarbonate). Lyophilization of the pure fractions afforded the title compound (97.0%, 86.2 mg, 81.0 µmol, yield: 11%). MS ($ESI^+$) *m/z* 518 $[M+2H]^{2+}$. HRMS *m/z* calculated 1035.4874, found 1035.4890 for $[M+H]^+$. $^1H$ NMR (400 MHz, DMSO-$d_6$) δ 8.97 (s, 1H), 8.55 (t, *J* = 6.1 Hz, 1H), 7.84 (d, *J* = 9.3 Hz, 1H), 7.48 – 7.33 (m, 7H), 5.71 (s, 2H), 5.11 (d, *J* = 3.5 Hz, 1H), 4.54 (d, *J* = 9.4 Hz, 1H), 4.47 – 4.39 (m, 2H), 4.37 – 4.31 (m, 1H), 4.21 (dd, *J* = 15.9, 5.4 Hz, 1H), 3.88 (s, 3H), 3.68 – 3.62 (m, 2H), 3.57 (d, *J* = 6.8 Hz, 4H), 3.40 (s, 3H), 3.29 – 3.19 (m, 4H), 3.15 (s, 3H), 2.44 (s, 3H), 2.35 – 2.24 (m, 3H), 2.23 (s, 3H), 2.16 – 1.98 (m, 2H), 1.95 – 1.85 (m, 1H), 1.57 – 1.37 (m, 4H), 1.33 – 1.17 (m, 4H), 0.93 (s, 9H). $^{13}C$ NMR (101 MHz, DMSO-$d_6$) δ 172.1, 171.9, 170.9, 169.7, 164.4, 162.4, 157.7, 155.9, 153.8, 151.4, 150.9, 147.7, 147.2, 139.5, 131.4, 131.2, 130.7, 129.6, 128.6, 127.4, 121.8, 118.7, 107.5, 104.1, 68.9, 58.7, 56.4, 56.3, 55.5, 50.0, 49.7, 44.3, 41.6, 40.4, 38.0, 35.2, 34.8, 32.2, 29.5, 28.5, 27.4, 26.4, 25.3, 24.6, 16.2, 15.9. Two C overlapping somewhere at lower shift than DMSO.

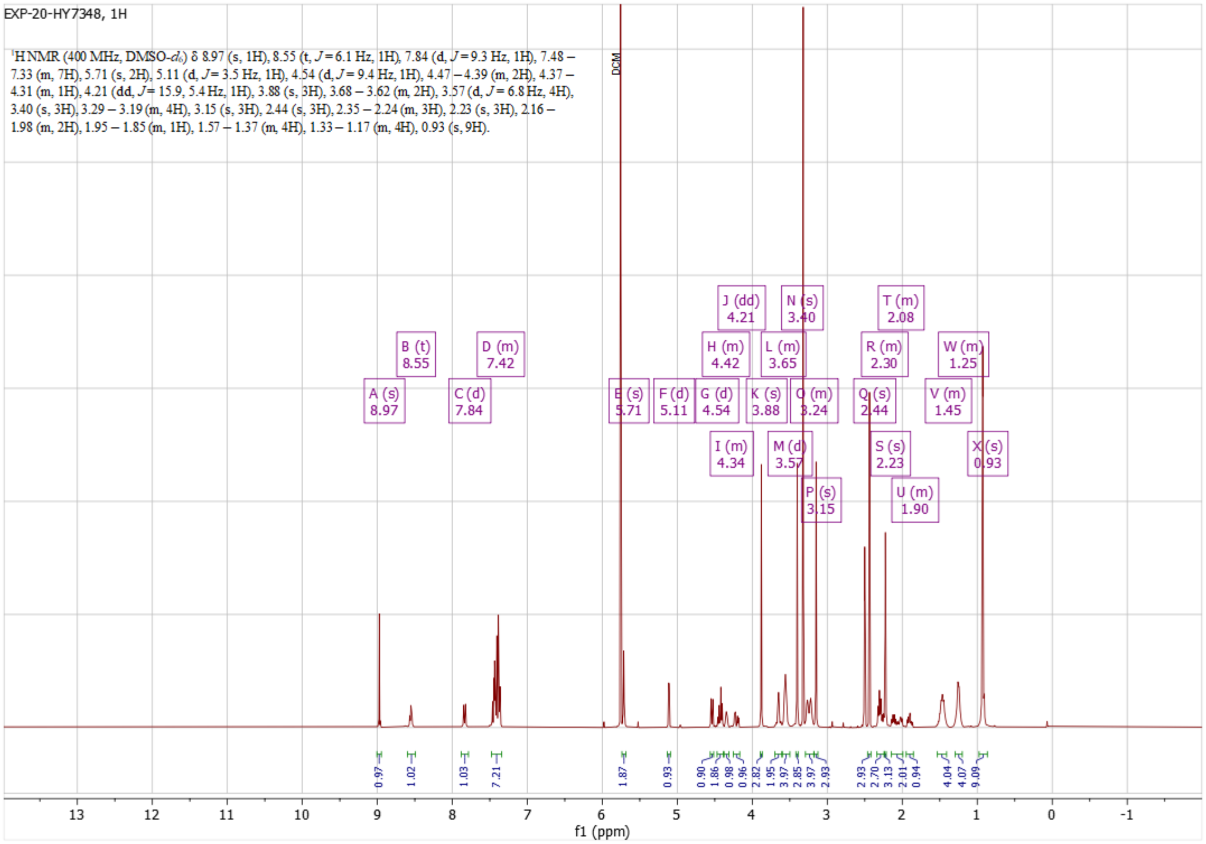

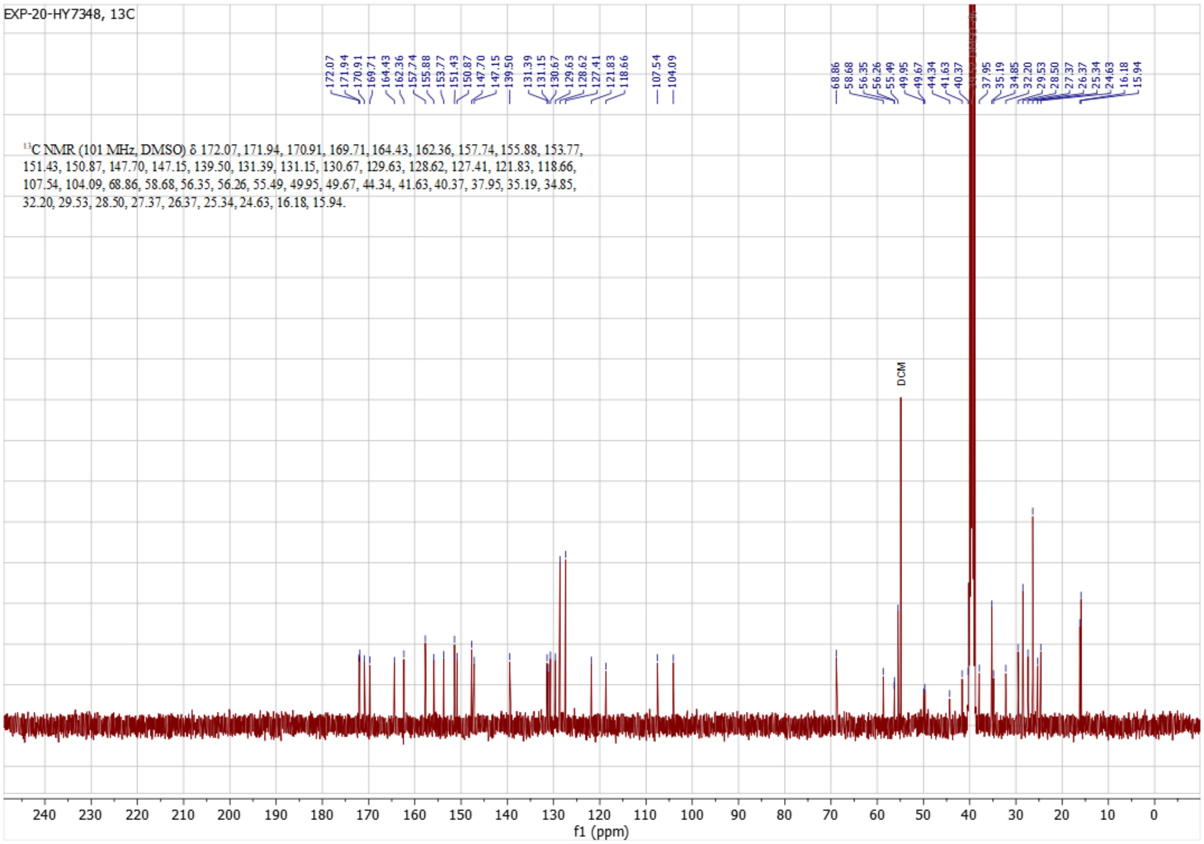
EXP-20-HY7348, 13C
$^{13}$C NMR (101 MHz, DMSO) δ 172.07, 171.94, 170.91, 169.71, 164.43, 162.36, 157.74, 155.88, 153.77, 151.43, 150.87, 147.70, 147.15, 139.50, 131.39, 131.15, 130.67, 129.63, 128.62, 127.41, 121.83, 118.66, 107.54, 104.09, 68.86, 58.68, 56.35, 56.26, 55.49, 49.95, 49.67, 44.34, 41.63, 40.37, 37.95, 35.19, 34.85, 32.20, 29.53, 28.50, 27.37, 26.37, 25.34, 24.63, 16.18, 15.94.
DCM
f1 (ppm)

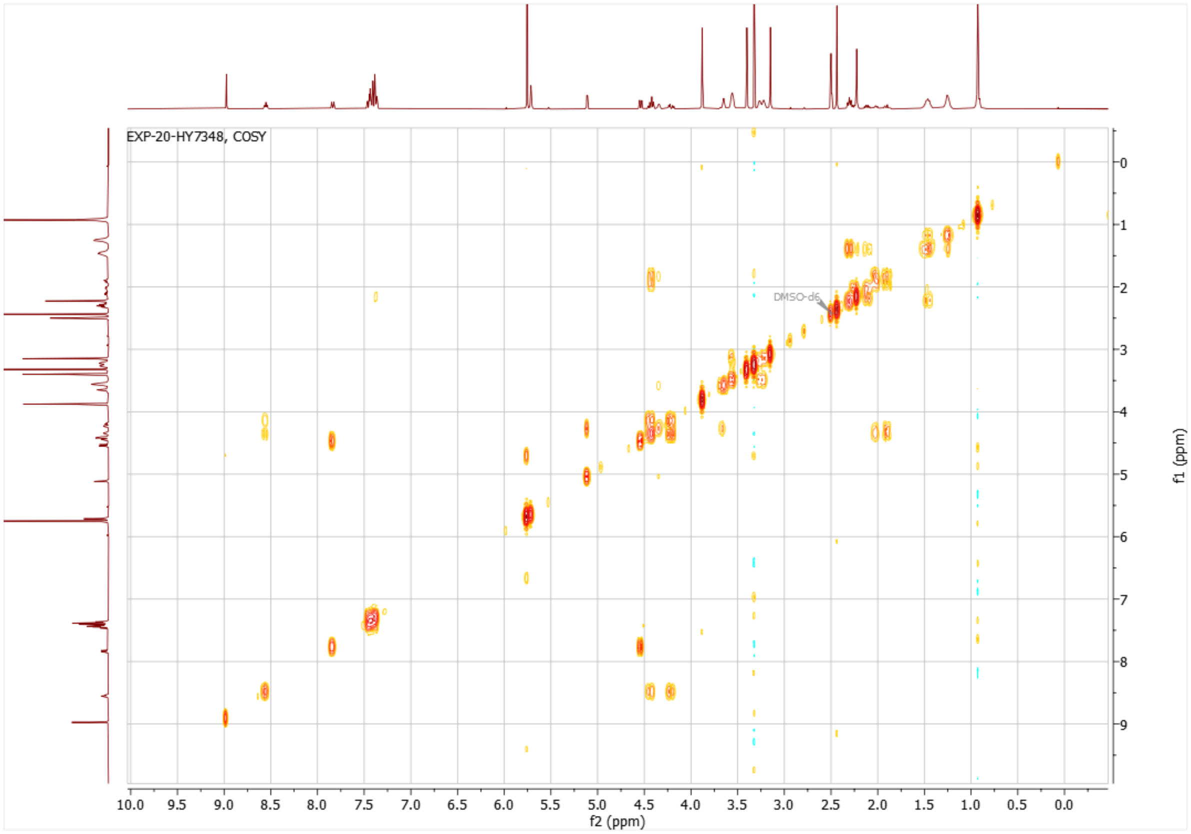
EXP-20-HY7348, COSY
DMSO-d6
f2 (ppm)
f1 (ppm)

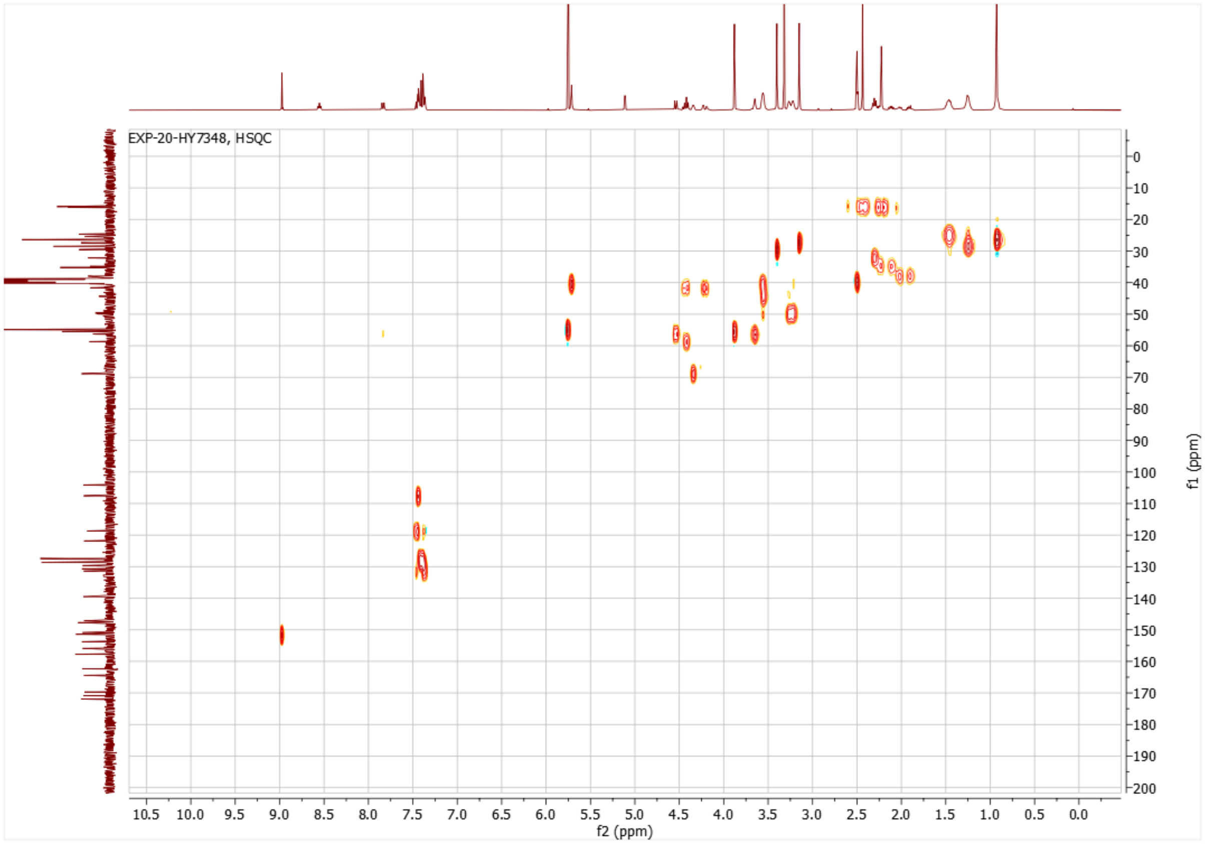
EXP-20-HY7348, HSQC
f2 (ppm)
f1 (ppm)

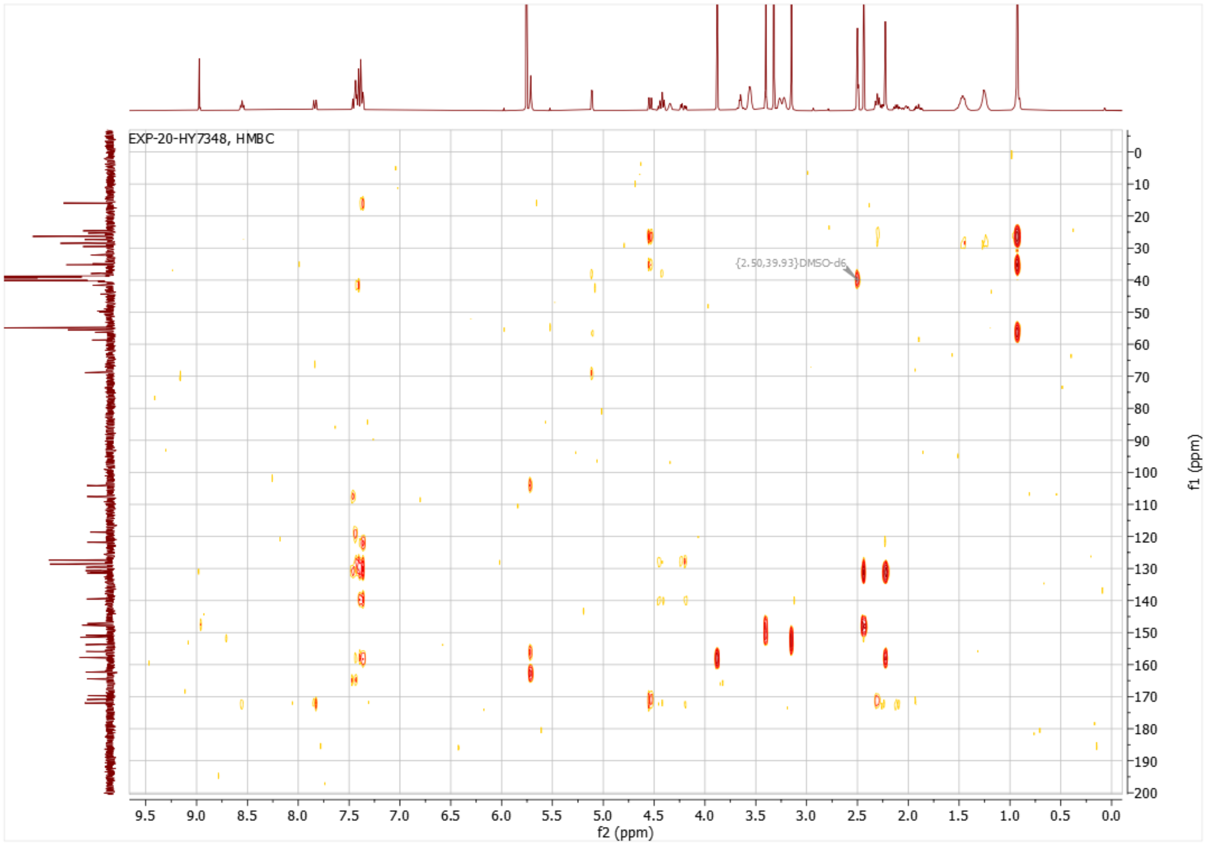

EXP-20-HY7348, HMBC
{2.50,39.93}DMSO-d6
f2 (ppm)
f1 (ppm)

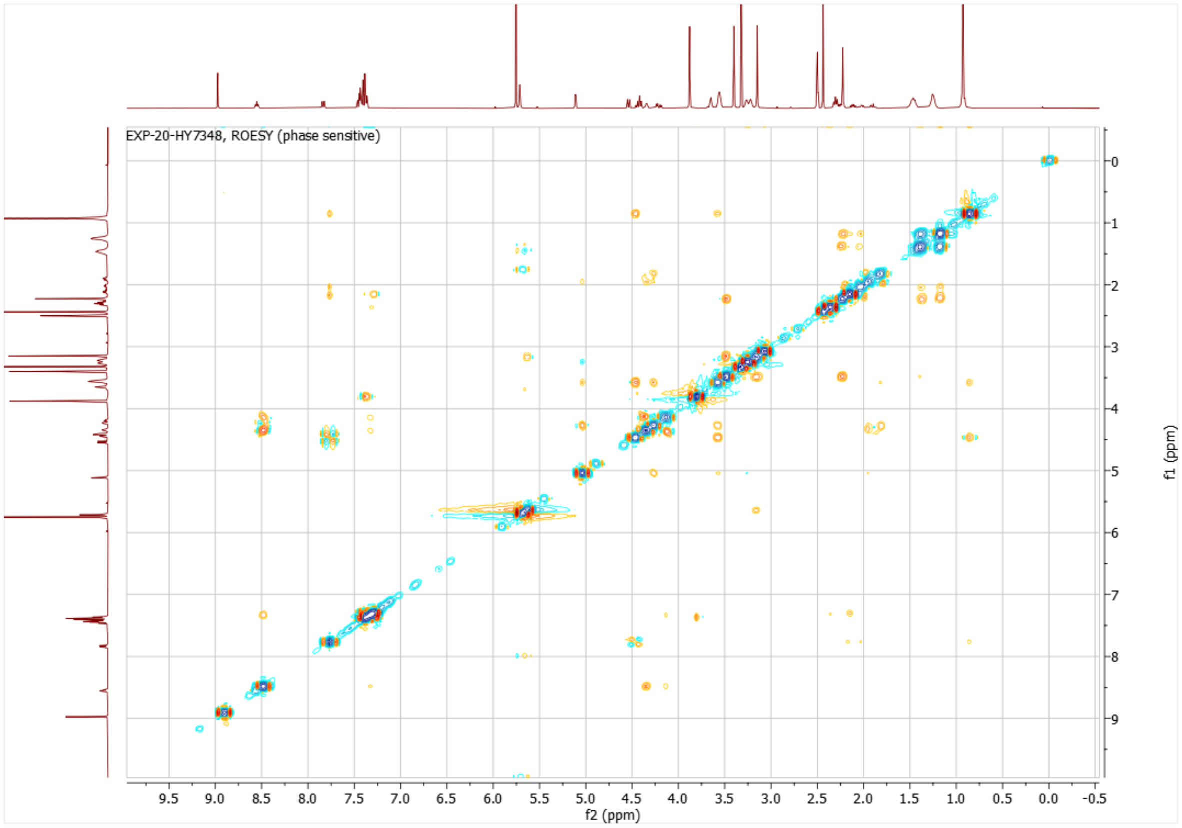

EXP-20-HY7348, ROESY (phase sensitive)
f2 (ppm)
f1 (ppm)

Preparation of **DDD3**

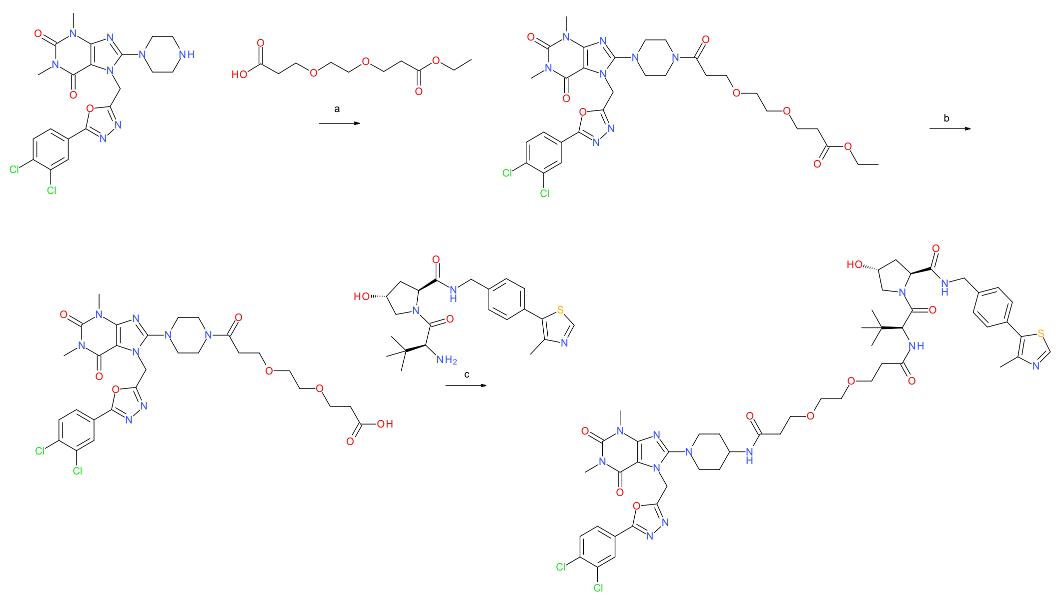

**Scheme S3.** Reagents and conditions: a) HATU, DIEA, DMF, rt, overnight; b) Bis(tributyltin) oxide, toluene, 115 °C for 12 h; c) HATU, DIEA, DMF, rt, overnight.

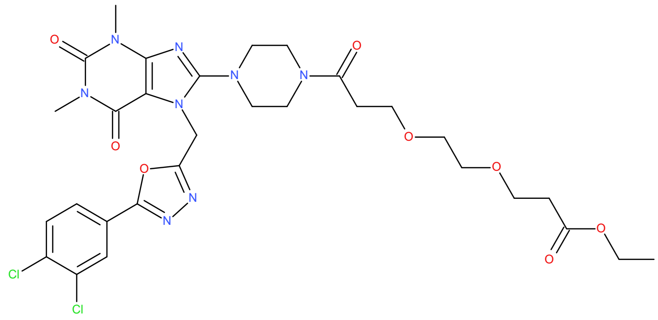

**Ethyl 3-[2-[3-[[1-[7-[[5-(3,4-dichlorophenyl)-1,3,4-oxadiazol-2-yl]methyl]-1,3-dimethyl-2,6-dioxo-purin-8-yl]-4-piperidyl]amino]-3-oxo-propoxy]ethoxy]propanoate (13a)**

3-[2-(3-Ethoxy-3- oxo-propoxy)ethoxy]propanoic acid (0.0381 g, 0.163 mmol), DIEA (0.0557 mL, 0.323 mmol) and compound **9a** (0.0800 g, 0.158 mmol) were dissolved in DMF (2 mL) and conditioned for 10 minutes at rt. HATU (0.0743 g, 0.195 mmol) was added and the reaction mixture left overnight. The mixture was diluted with EtOAc:water. The phases were separated and the organic phase washed with $NaHCO_3$ aqueous solution and brine and dried over $MgSO_4$. Removal of the solvent in vacuo yielded the title compound (83.0%, 0.0390 g, 0.0449 mmol, yield: 28%), which was used in the next step without further purification. MS (ESI$^+$) *m/z* 721 $[M+H]^+$.

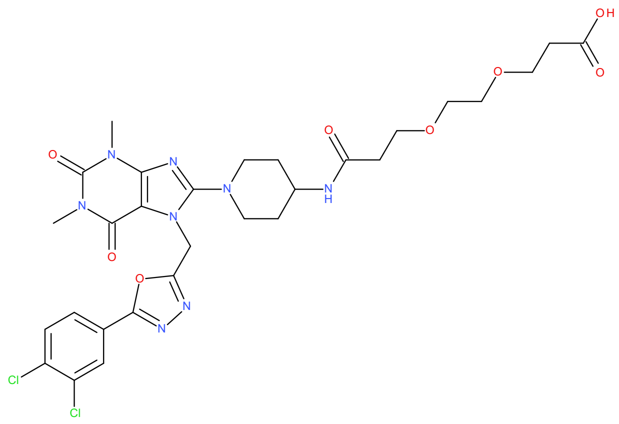

**3-[2-[3-[[1-[7-[[5-(3,4-Dichlorophenyl)-1,3,4-oxadiazol-2-yl]methyl]-1,3-dimethyl-2,6-dioxopurin-8-yl]-4-piperidyl]amino]-3-oxo-propoxy]ethoxy]propanoic acid (14a)**

Compound **13a** (0.0390 g, 0.0449 mmol) and bis(tributyltin) oxide (0.0932 mL, 0.174 mmol) were dissolved in toluene (1 mL) and the mixture stirred at 115 °C for 12 h. Excess toluene was removed in vacuo and the crude material was purified by flash column chromatography (0% to 10% MeOH in DCM), yielding the crude product back. Said crude product was then dissolved in a small volume of DMF and instead purified by preparative HPLC (Xbridge, gradient of 10% to 60% acetonitrile in 0.1% TFA). Lyophilization of the pure fractions yielded the title compound (73.0%, 0.0154 g, 0.0157 mmol, yield: 34%). MS (ESI$^-$) *m/z* 691 [M-H]$^-$.

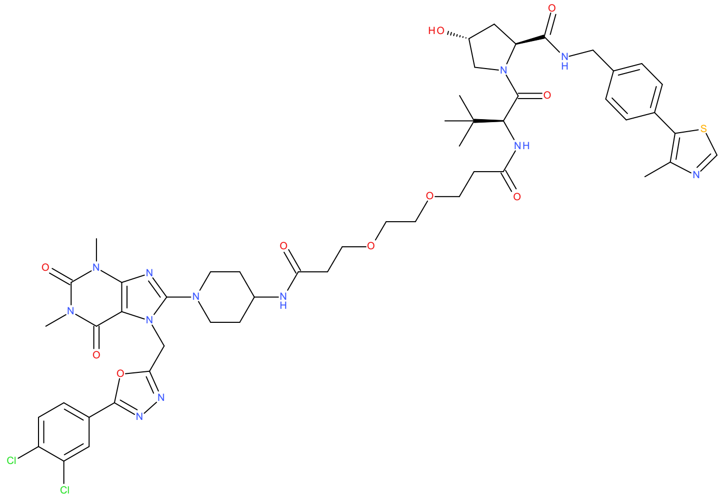

**(*2S*,*4R*)-1-[(*2S*)-2-[3-[2-[3-[[1-[7-[[5-(3,4-Dichlorophenyl)-1,3,4-oxadiazol-2-yl]methyl]-1,3-dimethyl-2,6-dioxo-purin-8-yl]-4-piperidyl]amino]-3-oxo-propoxy]ethoxy]propanoylamino]-3,3- dimethyl-butanoyl]-4-hydroxy-*N*-[[4-(4-methylthiazol-5-yl)phenyl]methyl]pyrrolidine-2-carboxamide (15a)**

Compound **14a** (0.0146 g, 0.0154 mmol), DIEA (0.00805 mL, 0.0461 mmol) and compound **2a** (0.00675 g, 0.0157 mmol) were dissolved in DMF (1.5 mL) and stirred for 10 minutes at rt. HATU (0.00716 g, 0.0188 mmol) was added and the reaction was left overnight. The mixture was diluted with EtOAc and washed with brine, concentrated, redissolved in a small volume of DMF and finally purified by preparative HPLC (gradient of 10% to 60% acetonitrile in 50 mM ammonium bicarbonate). Lyophilizing the pure fractions afforded the title compound (>99%, 2.10 mg, 1.90 μmol, yield: 12%). MS (ESI$^+$) *m/z* 554 $[M+2H]^{2+}$. HRMS: *m/z* calculated 1127.3708, found 1127.3717 for $[M+Na]^+$. $^1$H NMR (400 MHz, DMSO-$d_6$): δ 8.98 (s, 1H), 8.57 (t, J=6.0 Hz, 1H), 8.15 (d, J=1.8 Hz, 1H), 7.85-7.95 (m, 4H), 7.37-7.44 (m, 4H), 5.67 (s, 2H), 5.12 (d, J=3.2 Hz, 1H), 4.55 (d, J=9.4 Hz, 1H), 4.40-4.47 (m, 2H), 4.33-4.38 (m, 1H), 4.22 (dd, J=16 Hz, 5.2 Hz, 1H), 3.72-3.80 (m, 1H), 3.51-3.70 (m, 8H), 3.43- 3.49 (m, 4H), 3.41 (s, 3H), 3.16 (s, 3H), 3.08 (t, J=11.6 Hz, 2H), 2.54 (t, J=5.7 Hz, 2H), 2.44 (s, 3H), 2.30 (t, J=6.5 Hz, 1H), 1.98-2.05 (m, 1H), 1.87-1.99 (m, 1H), 1.75-1.82 (m, 2H), 1.44-1.55 (m, 2H), 0.93 (s, 9H).

<u>Preparation of **DDD4**</u>

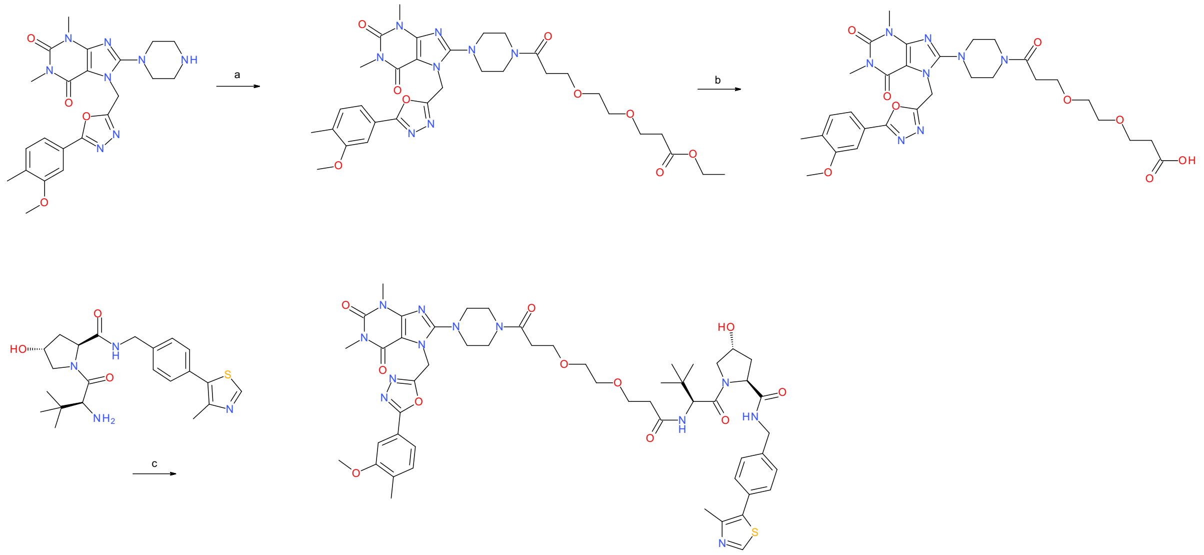

**Scheme S4.** Reagents and conditions: a) HATU, DIEA, DMF, rt, overnight; b) $LiOH \cdot H_2O$, water:THF (1:1), rt, 2.5 h; c) HATU, DIEA, DMF, rt, overnight.

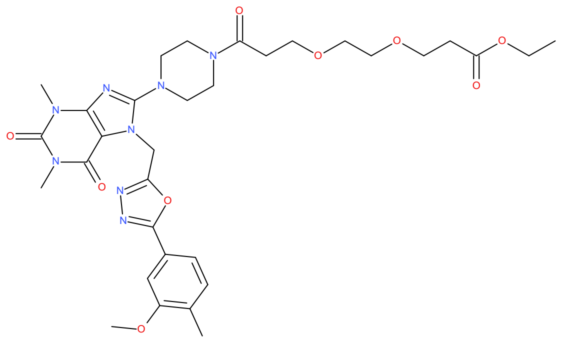

**Ethyl 3-[2-[3-[4-[7-[[5-(3-methoxy-4-methyl-phenyl)-1,3,4-oxadiazol-2-yl]methyl]-1,3-dimethyl-2,6-dioxo-purin-8-yl]piperazin-1-yl]-3-oxo-propoxy]ethoxy]propanoate (13b)**

3-[2-(3-Ethoxy-3- oxo-propoxy)ethoxy]propanoic acid (0.0251 g, 0.107 mmol), DIEA (0.0367 mL, 0.212 mol) and HATU (0.0489 g, 0.129 mmol) were dissolved in DMF (1 mL) and conditioned for 10 minutes at rt. Compound **9c** (0.0500 g, 0.107 mmol) was then added and the reaction mixture left overnight. The mixture was then diluted with EtOAc (25 mL) and the organic phase was washed with aqueous $NaHCO_3$ solution and brine. The product was dried over magnesium sulphate, and the solvent was removed under pressure. The crude product was purified by flash column chromatography with MeOH in DCM (0% to 10%) as eluent to afford the title compound (0.0440 g, 0.0644 mmol, yield: 60%). The crude material was used in the next step without further purification. MS ($ESI^+$) *m/z* 683 $[M+H]^+$.

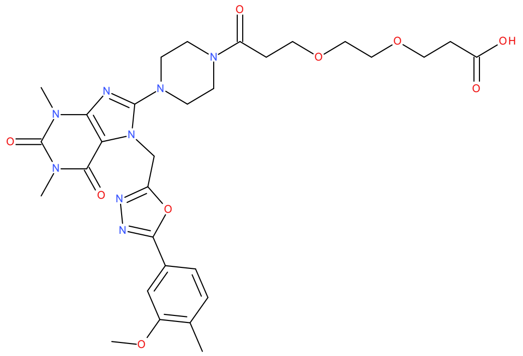

**3-[2-[3-[4-[7-[[5-(3-Methoxy-4-methyl-phenyl)-1,3,4-oxadiazol-2-yl]methyl]-1,3-dimethyl-2,6- dioxo-purin-8-yl]piperazin-1-yl]-3-oxo-propoxy]ethoxy]propanoic acid (14b)**

Compound **13b** (0.0440 g, 0.0644 mol) and $LiOH \cdot H_2O$ (0.00900 g, 0.211 mmol) were stirred in water:THF (1:1, 2 mL) in a closed vessel for 2.5 h. Workup was made via removal of the THF, acidification until pH=2. The material was extracted with EtOAc (4x), dried over $MgSO_4$ and solvent removal to afford the title compound (75%, 0.0180 g, 0.0275 mmol, yield: 43%), which was used in the next step without further purification. MS ($ESI^+$) *m/z* 655 $[M+H]^+$.

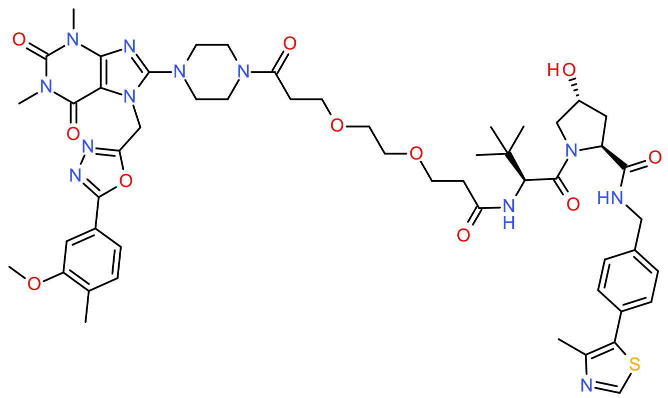

**(*2S*,*4R*)-4-Hydroxy-1-[(*2S*)-2-[3-[2-[3-[4-[7-[[5-(3-methoxy-4-methyl-phenyl)-1,3,4-oxadiazol-2- yl]methyl]-1,3-dimethyl-2,6-dioxo-purin-8-yl]piperazin-1-yl]-3-oxo-propoxy]ethoxy] propanoylamino]-3,3-dimethyl-butanoyl]-*N*-[[4-(4-methylthiazol-5-yl)phenyl]methyl] pyrrolidine-2-carboxamide (15b)**:

Compound **14b** (0.0180 g, 0.0206 mmol), DIEA (0.0106 mL, 0.0613 mmol) and HATU (0.00941 g, 0.0247 mmol) were dissolved in DMF (0.5 mL) and stirred for 10 minutes at rt. Compound **2a** (0.0197 g, 0.0458 mmol) was added and the reaction was left overnight. An additional 5 μL of DIEA and 5 mg of HATU were added and the mixture left for 5 h. The mixture was then diluted with DCM, washed with brine and dried in vacuo. The crude product was redissolved in acetonitrile and purified by preparative HPLC (gradient of 5% to 40% acetonitrile in 50 mM ammonium bicarbonate). The collected fractions were lyophilized to afford the title compound (>99%, 10.0 mg, 8.94 μmol, yield: 43%). HRMS: *m/z* calculated 1089.4593, found 1089.4522 for $[M+Na]^+$. $^1$H NMR (600 MHz, DMSO-$d_6$): δ 8.96 (s, 1H, H5), 8.55 (t, J=6.0 Hz, 1H), 7.90 (d, J=9.4 Hz, 1H), 7.37-7.48 (m, 7H), 5.72 (s, 2H), 5.12 (d, J=3.6 Hz, 1H), 4.55 (d, J=9.4 Hz, 1H), 4.41-4.46 (m, 2H), 4.33-4.37 (m, 1H), 4.22 (dd, J=16 Hz, 5.4 Hz, 1H), 3.89 (s, 3H), 3.55-3.69 (m, 10H), 3.44-3.51 (m, 4H), 3.41 (s, 3H), 3.21-3.30 (m, 4H), 3.16 (s, 3H), 2.60 (t, J=6.7 Hz, 2H), 2.51-2.55 (m, 1H), 2.44 (s, 3H), 2.36 (dt, J=15 Hz, 6.2 Hz, 1H), 2.23 (s, 3H), , 0.93 (s, 9H). $^{13}$C NMR (151 MHz, DMSO-$d_6$): δ 172.4, 170.4, 170.0, 169.6 (C39), 164.9, 162.8, 158.2, 156.4, 154.3, 151.9, 151.4 , 148.2, 147.6, 140.0, 131.9, 131.6 , 131.2, 130.1, 129.1 (two C overlapping), 127.9 (two C overlapping), 122.3 , 119.1, 108.1, 104.6, 70.1, 69.9 , 69.3 , 67.4, 67.2 , 59.2, 56.8, 56.7, 56.0, 50.4, 50.1, 44.9, 42.1, 40.9, 40.6, 38.4 , 36.1, 35.8, 33.2, 30.0, 27.9, 26.8 (three C overlapping), 16.6 , 16.4.

<u>Preparation of **DDD5**</u>

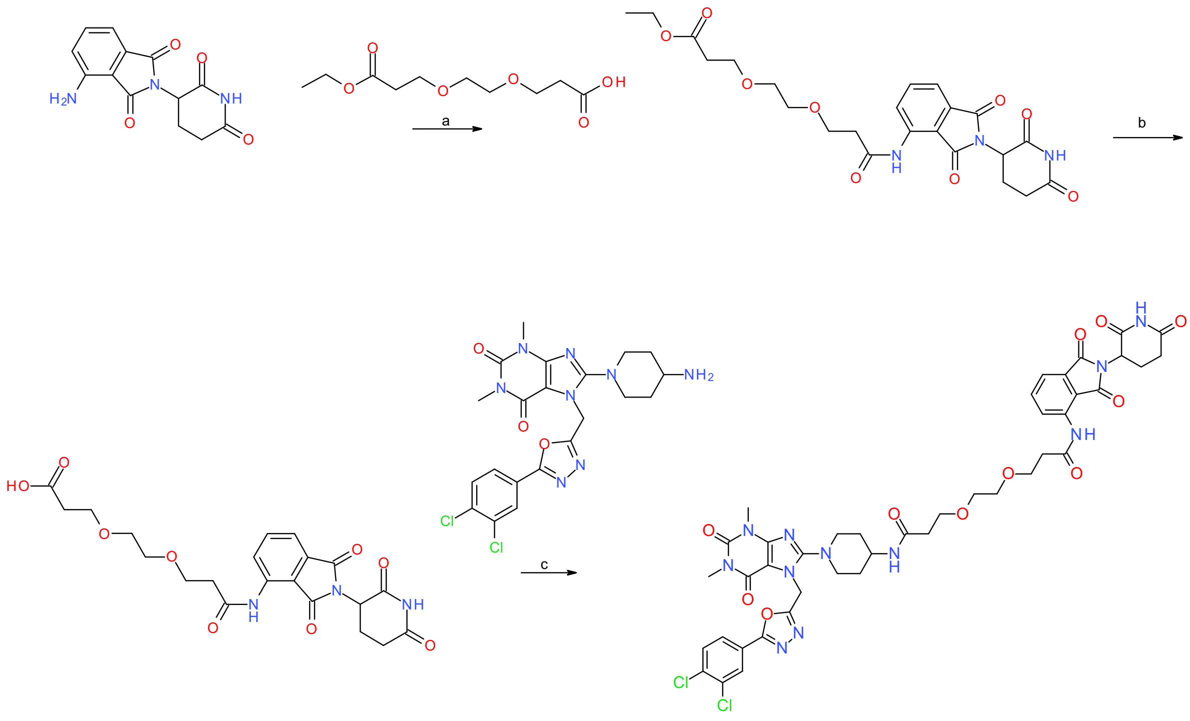

**Scheme S5.** Reagents and conditions: a) oxalyl chloride, DMF (catalytic amount), diethyl ether, THF, reflux, overnight; b) Bis(tributyltin) oxide, toluene, 115 °C, overnight; c) HATU, DIEA, DMF, rt, overnight.

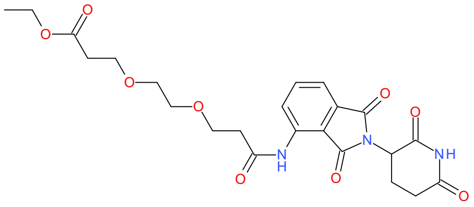

**Ethyl 3-[2-[3-[[2-(2,6-dioxo-3-piperidyl)-1,3-dioxo-isoindolin-4-yl]amino]-3-oxo-propoxy]ethoxy] propanoate (6)**

3-[2-(3-Ethoxy-3-oxo-propoxy)ethoxy]propanoic acid (0.193 g, 0.823 mmol), DMF (catalytic amount) and oxalyl chloride (0.0929 g, 0.717 mmol) were dissolved in diethyl ether (0.5 mL) and conditioned under a nitrogen atmosphere for a few minutes. Pomalidomide (**2b**) (0.100 g, 0.366 mmol) and THF (4 mL) were added to the mixture and the mixture was heated at reflux overnight. The mixture was added to a larger volume of methanol, EtOAc and brine, and the phases separated. The organic phase was washed twice with brine, dried over $MgSO_4$ and in vacuoto afford the title compound (94.0%, 0.236 g, 0.455 mmol, yield: >99%), which was used in the next step without further purification. MS (ESI$^+$) *m/z* 490 [M+H]$^+$.

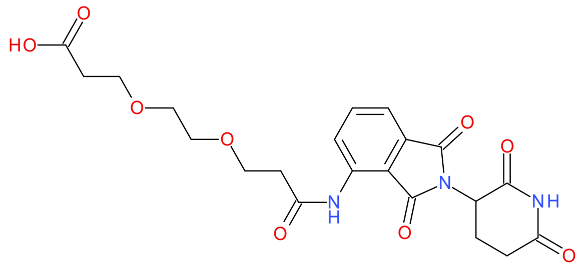

**3-[2-[3-[[2-(2,6-Dioxo-3-piperidyl)-1,3-dioxo-isoindolin-4-yl]amino]-3-oxo-propoxy]ethoxy]propanoic acid (7)**

Bis(tributyltin) oxide (0.343 g, 0.547 mmol) was dissolved in toluene (0.75 mL) and added to ethyl 3-[2-[3-[[2-(2,6-dioxo-3-piperidyl)-1,3-dioxo-isoindolin-4-yl]amino]-3-oxo-propoxy]ethoxy] propanoate (**6**) (0.0750 g, 0.144 mmol). The mixture was heated at 115 °C overnight. The toluene was evaporated in vacuo, after which the crude product was purified by flash column chromatography (MeOH in DCM, 5% to 15%), yielding the title compound (90.0%, 0.0890 g, 0.174 mmol, yield: >99%). MS (ESI$^-$) *m/z* 460 [M-H]$^-$.

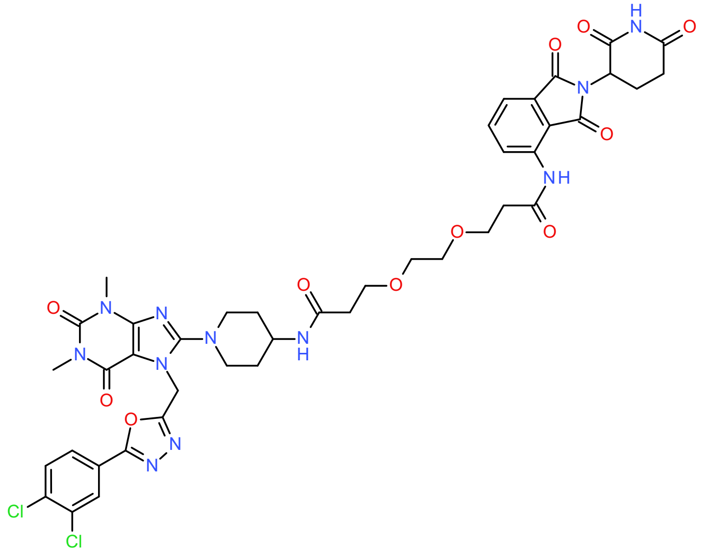

**3-[2-[3-[[1-[7-[[5-(3,4-Dichlorophenyl)-1,3,4-oxadiazol-2-yl]methyl]-1,3-dimethyl-2,6-dioxo-purin-8-yl]-4-piperidyl]amino]-3-oxo-propoxy]ethoxy]-N-[2-(2,6-dioxo-3-piperidyl)-1,3-dioxo-isoindolin-4-yl]propenamide**

8-(4-Amino-1-piperidyl)-7-[[5-(3,4-dichlorophenyl)-1,3,4-oxadiazol-2-yl]methyl]-1,3-dimethyl-purine-2,6-dione (**9a**) (0.0150 g, 0.0297 mmol), DIEA (0.0105 mL, 0.0605 mmol) and 3-[2-[3-[[2-(2,6-dioxo-3-piperidyl)-1,3-dioxo-isoindolin-4-yl]amino]-3-oxo propoxy]ethoxy]propanoic acid (**7**) (0.0157 g, 0.0306 mmol) were dissolved in DMF (0.5 mL) and conditioned for 10 minutes at rt. HATU (0.0139 g, 0.0367 mmol) was then added and the reaction mixture left at rt overnight. The mixture was diluted with EtOAc and washed with aqueous $NaHCO_3$ solution, brine, and water. The solvent was removed, and the crude product redissolved in about 1 mL of DMF and purified by preparative HPLC (gradient of 10% to 60% acetonitrile in 0.1% TFA). Lyophilization of the pure fractions yielded the title compound (>99%, 7.80 mg, 8.22 µmol, yield: 28%). HRMS: *m/z* calculated 948.2599, found 948.2598 for $[M+H]^+$. $^1$H NMR (400 MHz, DMSO-$d_6$): δ 11.15 (s, 1H), 9.85 (s, 1H), 8.53 (d, J=8.3 Hz, 1H), 8.14 (d, J=2.0 Hz, 1H), 7.79-7.94 (m, 4H), 7.60 (d, J=7.5 Hz, 1H), 5.68 (s, 2H), 5.15 (dd, J=13 Hz, 5.5 Hz, 1H), 3.70-3.76 (m, 3H), 3.48-3.62 (m, 8H), 3.41 (s, 3H), 3.15 (s, 3H), 3.02-3.11 (m, 2H), 2.90 (td, J=15 Hz, 5.7 Hz, 1H), 2.69 (t, J=6.0 Hz, 2H), 2.54-2.65 (m, 2H), 2.27 (t, J=6.4 Hz, 2H), 2.03-2.12 (m, 2H), 1.74-1.82 (m, 2H), 1.43-1.54 (m, 2H).

Preparation of **DDD6**

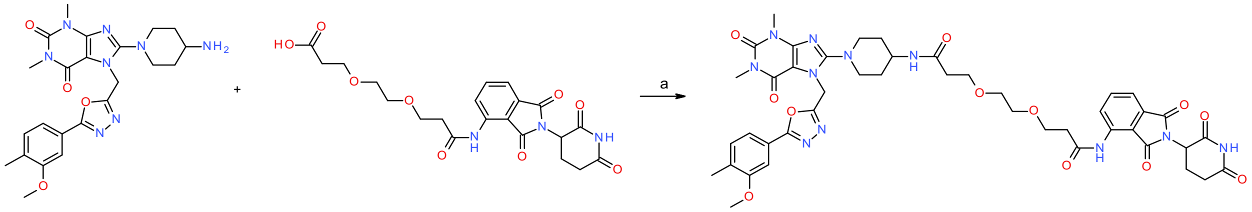

**Scheme S6.** Reagents and conditions: a) HATU, DIEA, DMF, rt, overnight.

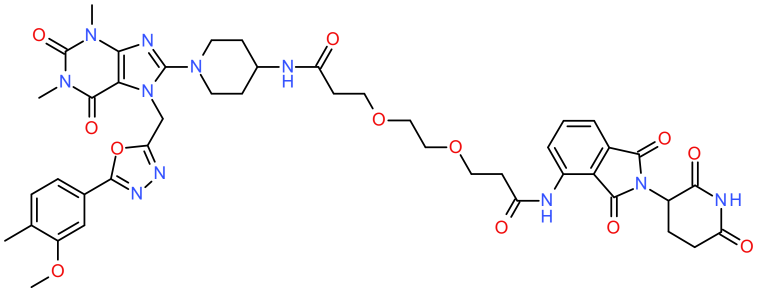

**N-[2-(2,6-Dioxo-3-piperidyl)-1,3-dioxo-isoindolin-4-yl]-3-[2-[3-[[1-[7-[[5-(3-methoxy-4-methyl-phenyl)-1,3,4-oxadiazol-2-yl]methyl]-1,3-dimethyl-2,6-dioxo-purin-8-yl]-4-piperidyl]amino]-3- oxo-propoxy]ethoxy]propanamide (15d)**

8-(4-Amino-1-piperidyl)-7-[[5-(3-methoxy-4-methyl-phenyl)-1,3,4-oxadiazol-2-yl]methyl]-1,3-dimethyl-purine-2,6-dione (**9b**) (0.0143 g, 0.0306 mmol), DIEA (0.0105 mL, 0.0605 mmol) and 3-[2-[3-[[2-(2,6-dioxo-3-piperidyl)-1,3-dioxo-isoindolin-4-yl]amino]-3-oxo-propoxy]ethoxy] propanoic acid (**7**) 0.0157 g, 0.0306 mmol) were dissolved in DMF (0.5 mL) and conditioned for 10 minutes at rt. HATU (0.0139 g, 0.0367 mmol) was then added and the reaction mixture left at rt overnight. The mixture was diluted with EtOAc and the mixture was washed with aqueous $NaHCO_3$ solution, brine and water. The solvent was removed, and the crude product redissolved in about 1 mL of DMF and purified by preparative HPLC (Xbridge, gradient of 10% to 60% acetonitrile in 0.1% TFA). Lyophilization of the pure fractions afforded the title compound (>99% 5.60 mg, 6.15 µmol, yield: 20%). MS ($ESI^+$) *m/z* 924 $[M+H]^+$. HRMS: *m/z* calculated 924.3640, found 924.3637 for $[M+H]^+$.

<u>Preparation of **DDD7**</u>

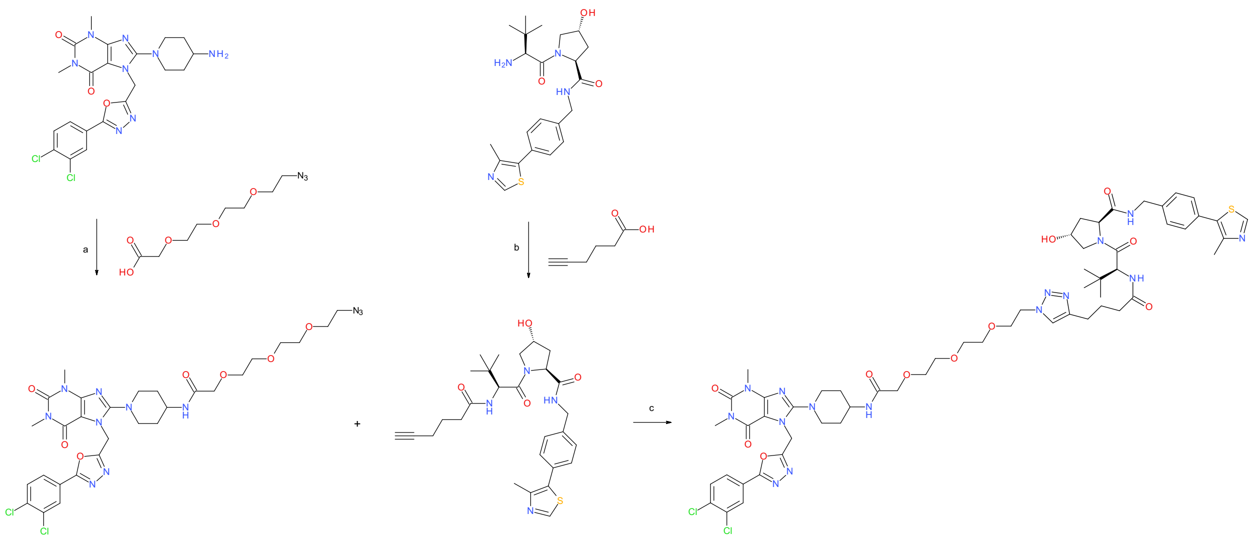

**Scheme S7.** Reagents and conditions: a) HATU, DIEA, DMF, rt, 3 h; b) HATU, DIEA, DMF, rt, overnight; c) sodium *L*-ascorbate, $CuSO_4 \cdot 5\ H_2O$, DCM:*t*-BuOH (1:2) mixture, water, rt overnight.

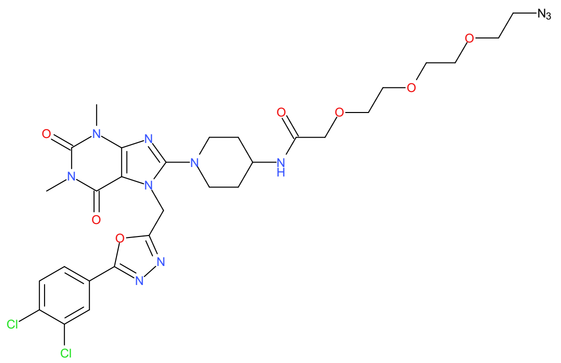

**2-[2-[2-(2-Azidoethoxy)ethoxy]ethoxy]-N-[1-[7-[[5-(3,4-dichlorophenyl)-1,3,4-oxadiazol-2-yl]methyl]-1,3-dimethyl-2,6-dioxo-purin-8-yl]-4-piperidyl]acetamide (16a)**

Compound **9a** (0.100 g, 0.198 mmol), DIEA (0.0697 mL, 0.403 mmol) and 2-[2-[2-[2-azidoethoxy]ethoxy]ethoxy]acetic acid (0.0477 g, 0.204 mmol) were dissolved in DMF (2 mL) and conditioned for 10 minutes at rt. HATU (0.0929 g, 0.244 mmol) was added and the reaction mixture stirred overnight. The mixture was then diluted with water and EtOAc and the phases were separated. The organic phase was washed with saturated aqueous $NaHCO_3$ and brine, dried over $MgSO_4$. Removal of the solvent in vacuo yielded the title compound (96.0%, 0.0810 g, 0.108 mmol, yield: 54%) which was used in the next step without further purification. MS ($ESI^+$) *m/z* 720 $[M+H]^+$.

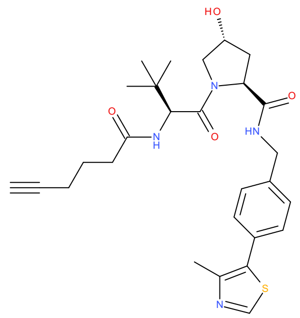

**(*2S,4R*)-1-[(*2S*)-2-(Hex-5-ynoylamino)-3,3-dimethyl-butanoyl]-4-hydroxy-*N*-[[4-(4-methylthiazol-5-yl)phenyl]methyl]pyrrolidine-2-carboxamide (5a)**

Hex-5-ynoic acid (0.0130 g, 0.113 mmol), DIEA (0.0596 mL, 0.344 mmol) and compound **2a** (0.0500 g, 0.116 mol) were dissolved in DMF (1.25 mL) and stirred for 10 minutes at rt. HATU (0.0530 g, 0.139 mmol) was added and the reaction was left overnight. The mixture was diluted with EtOAc. The organic phase was washed with saturated aqueous $NaHCO_3$ and brine and concentrated in vacuo. This yielded a small amount of product - LCMS analysis showed product still present in the aqueous extracts. The combined aqueous phases were thus washed with DCM (3x), and the combined organic phases were then washed with brine and concentrated in vacuo, yielding the title compound (73.0%, 0.0550 g, 0.0765 mmol, yield: 68%). The crude material was used in the next step without further purification. MS ($ESI^+$) *m/z* 525 $[M+H]^+$. $^1H$ NMR (400 MHz, DMSO-$d_6$): δ 8.99 (s, 1H), 8.57 (t, J=6.1 Hz, 1H), 7.92 (d, J=9.3 Hz, 1H), 7.37-7.45 (m, 4H), 5.13

(d, J=J.5 Hz, 1H), 4.54 (d, J=9.3 Hz, 1H), 4.41-4.47 (m, 2H), 4.33-4.48 (m, 1H), 4.22 (dd, J=16 Hz, 5.4 Hz, 1H), 3.62- 3.70 (m, 2H), 2.78 (t, J=2.7 Hz, 1H), 2.45 (s, 3H), 2.20-2.38 (m, 2H), 2.14 (td, J=7.0 Hz, 2.5 Hz, 2H), 2.01-2.07 (m, 1H), 1.86-1.94 (m, 1H), 1.60-1.73 (m, 2H), 0.94 (s, 9H).

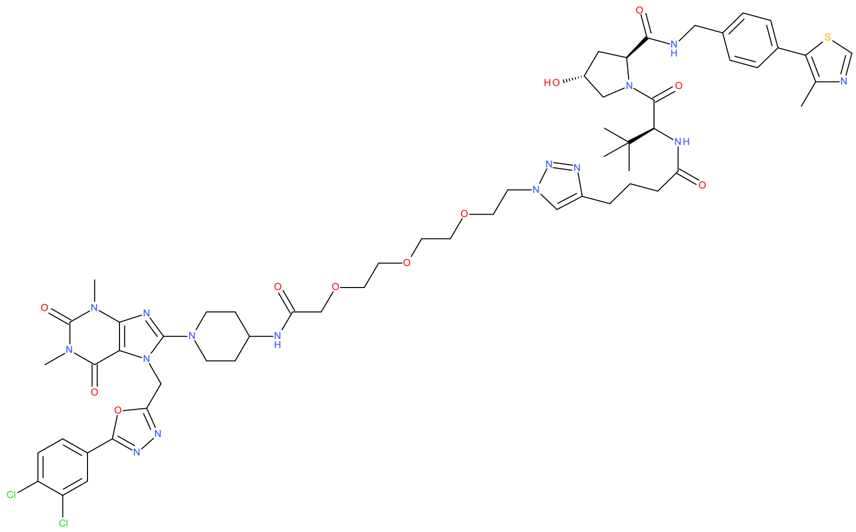

**(*2S*,*4R*)-1-[(*2S*)-2-[4-[1-[2-[2-[2-[2-[[1-[7-[[5-(3,4-Dichlorophenyl)-1,3,4-oxadiazol-2-yl]methyl]- 1,3-dimethyl-2,6-dioxo-purin-8-yl]-4-piperidyl]amino]-2-oxo-ethoxy]ethoxy]ethoxy]ethyl] triazol-4-yl]butanoylamino]-3,3-dimethyl-butanoyl]-4-hydroxy-*N*-[[4-(4-methylthiazol-5- yl)phenyl]methyl]pyrrolidine-2-carboxamide (17a)**

Compound **16a** (0.0276 g, 0.0368 mmol), compound **5a** (63.0%, 0.0260 g, 0.0312 mmol) and sodium *L*-ascorbate (0.0186 g, 0.0928 mmol) were dissolved in *t*-BuOH:DCM (2:1, 2.25 mL). $CuSO_4 \cdot 5\ H_2O$ (0.0156 g, 0.0618 mmol) in water (0.75 mL) was added and the mixture stirred at room temperature overnight. The reaction mixture was diluted with EtOAc and the resulting organic phase was washed with ammonia solution (28%), and a water:ammonia mixture. The solvent was removed and the resulting product redissolved in acetonitrile and purified by preparative HPLC (gradient of 10% to 60% acetonitrile in 50 mM ammonium bicarbonate buffer). Lyophilization of the pure fractions gave the title compound (92.0%, 2.70 mg, 1.99 µmol, yield: 6.4%). MS ($ESI^+$) *m/z* 623 $[M+2H]^{2+}$. HRMS: *m/z* calculated 1244.4633, found 1244.4670 for $[M+H]^+$.

---

---

Preparation of **DDD8**

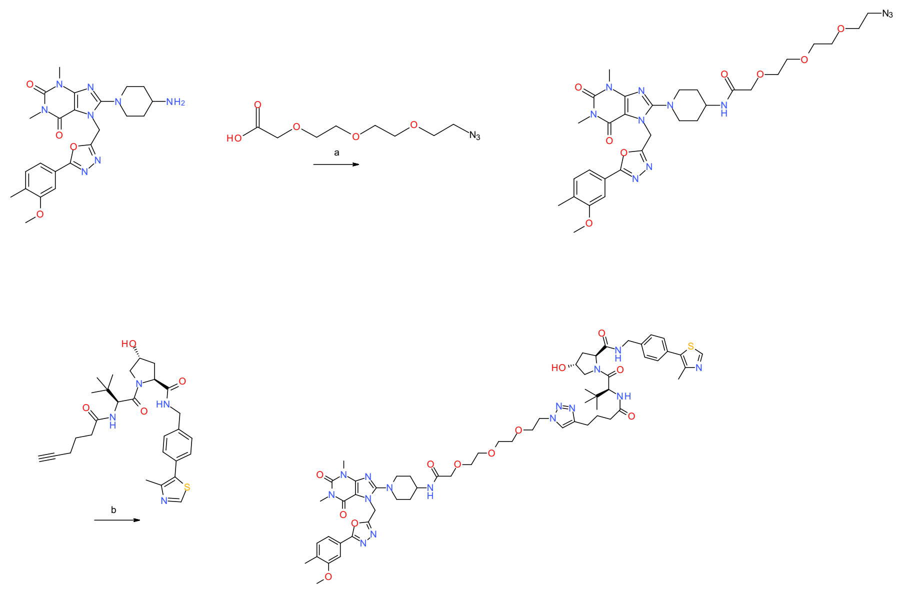

**Scheme S8.** Reagents and conditions: a) HATU, DIEA, DMF, rt, 3 h; b) sodium *L*-ascorbate, $CuSO_4 \cdot 5\ H_2O$, DCM:*t*-BuOH (1:2) mixture, water, rt overnight.

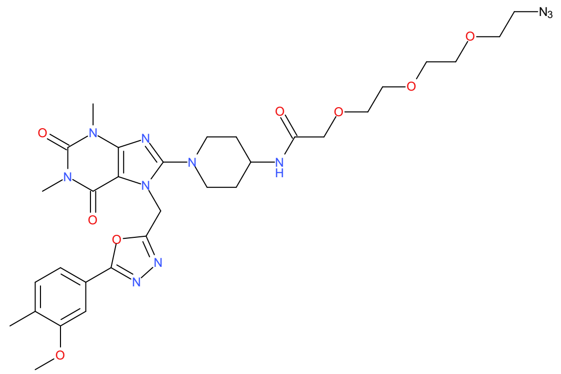

**2-[2-[2-(2-Azidoethoxy)ethoxy]ethoxy]-*N*-[1-[7-[[5-(3-methoxy-4-methyl-phenyl)-1,3,4-oxadiazol-2-yl]methyl]-1,3-dimethyl-2,6-dioxo-purin-8-yl]-4-piperidyl]acetamide (16b)**

Compound **9b** (0.0660 g, 0.121 mmol) prepared as described above, DIEA (0.0426 mL, 0.247 mmol) and 2-[2-[2-[2- azidoethoxy]ethoxy]ethoxy]acetic acid (0.0292 g, 0.125 mmol) were dissolved in DMF (5 mL) and conditioned for 10 minutes at rt. HATU (0.0568 g, 0.149 mmol) was added and the reaction mixture left over the weekend. The mixture was diluted with aqueous $NaHCO_3$ solution and EtOAc and the phases separated. The organic phase was washed further with aqueous $NaHCO_3$ and brine and dried over $MgSO_4$. Removal of the solvent in vacuo yielded the title compound (84.0%, 0.0780 g, 0.0942 mmol, yield: 78%), which was used in the next step without further purification. MS (ESI$^+$) *m/z* 696 $[M+H]^+$.

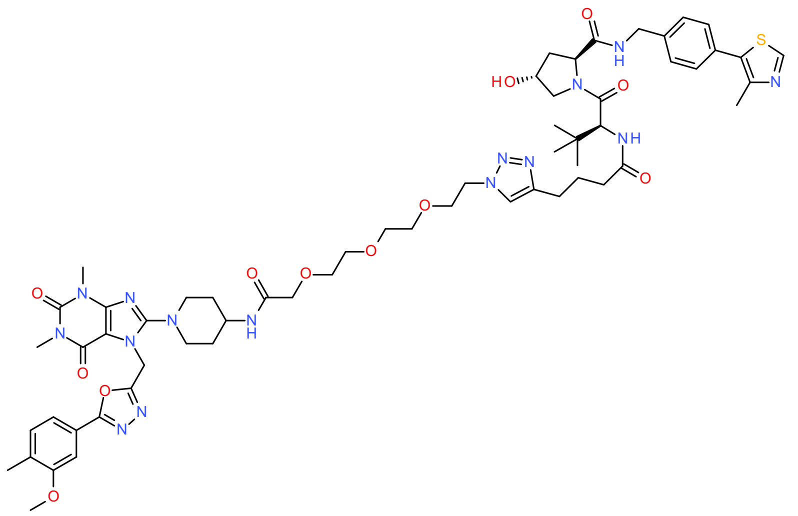

**(*2S*,*4R*)-4-Hydroxy-1-[(*2S*)-2-[4-[1-[2-[2-[2-[2-[[1-[7-[[5-(3-methoxy-4-methyl-phenyl)-1,3,4-oxadiazol-2-yl]methyl]-1,3-dimethyl-2,6-dioxo-purin-8-yl]-4-piperidyl]amino]-2-oxoethoxy]ethoxy]ethoxy]ethyl]triazol-4-yl]butanoylamino]-3,3-dimethyl-butanoyl]-*N*-[[4-(4- methylthiazol-5-yl)phenyl]methyl]pyrrolidine-2-carboxamide (17b)**

Compound **16b** (0.0244 g, 0.0301 mmol), compound **5a** prepared as described above in the preparation of compound DDD7 (73.0%, 0.0180 g, 0.0250 mmol) and sodium *L*-ascorbate (0.0149 g, 0.0743 mmol) were dissolved in *t*-BuOH:DCM (2:1, 1.5 mL). $CuSO_4 \cdot 5\ H_2O$ (0.0125 g, 0.0496 mmol) in water (0.5 mL) was added and the mixture stirred at room temperature overnight. The reaction mixture was diluted with EtOAc. The resulting organic phase was washed with ammonia solution (28%) with some water. The solvent was removed and the resulting product redissolved in acetonitrile and purified by preparative HPLC (gradient of 10% to 60% acetonitrile in 0.1% TFA). Lyophilization of the pure fractions yielded the title compound (99.0%, 3.80 mg, 3.12 µmol, yield: 12%). MS ($ESI^+$) *m/z* 611 $[M+2H]^{2+}$. HRMS: *m/z* calculated 1220.5675, found 1220.5693 for $[M+H]^+$. $^1H$ NMR (400 MHz, DMSO-$d_6$): δ 8.99 (s, 1H), 8.56 (t, J=6.1 Hz, 1H), 7.91 (d, J=9.3 Hz, 1H), 7.81 (s, 1H), 7.62 (d, J=8.0 Hz, 1H), 7.36-7.47 (m, 7H), 5.66 (s, 2H), 5.13 (d, J=3.6 Hz, 1H), 4.55 (d, J=9.4 Hz, 1H), 4.40-4.47 (m, 4H), 4.33-4.37 (m, 1H), 4.22 (dd, J=16 Hz, 5.4 Hz, 1H), 3.89 (s, 3H), 3.87 (s, 2H), 3.77 (t, J=5.4 Hz, 2H), 3.48-3.68 (m, 13H), 3.41 (s, 3H), 3.16 (s, 3H), 3.05-3.13 (m, 2H), 2.54-2.60 (m, 2H), 2.44 (s, 3H), 2.15-2.32 (m, 5H), 2.00-2.07 (m, 1H), 1.86-1.94 (m, 1H), 1.74-1.84 (m, 4H), 1.56-1.67 (m, 2H), 0.95 (s, 9H).

<u>Preparation of **DDD9**</u>

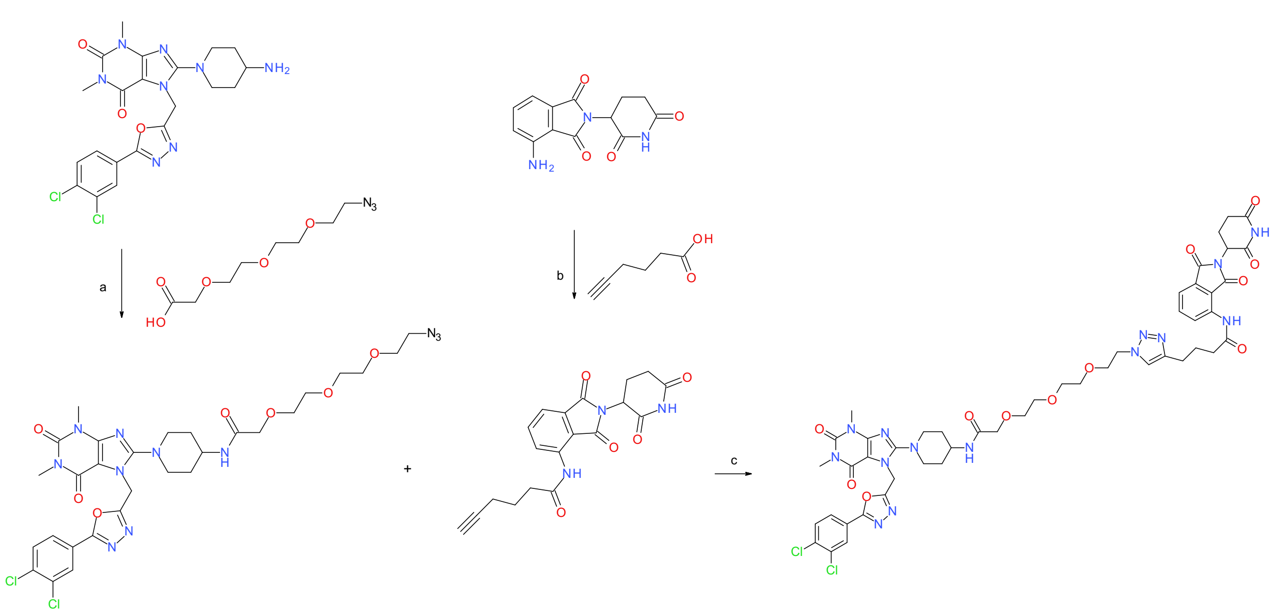

**Scheme S9.** Reagents and conditions: a) HATU, DIEA, DMF, rt, overnight; b) oxalyl chloride, DMF (catalytic amount), diethyl ether, THF, reflux, overnight; c) diethyl ether, THF, reflux, overnight; c) sodium *L*-ascorbate, $CuSO_4 \cdot 5\ H_2O$, DCM:tBuOH (1:2) mixture, water, rt, 30 min.

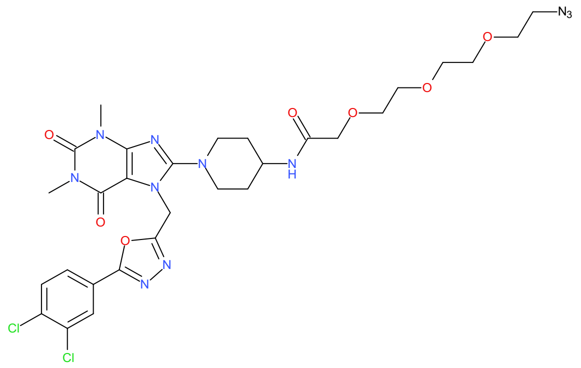

**2-[2-[2-(2-Azidoethoxy)ethoxy]ethoxy]-N-[1-[7-[[5-(3,4-dichlorophenyl)-1,3,4-oxadiazol-2-yl]methyl]-1,3-dimethyl-2,6-dioxo-purin-8-yl]-4-piperidyl]acetamide (16a)**

8-(4-Amino-1-piperidyl)-7-[[5-(3,4-dichlorophenyl)-1,3,4-oxadiazol-2-yl]methyl]-1,3-dimethyl-purine-2,6-dione (**9a**) (0.100 g, 0.198 mmol), DIEA (0.0697 mL, 0.403 mmol) and 2-[2-[2-[2-azidoethoxy]ethoxy]ethoxy]acetic acid (0.0477 g, 0.204 mmol) were dissolved in DMF (2 mL) and conditioned for 10 minutes at rt. HATU (0.0929 g, 0.244 mmol) was added and the reaction mixture left at rt overnight. The mixture was then diluted with water and EtOAc and the phases separated. The organic phase was washed with saturated aqueous $NaHCO_3$ and brine, after which the product was dried over $MgSO_4$. Removal of the solvent in vacuo yielded the title compound (96.0%, 0.0810 g, 0.108 mmol, yield: 54%) which was used in the next step without further purification. MS ($ESI^+$) *m/z* 720 $[M+H]^+$.

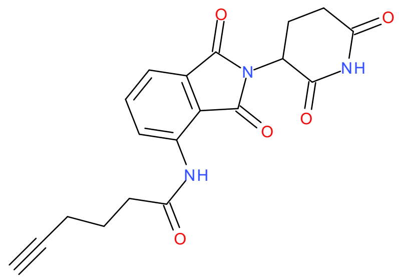

**N-[2-(2,6-Dioxo-3-piperidyl)-1,3-dioxo-isoindolin-4-yl]hex-5-ynamide (5b):**

Hex-5-ynoic acid (0.0923 g, 0.798 mmol), DMF (catalytic amount) and oxalyl chloride (0.0929 g, 0.717 mmol) were dissolved in diethyl ether (0.5 mL) and conditioned under a nitrogen atmosphere for 30 min. Pomalidomide (**2b**) (0.100 g, 0.366 mmol) and THF (4 mL) were added to the mixture and the mixture heated at reflux overnight. The crude product was suspended in water, filtered and dried in vacuo, yielding the title compound (98.0%, 0.104 g, 0.280 mmol, yield: 76%) which was used in the next step without further purification. MS ($ESI^+$) *m/z* 368 $[M+H]^+$. $^1H$ NMR (400 MHz, DMSO-$d_6$): δ 11.14 (s, 1H), 9.76 (s, 1H), 8.42 (d, J=8.4 Hz, 1H), 7.80-7.86 (m, 1H), 7.62 (d, J=7.7 Hz, 1H), 5.11-5.18 (m, 1H), 2.84-2.95 (m, 1H), 2.83 (t, J=2.6 Hz, 1H), 2.52-2.66 (m, 2H), 2.56 (t, J=7.4 Hz, 2H), 2.26 (td, 2H, J=7.4 Hz, 2.6 Hz), 2.02-2.11 (m, 1H), 1.80 (p, 2H).

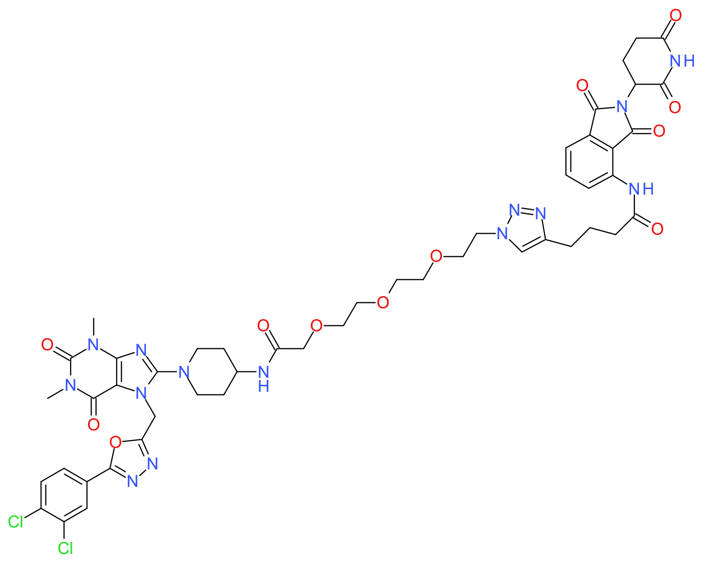

**4-[1-[2-[2-[2-[2-[[1-[7-[[5-(3,4-Dichlorophenyl)-1,3,4-oxadiazol-2-yl]methyl]-1,3-dimethyl-2,6-dioxo-purin-8-yl]-4-piperidyl]amino]-2-oxo-ethoxy]ethoxy] ethoxy]ethyl]triazol-4-yl]-*N*-[2-(2,6-dioxo-3-piperidyl)-1,3-dioxo-isoindolin-4-yl] butanamide (17c)**

2-[2-[2-(2-Azidoethoxy)ethoxy]ethoxy]-*N*-[1-[7-[[5-(3,4-dichlorophenyl)-1,3,4-oxadiazol-2-yl]methyl]-1,3-dimethyl-2,6-dioxo-purin-8-yl]-4-piperidyl]acetamide (**16a**) (0.0347 g, 0.0462 mmol), *N*-[2-(2,6-Dioxo-3-piperidyl)-1,3-dioxo-isoindolin-4-yl]hex-5-ynamide (**5b**) (0.0150 g, 0.0392 mmol) and sodium *L*-ascorbate (0.0233 g, 0.117 mmol) were dissolved in *t*-BuOH:DCM (2:1, 2.25 mL). $CuSO_4 \cdot 5\ H_2O$ (0.0196 g, 0.0776 mmol) in water (0.75 mL) was added and the mixture stirred at room temperature for 30 minutes. EtOAc was added, and the resulting organic phase was washed with ammonia solution (28%), brine, and water. The organic solvent was removed and the resulting product redissolved in DMF and purified by preparative HPLC (gradient of 10% to 60% acetonitrile in 0.1% TFA). Lyophilization of the pure fractions yielded the title compound (91.0%, 11.4 mg, 9.62 µmol, yield: 24%). MS (ESI$^+$) *m/z* 1087 [M+H]$^+$. HRMS: *m/z* calculated 1087.3344, found 1087.3358 for [M+H]$^+$. $^1$H NMR (600 MHz, DMSO-$d_6$): δ 11.14 (s, 1H), 9.70 (s, 1H), 8.45 (d, J=8.4 Hz, 1H), 8.14 (d, J=2.0 Hz, 1H), 7.92 (dd, J=8.4, 2.0 Hz, 1H), 7.88 (d, J=8.4 Hz, 1H), 7.85 (s, 1H), 7.82 (dd, J=8.3, 7.5 Hz, 1H), 7.61 (dd, J=7.3, 0.46 Hz, 1H), 7.59 (d, J=6.0 Hz, 1H), 5.67 (s, 2H), 5.14 (dd, J=13, 5.4 Hz, 1H), 4.46 (t, J=5.3 Hz, 1H), 3.86 (s, 2H), 3.81-3.85 (m, 1H), 3.79 (t, J=5.3 Hz, 2H), 3.46-3.68 (m, 10H), 3.41 (s, 3H), 3.15 (s, 3H), 3.07-3.11 (m, 2H), 2.86-2.94 (m, 1H), 2.68 (t, J=7.6 Hz, 2H), 2.51-2.58 (m, 4H), 2.04-2.10 (m, 1H), 1.94 (p, J=7.5 Hz, 2H), 1.74-1.81 (m, 2H), 1.55-1.64 (m, 2H). $^{13}$C NMR (151 MHz, DMSO-$d_6$): δ 173.2, 172.2, 170.2, 169.0 , 168.1, 167.1, 163.6, 163.1, 157.0, 154.2, 151.4, 147.8, 146.5 , 137.0, 136.5, 135.4, 132.9, 132.5, 131.9, 128.6, 127.1 , 126.8, 124.0, 122.8, 118.8, 117.5, 104.3, 70.6, 70.3 , 70.0 (three C overlapping), 69.2, 49.7, 49.4 (two C overlapping), 45.5, 41.0, 40.6, 36.3, 31.4, 31.3 (two C overlapping), 30.0, 27.8, 25.1, 24.9, 22.5.

Preparation of **DDD10**

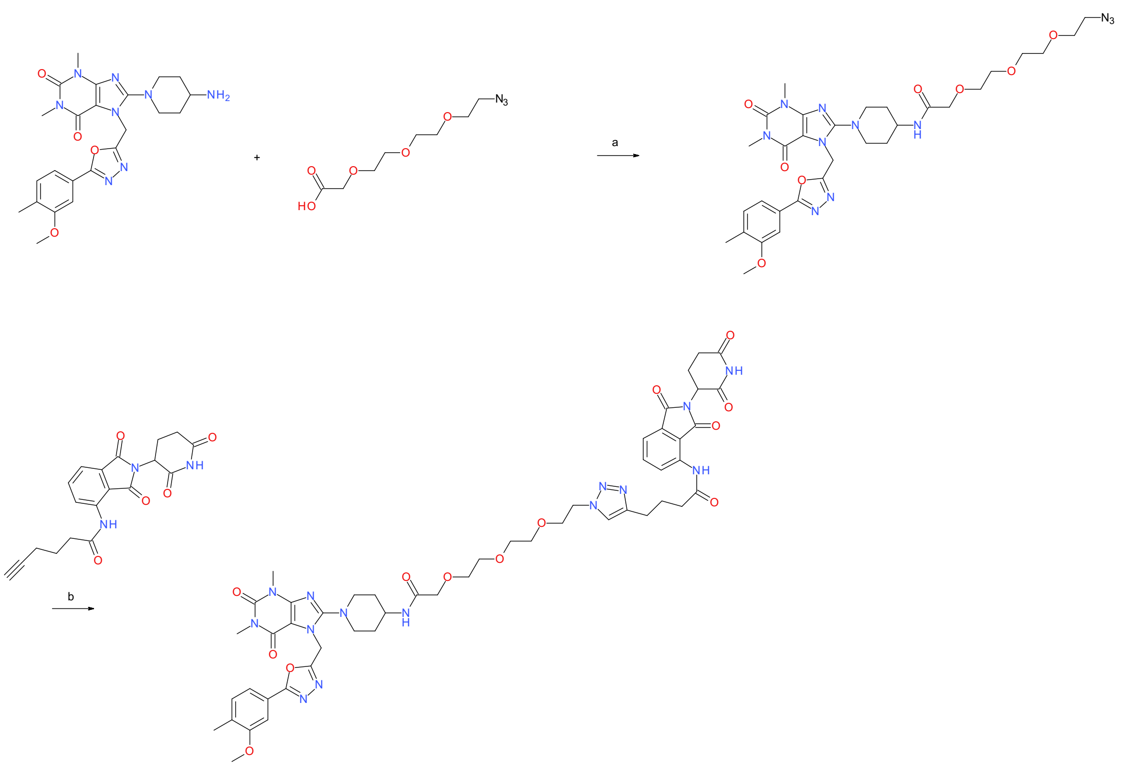

**Scheme S10.** Reagents and conditions: a) HATU, DIEA, DMF, rt, overnight; b) sodium *L*-ascorbate, $CuSO_4 \cdot 5\ H_2O$, DCM:*t*-BuOH (1:2) mixture, water, rt, 2 h.

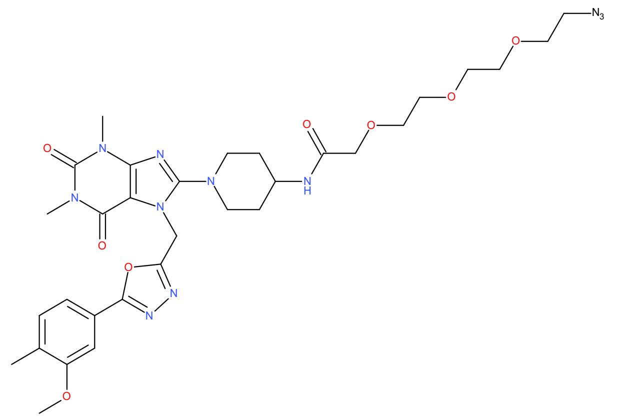

**2-[2-[2-(2-Azidoethoxy)ethoxy]ethoxy]-*N*-[1-[7-[[5-(3-methoxy-4-methyl-phenyl)-1,3,4-oxadiazol-2-yl]methyl]-1,3-dimethyl-2,6-dioxo-purin-8-yl]-4-piperidyl]acetamide (16b)**

8-(4-Amino-1-piperidyl)-7-[[5-(3-methoxy-4-methyl-phenyl)-1,3,4-oxadiazol-2-yl]methyl]-1,3-dimethyl-purine-2,6-dione (**9b**) (0.0660 g, 0.121 mmol), DIEA (0.0426 mL, 0.247 mmol) and 2-[2-[2-[2- azidoethoxy]ethoxy]ethoxy]acetic acid (0.0292 g, 0.125 mmol) were dissolved in DMF (5 mL) and conditioned for 10 minutes at rt. HATU (0.0568 g, 0.149 mmol) was added and the reaction mixture left at rt over the weekend. The mixture was diluted with water/$NaHCO_3$ and EtOAc and the phases separated. The organic phase was washed with aqueous $NaHCO_3$ solution and brine, after which the product was dried over $MgSO_4$. Removal of the solvent in vacuo yielded the title compound (84.0%, 0.0780 g, 0.0942 mmol, yield: 78%) which was used in the next step without further purification. MS (ESI$^+$) *m/z* 696 $[M+H]^+$.

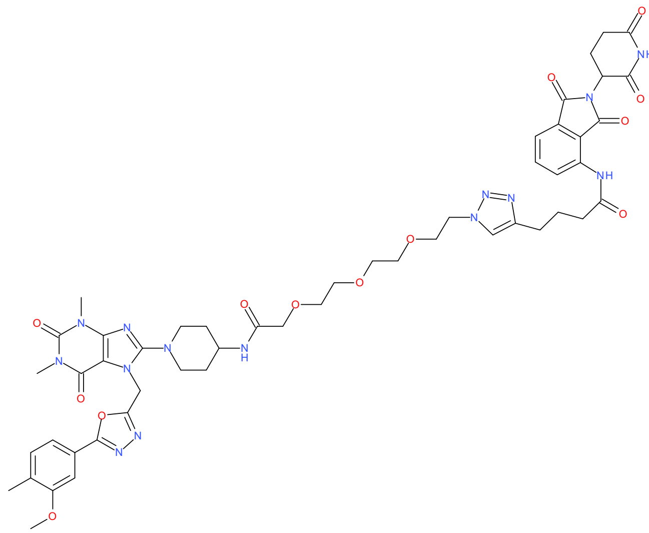

**N-[2-(2,6-Dioxo-3-piperidyl)-1,3-dioxo-isoindolin-4-yl]-4-[1-[2-[2-[2-[2-[[1-[7-[[5-(3-methoxy-4- methyl-phenyl)-1,3,4-oxadiazol-2-yl]methyl]-1,3-dimethyl-2,6-dioxo-purin-8-yl]-4-piperidyl] amino]-2-oxo-ethoxy]ethoxy]ethoxy]ethyl]triazol-4-yl]butanamide (17d)**

2-[2-[2-(2-Azidoethoxy)ethoxy]ethoxy]-*N*-[1-[7-[[5-(3-methoxy-4-methyl-phenyl)-1,3,4- oxadiazol-2-yl]methyl]-1,3-dimethyl-2,6-dioxo-purin-8-yl]-4-piperidyl]acetamide (**16b**) (0.0382 g, 0.0461 mmol) and sodium *L*-ascorbate (0.0233 g, 0.117 mmol) were dissolved in in *t*-BuOH:DCM (2:1, 2.25 mL). $CuSO_4 \cdot 5\ H_2O$ (0.0196 g, 0.0776 mmol) in water (0.75 mL) was added and the mixture stirred at room temperature for 30 minutes. Then, *N*-[2-(2,6-Dioxo-3-piperidyl)-1,3-dioxo-isoindolin-4-yl]hex-5-ynamide (**5b**) (0.0150 g, 0.0392 mmol) was added together with some more DCM, *t*-BuOH and a few mgs of sodium *L*-ascorbate. EtOAc was added, and the resulting organic phase was washed with ammonia solution (28%), brine, and water. The solvent was removed and the resulting product redissolved in DMF and purified by preparative HPLC (gradient of 10% to 60% acetonitrile in 0.1% TFA). Lyophilization of the pure fractions afforded the title compound (91.0%, 2.50 mg, 2.14 µmol, yield: 5.5%). MS (ESI$^+$) *m/z* 538 $[M+2H]^{2+}$. HRMS: *m/z* calculated 1063.4386, found 1063.4380 for $[M+H]^+$.

Preparation of **DDD11**

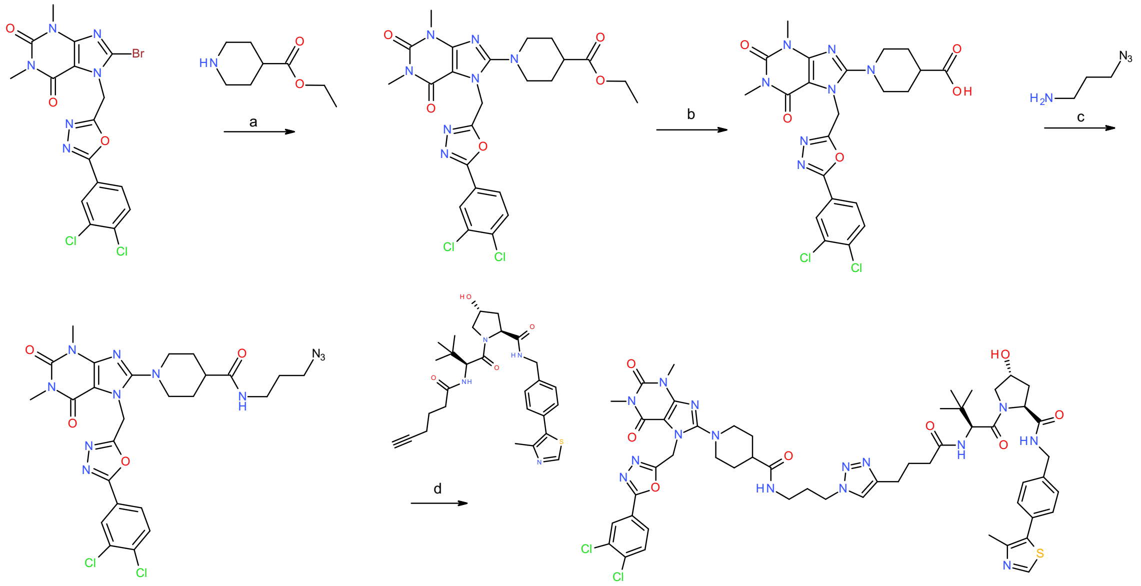

**Scheme S11.** Reagents and conditions: a) $K_2CO_3$, DMF, 70 °C, overnight; b) Bis(tributyltin) oxide, toluene, 115 °C, 12 h; c) HATU, DIEA, DMF, rt, 1 h; d) sodium *L*-ascorbate, $CuSO_4 \cdot 5 H_2O$, DCM:*t*-BuOH (1:2) mixture, water, rt, 90 min.

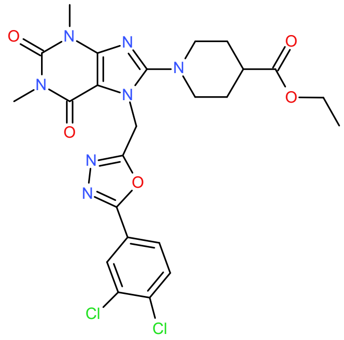

**Ethyl 1-[7-[[5-(3,4-dichlorophenyl)-1,3,4-oxadiazol-2-yl]methyl]-1,3-dimethyl-2,6-dioxo-purin-8-yl]piperidine-4-carboxylate (18a)**

Compound **1a** (0.270 g, 0.555 mmol), ethyl piperidine-4-carboxylate (0.109 g, 0.680 mmol) and $K_2CO_3$ (0.154 g, 1.10 mmol) were dissolved in DMF (5 mL). The mixture was stirred at 70 °C overnight. The mixture was diluted with EtOAc and water and the layers separated. The organic phase was washed with brine, leading to the formation of a precipitate. The precipitate was collected by filtration, and the organic phase dried over $MgSO_4$ and concentrated in vacuo. The organic phase and the filtrate were pooled and purified using flash column chromatography

(MeOH in DCM, 0% to 10%) to afford the title compound (92.0%, 0.174 g, 0.286 mmol, yield: 51%). MS (ESI$^+$) *m/z* 562 [M+H]$^+$.

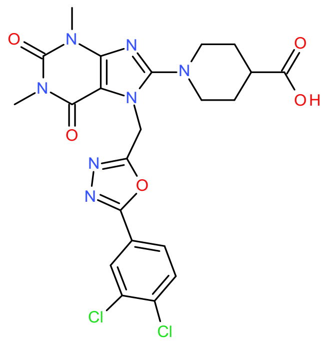

**1-[7-[[5-(3,4-Dichlorophenyl)-1,3,4-oxadiazol-2-yl]methyl]-1,3-dimethyl-2,6-dioxo-purin-8-yl]piperidine-4-carboxylic acid (19a)**

Compound **18a** (0.170 g, 0.278 mmol) and bis(tributyltin) oxide (0.567 mL, 1.05 mmol) were dissolved in toluene (2 mL) and heated to 115 °C in a closed vessel for 12 h. The toluene was removed in vacuo and the crude material was purification by flash column chromatography on silica gel (MeOH in DCM, 0% to 15%) to afford the title compound (91.0%, 0.0560 g, 0.0954 mmol, yield: 34%). MS (ESI$^-$) *m/z* 532 [M-H]$^-$.

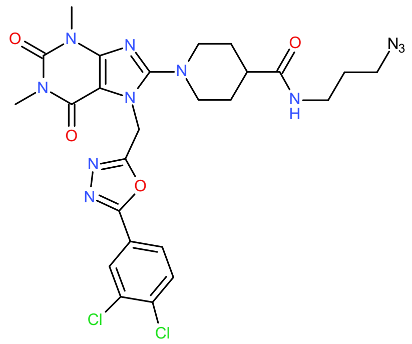

***N*-(3-Azidopropyl)-1-[7-[[5-(3,4-dichlorophenyl)-1,3,4-oxadiazol-2-yl]methyl]-1,3-dimethyl-2,6- dioxo-purin-8-yl]piperidine-4-carboxamide (20a)**

Compound **19a** (0.0560 g, 0.0954 mmol), DIEA (0.0490 mL, 0.283 mmol) and 3-azidopropan-1-amine (0.0116 g, 0.108 mmol) were dissolved in DMF (1 mL) and stirred for 10 minutes at rt. HATU (0.0435 g, 0.114 mmol) was added and the reaction was left at rt for 1 h. The mixture was then diluted with EtOAc and washed with saturated aqueous $NaHCO_3$ and brine, dried over $MgSO_4$ and concentrated in vacuo to afford the title compound (73.0%, 0.0700 g, 0.0829 mmol, yield: 87%). The crude material was used in the next step without further purification. MS (ESI$^+$) *m/z* 616 [M+H]$^+$.

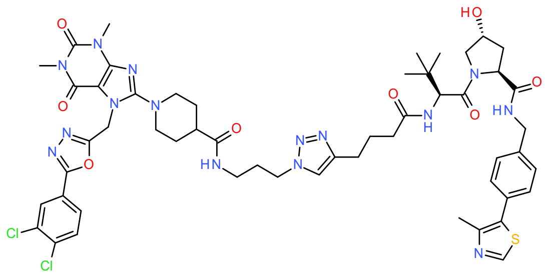

**1-[7-[[5-(3,4-Dichlorophenyl)-1,3,4-oxadiazol-2-yl]methyl]-1,3-dimethyl-2,6-dioxo-purin-8-yl]-*N*-[3-[4-[4-[[(*1S*)-1-[(*2S*,*4R*)-4-hydroxy-2-[[4-(4-methylthiazol-5-yl)phenyl]methylcarbamoyl] pyrrolidine-1-carbonyl]-2,2-dimethyl-propyl]amino]-4-oxo-butyl]triazol-1-yl]propyl]piperidine-4-carboxamide (21a)**

Compound **20a** (0.0233 g, 0.0275 mmol), compound **5a** (73.0%, 0.0165 g, 0.0229 mmol) and sodium *L*-ascorbate (0.0136 g, 0.0681 mmol) were dissolved in *t*-BuOH:DCM (2:1, 1.5 mL). $CuSO_4 \cdot 5\ H_2O$ (0.0114 g, 0.0454 mmol) in water (0.5 mL) was added and the mixture stirred at room temperature for 90 minutes. The mixture was diluted with ammonia solution (28%) and extracted with DCM. The organic phase was concentrated and redissolved in a small volume of DMF and purified by preparative HPLC (gradient of 10% to 60% acetonitrile in 0.1% TFA). Lyophilization of the pure fractions afforded the title compound (95.0%, 4.20 mg, 3.50 µmol, yield: 15%). MS ($ESI^+$) *m/z* 571 $[M+2H]^{2+}$. HRMS: *m/z* calculated 1140.4160, found 1140.4180 for $[M+H]^+$.

Preparation of **DDD12**

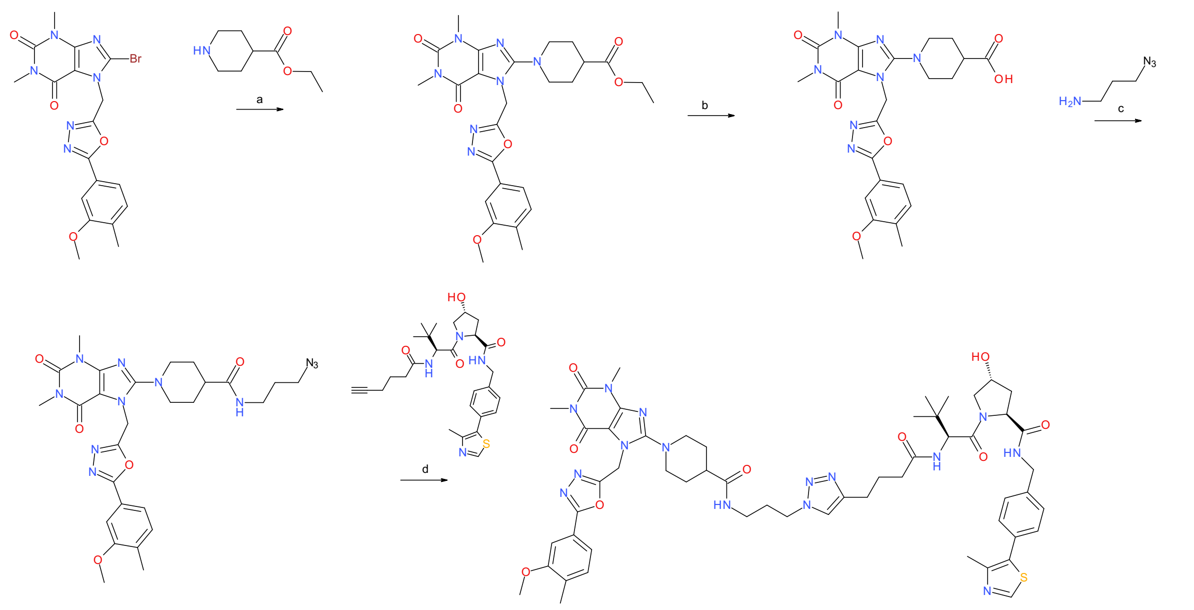

**Scheme S12.** Reagents and conditions: a) $K_2CO_3$, DMF, 70 °C, overnight; b) Bis(tributyltin) oxide, toluene, 115 °C, 12 h; c) HATU, DIEA, DMF, rt, 1 h; d) sodium *L*-ascorbate, $CuSO_4$·5 $H_2O$, DCM:*t*-BuOH (1:2) mixture, water, rt, 1.5 h.

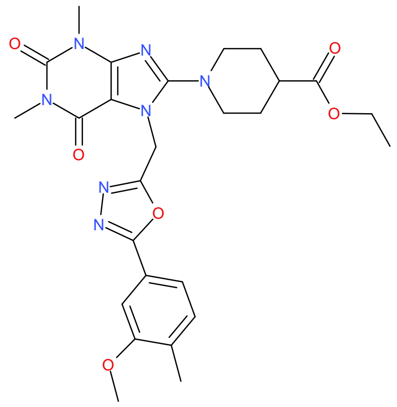

**Ethyl 1-[7-[[5-(3-methoxy-4-methyl-phenyl)-1,3,4-oxadiazol-2-yl]methyl]-1,3-dimethyl-2,6-dioxo-purin-8-yl]piperidine-4-carboxylate (18b)**

Compound **1b** (0.275 g, 0.596 mmol), ethyl piperidine-4-carboxylate (0.117 g, 0.731 mmol) and $K_2CO_3$ (0.165 g, 1.18 mmol) were dissolved in DMF (3 mL). The mixture was stirred at 70 °C overnight. The mixture was diluted with EtOAc and water, leading to the formation of a precipitate. The precipitate was filtered off, and the organic phase washed with brine, dried over $MgSO_4$ and concentrated in vacuo. The organic phase and the filtrate were pooled and purified by flash column chromatography (MeOH in DCM, 0% to 10%), yielding the title compound (94.0%, 0.240 g, 0.421 mmol, yield: 71%). MS (ESI$^+$) *m/z* 538 $[M+H]^+$.

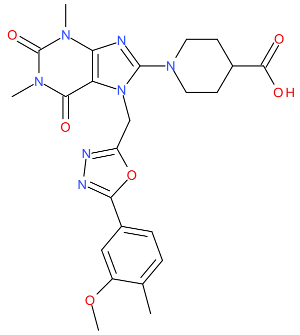

**1-[7-[[5-(3-Methoxy-4-methyl-phenyl)-1,3,4-oxadiazol-2-yl]methyl]-1,3-dimethyl-2,6-dioxo-purin-8-yl]piperidine-4-carboxylic acid (19b)**

Compound **18b** (0.200 g, 0.350 mmol) and bis(tributyltin) oxide (0.713 mL, 1.33 mmol) were dissolved in toluene (2 mL) and heated to 115 °C in a closed vessel for 12 h. Toluene was removed in vacuo and the material was purification by flash column chromatography on silica gel (MeOH in DCM, 0% to 10%)to afford the title compound (90.0%, 0.196 g, 0.346 mmol, yield: 99%). MS (ESI$^-$) *m/z* 508 [M-H]$^-$.

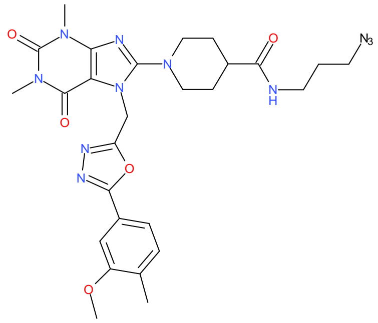

***N*-(3-Azidopropyl)-1-[7-[[5-(3-methoxy-4-methyl-phenyl)-1,3,4-oxadiazol-2-yl]methyl]-1,3-dimethyl-2,6-dioxo-purin-8-yl]piperidine-4-carboxamide (20b)**

Compound **19b** (0.100 g, 0.177 mmol), DIEA (0.0907 mL, 0.525 mmol) and 3-azidopropan-1-amine (0.0214 g, 0.201 mmol) were dissolved in DMF (2 mL) and left to stir for 10 minutes at rt. HATU (0.0806 g, 0.212 mmol) was added and the reaction was left for 1 h. The mixture was diluted with EtOAc and saturated aqueous $NaHCO_3$, resulting in the formation of a precipitate. The precipitate was collected by filtration and dried to afford the title compound (73.0%, 0.0500 g, 0.0617 mmol, yield: 35%), which was used in the next step without further purification. MS (ESI$^+$) *m/z* 592 [M+H]$^+$.

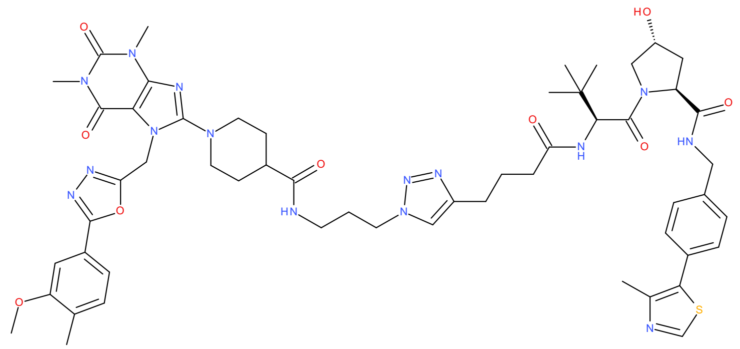

**1-[7-[[5-(3-Methoxy-4-methyl-phenyl)-1,3,4-oxadiazol-2-yl]methyl]-1,3-dimethyl-2,6-dioxo-purin-8-yl]-N-[3-[4-[4-[[(*1S*)-1-[(*2S*,*4R*)-4-hydroxy-2-[[4-(4-methylthiazol-5-yl)phenyl]methylcarbamoyl]pyrrolidine-1-carbonyl]-2,2-dimethyl-propyl]amino]-4-oxo-butyl]triazol-1-yl]propyl]piperidine-4-carboxamide (21b)**

Compound **20b** (0.0223 g, 0.0275 mmol), compound **5a** (0.0165 g, 0.0229 mmol) and sodium *L*-ascorbate (0.0136 g, 0.0681 mmol) were dissolved in *t*-BuOH:DCM (2:1, 1.5 mL). $CuSO_4 \cdot 5\ H_2O$ (0.0114 g, 0.0454 mmol) in water (0.5 mL) was added and the mixture stirred at room temperature for 1.5 h. The mixture was diluted with ammonia solution (28%) and extracted with DCM. The organic phase was dried in vacuo and redissolved in a small volume of DMF, filtered, and purified by preparative HPLC (gradient of 10% to 60% acetonitrile in 0.1% TFA). Lyophilization of the pure fractions yielded the title compound (94.0%, 2.20 mg, 1.85 µmol, yield: 8.1%). MS ($ESI^+$) *m/z* 559 $[M+2H]^{2+}$. HRMS: *m/z* calculated 1116.5201, found 1116.5222 for $[M+H]^+$.

<u>Preparation of **DDD13**</u>

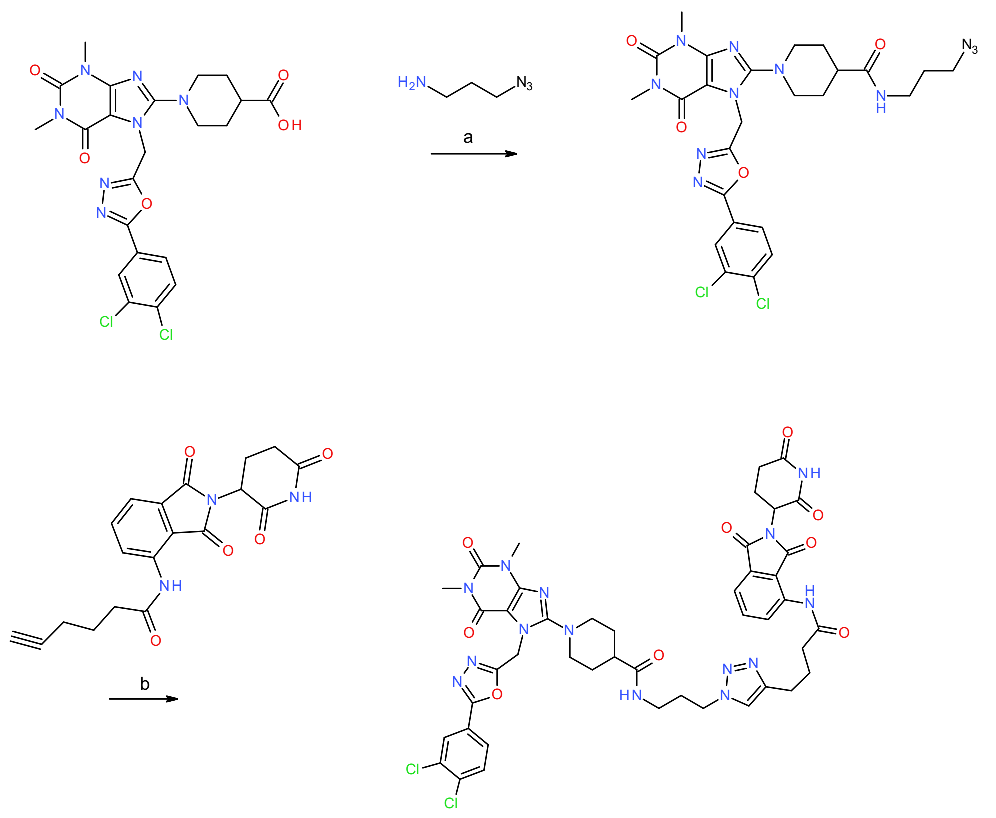

**Scheme S13.** Reagents and conditions: a) HATU, DIEA, DMF, rt, 1 h; b) sodium *L*-ascorbate, $CuSO_4 \cdot 5\ H_2O$, DCM:*t*-BuOH (1:2) mixture, water, rt, 1.5 h.

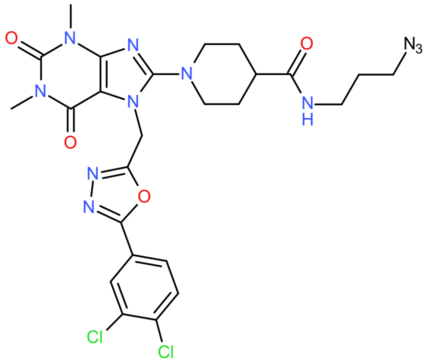

**_N_-(3-Azidopropyl)-1-[7-[[5-(3,4-dichlorophenyl)-1,3,4-oxadiazol-2-yl]methyl]-1,3-dimethyl-2,6- dioxo-purin-8-yl]piperidine-4-carboxamide (20a)**

1-[7-[[5-(3,4-Dichlorophenyl)-1,3,4-oxadiazol-2-yl]methyl]-1,3-dimethyl-2,6-dioxo-purin-8-yl]piperidine-4-carboxylic acid (**19a**) (0.0560 g, 0.0954 mmol), DIEA (0.0490 mL, 0.283 mmol) and 3-azidopropan-1-amine (0.0116 g, 0.108 mmol) were dissolved in DMF (1 mL) and left to stir for 10 minutes at rt. HATU (0.0435 g, 0.114 mmol) was added and the reaction was stirred at rt for 1

h. EtOAc was added and the and the mixture was washed with saturated aqueous $NaHCO_3$ and brine, dried over $MgSO_4$ and concentrated in vacuo, yielding the title compound (73.0%, 0.0700 g, 0.0829 mmol, yield: 87%). The crude material was used in the next step without further purification. MS ($ESI^+$) *m/z* 616 $[M+H]^+$.

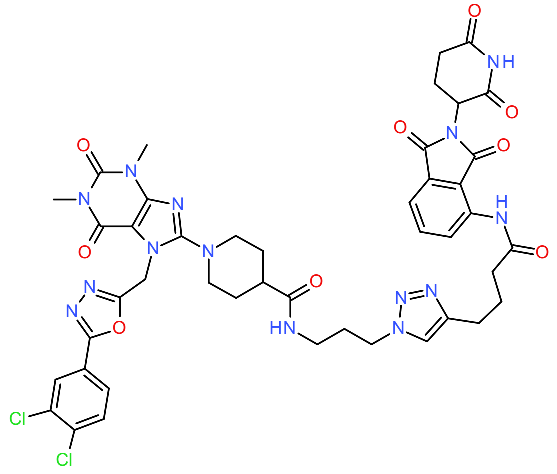

**1-[7-[[5-(3,4-Dichlorophenyl)-1,3,4-oxadiazol-2-yl]methyl]-1,3-dimethyl-2,6-dioxo-purin-8-yl]-*N*- [3-[4-[4-[[2-(2,6-dioxo-3-piperidyl)-1,3-dioxo-isoindolin-4-yl]amino]-4-oxo-butyl]triazol-1-yl] propyl]piperidine-4-carboxamide (DDD13)**

*N*-(3-Azidopropyl)-1-[7-[[5-(3,4-dichlorophenyl)-1,3,4-oxadiazol-2-yl]methyl]-1,3-dimethyl-2,6-dioxo-purin-8-yl]piperidine-4-carboxamide (**20a**) (0.0233 g, 0.0275 mmol), *N*-[2-(2,6-Dioxo-3-piperidyl)-1,3-dioxo-isoindolin-4-yl]hex-5-ynamide (**5b**) (0.00868 g, 0.0229 mmol) and sodium *L*-ascorbate (0.0136 g, 0.0681 mmol) were dissolved in *t*-BuOH:DCM (2:1, 1.5 mL). $CuSO_4 \cdot 5\ H_2O$ (0.0114 g, 0.0454 mmol) in water (0.5 mL) was added and the mixture stirred at room temperature for 30 minutes. After that, LC-MS indicated that unreacted hexynamide was still present, so an additional few mgs of azide in DCM/*t*-BuOH solution was added, and the mixture left for 1 h. Ammonia solution (28%) and EtOAc were added to the reaction mixture. The phases were separated, and the organic phase was dried over $MgSO_4$ and concentrated in vacuo. The crude product was dissolved in DMF and purified by preparative HPLC (gradient of 10% to 60% can in 0.1% TFA). Lyophilization of the pure fractions afforded the title compound (83.0%, 2.70 mg, 2.28 mmol, yield: 9.9%). MS ($ESI^+$) *m/z* 983 $[M+H]^+$. HRMS: *m/z* calculated 983.2871, found 983.2869 for $[M+H]^+$. $^1H$ NMR (400 MHz, DMSO-$d_6$): δ 11.15 (s, 1H), 9.73 (s, 1H), 8.45 (d, J=8.4 Hz, 1H), 8.15 (d, J=1.9 Hz, 1H), 7.87-7.97 (m, 4H), 7.83 (dd, J=8.0 Hz, 7.5 Hz, 1H), 7.60 (d, J=7.3 Hz, 0.52 Hz, 1H), 5.67 (s, 2H), 36 5.15 (dd, J=13 Hz, 5.3 Hz, 1H), 4.30 (t, J=7.0 Hz, 2H), 3.56-3.64 (m, 2H), 3.41 (s, 3H), 3.16 (s, 3H), 2.93-3.08 (m, 5H), 2.54-2.72 (m, 7H), 2.02-2.10 (m, 1H), 1.90-2.00 (m, 4H), 1.63-1.78 (m, 4H).

<u>Preparation of **DDD14**</u>

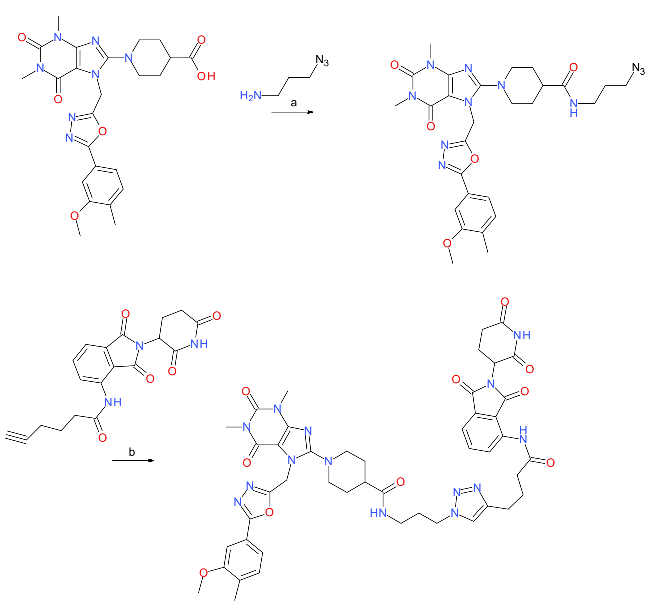

**Scheme S14.** Reagents and conditions: a) HATU, DIEA, DMF, rt, 1 h; b) sodium *L*-ascorbate, $CuSO_4 \cdot 5\ H_2O$, DCM:*t*-BuOH (1:2) mixture, water, rt, 1.5 h.

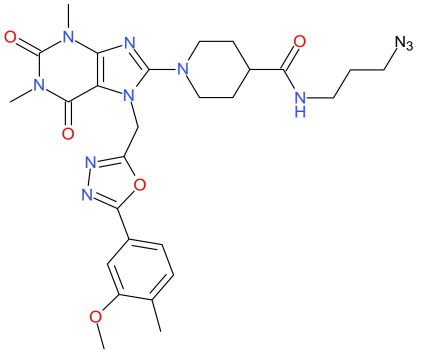

**_N_-(3-Azidopropyl)-1-[7-[[5-(3-methoxy-4-methyl-phenyl)-1,3,4-oxadiazol-2-yl]methyl]-1,3-dimethyl-2,6-dioxo-purin-8-yl]piperidine-4-carboxamide (20b)**

1-[7-[[5-(3-Methoxy-4-methyl-phenyl)-1,3,4-oxadiazol-2-yl]methyl]-1,3-dimethyl-2,6-dioxo-purin-8-yl]piperidine-4-carboxylic acid (**19b**) (0.100 g, 0.177 mmol), DIEA (0.0907 mL, 0.525 mmol) and 3-azidopropan-1-amine (0.0214 g, 0.201 mmol) were dissolved in DMF (2 mL) and stirred for 10 minutes at rt. HATU (0.0806 g, 0.212 mmol) was added and the reaction was left at rt for 1 h. The mixture was diluted with EtOAc and saturated aqueous $NaHCO_3$ solution, resulting in the formation of a precipitate. The precipitate was collected by filtration and dried to afford the title compound (73.0%, 0.0500 g, 0.0617 mmol, yield: 35%) which was used in the next step without further purification. MS (ESI$^+$) *m/z* 592 [M+H]$^+$.

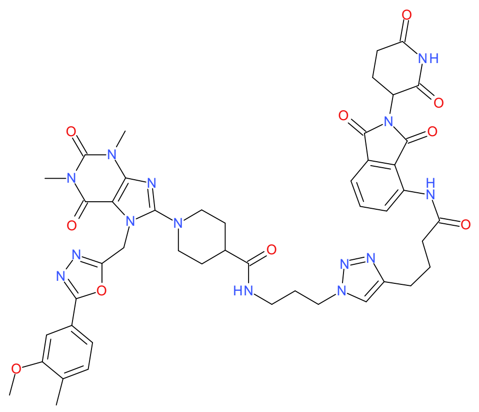

**1-[7-[[5-(3-Methoxy-4-methyl-phenyl)-1,3,4-oxadiazol-2-yl]methyl]-1,3-dimethyl-2,6-dioxopurin-8-yl]-*N*-[3-[4-[4-[[2-(2,6-dioxo-3-piperidyl)-1,3-dioxo-isoindolin-4-yl]amino]-4-oxobutyl]triazol-1-yl]propyl]piperidine-4-carboxamide (DDD14)**

*N*-(3-Azidopropyl)-1-[7-[[5-(3-methoxy-4-methyl-phenyl)-1,3,4-oxadiazol-2-yl]methyl]-1,3-dimethyl-2,6-dioxo-purin-8-yl]piperidine-4-carboxamide (**20b**) (0.0223 g, 0.0275 mmol), *N*-[2-(2,6-Dioxo-3-piperidyl)-1,3-dioxo-isoindolin-4-yl]hex-5-ynamide (**5b**) (0.00868 g, 0.0229 mmol) and sodium *L*-ascorbate (0.0136 g, 0.0681 mmol) were dissolved in in *t*-BuOH:DCM (2:1, 1.5 mL). $CuSO_4 \cdot 5\ H_2O$ (0.0114 g, 0.0454 mmol) in water (0.5 mL) was added and the mixture stirred at room temperature for 1.5 h. Ammonia solution was addedand the mixture was extracted with DCM and the phases separated. The organic phase was removed in vacuo. The crude product was dissolved in DMF and purified by preparative HPLC (gradient of 10% to 60% ACN in 0.1% TFA). Lyophilization of the pure fractions afforded the title compound (81.0%, 3.30 mg, 2.79 µmol, yield: 12%). MS (ESI$^+$) *m/z* 959 [M+H]$^+$. HRMS: *m/z* calculated 959.3912, found 959.3910 for [M+H]$^+$. $^1$H NMR (400 MHz, DMSO-$d_6$): δ 11.15 (s, 1H), 9.72 (s, 1H), 8.45 (d, J=8.4 Hz, 1H), 7.89-97 (m, 2H), 7.83 (dd, J=8.1 Hz, 7.5 Hz, 1H), 7.62 (d, J=7.3 Hz, 0.45 Hz, 1H), 7.35-7.48 (m, 3H), 5.66 (s, 2H), 5.15 (dd, J=13 Hz, 5.4 Hz, 1H), 4.30 (t, J=7.0 Hz, 2H), 3.87 (s, 3H), 3.57-3.64 (m, 2H), 3.41 (s, 3H), 3.16 (s, 3H), 2.93-3.12 (m, 5H), 2.53-2.72 (m, 7H), 2.23 (s, 3H), 2.03-2.10 (m, 1H), 1.89-1.99 (m, 4H), 1.61-1.78 (m, 4H).

<u>Preparation of **DDD15**</u>

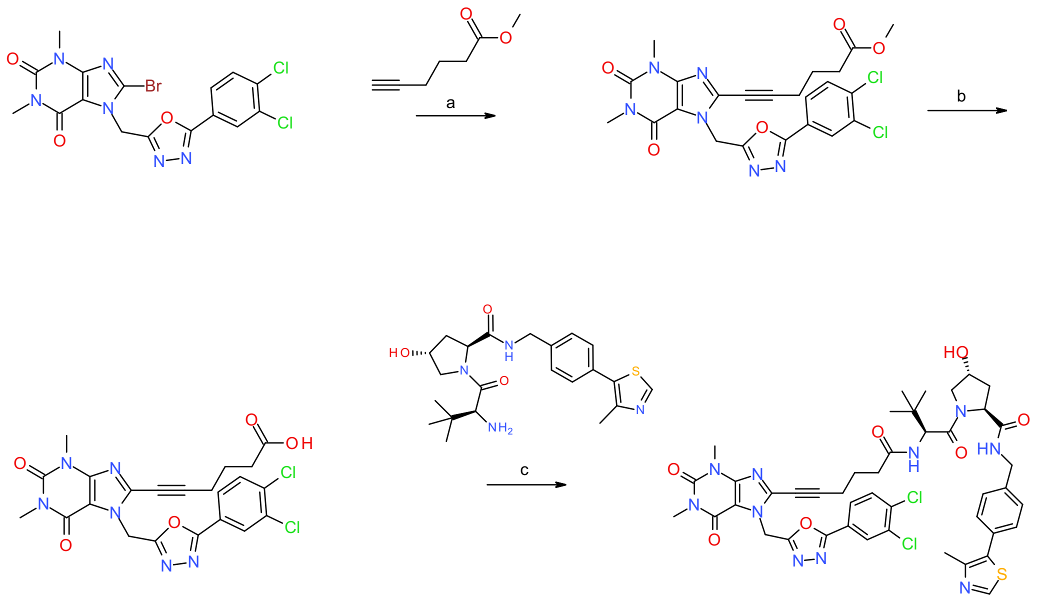

**Scheme S15.** Reagents and conditions: a) tetrakis(triphenylphosphine)palladium(0), CuI, TEA, DMF, 50 °C, overnight; b) HATU, DIEA, DMF, rt, 3 h; b) LiOH·$H_2O$, THF:water 3:7, rt, 2.5 h; c) HATU, DIEA, DMF, rt, over the weekend.

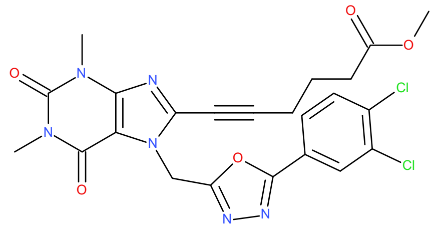

**Methyl 6-[7-[[5-(3,4-dichlorophenyl)-1,3,4-oxadiazol-2-yl]methyl]-1,3-dimethyl-2,6-dioxo-purin-8-yl]hex-5-ynoate (22a)**

Compound **1a** (0.200 g, 0.411 mmol), compound **22c** (0.457 g, 3.62 mmol) and TEA (2.29 mL, 16.3 mmol) were dissolved in DMF (8 mL) under a nitrogen atmosphere for a few minutes. Then, tetrakis (triphenylphosphine)palladium(0) (0.0333 g, 0.0285 mmol) and CuI (0.0533 g, 0.274 mmol) were added, and the reaction mixture was heated under a nitrogen atmosphere at 50 °C overnight. The mixture was then cooled, filtered and washed with EtOAc. The filtrate was washed with brine, concentrated and purified by flash column chromatography on silica gel, using MeOH in DCM (0% to 100%) to afford the title compound (0.0150 g, 0.282 mmol, yield: 69%). MS (ESI$^+$) *m/z* 531 $[M+H]^+$.

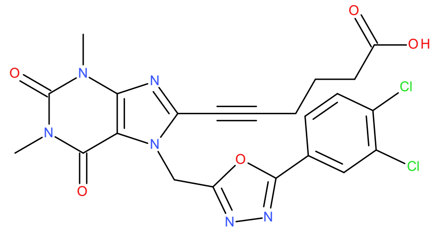

**6-[7-[[5-(3,4-Dichlorophenyl)-1,3,4-oxadiazol-2-yl]methyl]-1,3-dimethyl-2,6-dioxo-purin-8-yl] hex-5-ynoic acid (23a)**

Compound (**22a**) (0.150 g, 0.282 mmol) was dissolved in THF:water (3:7, 10 mL). LiOH·$H_2O$ (0.0355 g, 0.830 mmol) was added and the mixture stirred at rt for 2.5 h. Then THF was removed in vacuo and the remaining water acidified to pH=2 using 1 M HCl. The product was extracted with DCM (4 × 25 mL), the organic phase was washed with brine and dried over $MgSO_4$. The solvent was removed in vacuo and the product dried to afford the title compound (70.0%, 0.0580 g, 0.0785 mmol, yield: 28%), which was used in the next step without further purification. MS ($ESI^+$) *m/z* 517 $[M+H]^+$.

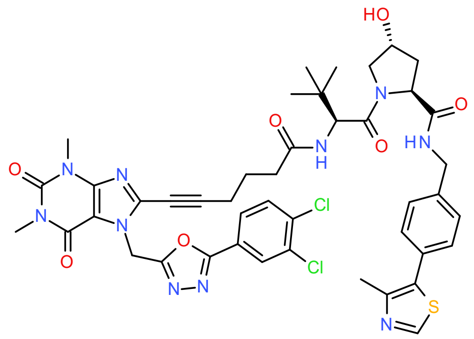

**(*2S*,*4R*)-1-[(*2S*)-2-[6-[7-[[5-(3,4-Dichlorophenyl)-1,3,4-oxadiazol-2-yl]methyl]-1,3-dimethyl-2,6- dioxo-purin-8-yl]hex-5-ynoylamino]-3,3-dimethyl-butanoyl]-4-hydroxy-*N*-[[4-(4-methylthiazol-5-yl)phenyl]methyl]pyrrolidine-2-carboxamide (DDD15)**

Compound **23a** (0.0290 g, 0.0392 mmol), DIEA (0.0202 mL, 0.117 mmol) and HATU (0.0179 g, 0.0471 mmol) were dissolved in DMF (0.5 mL) and stirred for 10 minutes at rt. Compound Compound **2a** (0.0169 g, 0.0392 mmol) was added and the reaction was left over the weekend. An additional 9 mg of HATU, 10 µL of DIEA and 8 mg of compound **2a** were added and the reaction left at rt for 5 h. The reaction mixture was diluted with DCM and washed with brine (2x) and water. The solvent was then removed and the crude product redissolved in acetonitrile and purified by preparative HPLC (Xbridge, gradient of 20% to 50% acetonitrile in 50 mM ammonium bicarbonate). Lyophilizing the pure fractions yielded the title compound (>99%, 11.5 mg, 12.5 µmol, yield: 32%). MS ($ESI^+$) *m/z* 465 $[M+2H]^{2+}$. HRMS: *m/z* calculated 929.2727, found 929.2671 for $[M+H]^+$. $^1H$ NMR (400 MHz, DMSO-$d_6$): δ 8.99 (s, 1H), 8.56 (t, J=6.1 Hz, 1H), 8.15 (d, J=1.9 Hz, 1H), 7.97 (d, J=9.3 Hz, 1H), 7.93 (dd, J=8.4, 1.9 Hz, 1H), 7.89 (d, J=8.4 Hz, 1H), 7.36-7.44 (m, 4H), 5.96 (s, 2H), 5.12 (d, J=3.5 Hz, 1H), 4.53 (d, J=9.4 Hz, 1H), 4.40-4.48 (m, 2H), 4.32-4.38

(m, 1H), 4.21 (dd, J=16 Hz, 5.4 Hz, 1H), 3.60-3.69 (m, 2H), 3.44 (s, 3H), 3.21 (s, 3H), 2.55 (t, J=7.0 Hz, 2H), 2.45 (s, 3H), 2.25-2.42 (m, 2H), 1.99-2.07 (m, 1H), 1.88-1.94 (m, 1H), 1.79 (p, J=6.6 Hz, 2H), 0.91 (s, 9H).

Preparation of **DDD16**

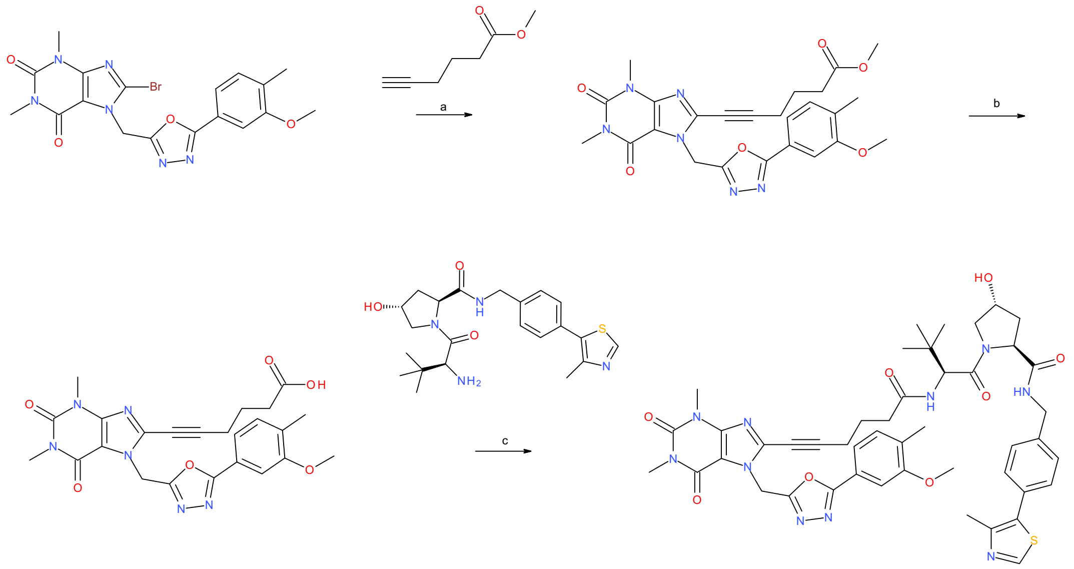

**Scheme S16.** Reagents and conditions: a) Tetrakis(triphenylphosphine)palladium(0), CuI, TEA, DMF, 50 °C, overnight; b) HATU, DIEA, DMF, rt, 3 h; b) LiOH·$H_2O$, THF:water 1:1, rt, 2.5 h; c) HATU, DIEA, DMF, rt, over the weekend.

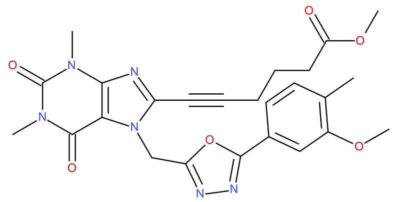

**Methyl 6-[7-[[5-(3-methoxy-4-methyl-phenyl)-1,3,4-oxadiazol-2-yl]methyl]-1,3-dimethyl-2,6-dioxo-purin-8-yl]hex-5-ynoate (22b)**

Compound **1b** (0.150 g, 0.315 mmol), compound **22c** (0.233 g, 1.85 mmol) and TEA (1.76 mL, 12.5 mmol) were dissolved in DMF (8 mL) and the mixture conditioned under nitrogen atmosphere for a few minutes. Tetrakis(triphenylphosphine)palladium(0) (0.0255 g, 0.0219 mmol) and CuI (0.0409 g, 0.210 mmol) were added and the reaction heated under a nitrogen atmosphere at 50 °C overnight. The mixture was then diluted with EtOAc:water, and the organic phase washed with brine and concentrated in vacuo. The crude product was purified twice by flash column chromatography on silica gel, using MeOH in DCM (0% to 100%). Finally, the material was triturated with EtOAc to remove impurities and dried to afford the title compound (0.0600 g, 0.118 mmol, yield: 38%). MS (ESI$^+$) *m/z* 507 $[M+H]^+$.

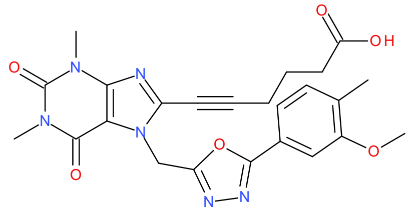

**6-[7-[[5-(3-Methoxy-4-methyl-phenyl)-1,3,4-oxadiazol-2-yl]methyl]-1,3-dimethyl-2,6-dioxo-purin-8-yl]hex-5-ynoic acid (23b)**

Compound **22b** (0.0590 g, 0.116 mmol) was dissolved in THF:water (1:1, 2 mL). $LiOH \cdot H_2O$ (0.0147 g, 0.342 mmol) was added and the mixture stirred at rt for 2.5 h. THF was removed in vacuo and the remaining water was acidified to pH=2 using 1 M HCl. The product was extracted with DCM (4x). The combined organic phases were washed with brine and dried over $MgSO_4$. The solvent was removed in vacuo and the material dried to give the title compound (71.0%, 0.0240 g, 0.0346 mmol, yield: 30%), which was used in the next step without further purification. MS (ESI$^-$) *m/z* 491 $[M-H]^-$.

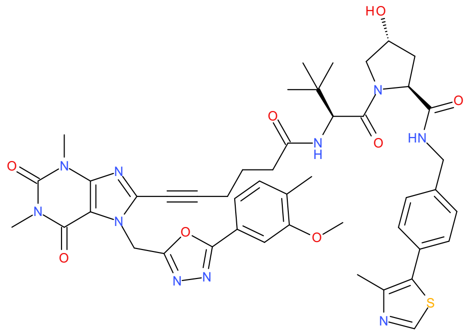

**(*2S*,*4R*)-4-hydroxy-1-[(*2S*)-2-[6-[7-[[5-(3-Methoxy-4-methyl-phenyl)-1,3,4-oxadiazol-2-yl]methyl]-1,3-dimethyl-2,6-dioxo-purin-8-yl]hex-5-ynoylamino]-3,3-dimethyl-butanoyl]-*N*-[[4- (4-methylthiazol-5-yl)phenyl]methyl]pyrrolidine-2-carboxamide (DDD16)**

Compound **23b** (0.0200 g, 0.0288 mmol), DIEA (0.0209 mL, 0.121 mmol) and compound **2a** (0.0175 g, 0.0406 mmol) were dissolved in DMF (0.5 mL) and stirred for 10 minutes at rt. HATU (0.0185 g, 0.0487 mmol) was added and the reaction was left over the weekend. The mixture was diluted with water and EtOAc and the phases separated. The organic phase was washed with saturated aqueous $NaHCO_3$, brine and water. The solvent was then removed and the crude product redissolved in acetonitrile and purified by preparative HPLC (gradient of 10% to 60% acetonitrile in 50 mM ammonium bicarbonate). Lyophilizing the pure fractions yielded the title compound (>99%, 8.20 mg, 9.06 µmol, yield: 31%). MS (ESI+) *m/z* 453 $[M+2H]^{2+}$. HRMS: *m/z* calculated 905.3768, found 905.3770 for $[M+H]^+$. $^1H$ NMR (400 MHz, DMSO-$d_6$): δ 8.98 (s, 1H), 8.56 (t, J=6.0 Hz, 1H), 7.98 (d, J=9.4 Hz, 1H), 7.35-7.47 (m, 7H), 5.94 (s, 2H), 5.12 (d, J=3.6 Hz, 1H), 4.53 (d, J=9.4 Hz, 1H), 4.40-4.47 (m, 2H), 4.32-4.37 (m, 1H), 4.22 (dd, J=16 Hz, 5.4 Hz, 1H), 3.89 (s, 3H), 3.60-3.68 (m, 2H), 3.43 (s, 3H), 3.20 (s, 3H), 2.55 (t, J=9.4 Hz, 2H), 2.44 (s, 3H), 2.25-2.43 (m, 2H), 2.23 (s, 3H), 1.99-2.07 (m, 1H), 1.86-1.94 (m, 1H), 1.79 (p, J=6.9 Hz, 2H), 0.92 (s, 9H).

<u>Preparation of **DDD17**</u>

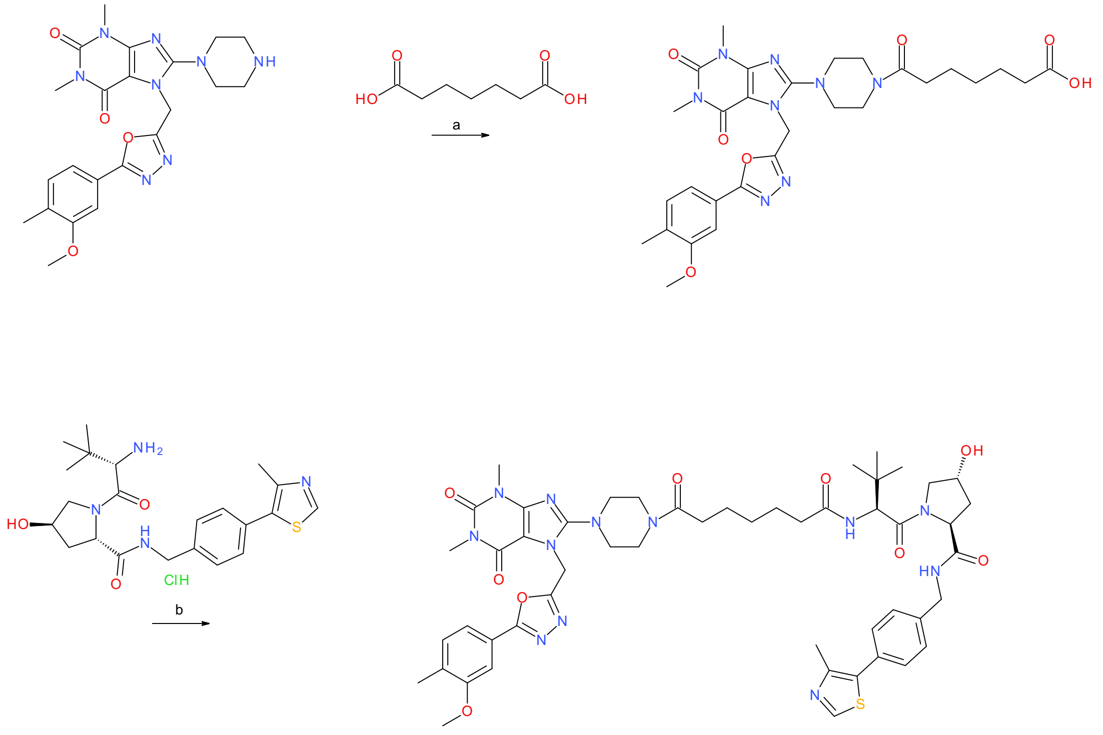

**Scheme S17.** Reagents and conditions: a) HATU, DIEA, DMF, rt, overnight; b) HATU, DIEA, DMF, rt, overnight.

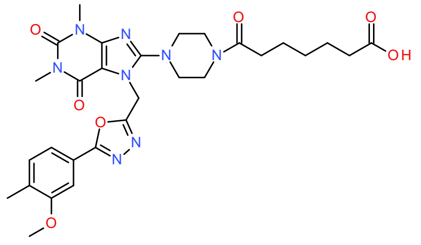

**7-[4-[7-[[5-(3-methoxy-4-methyl-phenyl)-1,3,4-oxadiazol-2-yl]methyl]-1,3-dimethyl-2,6-dioxo-purin-8-yl]piperazin-1-yl]-7-oxo-heptanoic acid**

7-[[5-(3-Methoxy-4-methyl-phenyl)-1,3,4-oxadiazol-2-yl]methyl]-1,3-dimethyl-8-piperazin-1-yl-purine-2,6-dione (93.0 %, 0.120 g, 0.239 mmol), heptanedioic acid (0.0383 g, 0.239 mmol) and DIEA (0.0837 mL, 0.489 mmol) were dissolved in DMF (1.5 mL) and conditioned at rt for 10 min. HATU (0.109 g, 0.287 mmol) was added and the mixture left at rt for 2.5 h. The mixture was treated with saturated $NaHCO_3$ (10 mL) and organic impurities extracted with EtOAc (15 mL). The aqueous phase was then treated with 1 M HCl until pH=1. The aqueous phase was extracted with EtOAc (25 mL), The organic phase was washed with brine (25 mL) and dried over $MgSO_4$. The solvent was removed in vacuo and the product was dried to give the title compound (60.0 %, 0.0460 g, 0.0453 mmol, yield: 19%) which was used in the next step without further purification. MS ($ESI^-$) *m/z* 607 $[M-H]^-$.

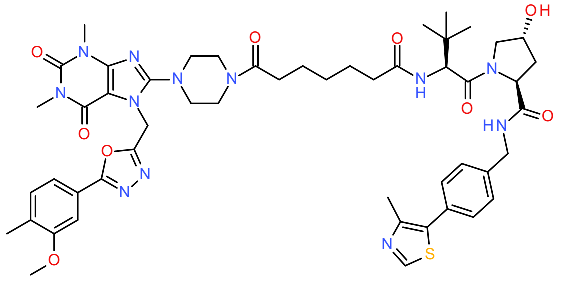

**(*2S*,*4R*)-4-hydroxy-1-[(*2S*)-2-[[7-[4-[7-[[5-(3-methoxy-4-methyl-phenyl)-1,3,4-oxadiazol-2-yl]methyl]-1,3-dimethyl-2,6-dioxo-purin-8-yl]piperazin-1-yl]-7-oxo-heptanoyl]amino]-3,3-dimethyl-butanoyl]-*N*-[[4-(4-methylthiazol-5-yl)phenyl]methyl]pyrrolidine-2-carboxamide (DDD17)**

7-[4-[7-[[5-(3-Methoxy-4-methyl-phenyl)-1,3,4-oxadiazol-2-yl]methyl]-1,3-dimethyl-2,6-dioxo-purin-8-yl]piperazin-1-yl]-7-oxo-heptanoic acid (60.0 %, 0.0460 g, 0.0453 mmol), (*2S*,*4R*)-1-[(*2S*)-2-amino-3,3-dimethyl-butanoyl]-4-hydroxy-*N*-[[4-(4-methylthiazol-5-yl)phenyl]methyl] pyrrolidine-2-carboxamide;hydrochloride (0.0212 g, 0.0453 mmol) and DIEA (0.0233 mL, 1.36 mmol) were dissolved in DMF (0.5 mL) and stirred at rt for 10 min. HATU (0.0207 g, 0.0544 mmol) was added and the mixture left overnight. The reaction mixture was diluted with acetonitrile and purified by preparative HPLC (gradient of 30% to 60% acetonitrile in 10 mM ammonium bicarbonate buffer). Lyophilization of the relevant fractions yielded the target compound (98.0%, 0.0154 g, 0.0148 mmol, yield: 33%). MS ($ESI^+$) *m/z* 511 $[M+2H]^{2+}$. HRMS *m/z* calculated 1021.4719, found 1021.4736 for $[M+H]^+$. $^1H$ NMR (400 MHz, DMSO-$d_6$) δ 8.97 (s, 1H), 8.55 (t, *J* = 6.0 Hz, 1H), 7.84 (d, *J* = 9.3 Hz, 1H), 7.47 – 7.35 (m, 7H), 5.71 (s, 2H), 5.11 (d, *J* = 3.5 Hz, 1H), 4.53 (d, *J* = 9.4 Hz, 1H), 4.47 – 4.39 (m, 2H), 4.37 – 4.31 (m, 1H), 4.21 (dd, *J* = 15.9, 5.4 Hz, 1H), 3.88 (s, 3H), 3.67 – 3.62 (m, 1H), 3.60 – 3.52 (m, 4H), 3.40 (s, 3H), 3.28 – 3.19 (m, 4H), 3.15 (s, 3H), 2.44 (s, 3H), 2.35 – 2.25 (m, 3H), 2.23 (s, 3H), 2.16 – 2.06 (m, 1H), 2.06 – 1.97 (m, 1H), 1.95 – 1.85 (m, 1H), 1.55 – 1.41 (m, 4H), 1.30 – 1.21 (m, 2H), 0.93 (s, 9H).

Preparation of **DDD18**

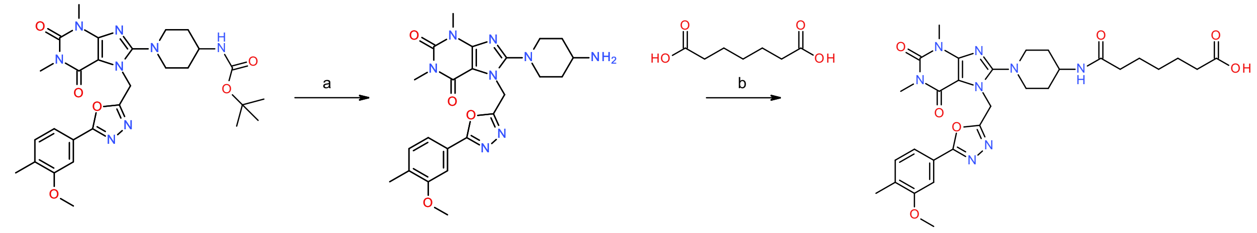

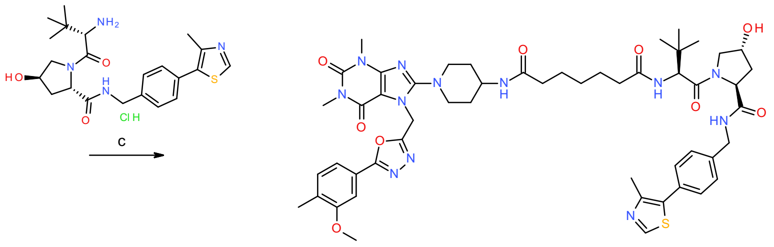

**Scheme S18.** Reagents and conditions: a) TFA in DCM (20%), rt; b) HATU, DIEA, DMF, rt, overnight; c) HATU, DIEA, DMF, rt, overnight.

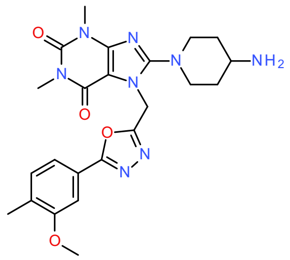

**8-(4-amino-1-piperidyl)-7-[[5-(3-methoxy-4-methyl-phenyl)-1,3,4-oxadiazol-2-yl]methyl]-1,3-dimethyl-purine-2,6-dione**

*tert*-Butyl *N*-[1-[7-[[5-(3-methoxy-4-methyl-phenyl)-1,3,4-oxadiazol-2-yl]methyl]-1,3-dimethyl-2,6-dioxo-purin-8-yl]-4-piperidyl]carbamate (95.0 %, 1.50 g, 2.45 mmol) was treated with 20% TFA in DCM solution (15 mL) and left to stir overnight. The mixture was neutralized with 2 M potassium carbonate solution and organics extracted with DCM (30 mL). The organic phase was dried over magnesium sulphate and concentrated in vacuo to afford the title compound (90.0 %, 0.867 g, 1.62 mmol, yield: 66%). MS (ESI$^+$) *m/z* 481 [M+H]$^+$.

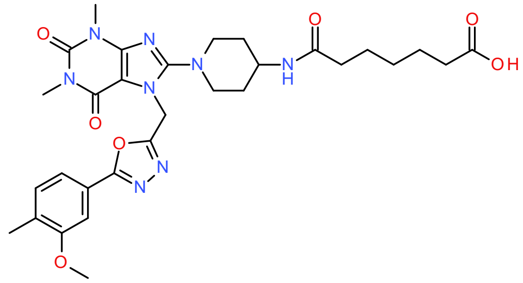

**7-[[1-[7-[[5-(3-Methoxy-4-methyl-phenyl)-1,3,4-oxadiazol-2-yl]methyl]-1,3-dimethyl-2,6-dioxo-purin-8-yl]-4-piperidyl]amino]-7-oxo-heptanoic acid**

7-[[5-(3-Methoxy-4-methyl-phenyl)-1,3,4-oxadiazol-2-yl]methyl]-1,3-dimethyl-8-piperazin-1-yl-purine-2,6-dione (93.0 %, 0.120 g, 0.239 mmol), heptanedioic acid (0.0383 g, 0.239 mmol) and DIEA (0.0837 mL, 0.489 mmol) were dissolved in DMF (1.5 mL) and stirred at RT for 10 min. HATU (0.109 g, 0.287 mmol) was added and the mixture left at RT for 2.5 h. The mixture was treated with saturated $NaHCO_3$ (10 mL) and organic impurities extracted with EtOAc (2x15 mL). The aqueous phase was then treated with 1 M HCl until pH=1, organics extracted with DCM (2x15 mL). The combined organic extracts were dried over $MgSO_4$, after which removal of the solvent in vacuo yielded the title compound (67.0%, 0.0590 g, 0.0635 mmol, yield: 26%), which was used in the next step without further purification. MS (ESI$^-$) *m/z* 621 [M-H]$^-$.

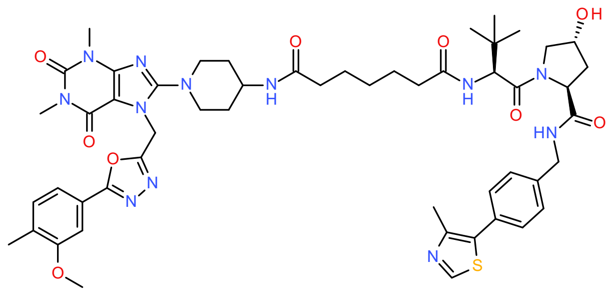

***N'*-[(*1S*)-1-[(*2S*,*4R*)-4-Hydroxy-2-[[4-(4-methylthiazol-5-yl)phenyl]methylcarbamoyl]pyrrolidine-1-carbonyl]-2,2-dimethyl-propyl]-*N*-[1-[7-[[5-(3-methoxy-4-methyl-phenyl)-1,3,4-oxadiazol-2-yl]methyl]-1,3-dimethyl-2,6-dioxo-purin-8-yl]-4-piperidyl]heptanediamide (DDD18)**

7-[[1-[7-[[5-(3-Methoxy-4-methyl-phenyl)-1,3,4-oxadiazol-2-yl]methyl]-1,3-dimethyl-2,6-dioxo-purin-8-yl]-4-piperidyl]amino]-7-oxo-heptanoic acid (67.0 %, 0.0500 g, 0.0538 mmol), (*2S*,*4R*)-1-[(*2S*)-2-amino-3,3-dimethyl-butanoyl]-4-hydroxy-*N*-[[4-(4-methylthiazol-5-yl)phenyl]methyl] pyrrolidine-2-carboxamide;hydrochloride (0.0251 g, 0.0538 mmol) and DIEA (0.0276 mL, 0.161 mmol) were dissolved in DMF (0.5 mL) and conditioned at RT for 10 min. HATU (0.0245 g, 0.0646 mmol) was added and the mixture left overnight. The reaction mixture was diluted with acetonitrile and purified via direct injection onto the preparatory HPLC system (gradient of 30% to 60% acetonitrile in 50 mM ammonium bicarbonate buffer). Lyophilization of the relevant fractions yielded the title compound (81.0%, 0.0154 g, 0.00923 mmol, yield: 17%). MS (ESI$^+$) *m/z* 518

[M+2H]$^{2+}$. HRMS *m/z* calculated 1035.4875, found 1035.4886 for [M+H]$^+$. $^1$H NMR (400 MHz, DMSO-$d_6$) δ 8.98 (s, 1H), 8.55 (t, *J* = 6.1 Hz, 1H), 7.83 (d, *J* = 9.3 Hz, 1H), 7.78 (d, *J* = 7.6 Hz, 1H), 7.47 – 7.34 (m, 7H), 5.65 (s, 2H), 5.12 (d, *J* = 3.5 Hz, 1H), 4.53 (d, *J* = 9.4 Hz, 1H), 4.47 – 4.38 (m, 2H), 4.37 – 4.31 (m, 1H), 4.26 – 4.17 (m, 1H), 3.88 (s, 3H), 3.80 – 3.68 (m, 1H), 3.70 – 3.60 (m, 2H), 3.60 – 3.50 (m, 2H), 3.41 (s, 3H), 3.15 (s, 3H), 3.11 – 3.01 (m, 2H), 2.44 (s, 3H), 2.28 – 2.18 (m, 4H), 2.15 – 2.07 (m, 1H), 2.07 – 1.97 (m, 3H), 1.95 – 1.85 (m, 1H), 1.83 – 1.72 (m, 2H), 1.55 – 1.39 (m, 4H), 1.34 – 1.14 (m, 4H), 0.92 (s, 9H).

Preparation of **DDD19**

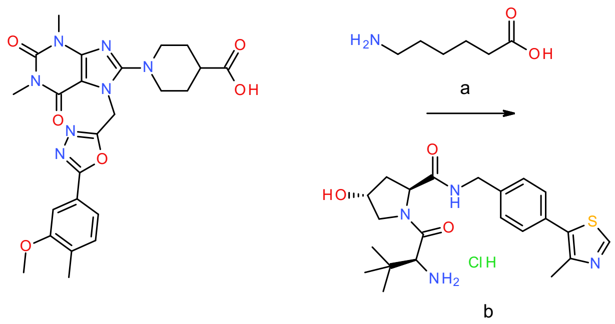

**Scheme S19.** Reagents and conditions: a) oxalyl chloride, DCM, cat. DMF, rt, 30 min; b) HATU, DIEA, DMF, rt, overnight.

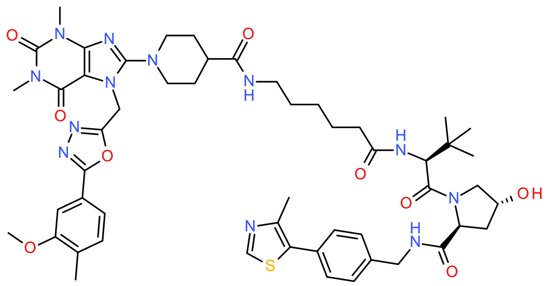

***N*-[6-[[(1S)-1-[(2S,4R)-4-Hydroxy-2-[[4-(4-methylthiazol-5-yl)phenyl]methylcarbamoyl]pyrrolidine-1-carbonyl]-2,2-dimethyl-propyl]amino]-6-oxo-hexyl]-1-[7-[[5-(3-methoxy-4-methyl-phenyl)-1,3,4-oxadiazol-2-yl]methyl]-1,3-dimethyl-2,6-dioxo-purin-8-yl]piperidine-4-carboxamide (DDD19)**

2-[4-[7-[[5-(3-Methoxy-4-methyl-phenyl)-1,3,4-oxadiazol-2-yl]methyl]-1,3-dimethyl-2,6-dioxo-purin-8-yl]piperazin-1-yl]acetic acid (95.0 %, 0.170 g, 0.308 mmol) was dissolved in DCM (10 mL) and oxalyl chloride (0.0521 mL, 0.616 mmol) and DMF (catalytic amount) added. The mixture was left at RT for 30 minutes, then the mixture was concentrated in vacuo. 5-Aminovaleric acid (0.0361 g, 0.308 mmol) in DMF (10 mL) was then added and the mixture left overnight after which no reaction was observed. At this point, HATU (100 mg) and DIEA (300 uL) were added. After the reaction was checked after 3 h, product was present and the reaction was continued via addition of (2S,4R)-1-[(2S)-2-amino-3,3-dimethyl-butanoyl]-4-hydroxy-N-[[4-(4-methylthiazol-5-yl)phenyl]methyl]pyrrolidine-2-carboxamide;hydrochloride (0.0400 g, 0.0856 mmol), HATU (40 mg) and DIEA (100 uL). After another night, workup was done via addition of water (15 mL) and EtOAc (25 mL). The organic phase was washed with saturated $NaHCO_3$ (25 mL) and water (25 mL), after which it was concentrated in vacuo and purified by preparative HPLC (gradient of 30% to 60% acetonitrile in 50 mM ammonium bicarbonate buffer). Lyophilization of the relevant fractions yielded the title compound (80.0%, 7.30 mg, 5.64 µmol, yield: 1.8%). MS (ESI$^+$) *m/z* 518 $[M+2H]^{2+}$. HRMS *m/z* calculated 1035.4875, found 1035.4895 for $[M+H]^+$. $^1$H NMR (400 MHz, DMSO-$d_6$) δ 8.98 (s, 1H), 8.55 (t, *J* = 6.0 Hz, 1H), 7.84 (d, *J* = 9.3 Hz, 1H), 7.77 (t, *J* = 5.6 Hz, 1H), 7.49 – 7.34 (m, 7H), 5.65 (s, 2H), 5.11 (d, *J* = 3.6 Hz, 1H), 4.53 (d, *J* = 9.4 Hz, 1H), 4.48 – 4.38 (m, 2H), 4.37 – 4.31 (m, 1H), 4.21 (dd, *J* = 15.8, 5.4 Hz, 1H), 3.88 (s, 3H), 3.70 – 3.54 (m, 4H), 3.40 (s, 3H), 3.15 (s, 3H), 3.05 – 2.91 (m, 4H), 2.44 (s, 3H), 2.31 – 2.18 (m, 5H), 2.14 – 1.97 (m, 2H), 1.95 – 1.85 (m, 1H), 1.75 – 1.66 (m, 4H), 1.54 – 1.41 (m, 3H), 1.40 – 1.32 (m, 2H), 1.27 – 1.18 (m, 2H), 0.92 (s, 9H).

Preparation of **DDD20**

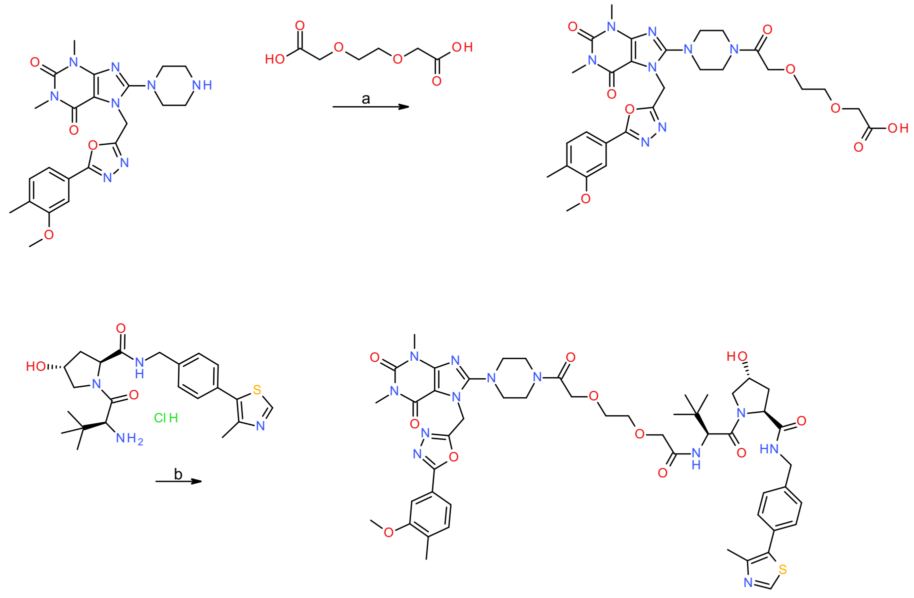

**Scheme S20.** Reagents and conditions: a) HATU, DIEA, DMF, rt, 3 h; b) HATU, DIEA, DMF, rt, overnight.

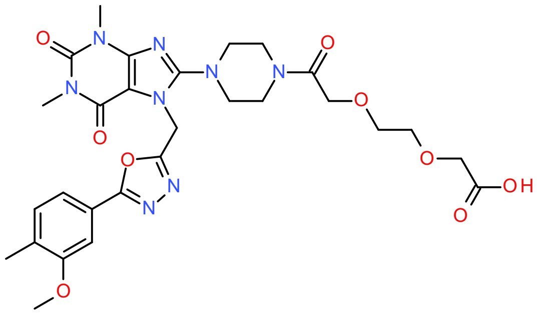

**2-[2-[2-[4-[7-[[5-(3-Methoxy-4-methyl-phenyl)-1,3,4-oxadiazol-2-yl]methyl]-1,3-dimethyl-2,6-dioxo-purin-8-yl]piperazin-1-yl]-2-oxo-ethoxy]ethoxy]acetic acid**

7-[[5-(3-Methoxy-4-methyl-phenyl)-1,3,4-oxadiazol-2-yl]methyl]-1,3-dimethyl-8-piperazin-1-yl-purine-2,6-dione (93.0 %, 0.120 g, 0.239 mmol), 2-[2-(carboxymethoxy)ethoxy]acetic acid (0.0426 g, 0.239 mmol) and DIEA (0.0837 mL, 0.489 mmol) were dissolved in DMF (1.5 mL) and conditioned at RT for 10 min. HATU (0.109 g, 0.287 mmol) was added and the mixture left at RT for 3 h. The mixture was treated with saturated $NaHCO_3$ (25 mL) and organics extracted to EtOAc (25 mL). The organic phase was then washed with brine (2x25 mL), at which point LCMS analysis indicated no product present in the organic phase. The aqueous waste was then treated with 1 M HCl, organics extracted with DCM and the organic phase dried over $MgSO_4$. Removal of the solvent in vacuo yielded the title compound as a liquid - indicating solvent still present - which was used in the next step without further purification. MS (ESI$^-$) *m/z* 625 $[M-H]^-$.

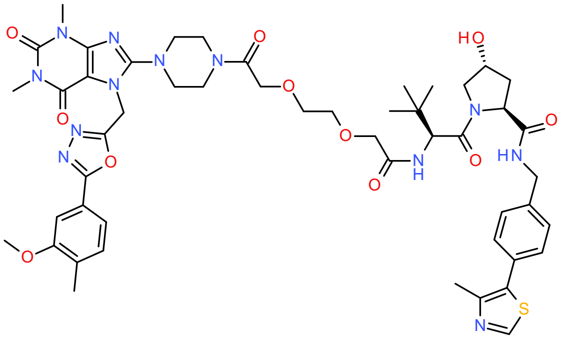

**(*2S*,*4R*)-4-Hydroxy-1-[(*2S*)-2-[[2-[2-[2-[4-[7-[[5-(3-methoxy-4-methyl-phenyl)-1,3,4-oxadiazol-2-yl]methyl]-1,3-dimethyl-2,6-dioxo-purin-8-yl]piperazin-1-yl]-2-oxo-ethoxy]ethoxy]acetyl]amino]-3,3-dimethyl-butanoyl]-*N*-[[4-(4-methylthiazol-5-yl)phenyl]methyl]pyrrolidine-2-carboxamide (DDD20)**

2-[2-[2-[4-[7-[[5-(3-Methoxy-4-methyl-phenyl)-1,3,4-oxadiazol-2-yl]methyl]-1,3-dimethyl-2,6-dioxo-purin-8-yl]piperazin-1-yl]-2-oxo-ethoxy]ethoxy]acetic acid (80.0 %, 0.0400 g, 0.00511 mmol), (*2S*,*4R*)-1-[(*2S*)-2-amino-3,3-dimethyl-butanoyl]-4-hydroxy-*N*-[[4-(4-methylthiazol-5-yl)phenyl]methyl]pyrrolidine-2-carboxamide;hydrochloride (0.0238 g, 0.00511 mmol) and DIEA

(0.0262 mL, 0.153 mmol) were dissolved in DMF (0.5 mL) and stirred at RT for 10 min. HATU (0.0233 g, 0.0613 mmol) was added and the mixture left overnight. The reaction mixture was diluted with acetonitrile and purified by preparative HPLC (gradient of 20% to 30% acetonitrile in 0.1% TFA). Lyophilization of the relevant fractions yielded the target compound (95.0%, 2.50 mg, 2.29 µmol, yield: 4.5%). MS (ESI$^+$) *m/z* 329, 493, 688 $[M+H]^+$. HRMS *m/z* calculated 1039.4461, found 1039.4479 for $[M+H]^+$. $^1$H NMR (400 MHz, DMSO-$d_6$) δ 8.96 (s, 1H), 8.56 (t, *J* = 6.0 Hz, 1H), 7.49 – 7.34 (m, 8H), 5.70 (s, 2H), 5.15 (s, 1H), 4.56 (d, *J* = 9.6 Hz, 1H), 4.43 (t, *J* = 8.1 Hz, 1H), 4.40 – 4.32 (m, 2H), 4.29 – 4.20 (m, 4H), 3.98 (s, 2H), 3.88 (s, 3H), 3.69 – 3.48 (m, 12H), 3.29 – 3.19 (m, 4H, partially obscured by the water peak), 3.14 (s, 3H), 2.43 (s, 3H), 2.22 (s, 3H), 2.09 – 2.01 (m, 1H), 1.96 – 1.83 (m, 1H), 0.92 (s, 9H).

Preparation of **DDD21**

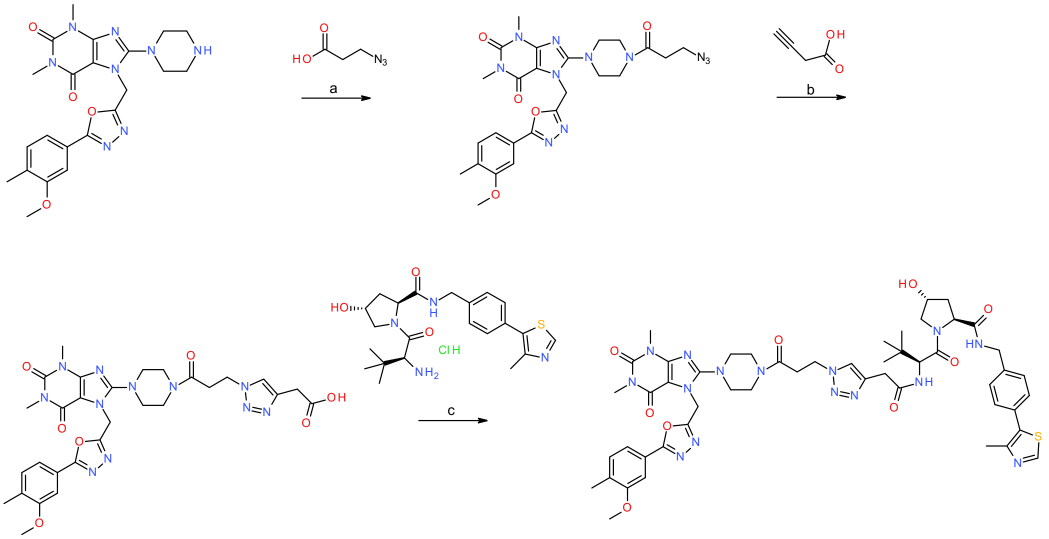

**Scheme S21.** Reagents and conditions: a) HATU, DIEA, DMF, rt, 3 h; b) Sodium *L*-ascorbate, $CuSO_4 \cdot 5\ H_2O$, DCM:*t*-BuOH (1:2) mixture, water, rt overnight; c) HATU, DIEA, DMF, rt, overnight.

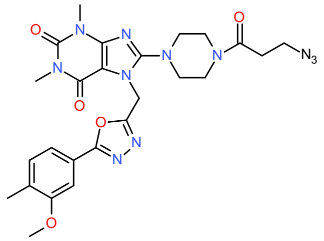

**8-[4-(3-Azidopropanoyl)piperazin-1-yl]-7-[[5-(3-methoxy-4-methyl-phenyl)-1,3,4-oxadiazol-2-yl]methyl]-1,3-dimethyl-purine-2,6-dione**

7-[[5-(3-Methoxy-4-methyl-phenyl)-1,3,4-oxadiazol-2-yl]methyl]-1,3-dimethyl-8-piperazin-1-yl-purine-2,6-dione (93.0 %, 0.120 g, 0.000239 mol), 3-azidopropanoic acid (0.0275 g, 0.239 mmol) and DIEA (0.0837 mL, 0.489 mmol) were dissolved in DMF (1.5 mL) and conditioned at RT for 10 min. HATU (0.109 g, 0.287 mmol) was added and the mixture left at RT for 3 h. The mixture was treated with saturated $NaHCO_3$ (25 mL) and organics extracted to EtOAc (25 mL). The organic phase was then washed with brine (3x25 mL) and dried over $MgSO_4$. Removal of the solvent in vacuo yielded the title compound (90.0%, 0.129 g, 0.206 mmol, yield: 86%) which was used in the next step without further purification. MS ($ESI^+$) *m/z* 564 $[M+H]^+$.

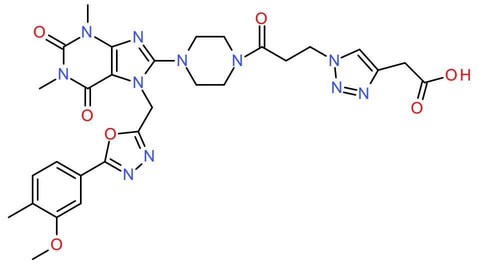

**2-[1-[3-[4-[7-[[5-(3-Methoxy-4-methyl-phenyl)-1,3,4-oxadiazol-2-yl]methyl]-1,3-dimethyl-2,6-dioxo-purin-8-yl]piperazin-1-yl]-3-oxo-propyl]triazol-4-yl]acetic acid**

8-[4-(3-Azidopropanoyl)piperazin-1-yl]-7-[[5-(3-methoxy-4-methyl-phenyl)-1,3,4-oxadiazol-2-yl]methyl]-1,3-dimethyl-purine-2,6-dione (90.0 %, 0.120 g, 0.192 mmol), but-3-ynoic acid (0.0161 g, 0.192 mmol) and sodium *L*-ascorbate (0.114 g, 0.575 mmol) were dissolved in DCM:*t*-BuOH (1:2) mixture (15 mL) and stirred for a few minutes. $CuSO_4 \cdot 5\ H_2O$ (0.0957 g, 0.383 mmol) in water (5 mL) was added and the mixture stirred at rt. After 30 minutes, another 5 mL of DCM was added, and after 90 min, another 17 mg of but-3-ynoic acid and 114 mg of sodium *L*-ascorbate were added and the mixture left overnight. The reaction mixture was diluted with EtOAc (25 mL) and the resulting organic phase washed with ammonia (28%) (25 mL) and brine (2x25 mL). At this stage, the product was found in the aqueous phase. The aqueous phase was acidified with concentrated HCl and organics extracted again with EtOAc (70 mL). The new organic phase was dried over $MgSO_4$ and the solvent removed in vacuo to afford the title compound (95.0%, 0.120 g, 0.176 mmol, yield: 92%) which was used in the next step without further purification. MS ($ESI^-$) *m/z* 646 $[M-H]^-$.

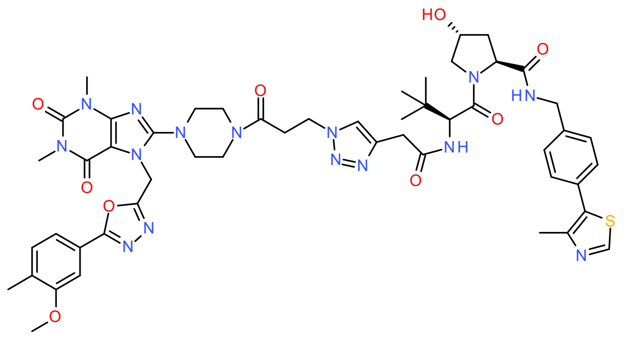

**(*2S*,*4R*)-4-Hydroxy-1-[(*2S*)-2-[[2-[1-[3-[4-[7-[[5-(3-methoxy-4-methyl-phenyl)-1,3,4-oxadiazol-2-yl]methyl]-1,3-dimethyl-2,6-dioxo-purin-8-yl]piperazin-1-yl]-3-oxo-propyl]triazol-4-yl]acetyl]amino]-3,3-dimethyl-butanoyl]-*N*-[[4-(4-methylthiazol-5-yl)phenyl]methyl]pyrrolidine-2-carboxamide (DDD21)**

2-[1-[3-[4-[7-[[5-(3-Methoxy-4-methyl-phenyl)-1,3,4-oxadiazol-2-yl]methyl]-1,3-dimethyl-2,6-dioxo-purin-8-yl]piperazin-1-yl]-3-oxo-propyl]triazol-4-yl]acetic acid (95.0 %, 0.0500 g, 0.0733 mmol), (*2S*,*4R*)-1-[(*2S*)-2-amino-3,3-dimethyl-butanoyl]-4-hydroxy-*N*-[[4-(4-methylthiazol-5-yl)phenyl]methyl]pyrrolidine-2-carboxamide;hydrochloride (0.0343 g, 0.0733 mmol) and DIEA (0.0377 mL, 0.220 mmol) were dissolved in DMF (0.5 mL) and the mixture stirred a few minutes at rt. HATU (0.0335 g, 0.0880 mmol) was added and the mixture left overnight. Additional 34 mg of VHL ligand, 37.8 uL of DIEA and 34 mg of HATU were added and the mixture left over the weekend. The reaction mixture was diluted with acetonitrile and purified by preparative HPLC (gradient of 15% to 30% acetonitrile in 0.1% TFA). Lyophilization of the relevant fractions yielded the title compound (88.0%, 19.9 mg, 16.5 µmol, yield: 22%). MS (ESI$^+$) *m/z* 531 $[M+2H]^{2+}$. HRMS *m/z* calculated 1060.4576, found 1060.4589 for $[M+H]^+$. $^1$H NMR (400 MHz, DMSO-$d_6$) δ 8.98 (s, 1H), 8.57 (t, *J* = 6.1 Hz, 1H), 8.15 (d, *J* = 9.3 Hz, 1H), 7.88 (s, 1H), 7.48 – 7.35 (m, 7H), 5.71 (s, 2H), 4.58 – 4.49 (m, 3H), 4.46 – 4.38 (m, 2H), 4.36 – 4.31 (m, 1H), 4.22 (dd, *J* = 15.8, 5.4 Hz, 1H), 3.88 (s, 3H), 3.69 – 3.51 (m, 8H, overlapping with water), 3.41 (s, 3H), 3.28 – 3.20 (m, 4H), 3.15 (s, 3H), 3.00 (t, *J* = 6.9 Hz, 2H), 2.44 (s, 3H), 2.23 (s, 3H), 2.07 – 1.97 (m, 1H), 1.94 – 1.85 (m, 1H), 0.93 (s, 9H). -OH proton is missing.

<u>Preparation of **DDD22**</u>

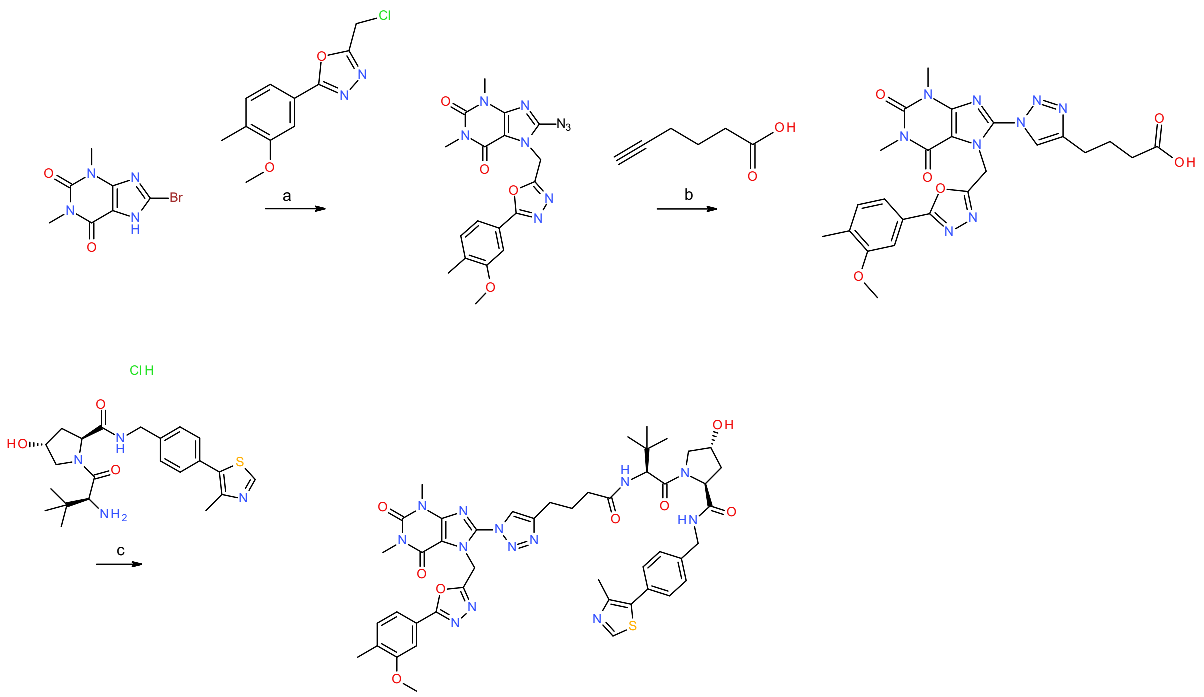

**Scheme S22.** Reagents and conditions: a) $K_2CO_3$, DMF, 65 °C, 2 h then $NaN_3$, 1 h; b) Sodium *L*-ascorbate, $CuSO_4 \cdot 5\ H_2O$, acetone:water (2.5:1) mixture, *t*-BuOH, rt, 4 h; c) HATU, DIEA, DMF, rt, overnight.

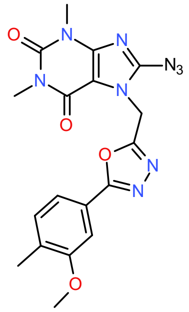

**8-Azido-7-[[5-(3-methoxy-4-methyl-phenyl)-1,3,4-oxadiazol-2-yl]methyl]-1,3-dimethyl-purine-2,6-dione**

2-(Chloromethyl)-5-(3-methoxy-4-methyl-phenyl)-1,3,4-oxadiazole (0.100 g, 0.390 mmol), 8-bromo-1,3-dimethyl-7H-purine-2,6-dione (0.111 g, 0.429 mmol) and $K_2CO_3$ (0.0646 g, 0.468 mmol) were mixed in DMF (2 mL) and heated to 65 °C for 2 h. Then, $NaN_3$ (0.0380 g, 0.584 mmol) was added and the mixture left for 1 h. The mixture was diluted in water (10 mL) and the mixture extracted with EtOAc (20 mL). The organic phase was washed with brine (2x20 mL) and dried over $MgSO_4$. Removal of the solvent in vacuo yielded the title compound (81.0%, 0.0550 g, 0.105 mmol, yield: 27%). MS ($ESI^+$) *m/z* 424 $[M+H]^+$.

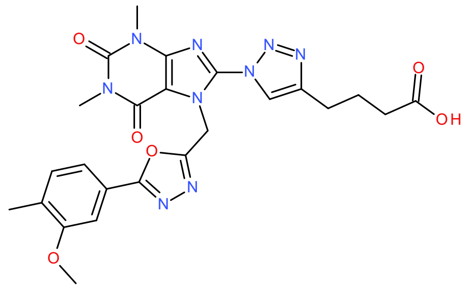

**4-[1-[7-[[5-(3-Methoxy-4-methyl-phenyl)-1,3,4-oxadiazol-2-yl]methyl]-1,3-dimethyl-2,6-dioxo-purin-8-yl]triazol-4-yl]butanoic acid**

To a suspension of hex-5-ynoic acid (0.0160 g, 0.143 mmol), 8-azido-7-[[5-(3-methoxy-4-methyl-phenyl)-1,3,4-oxadiazol-2-yl]methyl]-1,3-dimethyl-purine-2,6-dione (0.0550 g, 0.130 mmol) in acetone (2.50 mL) and water (1.00 mL) were added L-ascorbic acid sodium salt (0.0257 g, 0.130 mmol) and copper(II)sulfatepentahydrate (0.00487 g, 0.0195 mmol). The reaction was stirred at rt for 15 min. *tert*-Butanol (0.500 mL) was added to increase solubility and the reaction stirred at rt for 3 h. About 10% of starting material remained so additional alkyne was added (3 µL, 0.2 eqiuv). The reaction was stirred for another 1 h. EtOAc and saturated aqueous $NaHCO_3$ were added and the phases separated. The aqueous phase was acidified with 1 M aqueous HCl and extracted with EtOAc (3x). The combined organic phases were dried over magnesium sulfate and evaporated and dried under vacuum to give 90 mg of the crude title compound. The crude material was taken to the next step without further purification. MS ($ESI^+$) *m/z* 536 $[M+H]^+$.

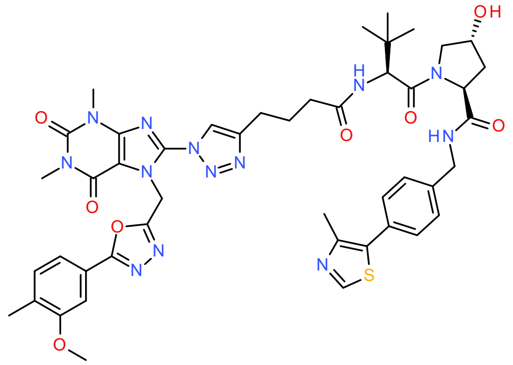

**(*2S*,*4R*)-4-Hydroxy-1-[(*2S*)-2-[4-[1-[7-[[5-(3-methoxy-4-methyl-phenyl)-1,3,4-oxadiazol-2-yl]methyl]-1,3-dimethyl-2,6-dioxo-purin-8-yl]triazol-4-yl]butanoylamino]-3,3-dimethyl-butanoyl]-*N*-[[4-(4-methylthiazol-5-yl)phenyl]methyl]pyrrolidine-2-carboxamide (DDD22)**

4-[1-[7-[[5-(3-Methoxy-4-methyl-phenyl)-1,3,4-oxadiazol-2-yl]methyl]-1,3-dimethyl-2,6-dioxo-purin-8-yl]triazol-4-yl]butanoic acid (0.0400 g, 0.0672 mmol), (2*S*,4*R*)-1-[(2*S*)-2-amino-3,3-dimethyl-butanoyl]-4-hydroxy-*N*-[[4-(4-methylthiazol-5-yl)phenyl]methyl]pyrrolidine-2-carboxamide;hydrochloride (0.0314 g, 0.0672 mmol) and DIEA (0.0345 mL, 0.202 mmol) were dissolved in DMF (0.5 mL) and conditioned at rt for 10 min. HATU (0.0307 g, 0.0807 mmol) was added and the mixture left overnight. The reaction mixture was diluted with acetonitrile and purified by preparatory HPLC (gradient of 30% to 60% acetonitrile in 50 mM ammonium bicarbonate buffer). Lyophilization of the pure fractions afforded the title compound (98.0%, 15.4 mg, 14.8 µmol, yield: 33%). HRMS: *m/z* calculated 948.3939, found 948.3954 for $[M+H]^+$. $^1H$ NMR (400 MHz, DMSO-$d_6$) δ 8.97 (s, 1H), 8.69 (s, 1H), 8.55 (t, *J* = 5.9 Hz, 1H), 7.92 (d, *J* = 9.2 Hz, 1H), 7.46 – 7.33 (m, 7H), 6.20 (s, 2H), 5.11 (d, *J* = 3.6 Hz, 1H), 4.54 (d, *J* = 9.3 Hz, 1H), 4.47 – 4.40 (m, 2H), 4.35 (s, 1H), 4.20 (dd, *J* = 16.0, 5.4 Hz, 1H), 3.88 (s, 3H), 3.68 – 3.64 (m, 2H), 3.48 (s, 3H), 3.25 (s, 3H), 2.72 (t, *J* = 7.6 Hz, 2H), 2.43 (s, 3H), 2.38 – 2.28 (m, 1H), 2.27 – 2.17 (m, 4H), 2.08 – 1.95 (m, 1H), 1.95 – 1.82 (m, 3H), 0.94 (s, 9H).

Preparation of **DDD23**

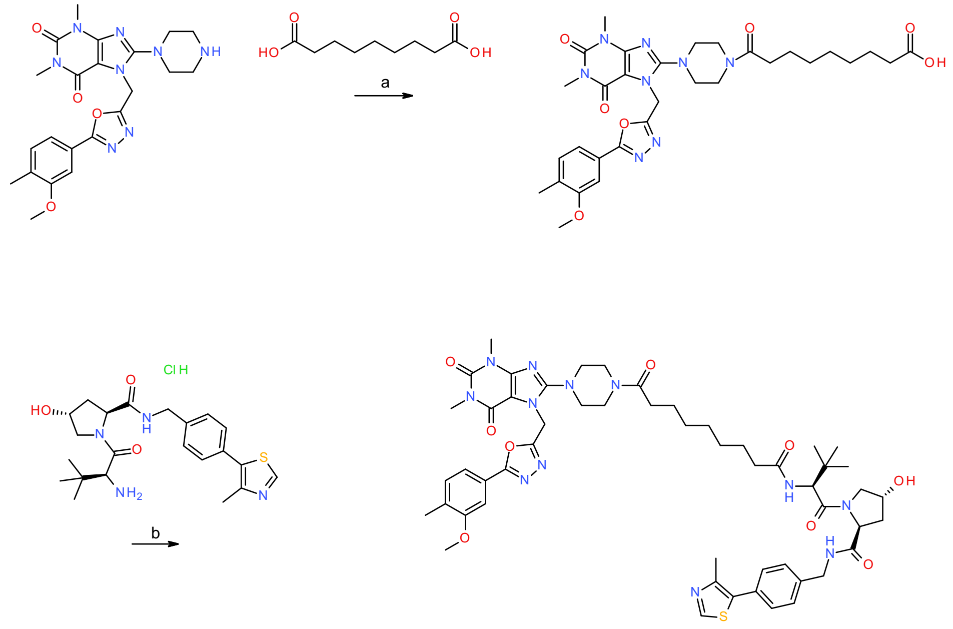

**Scheme S23.** Reagents and conditions: a) HATU, DIEA, DMF, rt, 4 h; b) HATU, DIEA, DMF, rt, overnight.

**9-[4-[7-[[5-(3-Methoxy-4-methyl-phenyl)-1,3,4-oxadiazol-2-yl]methyl]-1,3-dimethyl-2,6-dioxo-purin-8-yl]piperazin-1-yl]-9-oxo-nonanoic acid**

7-[[5-(3-Methoxy-4-methyl-phenyl)-1,3,4-oxadiazol-2-yl]methyl]-1,3-dimethyl-8-piperazin-1-yl-purine-2,6-dione (0.120 g, 0.239 mmol), nonanedioic acid (0.0450 g, 0.239 mmol) and DIEA (0.0837 mL, 0.489 mmol) were dissolved in DMF (1.5 mL) and conditioned at rt for 10 min. HATU (0.109 g, 0.287 mmol) was added and the mixture left at rt for 4 h. The mixture was treated with saturated aqueous $NaHCO_3$ solution (20 mL) and organic impurities extracted with DCM

(20 mL). The aqueous phase was then acidified with 1 M HCl and organics extracted to EtOAc (20 mL). The organic phase was washed with brine (20 mL) and dried over $MgSO_4$. Removal of the solvent in vacuo afforded the title compound. (0.0630 g, 0.0178 mmol, yield: 7.4%). The crude material was used in the next step without further purification. MS ($ESI^-$) *m/z* 634 $[M-H]^-$.

**(*2S*,*4R*)-4-Hydroxy-1-[(*2S*)-2-[[9-[4-[7-[[5-(3-methoxy-4-methyl-phenyl)-1,3,4-oxadiazol-2-yl]methyl]-1,3-dimethyl-2,6-dioxo-purin-8-yl]piperazin-1-yl]-9-oxo-nonanoyl]amino]-3,3-dimethyl-butanoyl]-*N*-[[4-(4-methylthiazol-5-yl)phenyl]methyl]pyrrolidine-2-carboxamide (DDD23)**

9-[4-[7-[[5-(3-Methoxy-4-methyl-phenyl)-1,3,4-oxadiazol-2-yl]methyl]-1,3-dimethyl-2,6-dioxo-purin-8-yl]piperazin-1-yl]-9-oxo-nonanoic acid (0.0630 g, 0.0178 mmol), (*2S*,*4R*)-1-[(*2S*)-2-amino-3,3-dimethyl-butanoyl]-4-hydroxy-*N*-[[4-(4-methylthiazol-5-yl)phenyl]methyl]pyrrolidine-2-carboxamide;hydrochloride (0.00832 g, 0.0178 mmol) and DIEA (0.00915 mL, 0.0534 mmol) were dissolved in DMF (0.5 mL) and conditioned at rt for 10 min. HATU (0.00813 g, 0.0214 mmol) was added and the mixture left overnight. The reaction mixture was diluted with acetonitrile and purified by preparatory HPLC (gradient of 30% to 60% acetonitrile in 0.1% TFA). Lyophilization of the pure fractions afforded the title compound (84.0%, 2.50 mg, 2.00 µmol, yield: 11%). MS ($ESI^+$) *m/z* 525 $[M+2H]^{2+}$. HRMS: *m/z* calculated 1049.5031, found 1049.5041 for $[M+H]^+$.

<u>Preparation of **DDD24**</u>

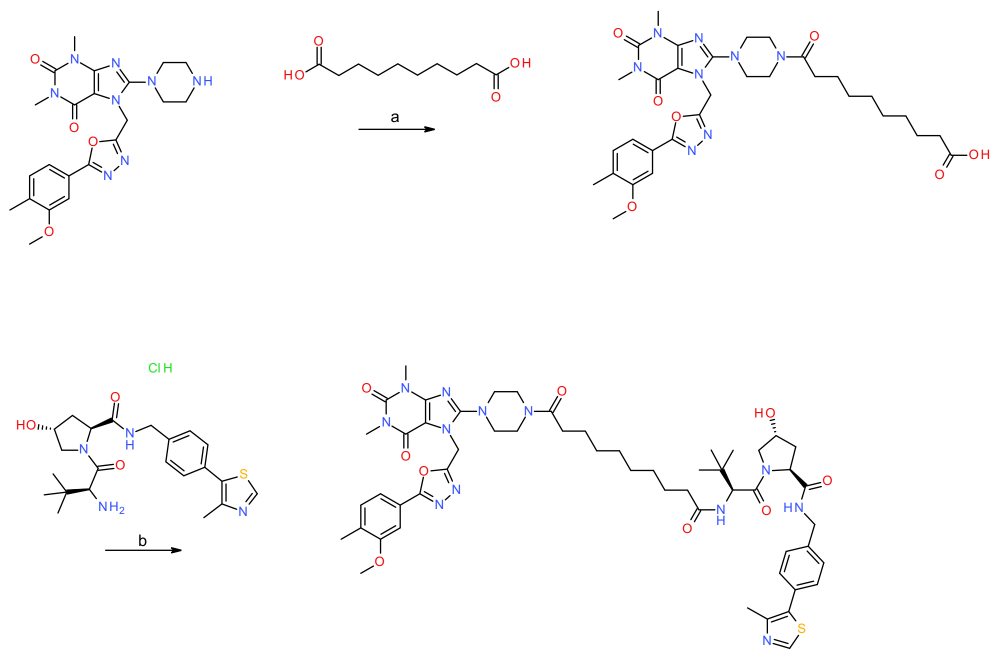

**Scheme S24.** Reagents and conditions: a) HATU, DIEA, DMF, rt, 4 h; b) HATU, DIEA, DMF, rt, overnight.

**10-[4-[7-[[5-(3-Methoxy-4-methyl-phenyl)-1,3,4-oxadiazol-2-yl]methyl]-1,3-dimethyl-2,6-dioxo-purin-8-yl]piperazin-1-yl]-10-oxo-decanoic acid**

7-[[5-(3-Methoxy-4-methyl-phenyl)-1,3,4-oxadiazol-2-yl]methyl]-1,3-dimethyl-8-piperazin-1-yl-purine-2,6-dione (0.120 g, 0.239 mmol), decanedioic acid (0.0484 g, 0.239 mmol) and DIEA (0.0837 mL, 0.489 mmol) were dissolved in DMF (1.5 mL) and conditioned at rt for 10 min. HATU (0.109 g, 0.287 mmol) was added and the mixture left at rt for 4 h. The mixture was treated with saturated aqueous $NaHCO_3$ solution (20 mL) and organic impurities extracted with DCM (20 mL). The aqueous phase was then acidified with 1 M HCl and organics extracted to EtOAc (20 mL). The organic phase was washed with brine (20 mL) and dried over $MgSO_4$. Removal of the solvent in vacuo yielded the title compound (0.0120 g, 0.0129 mmol, yield: 5.4%) The crude material was used in the next step without further purification. MS (ESI$^+$) *m/z* 651 $[M+H]^+$.

**(*2S*,*4R*)-4-Hydroxy-1-[(*2S*)-2-[[10-[4-[7-[[5-(3-methoxy-4-methyl-phenyl)-1,3,4-oxadiazol-2-yl]methyl]-1,3-dimethyl-2,6-dioxo-purin-8-yl]piperazin-1-yl]-10-oxo-decanoyl]amino]-3,3-dimethyl-butanoyl]-*N*-[[4-(4-methylthiazol-5-yl)phenyl]methyl]pyrrolidine-2-carboxamide (DDD24)**

10-[4-[7-[[5-(3-Methoxy-4-methyl-phenyl)-1,3,4-oxadiazol-2-yl]methyl]-1,3-dimethyl-2,6-dioxo-purin-8-yl]piperazin-1-yl]-10-oxo-decanoic acid (0.0120 g, 1.29e-5 mol), (*2S*,*4R*)-1-[(*2S*)-2-amino-3,3-dimethyl-butanoyl]-4-hydroxy-*N*-[[4-(4-methylthiazol-5-yl)phenyl]methyl] pyrrolidine-2-carboxamide;hydrochloride (0.00616 g, 0.0132 mmol)) and DIEA (0.00678 mL, 3.96e-5 mol) were dissolved in DMF (0.5 mL) and conditioned at rt for 10 min. HATU (0.00602 g, 1.58e-5 mol) was added and the mixture left overnight. The reaction mixture was diluted with acetonitrile and purified via direct injection onto the preparatory HPLC system (gradient of 30% to 60% acetonitrile in 0.1% TFA). Lyophilization of the pure fractions yielded the target compound (93.0%, 2.00 mg, 1.75 µmol, yield: 14%). MS (ESI$^+$) *m/z* 532 $[M+2H]^{2+}$. HRMS: *m/z* calculated 1063.5187, found 1063.5208 for $[M+H]^+$. $^1$H NMR (400 MHz, DMSO-$d_6$) δ 8.98 (s, 1H), 8.55 (t, *J* = 6.1 Hz, 1H), 7.83 (d, *J* = 9.2 Hz, 1H), 7.47 – 7.42 (m, 3H), 7.41 – 7.36 (m, 4H), 5.71 (s, 2H), 5.11 (br s, 1H), 4.54 (d, *J* = 9.3 Hz, 1H), 4.46 – 4.39 (m, 2H), 4.36 – 4.32 (m, 1H), 4.21 (dd, *J* = 15.8, 5.4 Hz, 1H), 3.88 (s, 3H), 3.68 – 3.62 (m, 2H), 3.59 – 3.52 (m, 4H), 3.41 (s, 3H), 3.28 – 3.19 (m, 4H), 3.15 (s, 3H), 2.44 (s, 3H), 2.33 – 2.27 (m, 3H), 2.23 (s, 3H), 2.15 – 2.05 (m, 1H), 2.05 – 1.97 (m, 1H), 1.93 – 1.86 (m, 1H), 1.53 – 1.40 (m, 4H), 1.26 – 1.20 (m, 8H), 0.93 (s, 9H).

<u>Preparation of **DDD25**</u>

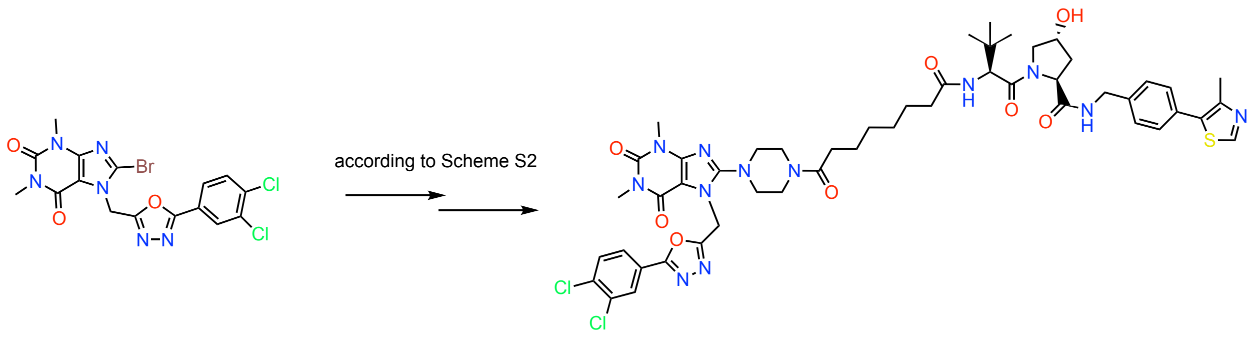

**1a** **DDD25**

*prepared according to ref*

**Scheme S25.** Reagents and conditions according to Scheme S2.

**DDD25** was prepared from 1a following the synthetic sequence outlined in Scheme S2 to yield (*2S*,*4R*)-1-((*S*)-2-(8-(4-(7-((5-(3,4-dichlorophenyl)-1,3,4-oxadiazol-2-yl)methyl)-1,3-dimethyl-2,6-dioxo-2,3,6,7-tetrahydro-1*H*-purin-8-yl)piperazin-1-yl)-8-oxooctanamido)-3,3-dimethylbutanoyl)-4-hydroxy-*N*-(4-(4-methylthiazol-5-yl)benzyl)pyrrolidine-2-carboxamide (**DDD25**) after preparatory HPLC system (gradient of 30% to 70% acetonitrile in 0.1% TFA). Lyophilization of the pure fractions yielded the target compound (6.8 mg, 0.006 mmol, yield: 23%). MS ($ESI^+$) *m/z* $[M+2H]^{2+}$. HRMS: *m/z* calculated 1058.3755, found for $[M+H]^+$. $^1H$ NMR (400 MHz, DMSO-$d_6$) δ 8.88 (s, 1H), 8.19 (s, 1H), 7.96 (d, 1H), 7.77 (d, 1H), 7.47 – 7.43 (m, 4H), 5.78 (s, 2H), 4.56 – 4.36 (m, 6H), 3.91 (m, 1H), 3.81 (m, 1H), 3.73 (m, 4H), 3.53 (s, 3H), 3.41 (s, 3H), 3.37 (m, 2H), 3.30 (s, 3H), 2.48 (s, 3H), 2.42 – 2.09 (m, 4H), 1.62 (m, br, 4H), 1.38 (m, br, 4H), 1.05 (s, 9H).

#### *Synthesis of compounds in Figure 5H*

The CRBN-based compounds **DDD26-DDD50** were obtained in >95% purity from Julia Eklund and Fredrik Klingegård (unpublished).

### Supplemental Figures

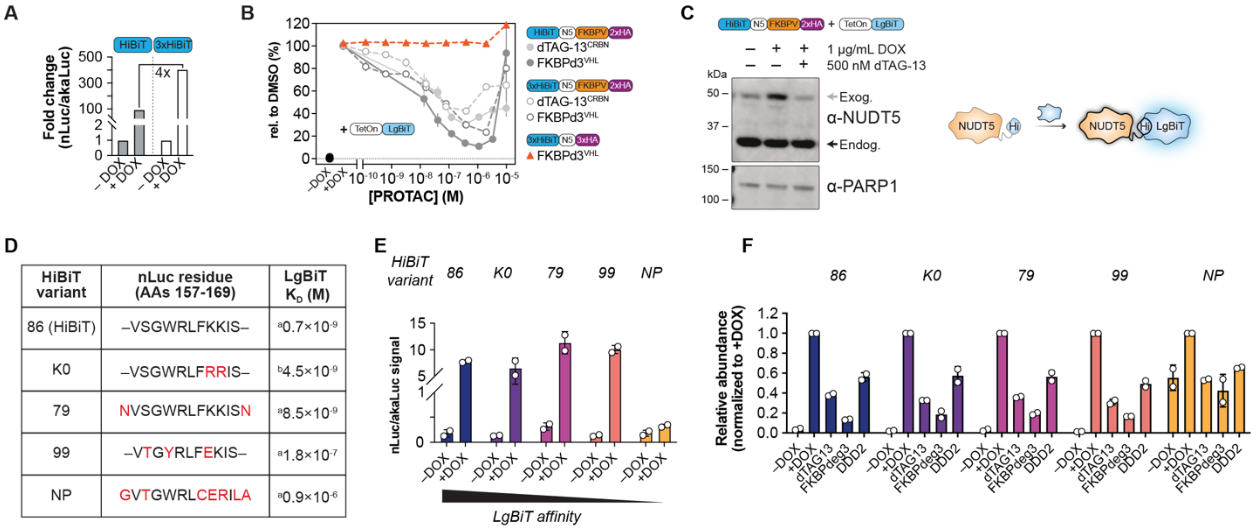

| HiBiT variant | nLuc residue (AAs 157-169) | LgBiT $K_D$ (M) |
|---|---|---|
| 86 (HiBiT) | –VSGWRLFKKIS– | [a]0.7×10⁻⁹ |
| K0 | –VSGWRLFRRIS– | [b]4.5×10⁻⁹ |
| 79 | NVSGWRLFKKISN | [a]8.5×10⁻⁹ |
| 99 | –VTGYRLFEKIS– | [a]1.8×10⁻⁷ |
| NP | GVTGWRLCERILA | [a]0.9×10⁻⁶ |

**Supplemental Figure 1. Optimization and characterization of HiBiT tags in luminescent degradation reporters using U-2 OS cells. A.** Luminescence fold change (nLuc/akaLuc reference) measured in U-2 OS cells expressing NUDT5 degradation reporters tagged with either a single HiBiT or a tandem 3× HiBiT tag. Cells were treated with doxycycline (DOX; 1 µg/mL) for 24h for induction of LgBiT expression. Results from a single experiment. **B.** Dose-response curves for degradation of reporters with single or 3× HiBiT tags (±FKBPV) after 24 h treatment with increasing concentrations of PROTACs dTAG-13 or FKBPd3. LgBiT expression was induced by doxycycline (DOX; 0.75 µg/mL) 24 h prior to compound treatment. nLuc signals are relative to DMSO control after normalization to akaLuc. Data represent mean ± SD (n=2). **C.** Western blot of NUDT5 protein levels in U-2 OS cells expressing the finalized single HiBiT reporter following DOX induction and treatment with 500 nM dTAG-13 for 24 h (single experiment). Exogenous (HiBiT-tagged) and endogenous NUDT5 are indicated by gray and black arrows, respectively. Included is a schematic of how the HiBiT–NUDT5 fusion protein may be stabilized by complementation with LgBiT to generate luminescence signal. **D.** Table listing HiBiT peptide variants (86 - canonical HiBiT, 79, 99, and NP) with their NanoLuc (nLuc) residue sequences. Amino acid differences from 86 are highlighted in red, and LgBiT binding affinities (KD in molar units) from literature references, denoted as a[5] and b[6]. **E.** Luminescence fold change measured in U-2 OS cells expressing different HiBiT variants before and after DOX induction (1 µg/mL), showing the effect of LgBiT complementation on signal. Data are mean ± SD (n=2). **F.** Degradation assays in cells expressing HiBiT variants treated with PROTACs dTAG-13 (1 µM), FKBPd3 (0.5 µM), or DDD2 (1 µM), normalized to akaLuc and relative to DOX. Data are mean ± SD (n=2).

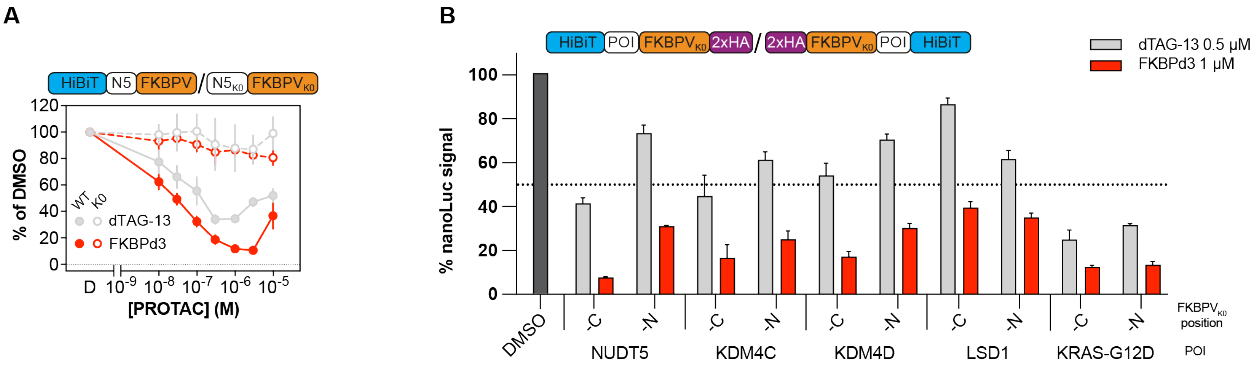

**Supplemental Figure 2. Confirmation of degradation reporter. A.** Dose-response analysis of dTAG-13 (0.5 µM) and FKBPd3 (1 µM) treatment for 24 hours in U-2 OS cells expressing wild-type NUDT5 (N5) and FKBP-tag or its lysine-free variant ($N5_{K0}$)-FKBPV $_{K0}$). Doxycycline (0.75 µg/mL) was used for induction prior to treatment. Luminescence signals were normalized to akaLuc and expressed relative to DMSO controls. Data represent mean ± SD (n=3). **B.** Comparison of C- versus N-terminal tagging of reporters across different proteins of interest (POIs). NanoLuc signal was measured in U-2 OS cells expressing dual-luminescent degradation reporters with $FKBPV_{K0}$ tags on either terminus, following doxycycline (0.75 µg/mL) induction of LgBiT expression 24 hours prior to treatment with dTAG-13 (0.5 µM) or FKBPd3 for 24 hours. Data are normalized to akaLuc and expressed relative to DMSO control. Means ± SD (n=3).

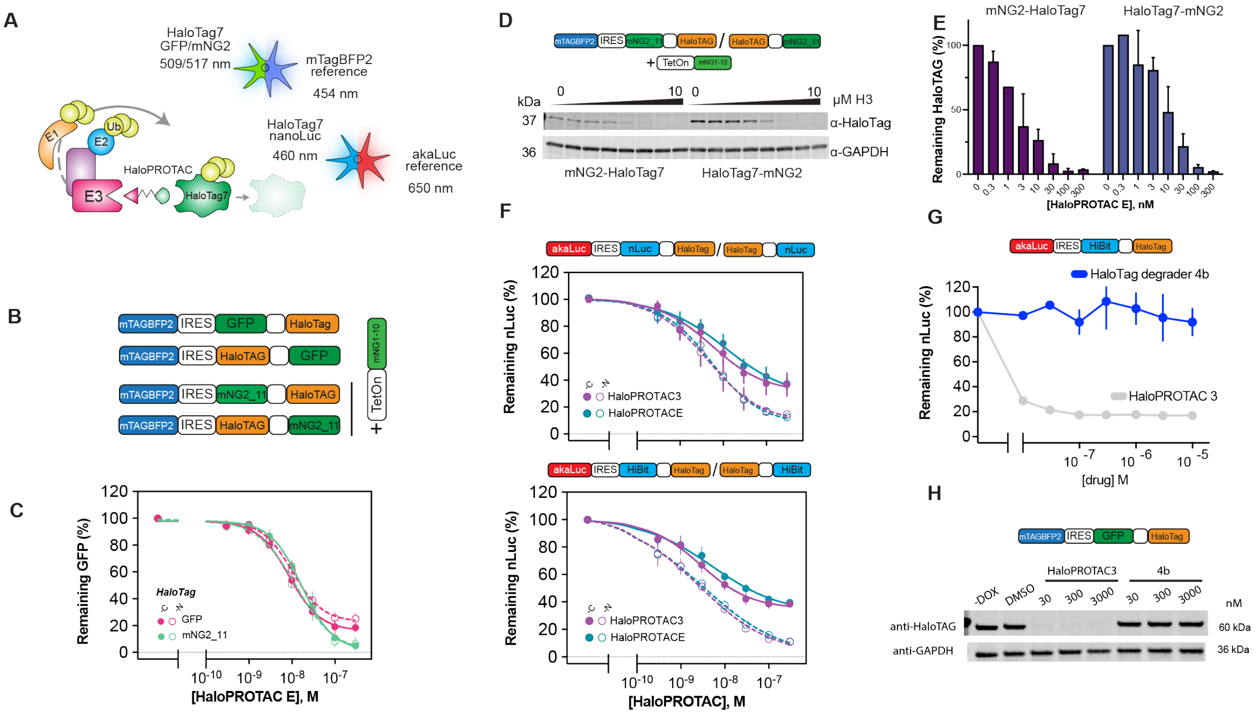

**Supplemental Figure 3. Development and evaluation of HaloPROTAC degradation assays using fluorescent and luminescent reporters. A.** Illustration of the dual-reporter system for HaloTag7 fusion proteins. HaloTag7, which binds E3 ligases via HaloPROTACs for degradation, is tagged with GFP or mNeonGreen2 (mNG2; excitation/emission 509/517 nm) to monitor abundance, with mTagBFP2 (excitation 454 nm) as a reference. For luminescence, HaloTag7 is tagged with HiBiT and combined with akaLuc (nLuc emission 460 nm, akaLuc 650 nm) to sensitively detect degradation. **B.** Schematic of four reporter constructs adapted from pENTR2X, containing mTagBFP2 and HaloTag7 fused to GFP or split mNG2_11 tags. Inducible mNG2_1–10 complementation enables conditional fluorescence. Variants differ by N- or C-terminal tag placement to assess positional effects on degradation. **C.** Dose–response flow cytometry of U-2 OS cells expressing HaloTag7-GFP or -mNG2_11 (N- or C-terminal), with DOX-induced mNG2_1–10 and 24 h HaloPROTAC-E treatment. Data shown as % of DMSO control (mean ± SD, n=2) gated on mTagBFP2-positive cells. **D.** Representative Western blot of U-2 OS cells expressing HaloTag7-mNG2_11 variants treated with 0–10 µM HaloPROTAC-3 for 24 h post-DOX induction (0.75 µg/mL), probed with anti-HaloTag and anti-GAPDH. **E.** Quantification of HaloTag degradation from two independent blots normalized to GAPDH and DMSO control (as in **D**). **F.** Comparison of degradation efficiency of HaloTag7 luminescent reporters (NanoLuc or HiBiT; varying tag position), treated with HaloPROTAC-E and -3 over a concentration range. Luminescence normalized to akaLuc and DMSO. Data represent two independent experiments, each in triplicate (mean ± SD). **G.** Luminescence dose-response in U-2 OS cells expressing HiBiT-HaloTag7 reporter treated for 24 h with HaloPROTAC-3 or HaloTag degrader 4b after DOX induction (0.75 µg/mL). Data normalized to akaLuc and expressed relative to DMSO (mean ± SD, two independent experiments, triplicates). **H.** Western blot of HEK293T cells expressing N-terminal GFP-HaloTag7 fusion treated with HaloTag degrader 4b for 24 h, probed with anti-HaloTag and anti-GAPDH. A single experiment was performed.

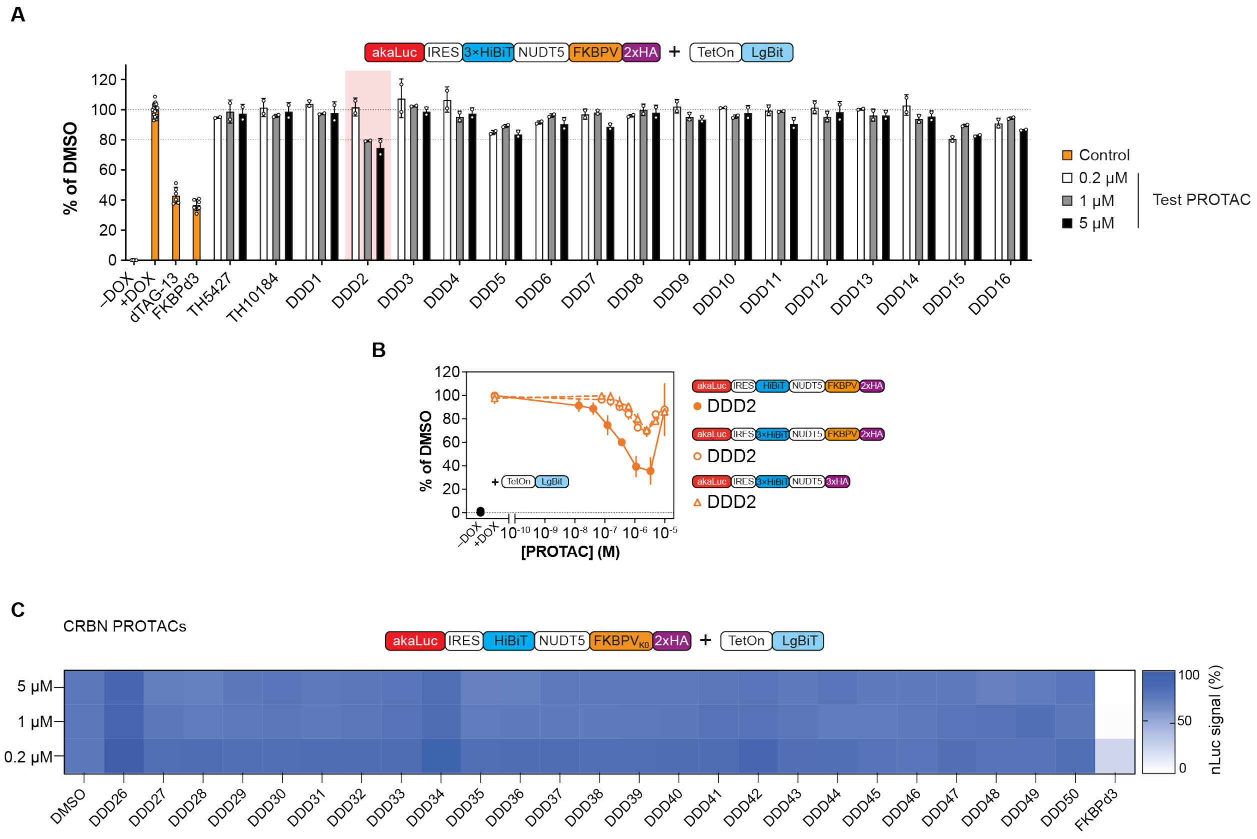

**Supplemental Figure 4. NUDT5 PROTAC screen using luminescence degradation reporters. A.** Initial screening of 16 NUDT5-targeting PROTACs (DDD1-16) in U-2 OS cells expressing 3×HiBiT-tagged NUDT5 degradation reporter and LgBiT. Cells were treated for 24 hours with three concentrations (0.2, 1, and 5 µM), following doxycycline (0.75 µg/mL) induction of LgBiT. dTAG-13 (0.5 µM) and FKBPd3 (1 µM) were included as degradation controls. NanoLuc signals were normalized to the akaLuc internal reference and expressed relative to DMSO. Data represent mean ± SD (n=3). **B.** Dose-response curves for degradation of reporters with single or 3×HiBiT tags (±FKBPV) after 24 h treatment with increasing concentrations of DDD2. LgBiT expression was induced by doxycycline (DOX; 0.75 µg/mL) 24 h prior to compound treatment. nLuc signals are relative to DMSO control after normalization to akaLuc. Data represent mean ± SD (n=2). **C.** Screening of additional CRBN-based NUDT5 PROTACs synthesized (DDD25-50) using the optimized 1×HiBiT degradation reporter in U-2 OS cells. Cells were treated compounds at 0.2, 1, and 5 µM for 24 h. Doxycycline (0.75 µg/mL) was added 24 h prior to treatment to induce LgBiT expression. Luminescence signals were normalized to akaLuc and expressed relative to DMSO controls. FKBPd3 was included as a positive degradation control. Heat map colors represent normalized NanoLuc signal (mean of n=3), from white (low) to dark blue (high).

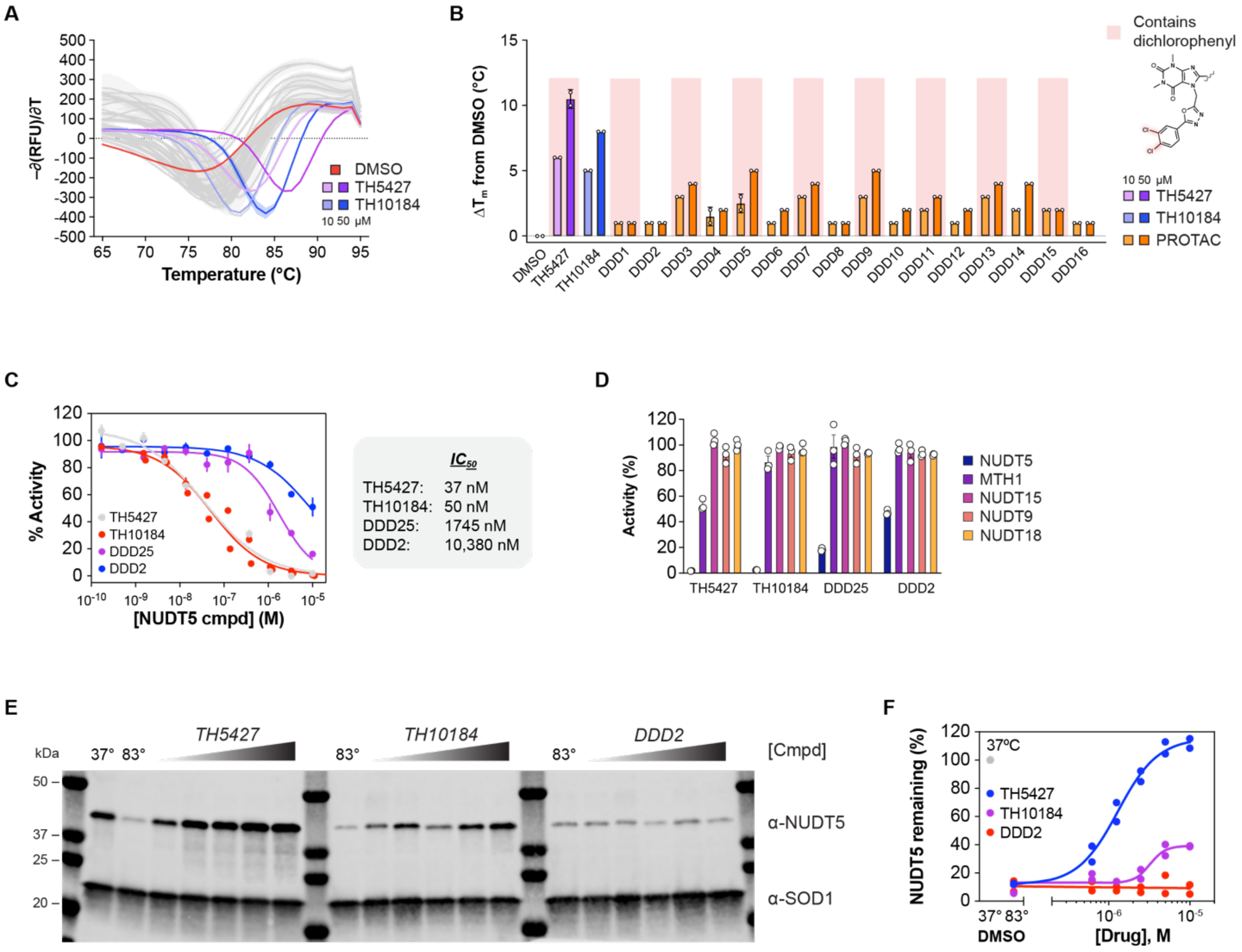

**Supplemental Figure 5. Biophysical and biochemical characterization of NUDT5 warheads and PROTACs. A.** Differential scanning fluorimetry (DSF) analysis of purified NUDT5 thermal denaturation with DMSO (red), TH5427 (purple), TH10184 (blue), or test NUDT5 PROTACs (grey) at 10 (light) or 50 µM (dark). Means values representing the negative first derivative (–d[RFU]/dT) from n=2 experiments ± SD (shading) are shown. **B.** Melting temperature change ($DT_m$) with respect to DMSO treatment from experiments in A. As before 10 and 50 µM groups are indicated by light and dark colors, respectively. Test PROTACs are indicated in orange. Molecules containing TH5427 are highlighted in red. Means from n=2 experiments ± SD are shown. **C.** Enzyme-coupled malachite green NUDT5 inhibition assay comparing inhibition of ADP-ribose conversion to ribose-5-phosphate with TH5427 (grey), TH10184 (red), DDD2 (blue), and DDD25 (purple). Means with lines-of-best-fit from two technical replicates shown ± SD (except for TH10184, which is n=2 independent experiments [in duplicate]). Calculated $IC_{50}$ values are also indicated. **D.** Selectivity of TH5427, TH10184, DDD2, and DDD25 towards purified NUDT5 (blue), MTH1 (purple), NUDT15 (light purple), NUDT9 (orange), and NUDT18 (yellow) at 10 µM. Means from three technical replicates are shown ± SD. **E.** Representative ITDRF CETSA experiment (from n=2) comparing intracellular NUDT5 binding of TH5427, TH10184, and DDD25 at concentrations ranging from 0.625 to 10 µM. SOD1 was probed as a loading control for temperature pulses. DMSO samples heated to 37°C or 83°C were included as reference controls. Molecular weight markers (in kDa) are also indicated. **F.** Quantification of NUDT5 stabilization at

83°C by ITDRF CETSA with TH5427 (blue), TH10184 (purple), and DDD2 (red). n=2 experiments shown (individual points) with lines-of-best-fit. NUDT5 abundance at 37°C was used to set normalization at 100% (grey).

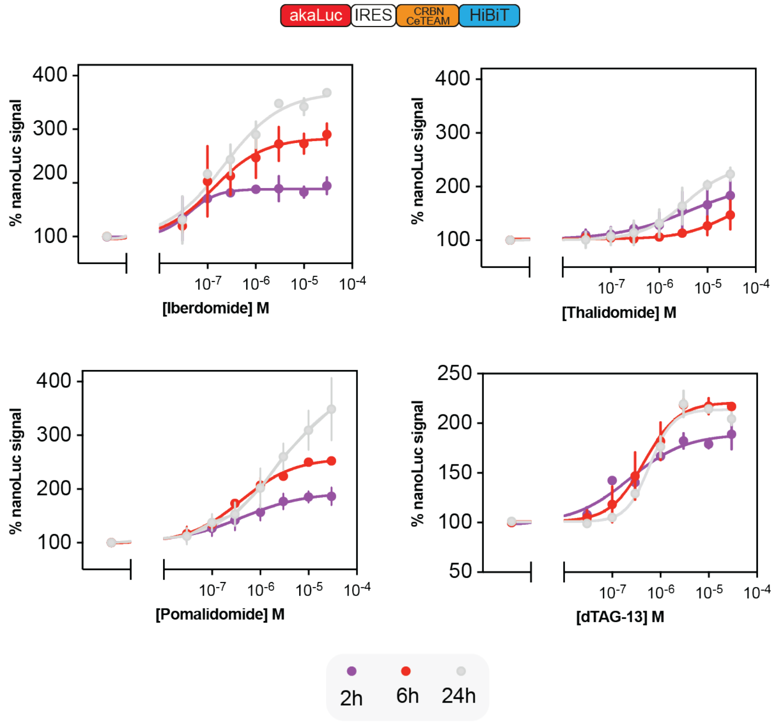

**Supplemental Figure 6. Time-dependent dose-response analysis of ligand-binding to the luminescent CRBN-CeTEAM biosensor.** U-2 OS cells expressing CRBN-CeTEAM reporter were treated with iberdomide, thalidomide, pomalidomide, or dTAG-13 at concentrations from 3 nM to 30 μM for 2, 6, or 24 hours. NanoLuc signal was measured and normalized to DMSO controls after akaLuc signal normalization. Data represent mean ± SD (n=3) with line of best fitted dose–response curves. Doxycycline (0.75 μg/mL) was added 24 hours prior to ligand treatment to induce LgBiT expression.

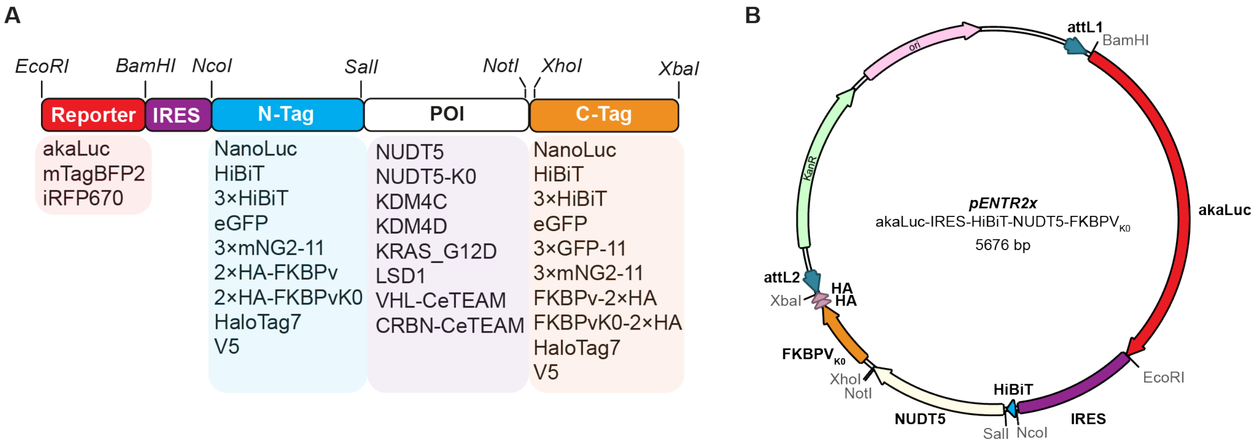

**Supplemental Figure 7. Modular cloning strategy for reporter construct generation. A.** Schematic representation of the multiple cloning site (MCS) from the pENTR2x-based bicistronic expression vector used for generating reporter constructs. The MCS includes an internal ribosomal entry site (IRES) to allow co-expression of a reference protein and fusion protein with the protein of interest (POI). Restriction sites are annotated for modular insertion: BamHI–EcoRI for reference proteins, NcoI–SalI for N-terminal tags, SalI–NotI for POIs and XhoI-XbaI for C-terminal tags. Genes inserted at each site for all expression constructs used in this study are listed below the corresponding restriction pairs. This design enables rapid assembly and variation of reporter constructs for degradation and CeTEAM assays. **B.** Vector map of the pENTR2x vector encoding the luminescence degradation reporter towards NUDT5.

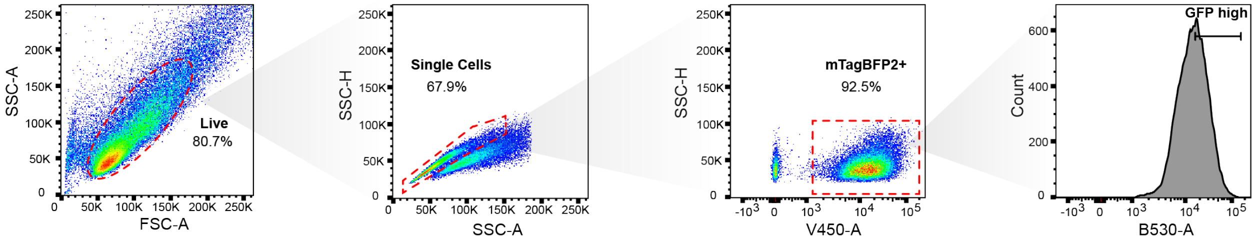

**Supplemental Figure 8. Gating strategy for flow cytometry analysis of HaloTag reporters.** Live, single cells were first gated to exclude debris and doublets. Transduced cells were identified by mTagBFP2 fluorescence, and within this population, GFP+ cells (or 3×mNG2_11 complemented with mNG2_1-10) were gated based on fluorescence in the B530 (GFP) channel. For degradation analysis, the top 50% of GFP+ cells (=GFP high), by fluorescence intensity, were selected.

### Supplemental Tables

#### Supplemental Table 1. NUDT5 PROTACs with chemical structure and SMILEs

| NAME | STRUCTURE | SMILE |
|---|---|---|
| **DDD1** | 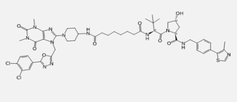 | ClC1=C(Cl)C=CC(C2=NN=C(CN3C(N4CCC(NC(=O)CCCCCCC(=O)N[C@H](C(=O)N5[C@H](C(=O)NCC6=CC=C(C7=C(C)N=CS7)C=C6)C[C@@H](O)C5)C(C)(C)C)CC4)=NC4=C3C(=O)N(C)C(=O)N4C)O2)=C1 |
| **DDD2** | 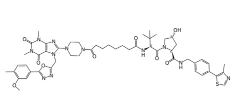 | S1C(C2=CC=C(CNC(=O)[C@H]3N(C(=O)[C@@H](NC(=O)CCCCCCC(=O)N4CCN(C5=NC6=C(N5CC5=NN=C(C7=CC(OC)=C(C)C=C7)O5)C(=O)N(C)C(=O)N6C)CC4)C(C)(C)C)C[C@H](O)C3)C=C2)=C(C)N=C1 |
| **DDD3** | 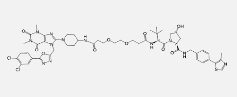 | ClC1=C(Cl)C=CC(C2=NN=C(CN3C(N4CCC(NC(=O)CCOCCOCCC(=O)N[C@H](C(=O)N5[C@H](C(=O)NCC6=CC=C(C7=C(C)N=CS7)C=C6)C[C@@H](O)C5)C(C)(C)C)CC4)=NC4=C3C(=O)N(C)C(=O)N4C)O2)=C1 |
| **DDD4** | 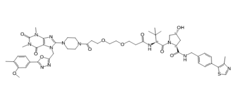 | S1C(C2=CC=C(CNC(=O)[C@H]3N(C(=O)[C@@H](NC(=O)CCOCCOCCC(=O)N4CCN(C5=NC6=C(N5CC5=NN=C(C7=CC(OC)=C(C)C=C7)O5)C(=O)N(C)C(=O)N6C)CC4)C(C)(C)C)C[C@H](O)C3)C=C2)=C(C)N=C1 |
| **DDD5** | 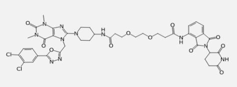 | ClC1=C(Cl)C=CC(C2=NN=C(CN3C(N4CCC(NC(=O)CCOCCOCCC(=O)NC5=C6C(=O)N(C7C(=O)NC(=O)CC7)C(=O)C6=CC=C5)CC4)=NC4=C3C(=O)N(C)C(=O)N4C)O2)=C1 |
| **DDD6** | 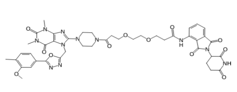 | O=C1N(C)C(=O)C2=C(N1C)N=C(N1CCN(C(=O)CCOCCOCCC(=O)NC3=C4C(=O)N(C5C(=O)NC(=O)CC5)C(=O)C4=CC=C3)CC1)N2CC1=NN=C(C2=CC(OC)=C(C)C=C2)O1 |
| **DDD7** | 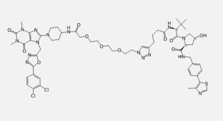 | ClC1=C(Cl)C=CC(C2=NN=C(CN3C(N4CCC(NC(=O)COCCOCCOCCN5N=NC(CCCC(=O)N[C@H](C(=O)N6[C@H](C(=O)NCC7=CC=C(C8=C(C)N=CS8)C=C7)C[C@@H](O)C6)C(C)(C)C)=C5)CC4)=NC4=C3C(=O)N(C)C(=O)N4C)O2)=C1 |
| **DDD8** | 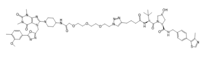 | S1C(C2=CC=C(CNC(=O)[C@H]3N(C(=O)[C@@H](NC(=O)CCCC4=CN(CCOCCOCCOCC(=O)NC5CCN(C6=NC7=C(N6CC6=NN=C(C8=CC(OC)=C(C)C=C8)O6)C(=O)N(C)C(=O)N7C)CC5)N=N4)C(C)(C)C)C[C@H](O)C3)C=C2)=C(C)N=C1 |
| **DDD9** | 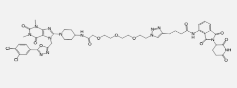 | ClC1=C(Cl)C=CC(C2=NN=C(CN3C(N4CCC(NC(=O)COCCOCCOCCN5N=NC(CCCC(=O)NC6=C7C(=O)N(C8C(=O)NC(=O)CC8)C(=O)C7=CC=C6)=C5)CC4)=NC4=C3C(=O)N(C)C(=O)N4C)O2)=C1 |
| **DDD10** | 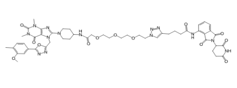 | O=C1N(C)C(=O)C2=C(N1C)N=C(N1CCC(NC(=O)COCCOCCOCCN3N=NC(CCCC(=O)NC4=C5C(=O)N(C6C(=O)NC(=O)CC6)C(=O)C5=CC=C4)=C3)CC1)N2CC1=NN=C(C2=CC(OC)=C(C)C=C2)O1 |

| | | |
|---|---|---|
| **DDD11** | 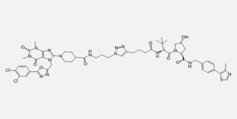 | ClC1=C(Cl)C=CC(C2=NN=C(CN3C(N4CCC(C(=O)NCCCN5N=NC(CCCC(=O)N[C@H](C(=O)N6[C@H](C(=O)NCC7=CC=C(C8=C(C)N=CS8)C=C7)C[C@@H](O)C6)C(C)(C)C)=C5)CC4)=NC4=C3C(=O)N(C)C(=O)N4C)O2)=C1 |
| **DDD12** | 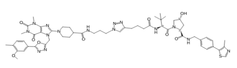 | S1C(C2=CC=C(CNC(=O)[C@H]3N(C(=O)[C@@H](NC(=O)CCCC4=CN(CCCNC(=O)C5CCN(C6=NC7=C(N6CC6=NN=C(C8=CC(OC)=C(C)C=C8)O6)C(=O)N(C)C(=O)N7C)CC5)N=N4)C(C)(C)C)C[C@H](O)C3)C=C2)=C(C)N=C1 |
| **DDD13** | 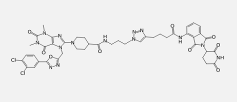 | ClC1=C(Cl)C=CC(C2=NN=C(CN3C(N4CCC(C(=O)NCCCN5N=NC(CCCC(=O)NC6=C7C(=O)N(C8C(=O)NC(=O)CC8)C(=O)C7=CC=C6)=C5)CC4)=NC4=C3C(=O)N(C)C(=O)N4C)O2)=C1 |
| **DDD14** | 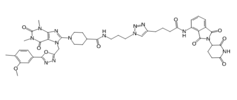 | O=C1N(C)C(=O)C2=C(N1C)N=C(N1CCC(C(=O)NCCCN3N=NC(CCCC(=O)NC4=C5C(=O)N(C6C(=O)NC(=O)CC6)C(=O)C5=CC=C4)=C3)CC1)N2CC1=NN=C(C2=CC(OC)=C(C)C=C2)O1 |
| **DDD15** | 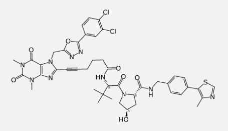 | ClC1=C(Cl)C=CC(C2=NN=C(CN3C(C#CCCCC(=O)N[C@H](C(=O)N4[C@H](C(=O)NCC5=CC=C(C6=C(C)N=CS6)C=C5)C[C@@H](O)C4)C(C)(C)C)=NC4=C3C(=O)N(C)C(=O)N4C)O2)=C1 |
| **DDD16** | 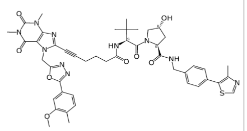 | S1C(C2=CC=C(CNC(=O)[C@H]3N(C(=O)[C@@H](NC(=O)CCCC#CC4=NC5=C(N4CC4=NN=C(C6=CC(OC)=C(C)C=C6)O4)C(=O)N(C)C(=O)N5C)C(C)(C)C)C[C@H](O)C3)C=C2)=C(C)N=C1 |
| **DDD17** | 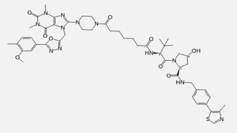 | S1C(C2=CC=C(CNC(=O)[C@H]3N(C(=O)[C@@H](NC(=O)CCCCCC(=O)N4CCN(C5=NC6=C(N5CC5=NN=C(C7=CC(OC)=C(C)C=C7)O5)C(=O)N(C)C(=O)N6C)CC4)C(C)(C)C)C[C@H](O)C3)C=C2)=C(C)N=C1 |
| **DDD18** | 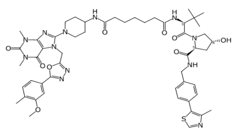 | S1C(C2=CC=C(CNC(=O)[C@H]3N(C(=O)[C@@H](NC(=O)CCCCCC(=O)NC4CCN(C5=NC6=C(N5CC5=NN=C(C7=CC(OC)=C(C)C=C7)O5)C(=O)N(C)C(=O)N6C)CC4)C(C)(C)C)C[C@H](O)C3)C=C2)=C(C)N=C1 |
| **DDD19** | 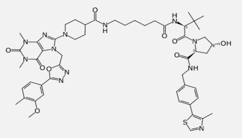 | S1C(C2=CC=C(CNC(=O)[C@H]3N(C(=O)[C@@H](NC(=O)CCCCCNC(=O)C4CCN(C5=NC6=C(N5CC5=NN=C(C7=CC(OC)=C(C)C=C7)O5)C(=O)N(C)C(=O)N6C)CC4)C(C)(C)C)C[C@H](O)C3)C=C2)=C(C)N=C1 |
| **DDD20** | 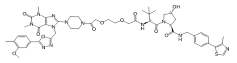 | S1C(C2=CC=C(CNC(=O)[C@H]3N(C(=O)[C@@H](NC(=O)COCCOCC(=O)N4CCN(C5=NC6=C(N5CC5=NN=C(C7=CC(OC)=C(C)C=C7)O5)C(=O)N(C)C(=O)N6C)CC4)C(C)(C)C)C[C@H](O)C3)C=C2)=C(C)N=C1 |
| **DDD21** | 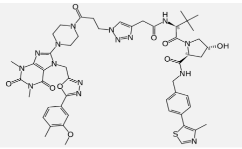 | S1C(C2=CC=C(CNC(=O)[C@H]3N(C(=O)[C@@H](NC(=O)CC4=CN(CCC(=O)N5CCN(C6=NC7=C(N6CC6=NN=C(C8=CC(OC)=C(C)C=C8)O6)C(=O)N(C)C(=O)N7C)CC5)N=N4)C(C)(C)C)C[C@H](O)C3)C=C2)=C(C)N=C1 |

| | | |
|---|---|---|
| **DDD22** | 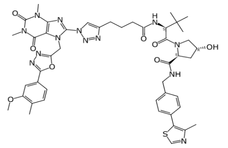 | S1C(C2=CC=C(CNC(=O)[C@H]3N(C(=O)[C@@H](NC(=O)CCCC4=CN(C5=NC6=C(N5CC5=NN=C(C7=CC(OC)=C(C)C=C7)O5)C(=O)N(C)C(=O)N6C)N=N4)C(C)(C)C)C[C@H](O)C3)C=C2)=C(C)N=C1 |
| **DDD23** | 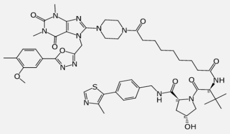 | S1C(C2=CC=C(CNC(=O)[C@H]3N(C(=O)[C@@H](NC(=O)CCCCCCCC(=O)N4CCN(C5=NC6=C(N5CC5=NN=C(C7=CC(OC)=C(C)C=C7)O5)C(=O)N(C)C(=O)N6C)CC4)C(C)(C)C)C[C@H](O)C3)C=C2)=C(C)N=C1 |
| **DDD24** | 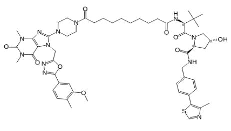 | S1C(C2=CC=C(CNC(=O)[C@H]3N(C(=O)[C@@H](NC(=O)CCCCCCCCC(=O)N4CCN(C5=NC6=C(N5CC5=NN=C(C7=CC(OC)=C(C)C=C7)O5)C(=O)N(C)C(=O)N6C)CC4)C(C)(C)C)C[C@H](O)C3)C=C2)=C(C)N=C1 |
| **DDD25** | 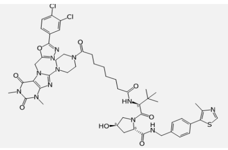 | O=C(CCCCCCC(=O)N[C@@H](C(C)(C)C)C(=O)N1C[C@H](O)C[C@H]1C(=O)NCC1=CC=C(C2=C(C)N=CS2)C=C1)N1CCN(C2=NC3=C(C(=O)N(C)C(=O)N3C)N2CC2=NN=C(C3=CC=C(Cl)C(Cl)=C3)O2)CC1 |
| **DDD26** | 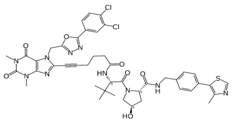 | ClC1=C(Cl)C=CC(C2=NN=C(CN3C(C#CCCCC(=O)N[C@H](C(=O)N4[C@H](C(=O)NCC5=CC=C(C6=C(C)N=CS6)C=C5)C[C@@H](O)C4)C(C)(C)C)=NC4=C3C(=O)N(C)C(=O)N4C)O2)=C1 |
| **DDD27** | 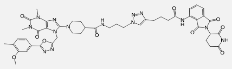 | O=C1N(C)C(=O)C2=C(N1C)N=C(N1CCC(C(=O)NCCCN3N=NC(CCCC(=O)NC4=C5C(=O)N(C6C(=O)NC(=O)CC6)C(=O)C5=CC=C4)=C3)CC1)N2CC1=NN=C(C2=CC(OC)=C(C)C=C2)O1 |
| **DDD28** | 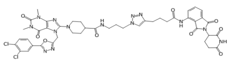 | ClC1=C(Cl)C=CC(C2=NN=C(CN3C(N4CCC(C(=O)NCCCN5N=NC(CCCC(=O)NC6=C7C(=O)N(C8C(=O)NC(=O)CC8)C(=O)C7=CC=C6)=C5)CC4)=NC4=C3C(=O)N(C)C(=O)N4C)O2)=C1 |
| **DDD29** | 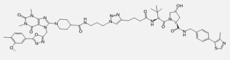 | S1C(C2=CC=C(CNC(=O)[C@H]3N(C(=O)[C@@H](NC(=O)CCCC4=CN(CCCNC(=O)C5CCN(C6=NC7=C(N6CC6=NN=C(C8=CC(OC)=C(C)C=C8)O6)C(=O)N(C)C(=O)N7C)CC5)N=N4)C(C)(C)C)C[C@H](O)C3)C=C2)=C(C)N=C1 |
| **DDD30** | 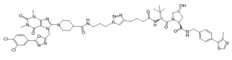 | ClC1=C(Cl)C=CC(C2=NN=C(CN3C(N4CCC(C(=O)NCCCN5N=NC(CCCC(=O)N[C@H](C(=O)N6[C@H](C(=O)NCC7=CC=C(C8=C(C)N=CS8)C=C7)C[C@@H](O)C6)C(C)(C)C)=C5)CC4)=NC4=C3C(=O)N(C)C(=O)N4C)O2)=C1 |
| **DDD31** | 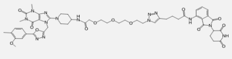 | O=C1N(C)C(=O)C2=C(N1C)N=C(N1CCC(NC(=O)COCCOCCOCCN3N=NC(CCCC(=O)NC4=C5C(=O)N(C6C(=O)NC(=O)CC6)C(=O)C5=CC=C4)=C3)CC1)N2CC1=NN=C(C2=CC(OC)=C(C)C=C2)O1 |
| **DDD32** | 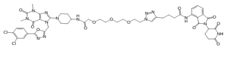 | ClC1=C(Cl)C=CC(C2=NN=C(CN3C(N4CCC(NC(=O)COCCOCCOCCN5N=NC(CCCC(=O)NC6=C7C(=O)N(C8C(=O)NC(=O)CC8)C(=O)C7=CC=C6)=C5)CC4)=NC4=C3C(=O)N(C)C(=O)N4C)O2)=C1 |
| **DDD33** | 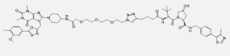 | S1C(C2=CC=C(CNC(=O)[C@H]3N(C(=O)[C@@H](NC(=O)CCCC4=CN(CCOCCOCCOCC(=O)NC5CCN(C6=NC7=C(N6CC6=NN=C(C8=CC(OC)=C(C)C=C8)O6)C(=O)N(C)C(=O)N7C)CC5)N=N4)C(C)(C)C)C[C@H](O)C3)C=C2)=C(C)N=C1 |

| | | |
|---|---|---|
| **DDD34** | 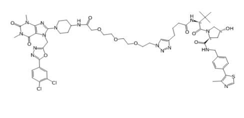 | ClC1=C(Cl)C=CC(C2=NN=C(CN3C(N4CCC(NC(=O)COCCOCCOCCN5N=NC(CCCC(=O)N[C@H](C(=O)N6[C@H](C(=O)NCC7=CC=C(C8=C(C)N=CS8)C=C7)C[C@@H](O)C6)C(C)(C)C)=C5)CC4)=NC4=C3C(=O)N(C)C(=O)N4C)O2)=C1 |
| **DDD35** | 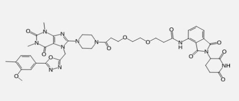 | O=C1N(C)C(=O)C2=C(N1C)N=C(N1CCN(C(=O)CCOCCOCCC(=O)NC3=C4C(=O)N(C5C(=O)NC(=O)CC5)C(=O)C4=CC=C3)CC1)N2CC1=NN=C(C2=CC(OC)=C(C)C=C2)O1 |
| **DDD36** | 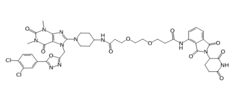 | ClC1=C(Cl)C=CC(C2=NN=C(CN3C(N4CCC(NC(=O)CCOCCOCCC(=O)NC5=C6C(=O)N(C7C(=O)NC(=O)CC7)C(=O)C6=CC=C5)CC4)=NC4=C3C(=O)N(C)C(=O)N4C)O2)=C1 |
| **DDD37** | 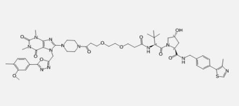 | S1C(C2=CC=C(CNC(=O)[C@H]3N(C(=O)[C@@H](NC(=O)CCOCCOCCC(=O)N4CCN(C5=NC6=C(N5CC5=NN=C(C7=CC(OC)=C(C)C=C7)O5)C(=O)N(C)C(=O)N6C)CC4)C(C)(C)C)C[C@H](O)C3)C=C2)=C(C)N=C1 |
| **DDD38** | 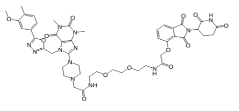 | C(OCCNC(CN1CCN(C2=NC3=C(C(=O)N(C)C(=O)N3C)N2CC2=NN=C(C3=CC=C(C)C(OC)=C3)O2)CC1)=O)COCCNC(COC1=C2C(=CC=C1)C(=O)N(C1CCC(=O)NC1=O)C2=O)=O |
| **DDD39** | 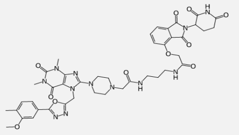 | C1=CC=C2C(=C1OCC(=O)NCCCNC(CN1CCN(C3=NC4=C(C(=O)N(C)C(=O)N4C)N3CC3=NN=C(C4=CC=C(C)C(OC)=C4)O3)CC1)=O)C(=O)N(C1CCC(=O)NC1=O)C2=O |
| **DDD40** | 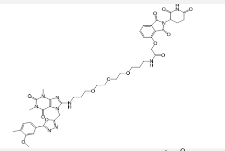 | N(CCCOCCOCCOCCCNC(COC1=C2C(=CC=C1)C(=O)N(C1CCC(=O)NC1=O)C2=O)=O)C1=NC2=C(C(=O)N(C)C(=O)N2C)N1CC1=NN=C(C2=CC=C(C)C(OC)=C2)O1 |
| **DDD41** | 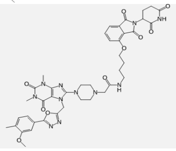 | O(C1=C2C(=CC=C1)C(=O)N(C1CCC(=O)NC1=O)C2=O)CCCCNC(CN1CCN(C2=NC3=C(C(=O)N(C)C(=O)N3C)N2CC2=NN=C(C3=CC=C(C)C(OC)=C3)O2)CC1)=O |
| **DDD42** | 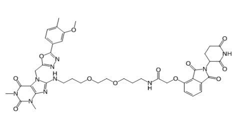 | C(CCOCCOCCCNC(COC1=C2C(=CC=C1)C(=O)N(C1CCC(=O)NC1=O)C2=O)=O)NC1=NC2=C(C(=O)N(C)C(=O)N2C)N1CC1=NN=C(C2=CC=C(C)C(OC)=C2)O1 |
| **DDD43** | 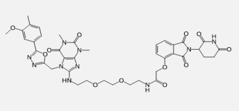 | C(OCCNC1=NC2=C(C(=O)N(C)C(=O)N2C)N1CC1=NN=C(C2=CC=C(C)C(OC)=C2)O1)COCCNC(COC1=C2C(=CC=C1)C(=O)N(C1CCC(=O)NC1=O)C2=O)=O |
| **DDD44** | 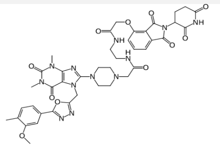 | C1=CC=C2C(=C1OCC(=O)NCCNC(CN1CCN(C3=NC4=C(C(=O)N(C)C(=O)N4C)N3CC3=NN=C(C4=CC=C(C)C(OC)=C4)O3)CC1)=O)C(=O)N(C1CCC(=O)NC1=O)C2=O |

| | | |
|---|---|---|
| **DDD45** | 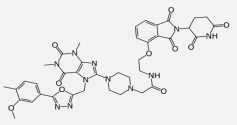 | O(C1=C2C(=CC=C1)C(=O)N(C1CCC(=O)NC1=O)C2=O)CCNC(CN1CCN(C2=NC3=C(C(=O)N(C)C(=O)N3C)N2CC2=NN=C(C3=CC=C(C)C(OC)=C3)O2)CC1)=O |
| **DDD46** | 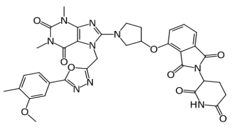 | O(C1=C2C(=CC=C1)C(=O)N(C1CCC(=O)NC1=O)C2=O)C1CN(C2=NC3=C(C(=O)N(C)C(=O)N3C)N2CC2=NN=C(C3=CC=C(C)C(OC)=C3)O2)CC1 |
| **DDD47** | 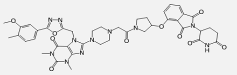 | C1(=O)N(C)C(=O)C2=C(N1C)N=C(N1CCN(CC(=O)N3CC(OC4=C5C(=CC=C4)C(=O)N(C4CCC(=O)NC4=O)C5=O)CC3)CC1)N2CC1=NN=C(C2=CC=C(C)C(OC)=C2)O1 |
| **DDD48** | 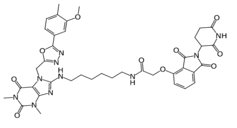 | C(CNC1=NC2=C(C(=O)N(C)C(=O)N2C)N1CC1=NN=C(C2=CC=C(C)C(OC)=C2)O1)CCCCNC(COC1=C2C(=CC=C1)C(=O)N(C1CCC(=O)NC1=O)C2=O)=O |
| **DDD49** | 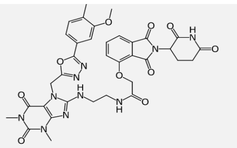 | C1(=O)N(C)C(=O)C2=C(N1C)N=C(NCCNC(COC1=C3C(=CC=C1)C(=O)N(C1CCC(=O)NC1=O)C3=O)=O)N2CC1=NN=C(C2=CC=C(C)C(OC)=C2)O1 |
| **DDD50** | 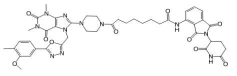 | O=C1N(C)C(=O)C2=C(N1C)N=C(N1CCN(C(=O)CCCCCCC(=O)NC3=C4C(=O)N(C5C(=O)NC(=O)CC5)C(=O)C4=CC=C3)CC1)N2CC1=NN=C(C2=CC(OC)=C(C)C=C2)O1 |

**Supplemental Table 2. Ubiquitin proteomic data on NUDT5**

| POSITION(S) | SOURCE | DESCRIPTION | Category |
|---|---|---|---|
| 14 | PTMeXchange | Ubiquitinated lysineCombined Sources | GOLD |
| 27 | PTMeXchange | Ubiquitinated lysineCombined Sources | BRONZE |
| 33 | PTMeXchange | Ubiquitinated lysineCombined Sources | GOLD |
| 42 | PTMeXchange | Ubiquitinated lysineCombined Sources | GOLD |
| 50 | PTMeXchange | Ubiquitinated lysineCombined Sources | GOLD |
| 55 | PTMeXchange | Ubiquitinated lysineCombined Sources | BRONZE |
| 81 | PTMeXchange | Ubiquitinated lysineCombined Sources | SILVER |
| 161 | PTMeXchange | Ubiquitinated lysineCombined Sources | SILVER |
| 175 | PTMeXchange | Ubiquitinated lysineCombined Sources | SILVER |
| 210 | PTMeXchange | Ubiquitinated lysineCombined Sources | GOLD |
| 218 | PTMeXchange | Ubiquitinated lysineCombined Sources | GOLD |

| Category | Criteria |
|---|---|
| Gold | Identified in *n* or more datasets at ≤1% FLR |
| Silver | Identified in at least *m* datasets, but less than *n*datasets at ≤1% FLR |
| Bronze | Identified in one or more datasets at ≤5% FLR, but not meeting Silver or Gold criteria |

Source (Uniprot: Q9UKK9: 2026-07-29)

**Supplemental Table 3. Luminescence signal in cell degradation data of NUDT5 variants by different PROTACs**

| | DDD2 | | | FKBPd3 | | | dTAG13 | | |
|---|---|---|---|---|---|---|---|---|---|
| | Average | SD | n= | Average | SD | n= | Average | SD | n= |
| **WT** | **100,00** | **11,10** | **3** | **100,00** | **1,43** | **3** | **100,00** | **7,07** | **3** |
| **K0** | **0,00** | **10,62** | **3** | **0,00** | **11,07** | **3** | **0,00** | **11,41** | **3** |
| H3 | 127,34 | 19,52 | 3 | 103,16 | 0,96 | 3 | 97,78 | 8,55 | 3 |
| H1 | 116,39 | 5,84 | 3 | 101,10 | 1,19 | 3 | 89,34 | 3,69 | 3 |
| H4 | 91,93 | 27,26 | 3 | 104,50 | 0,90 | 3 | 67,20 | 8,83 | 3 |
| H2 | 124,42 | 15,81 | 3 | 103,12 | 1,62 | 3 | 63,42 | 8,22 | 3 |

| | DDD2 | | | FKBPd3 | | | dTAG13 | | |
|---|---|---|---|---|---|---|---|---|---|
| | Average | SD | n= | Average | SD | n= | Average | SD | n= |
| **WT** | **100,00** | **22,71** | **14** | **100,00** | **3,38** | **12** | **100,00** | **8,66** | **14** |
| **K0** | **0,00** | **22,79** | **14** | **0,00** | **11,54** | **14** | **0,00** | **10,35** | **14** |
| K14 | 117,87 | 7,27 | 3 | 101,10 | 2,19 | 3 | 33,41 | 3,43 | 3 |
| K27 | 54,80 | 9,55 | 3 | 76,91 | 1,70 | 3 | 34,68 | 1,09 | 3 |
| K30 | 88,02 | 20,93 | 3 | 100,08 | 2,53 | 3 | 27,16 | 9,75 | 3 |
| K33 | 87,63 | 12,87 | 3 | 99,46 | 0,34 | 3 | 36,62 | 12,52 | 3 |
| K42 | 84,33 | 19,39 | 3 | 84,47 | 1,90 | 3 | 61,43 | 10,91 | 3 |
| K50 | 114,05 | 21,06 | 3 | 102,78 | 1,39 | 3 | 29,69 | 4,30 | 3 |
| K55 | 115,94 | 18,71 | 3 | 101,98 | 1,40 | 3 | 37,80 | 4,50 | 3 |
| K81 | 107,20 | 23,86 | 3 | 103,36 | 1,95 | 3 | 28,08 | 4,15 | 3 |
| K120 | 119,79 | 18,46 | 3 | 106,04 | 2,36 | 3 | 41,77 | 10,45 | 3 |
| K159 | 98,68 | 3,48 | 3 | 95,71 | 1,01 | 3 | 48,01 | 5,44 | 3 |
| K161 | 116,48 | 3,55 | 3 | 97,08 | 2,96 | 3 | 36,32 | 1,54 | 3 |
| K175 | 96,42 | 5,13 | 3 | 94,22 | 1,42 | 3 | 34,62 | 4,04 | 3 |

| | | | | | | | | | |
|---|---|---|---|---|---|---|---|---|---|
| K205 | 89,83 | 6,19 | 2 | 86,15 | 0,74 | 2 | 31,96 | 0,58 | 2 |
| K210 | 120,39 | 10,71 | 3 | 103,02 | 5,30 | 3 | 78,60 | 4,95 | 3 |
| K218 | 117,97 | 18,37 | 3 | 102,38 | 5,81 | 3 | 85,85 | 2,28 | 3 |

**Supplemental Table 4. Plasmids and nucleic acids used in this study**

| PLASMIDS | SOURCE |
|---|---|
| **ENTR plasmids** | |
| pENTR1A | Addgene # 17398 |
| pENTR2x | this work |
| **Other plasmids** | |
| pMuLE Lenti Dest eGFP | Addgene # 62175 |
| akaLuc | PMID: 39643609 |
| NanoLuc | PMID: 39643609 |
| mTagBFP2 | Addgene # 34632 |
| iRFP670 | PMID: 27918552 |
| shRNA plasmid | PMID: 27918552 |
| **DEST plasmids** | |
| *Empty* | |
| pCW57.1 | Addgene # 41393 |
| pLenti CMV Blast | Addgene # 17451 |
| Ef1a-Tta3G-P2A-Blast | PMID: 34348895 |
| *Tags* | |
| pCW57.1_mNG2_1-10 | this work |
| pCW57.1_LgBit | this work |
| pCW57.1_GFP1-10 | this work |
| **NUDT5** | |
| *pLenti CMV Blast* | |
| akaLuc-IRES-FKBPV -NUDT5- NanoLuc | this work |
| akaLuc-IRES-NanoLuc-NUDT5-FKBPV | this work |
| akaLuc_IRES_FKBPV-NUDT5-cHiBiT | this work |
| akaLuc-IRES-HiBiT-NUDT5-FKBPV | this work |
| akaLuc-IRES-HiBiT(K0)-NUDT5-FKBPV | this work |
| akaLuc-IRES-HiBiT(79)-NUDT5-FKBPV | this work |
| akaLuc-IRES-HiBiT(99)-NUDT5-FKBPV | this work |
| akaLuc-IRES-HiBiT(NP)-NUDT5-FKBPV | this work |
| akaLuc-IRES-$FKBPV_{K0}$-NUDT5-HiBiT | this work |
| akaLuc-IRES-HiBiT-NUDT5-$FKBPV_{K0}$ | this work |
| *Ef1a-Tta3G-P2A-Blast* | |
| akaLuc-IRES-HiBiT-NUDT5-$FKBPV_{K0}$ | this work |
| akaLuc-IRES-HiBiT-NUDT5_K0-$FKBPV_{K0}$ | this work |
| akaLuc-IRES-HiBiT-NUDT5R14K-$FKBPV_{K0}$ | this work |
| akaLuc-IRES-HiBiT-NUDT5R27K-$FKBPV_{K0}$ | this work |

| | |
|---|---|
| akaLuc-IRES-HiBiT-NUDT5R30K-FKBPV$_{K0}$ | this work |
| akaLuc-IRES-HiBiT-NUDT5R33K-FKBPV$_{K0}$ | this work |
| akaLuc-IRES-HiBiT-NUDT5R42K-FKBPV$_{K0}$ | this work |
| akaLuc-IRES-HiBiT-NUDT5R50K-FKBPV$_{K0}$ | this work |
| akaLuc-IRES-HiBiT-NUDT5R55K-FKBPV$_{K0}$ | this work |
| akaLuc-IRES-HiBiT-NUDT5R81K-FKBPV$_{K0}$ | this work |
| akaLuc-IRES-HiBiT-NUDT5R120K-FKBPV$_{K0}$ | this work |
| akaLuc-IRES-HiBiT-NUDT5R159K-FKBPV$_{K0}$ | this work |
| akaLuc-IRES-HiBiT-NUDT5R161K-FKBPV$_{K0}$ | this work |
| akaLuc-IRES-HiBiT-NUDT5R175K-FKBPV$_{K0}$ | this work |
| akaLuc-IRES-HiBiT-NUDT5R205K-FKBPV$_{K0}$ | this work |
| akaLuc-IRES-HiBiT-NUDT5R210K-FKBPV$_{K0}$ | this work |
| akaLuc-IRES-HiBiT-NUDT5R218K-FKBPV$_{K0}$ | this work |
| **Other genes** | |
| *pLenti CMV Blast* | |
| akaLuc_IRES_FKBPV$_{K0}$-KDM4C-cHiBiT | this work |
| akaLuc_IRES_FKBPV$_{K0}$ -KDM4D-cHiBiT | this work |
| akaLuc_IRES_FKBPV$_{K0}$-KRAS_G12D-cHiBiT | this work |
| akaLuc_IRES_FKBPV$_{K0}$-LSD1-cHiBiT | this work |
| akaLuc_IRES-HiBiT-KDM4C-FKBPV$_{K0}$ | this work |
| akaLuc_IRES-HiBiT-KDM4D-FKBPV$_{K0}$ | this work |
| akaLuc_IRES-HiBiT-KRASG12-FKBPV$_{K0}$ | this work |
| akaLuc_IRES_HiBiT-LSD1-FKBPV$_{K0}$ | this work |
| eGFP_IRES_CRBN L190F-V5 | this work |
| eGFP_IRES_VHL_R167Q-V5 | this work |
| akaLuc_IRES_V5-CRBN_L190F-HiBiT | this work |
| akaLuc_IRES_V5-VHL_R167Q-HiBiT | this work |
| mTagBFP2-IRES-eGFP-HaloTag | this work |
| mTagBFP2-IRES-HaloTag-eGFP | this work |
| mTagBFP2-IRES-3XmNG2_11-HaloTag | this work |
| mTagBFP2-IRES-HaloTag-3XmNG2_11 | this work |
| akaLuc-IRES-NanoLuc-HaloTag | this work |
| akaLuc-IRES-HaloTag-NanoLuc | this work |
| akaLuc-IRES-HiBiT-HaloTag | this work |
| akaLuc-IRES-HaloTag-HiBiT | this work |

| Synthetic | Gene Fragments Sequence (5'→3') |
|---|---|
| **MCS_p2x** | TCGAGgCCGGATCCACCTTTGAATTCGCCACCatggGAACCAATTCAGTCGACGGCGGCGGCGGCAGCGC |

| | |
|---|---|
| **IRES** | GCCCCTCTCCCTCCCCCCCCCCTAACGTTACTGGCCGAAGCCGCTTGGAATAAGGCCGGTGTG CGTTTGTCTATATGTTATTTTCCACCATATTGCCGTCTTTTGGCAATGTGAGGGCCCGGAAACCTG GCCCTGTCTTCTTGACGAGCATTCCTAGGGGTCTTTCCCCTCTCGCCAAAGGAATGCAAGGTCT GTTGAATGTCGTGAAGGAAGCAGTTCCTCTGGAAGCTTCTTGAAGACAAACAACGTCTGTAGCG ACCCTTTGCAGGCAGCGGAACCCCCCACCTGGCGACAGGTGCCTCTGCGGCCAAAAGCCACG TGTATAAGATACACCTGCAAAGGCGGCACAACCCCAGTGCCACGTTGTGAGTTGGATAGTTGTG GAAAGAGTCAAATGGCTCTCCTCAAGCGTATTCAACAAGGGGCTGAAGGATGCCCAGAAGGTAC CCCATTGTATGGGATCTGATCTGGGGCCTCGGTGCACATGCTTTACATGTGTTTAGTCGAGGTTA AAAAAACGTCTAGGCCCCCCGAACCACGGGGACGTGGTTTTCCTTTGAAAAACACGATGATAATA TGGCCACAAgcca |
| **LgBiT** | ATGATGGTGTTCACCCTGGAGGACTTCGTGGGCGACTGGGAGCAGACCGCCGCCTACAACCTG GACCAGGTGCTGGAGCAGGGCGGCGTGAGCAGCCTGCTGCAGAACCTGGCCGTGAGCGTGAC CCCCATCCAGAGGATCGTGAGGAGCGGCGAGAACGCCCTGAAGATCGACATCCACGTGATCAT CCCCTACGAGGGCCTGAGCGCCGACCAGATGGCCCAGATCGAGGAGGTGTTCAAGGTGGTGTA CCCCGTGGACGACCACCACTTCAAGGTGATCCTGCCCTACGGCACCCTGGTGATCGACGGCGT GACCCCCAACATGCTCAACTACTTCGGCAGGCCCTACGAGGGCATCGCCGTGTTCGACGGCAA GAAGATCACCGTGACCGGCACCCTGTGGAACGGCAACAAGATCATCGACGAGAGGCTGATCAC CCCCGACGGCAGCATGCTGTTCAGGGTGACCATCAACAGCTAA |
| **HiBiT** | ATGGTGAGCGGCTGGAGGCTGTTCAAGAAGATCAGC |
| **3xHiBiT** | ATGGTAAGTGGATGGCGCCTCTTCAAGAAAATTTCTGGAGGGTCTGGGGGAGTGTCTGGCTGGA GGCTTTTTAAAAAAATCTCCGGTGGGTCTGGGGGGGTATCTGGATGGAGACTGTTTAAGAAAATT TCAGGAGGTAGCGGAGGA |
| **HiBiT(K0)** | ATGGTGAGCGGCTGGAGGCTGTTCAGAAGGATC |
| **HiBiT(79)** | ATGGTGAGCGGCTGGAGGCTGTTCAAGAAGATC |
| **HiBiT(99)** | ATGGTGACCGGCTACAGGCTGTTCGAGAAGATC |
| **HiBiT(NP)** | ATGGGCGTGACCGGCTGGAGGCTGTGCGAGAGAATCCTGGCC |
| **GFP_1-10** | ATGTCCAAAGGAGAAGAACTGTTTACCGGTGTTGTGCCAATTTTGGTTGAACTCGATGGTGATGT CAACGGACATAAGTTCTCAGTGAGAGGCGAAGGAGAAGGTGACGCCACCATTGGAAAATTGACT CTTAAATTCATCTGTACTACTGGTAAACTTCCTGTACCgTGGCCGACTCTCGTAACAACGCTTACG TACGGAGTTCAGTGCTTTTCGAGATACCCAGACCATATGAAAAGACATGACTTTTTTAAGTCGGCT ATGCCTGAAGGTTACGTGCAAGAAAGAACAATTTCGTTCAAAGATGATGGAAAATATAAAACTAGA GCAGTTGTTAAATTTGAAGGAGATACTTTGGTTAACCGCATTGAACTGAAAGGAACAGATTTTAAA GAAGATGGTAATATTCTTGGACACAAACTCGAATACAATTTTAATAGTCATAACGTATACATCACTGC TGATAAGCAAAAGAACGGAATTAAAGCGAATTTCACAGTACGCCATAATGTAGAAGATGGCAGTGT TCAACTTGCCGACCATTACCAACAAAACACCCCTATTGGAGACGGTCCGGTACTTCTTCCTGATA ATCACTACCTCTCAACACAAACAGTCCTGAGCAAAGATCCAAATGAAAAATAA |
| **mNG2 _1-10** | ATGGTCAGTAAAGGCGAAGAGGACAACATGGCCAGCTTACCTGCGACGCACGAACTGCATATATT CGGCAGTATAAACGGGGTGGATTTTGACATGGTCGGCCAGGGCACGGGCAACCCTAATGATGGC TACGAGGAGCTAAATCTCAAGAGTACAAAAGGTGACCTGCAGTTTAGCCCGTGGATTCTGGTGC CTCATATCGGGTACGGATTTCACCAGTATCTCCCTTACCCAGACGGTATGTCGCCATTCCAGGCC GCAATGGTGGATGGTAGCGGATACCAAGTCCATCGCACAATGCAATTTGAAGATGGCGCTTCACT TACAGTTAACTACAGATACACGTATGAGGGCTCCCACATTAAGGGAGAGGCACAGGTGATGGGG ACCGGCTTTCCAGCAGACGGGCCCGTGATGACAAACACGCTTACCGCTGCAGACTGGTGCATG TCCAAGAAGACCTACCCCAACGATAAGACTATTATCTCAACCTTTAAGTGGTCTTATACTACTGTTA ATGGGAAACGGTATCGTTCTACAGCGCGAACCACCTACACCTTTGCTAAACCTATGGCGGCCAAT TATCTGAAAAATCAGCCCATGTATGTATTTAGGAAGACCGAGTTAAAGCATTCTATGTGA |
| **3xGFP_11** | ATGGTCCTTCATGAGTATGTAAATGCTGCTGGGATTACAGGTGGCTCTGGAGGTAGAGATCATATG GTTCTCCACGAATACGTTAACGCCGCAGGCATCACTGGCGGTAGTGGAGGACGCGACCATATGG TACTACATGAATATGTCAATGCAGCCGGAATAACC |

| | |
|---|---|
| **3xmNG2_11** | ATGGCAACCGAACTCAACTTCAAGGAATGGCAAAAAGCCTTTACTGATATGATGGGCGGCAGCG<br>GGGGAACAGAGCTTAATTTTAAGGAGTGGCAGAAGGCTTTTACCGACATGATGGGTGGGTCCGG<br>AGGCACAGAGCTGAACTTCAAAGAATGGCAGAAAGCCTTCACGGACATGATG |
| **2xHA_FKBPv-n** | GGAGGCGGAGGTTCTGGCGGCGGAGGGTCCGGCGGTGGGGGGAGCGGAGTGCAGGTGGAA<br>ACCATCTCCCCAGGAGACGGGCGCACCTTCCCCAAGCGCGGCCAGACCTGCGTGGTGCACTAC<br>ACCGGGATGCTTGAAGATGGAAAGAAAGTTGATTCCTCCCGGGACAGAAACAAGCCCTTTAAGT<br>TTATGCTAGGCAAGCAGGAGGTGATCCGAGGCTGGGAAGAAGGGGTTGCCCAGATGAGTGTGG<br>GTCAGAGAGCCAAACTGACTATATCTCCAGATTATGCCTATGGTGCCACTGGGCACCCAGGCATC<br>ATCCCACCACATGCCACTCTCGTCTTCGATGTGGAGCTTCTAAAACTGGAAGGCGGCTACCCCTA<br>CGACGTGCCCGACTACGCCGGCTATCCGTATGATGTCCCGGACTATGCAGGCTAA |
| **2xHA_FKBPvK0-n** | CCATGGCATACCCCTACGACGTGCCCGACTACGCCGGCTATCCGTATGATGTCCCGGACTATGCA<br>GGCGGCGGCGGAGTGCAGGTGGAAACCATCTCCCCAGGAGACGGGCGCACCTTCCCCAGACG<br>CGGCCAGACCTGCGTGGTGCACTACACCGGGATGCTTGAAGATGGAAgAAgAGTTGATTCCTCC<br>CGGGACAGAAACAgaCCCTTTAgaTTTATGCTAGGCAgaCAGGAGGTGATCCGAGGCTGGGAAGA<br>AGGGGTTGCCCAGATGAGTGTGGGTCAGAGAGCCAgACTGACTATATCTCCAGATTATGCCTATG<br>GTGCCACTGGGCACCCAGGCATCATCCCACCACATGCCACTCTCGTCTTCGATGTGGAGCTTCT<br>AAgACTGGAAGGAGGCGGAGGTTCTGGCGGCGGAGGGTCCGGCGGTGGGGGGAGCGTCGAC |
| **FKBPv_2xHA-c** | GGAGGCGGAGGTTCTGGCGGCGGAGGGTCCGGCGGTGGGGGGAGCGGAGTGCAGGTGGAA<br>ACCATCTCCCCAGGAGACGGGCGCACCTTCCCCAAGCGCGGCCAGACCTGCGTGGTGCACTAC<br>ACCGGGATGCTTGAAGATGGAAAGAAAGTTGATTCCTCCCGGGACAGAAACAAGCCCTTTAAGT<br>TTATGCTAGGCAAGCAGGAGGTGATCCGAGGCTGGGAAGAAGGGGTTGCCCAGATGAGTGTGG<br>GTCAGAGAGCCAAACTGACTATATCTCCAGATTATGCCTATGGTGCCACTGGGCACCCAGGCATC<br>ATCCCACCACATGCCACTCTCGTCTTCGATGTGGAGCTTCTAAAACTGGAAGGCGGCTACCCCTA<br>CGACGTGCCCGACTACGCCGGCTATCCGTATGATGTCCCGGACTATGCAGGCTAA |
| **FKBPvK0_2xHA-c** | CATATACTCGAGGGAGGCGGAGGTTCTGGCGGCGGAGGGTCCGGCGGTGGGGGGAGCGGAG<br>TGCAGGTGGAAACCATCTCCCCAGGAGACGGGCGCACCTTCCCCAGACGCGGCCAGACCTGC<br>GTGGTGCACTACACCGGGATGCTTGAAGATGGAAgAAgAGTTGATTCCTCCCGGGACAGAAACAg<br>aCCCTTTAgaTTTATGCTAGGCAgaCAGGAGGTGATCCGAGGCTGGGAAGAAGGGGTTGCCCAGA<br>TGAGTGTGGGTCAGAGAGCCAgACTGACTATATCTCCAGATTATGCCTATGGTGCCACTGGGCAC<br>CCAGGCATCATCCCACCACATGCCACTCTCGTCTTCGATGTGGAGCTTCTAAgACTGGAAGGCG<br>GCTACCCCTACGACGTGCCCGACTACGCCGGCTATCCGTATGATGTCCCGGACTATGCAGGCTA<br>ATCTAGATATACA |
| **V5** | GGTAAGCCTATCCCTAACCCTCTCCTCGGTCTCGATTCTACG |
| **NUDT5** | ATGGAGTCTCAAGAGCCCACTGAgAGTTCACAaAACGGCAAACAATACATCATCAGCGAGGAGCT<br>GATATCTGAAGGTAAGTGGGTCAAGTTGGAAAAGACCACATATATGGACCCAACTGGCAAGACAA<br>GAACTTGGGAAAGCGTCAAAAGGACCACCCGGAAAGAGCAGACCGCAGATGGTGTTGCGGTCA<br>TACCAGTATTACAaCGGACACTCCATTACGAATGCATCGTTCTGGTAAAGCAATTTCGGCCCCCTA<br>TGGGGGGATATTGCATAGAGTTTCCAGCGGGGTTAATTGACGACGGAGAAACCCCTGAAGCTGC<br>CGCATTAAGGGAACTGGAGGAAGAAACAGGTTACAAAGGCGACATAGCTGAGTGTTCCCCCGCG<br>GTCTGCATGGACCCGGGATTATCAAATTGTACAATTCACATaGTaACCGTTACTATcAAtGGGGATGA<br>CGCCGAAAACGCACGTCCAAAGCCTAAACCCGGAGATGGTGAGTTCGTtGAGGTtATCTCCCTGC<br>CCAAAAACGATCTATTGCAGCGGCTCGACGCCCTTGTGGCCGAAGAGCACCTGACCGTGGACG<br>CTCGTGTGTATAGTTAtGCGCTGGCTTTGAAgCATGCAAACGCTAAGCCTTTCGAGGTGCCCTTCC<br>TCAAATTC |

| | |
|---|---|
| **NUDT5-K0** | ATGGAGAGCCAAGAACCAACGGAATCTTCTCAGAATGGCAgACAGTATATCATTTCAGAGGAGTT<br>AATTTCAGAAGGAAgATGGGTCAgaCTTGAAAgAACAACGTACATGGAcCCTACTGGTAgAACTAGA<br>ACTTGGGAATCAGTGAgACGTACAACCAGGAgAGAGCAGACTGCGGATGGTGTCGCGGTCATCC<br>CCGTGCTGCAGAGAACACTTCACTATGAGTGTATCGTTCTGGTcAgACAGTTCCGACCACCAATG<br>GGGGGCTACTGCATAGAGTTCCCTGCAGGTCTCATAGATGATGGTGAAACCCCAGAAGCAGCTG<br>CTCTCCGGGAGCTTGAAGAAGAAACTGGCTACAgAGGGGACATTGCCGAATGTTCTCCAGCGGT<br>CTGTATGGACCCAGGCTTGTCAAACTGTACTATACACATCGTGACAGTCACCATTAACGGAGATG<br>ATGCCGAAAACGCAAGGCCGAgGCCAAgGCCAGGGGATGGAGAGTTTGTGGAAGTCATTTCTTT<br>ACCCAgGAATGACCTGCTGCAGAGACTTGATGCTCTGGTAGCTGAAGAACATCTCACAGTGGAC<br>GCCAGGGTCTATTCCTACGCTCTAGCACTGAgACAcGCAAATGCAAgGCCATTTGAAGTGCCCTT<br>CTTGAgATTT |
| **HaloTag7** | ATGGAAATCGGAACCGGGTTTCCGTTTGATCCACACTATGTGGAAGTACTTGGCGAGAGAATGCA<br>TTACGTCGATGTCGGACCTAGAGATGGCACACCTGTGCTGTTTCTTCATGGCAATCCGACATCAA<br>GCTACGTTTGGCGGAATATCATCCCGCACGTTGCTCCTACACATCGGTGTATCGCCCCGGATTTG<br>ATCGGTATGGGTAAAAGCGATAAACCAGACCTGGGTTATTTCTTCGACGATCACGTACGCTTCATG<br>GATGCGTTTATTGAGGCTCTCGGACTTGAGGAAGTTGTGTTGGTCATACACGACTGGGGTTCAG<br>CACTGGGATTTCATTGGGCTAAGCGGAACCCCGAGCGAGTGAAAGGTATTGCGTTCATGGAGTT<br>CATACGACCTATACCAACGTGGGACGAATGGCCCGAGTTCGCACGGGAAACATTCCAGGCGTTC<br>AGGACGACCGACGTTGGGCGAAAACTGATCATTGATCAAAACGTCTTCATAGAAGGAACACTCC<br>CTATGGGCGTTGTAAGGCCTTTGACTGAGGTCGAGATGGATCACTACCGCGAACCTTTCCTCAA<br>CCCCGTTGATCGCGAGCCCTTGTGGCGCTTTCCAAACGAACTGCCGATAGCGGGAGAACCTGC<br>AAATATCGTCGCGCTTGTCGAGGAGTATATGGACTGGCTTCATCAATCTCCCGTGCCCAAACTGC<br>TCTTTTGGGGGACTCCCGGAGTACTGATCCCACCCGCGGAGGCTGCACGACTCGCGAAAAGCC<br>TGCCGAATTGCAAAGCTGTTGACATTGGCCCTGGATTGAATTTGCTTCAGGAGGACAATCCTGAC<br>CTTATTGGTAGCGAAATCGCCAGGTGGCTCTCCACGCTGGAAATCTCCTAA |
| **KRAS** | ATGACAGAGTATAAGCTCGTCGTCGTTGGTGCCGATGGTGTCGGCAAATCAGCGCTTACGATACA<br>GCTCATACAGAATCACTTTGTTGATGAGTACGACCCCACTATTGAGGATTCTTATAGAAAACAAGT<br>AGTAATTGACGGGGAGACTTGTTTATTGGACATCCTTGACACCGCAGGGCAAGAGGAATATTCCG<br>CAATGAGGGACCAATACATGCGCACAGGAGAGGGATTCCTGTGCGTTTTTGCCATAAATAACACT<br>AAGTCGTTTGAAGATATTCACCATTACCGTGAACAGATCAAGCGCGTGAAAGATAGCGAGGATGT<br>CCCGATGGTGCTGGTGGGCAACAAGTGCGACCTGCCTAGTCGGACCGTGGATACAAAACAGGC<br>TCAGGACCTAGCCCGGTCTTATGGAATCCCATTCATTGAGACATCCGCCAAAACCAGACAGAGAG<br>TGGAGGACGCTTTCTACACCCTGGTGAGGGAAATTCGACAGTACAGGCTGAAAAAGATCAGCAA<br>GGAAGAAAAGACCCCCGGCTGTGTGAAAATCAAGAAGTGCATCATCATG |

| | |
|---|---|
| **KDM4C** | ATGGAGGTTGCAGAGGTGGAGAGCCCTCTGAATCCATCATGTAAGATTATGACTTTCAGACCGTC<br>AATGGAGGAGTTCCGTGAGTTTAATAAATATTTAGCATATATGGAATCGAAGGGCGCCCATCGTGC<br>TGGCCTAGCTAAAGTGATCCCCCCGAAGGAATGGAAGCCTCGGCAATGTTATGATGATATTGATAA<br>TCTGTTAATACCGGCACCGATCCAGCAGATGGTTACAGGGCAGAGCGGCTTATTTACGCAATACA<br>ACATCCAGAAAAAAGCAATGACTGTAAAGGAGTTCAGGCAGCTAGCTAATTCAGGGAAGTACTGT<br>ACCCCGAGGTACCTCGACTACGAAGACTTAGAGCGCAAATATTGGAAGAATCTAACCTTCGTCGC<br>CCCTATCTACGGTGCCGATATCAACGGCAGCATCTATGATGAAGGGGTTGACGAGTGGAACATAG<br>CTCGGCTTAACACTGTGCTAGACGTGGTGGAGGAGGAGTGCGGGATCAGTATCGAGGGAGTGA<br>ATACTCCTTACTTGTACTTCGGAATGTGGAAAACAACTTTTGCCTGGCACACTGAGGACATGGAT<br>CTCTATTCCATAAACTACCTGCATTTTGGTGAGCCAAAGTCGTGGTACGCCATTCCACCAGAACA<br>CGGCAAACGTCTGGAACGGCTAGCTCAAGGGTTTTTCCCAAGTAGTTCTCAGGGGTGCGACGC<br>CTTTCTACGTCACAAAATGACTCTTATCAGCCCAAGTGTCTTGAAAAAGTACGGTATTCCGTTTGA<br>TAAGATCACCCAGGAGGCCGGCGAGTTTATGATAACTTTTCCTTATGGATACCATGCTGGCTTTAA<br>CCACGGGTTTAATTGCGCTGAGTCCACCAACTTTGCTACCGTAAGGTGGATAGACTACGGCAAAG<br>TTGCTAAACTGTGTACCTGTCGTAAGGATATGGTGAAGATCTCAATGGACATCTTCGTCCGCAAAT<br>TTCAACCAGACAGATACCAGCTTTGGAAACAGGGGAAAGATATTTATACGATTGATCACACCAAAC<br>CAACCCCCGCCTCTACTCCTGAAGTCAAAGCGTGGCTTCAAAGGAGAAGGAAGGTGCGAAAAG<br>CAAGCCGTTCGTTCCAGTGCGCTCGATCGACGTCAAAAAGGCCCAAAGCTGACGAAGAAGAAG<br>AAGTCTCCGACGAGGTCGATGGAGCGGAAGTCCCAAACCCCGATAGCGTTACTGATGACCTCAA<br>GGTCAGTGAAAAATCTGAAGCTGCTGTGAAACTGAGAAATACAGAAGCGAGTAGTGAGGAGGAA<br>TCATCAGCATCACGGATGCAGGTGGAACAAAACTTATCCGACCACATCAAACTTAGCGGAAACTC<br>ATGCTTGAGTACCTCCGTCACGGAGGATATCAAAACCGAAGATGACAAGGCCTATGCCTACCGGT<br>CGGTACCATCTATCTCTTCCGAGGCCGACGATAGTATCCCTCTGTCTAGTGGTTACGAAAAGCCA<br>GAGAAGTCCGACCCCTCCGAACTGTCATGGCCTAAATCACCTGAGTCATGCTCCAGCGTGGCCG<br>AATCAAACGGAGTGCTCACGGAGGGGGAGGAATCAGACGTGGAATCTCACGGAAATGGATTGG<br>AGCCTGGCGAAATTCCGGCTGTGCCTTCTGGTGAGAGAAATTCCTTTAAGGTCCCTTCTATAGCC<br>GAAGGCGAAAACAAAACAAGTAAGAGTTGGCGTCATCCCCTCTCACGACCTCCTGCTCGATCAC<br>CTATGACGCTTGTGAAACAGCAGGCGCCCTCCGATGAAGAGCTGCCTGAGGTTCTTTCCATTGA<br>GGAGGAAGTCGAGGAGACGGAAAGCTGGGCCAAACCCCTCATACATCTTTGGCAAACAAAGTC<br>CCCCAATTTTGCTGCTGAGCAGGAGTATAATGCTACCGTGGCACGAATGAAACCTCACTGTGCGA<br>TTTGCACCCTTCTGATGCCATACCATAAACCTGACTCCTCCAATGAGGAAAACGACGCTAGATGG<br>GAGACAAAGTTGGATGAGGTGGTGACATCTGAAGGCAAAACAAAACCCTTGATTCCCGAGATGT<br>GTTTCATATATTCAGAAGAAAACATCGAGTACTCCCCTCCCAATGCGTTTTTGGAGGAGGATGGC<br>ACTTCACTGCTGATCAGCTGCGCAAAATGTTGCGTTCGCGTGCACGCCTCCTGCTATGGCATCC<br>CATCCCATGAGATTTGCGACGGCTGGTTGTGCGCACGATGTAAGCGGAACGCGTGGACAGCAG<br>AGTGCTGCCTCTGTAACCTCCGAGGCGGAGCCCTCAAGCAGACCAAGAACAATAAGTGGGCCC<br>ATGTGATGTGTGCCGTGGCCGTGCCCGAGGTACGGTTCACCAACGTCCCCGAGCGCACACAAA<br>TAGACGTCGGACGAATACCCCTGCAGCGGCTGAAGCTGAAGTGTATTTTTTGTCGCCATCGAGT<br>CAAGAGAGTGAGCGGGGCCTGTATCCAGTGTTCTTACGGTAGATGTCCCGCCTCCTTCCACGTT<br>ACATGTGCGCATGCCGCCGGAGTTCTGATGGAACCCGATGATTGGCCCTATGTTGTAAACATTAC<br>CTGCTTCAGGCACAAGGTTAATCCCAACGTGAAGAGCAAGGCATGCGAAAAGGTTATTAGCGTC<br>GGACAGACAGTTATTACAAAGCACCGGAACACTCGCTACTACAGCTGTCGGGTGATGGCTGTCA<br>CATCTCAGACTTTCTATGAGGTGATGTTTGATGATGGCTCTTTCAGCAGAGATACCTTCCCAGAG<br>GATATTGTTAGCAGAGACTGCCTCAAGCTGGGCCCCCCAGCCGAGGGGGAGGTAGTGCAGGTC<br>AAGTGGCCCGACGGAAAGCTTTACGGGGCAAAGTATTTCGGCAGCAATATAGCACACATGTACCA<br>GGTAGAGTTCGAAGACGGCAGCCAGATAGCTATGAAGCGGGAAGATATATATACACTCGACGAAG<br>AACTTCCAAAGCGCGTCAAGGCCCGCTTTTCGACAGCATCTGACATGAGATTCGAGGATACCTTC<br>TATGGTGCAGACATCATCCAGGGCGAGAGGAAGAGGCAGAGAGTTCTGTCTAGCAGGTTCAAGA<br>ACGAGTACGTCGCCGACCCAGTGTACAGGACCTTTCTGAAGAGCTCTTTTCAGAAGAAGTGCCA<br>GAAGAGGCAG |

**KDM4D**

ATGGGTTCTGAAGACCACGGTGCTCAAAATCCGTCCTGCAAAATTATGACCTTCCGACCCACGAT
GGAGGAGTTTAAGGATTTTAATAAGTACGTAGCTTACATAGAGTCTCAGGGGGCTCATAGAGCCG
GCCTTGCAAAAATTATCCCTCCCAAAGAGTGGAAACCCCGACAGACCTATGACGACATCGATGAT
GTGGTGATACCTGCGCCCATCCAGCAGGTGGTGACGGGTCAAAGTGGTCTCTTCACTCAATACA
ACATCCAGAAGAAGGCCATGACGGTTGGAGAATATAGAAGATTAGCGAACAGCGAAAAATATTGC
ACCCCCCGCCATCAGGACTTCGACGACCTAGAGAGAAAGTATTGGAAGAATTTGACCTTTGTGA
GCCCCATCTACGGTGCGGACATCTCCGGATCTCTCTATGATGACGATGTAGCCCAGTGGAACATC
GGATCACTGAGGACCATTTTAGATATGGTGGAAAGGGAATGCGGCACCATTATCGAGGGTGTGAA
CACGCCATACCTCTACTTCGGTATGTGGAAAACTACCTTCGCCTGGCACACCGAAGATATGGATC
TTTATTCCATTAATTACCTCCACTTTGGTGAACCGAAATCATGGTATGCTATCCCACCTGAGCACG
GGAAGAGACTGGAGCGCCTGGCGATCGGTTTCTTTCCTGGAAGCTCCCAGGGGTGCGATGCCT
TCCTGAGGCATAAAATGACATTAATTTCCCCGATCATTTTAAAGAAGTACGGGATACCATTCAGCC
GTATCACTCAGGAAGCCGGGGAGTTCATGATTACGTTTCCTTACGGATACCATGCGGGGTTCAAC
CACGGGTTTAACTGTGCGGAGTCTACAAATTTCGCTACATTGAGATGGATTGACTACGGTAAAGTT
GCAACCCAATGCACTTGCCGCAAGGATATGGTGAAAATTAGTATGGACGTATTTGTCCGCATTTTG
CAACCCGAACGCTATGAGTTGTGGAAACAGGGGAAGGATCTCACAGTGCTTGATCACACCCGCC
CCACTGCGCTCACATCACCGGAGCTTAGTTCATGGTCCGCGTCGCGGGCCAGTCTTAAGGCTAA
GTTGCTCCGGAGATCTCATCGCAAACGAAGCCAACCAAAAAAACCCAAACCAGAGGACCCCAAA
TTCCCTGGCGAAGGAACCGCAGGGGCTGCACTGCTGGAGGAAGCCGGAGGCTCCGTGAAGGA
GGAGGCTGGCCCTGAAGTTGACCCAGAAGAGGAGGAGGAGGAGCCTCAGCCACTTCCGCATG
GGCGGGAAGCTGAGGGCGCCGAAGAGGATGGTCGGGGCAAGCTGAGACCGACTAAGGCCAAG
AGTGAAAGGAAGAAGAAATCTTTTGGGCTGCTTCCCCCCCAGCTGCCGCCACCTCCCGCACACT
TTCCTTCTGAAGAGGCCCTGTGGCTTCCCTCTCCTTTAGAACCACCAGTGCTGGGTCCTGGACC
AGCCGCAATGGAAGAGAGTCCACTTCCCGCTCCCCTGAATGTCGTGCCACCCGAAGTACCTAGC
GAAGAGCTCGAAGCTAAACCGCGGCCTATCATTCCAATGCTATACGTCGTACCGCGACCGGGCA
AGGCTGCATTCAATCAGGAGCACGTCTCATGCCAACAGGCCTTCGAGCACTTTGCGCAAAAAGG
ACCTACATGGAAGGAACCTGTCAGCCCTATGGAGCTGACGGGCCCTGAGGACGGAGCCGCAAG
CTCTGGAGCCGGCCGGATGGAGACTAAGGCAAGAGCGGGCGAGGGTCAGGCCCCCTCAACAT
TCTCCAAATTGAAAATGGAAATCAAGAAGAGCAGGCGACACCCTTTAGGCCGACCCCCCACTAG
ATCGCCCCTGTCCGTGGTGAAGCAGGAGGCCTCTTCAGATGAGGAGGCCAGCCCTTTTAGCGG
GGAAGAGGATGTGAGTGATCCTGACGCTCTCCGACCACTGTTGAGCCTGCAGTGGAAAAACCG
CGCCGCTAGCTTTCAGGCCGAGAGGAAGTTTAACGCAGCTGCAGCCAGGACCGAACCTTACTG
TGCCATCTGTACATTGTTTTACCCGTATTGTCAAGCGCTCCAGACAGAAAAAGAGGCACCAATTG
CGTCCCTGGGTGAAGGATGTCCCGCTACCCTTCCGAGCAAGAGCCGACAGAAGACGAGGCCCC
TTATCCCAGAAATGTGTTTCACAAGCGGGGGCGAGAATACTGAGCCTCTACCAGCAAATTCGTAT
ATCGGCGATGACGGCACCAGTCCCCTAATTGCATGCGGAAAGTGCTGCCTGCAGGTGCACGCC
TCGTGTTATGGTATTCGCCCCGAGTTAGTTAATGAGGGCTGGACATGTTCCAGATGTGCGGCCCA
TGCATGGACCGCCGAGTGCTGTTTGTGTAATCTGAGAGGTGGGGCCCTACAAATGACAACTGAC
CGGAGATGGATTCATGTTATCTGTGCAATTGCCGTACCTGAGGCCCGCTTCCTTAATGTCATCGA
GCGGCACCCCGTGGACATTTCCGCTATACCAGAACAGCGTTGGAAACTGAAATGCGTCTATTGTC
GGAAACGGATGAAGAAAGTATCTGGAGCTTGTATCCAGTGTTCCTACGAGCATTGTAGTACTAGC
TTCCACGTAACCTGCGCCCACGCCGCCGGCGTTCTCATGGAGCCAGATGATTGGCCATACGTCG
TCTCTATTACGTGCCTCAAGCACAAAAGCGGCGGCCACGCAGTCCAGCTACTCCGCGCAGTGTC
CCTTGGACAGGTGGTTATCACCAAGAACCGCAATGGGCTCTATTATAGGTGCCGGGTAATCGGG
GCTGCCTCCCAAACCTGCTATGAAGTCAACTTTGACGATGGCAGTTATTCTGACAATTTATACCCT
GAGAGCATTACGAGCAGGGACTGTGTCCAACTCGGACCCCCTAGTGAAGGAGAATTGGTGGAG
CTTCGTTGGACCGATGGAAATCTGTACAAAGCCAAATTCATCAGTTCGGTAACGTCGCATATATAC
CAGGTGGAATTTGAGGACGGCTCACAGTTGACAGTCAAGAGGGGAGACATTTTCACACTGGAG
GAGGAACTGCCGAAGCGGGTGAGGAGTAGACTTAGCCTGTCCACTGGGGCACCCCAAGAACC
GGCCTTCTCGGGAGAGGAGGCCAAAGCAGCGAAAAGACCCAGGGTCGGGACACCGCTGGCGA
CGGAGGATAGCGGCAGGAGCCAGGACTATGTGGCATTCGTCGAGTCCCTTTTGCAAGTACAGG
GGAGGCCCGGTGCTCCCTTT

| | |
|---|---|
| **LSD1** | ATGCTATCGGGGAAAAAAGCTGCTGCGGCTGCAGCTGCTGCCGCGGCAGCCGCGACAGGGAC<br>TGAAGCCGGTCCAGGAACTGCAGGCGGATCGGAAAACGGGAGCGAGGTTGCCGCCCAACCGG<br>CCGGGCTGTCTGGTCCGGCAGAGGTCGGGCCTGGGGCTGTGGGTGAACGGACACCTCGCAAG<br>AAGGAGCCTCCCAGAGCTTCCCCTCCAGGCGGACTTGCGGAACCACCAGGATCAGCGGGTCC<br>GCAAGCGGGTCCAACCGTAGTGCCCGGTTCAGCCACGCCTATGGAGACCGGTATAGCCGAAAC<br>ACCAGAAGGTAGGCGAACGAGTAGGAGGAAGCGTGCCAAGGTCGAGTACAGAGAAATGGACGA<br>ATCATTAGCCAACTTAAGCGAGGATGAGTATTACAGCGAGGAGGAACGCAACGCGAAGGCAGAA<br>AAGGAGAAGAAGCTCCCACCGCCCCCCCCTCAGGCACCCCCCGAAGAGGAAAATGAGAGTGA<br>GCCCGAAGAACCAAGTGGCGTGGAGGGCGCAGCCTTTCAGTCCCGCCTGCCCCACGACCGGA<br>TGACATCTCAAGAGGCCGCGTGTTTTCCGGATATAATTTCAGGGCCTCAGCAAACTCAAAAGGTC<br>TTCCTTTTCATTCGGAATCGTACTCTGCAATTATGGTTGGATAACCCAAAGATCCAATTAACCTTTG<br>AAGCTACCCTACAGCAACTTGAGGCGCCTTACAATAGCGATACTGTATTGGTACATAGGGTCCACT<br>CATATTTGGAAAGGCACGGTTTGATTAATTTTGGAATTTATAAACGCATTAAGCCCCTTCCCACTAA<br>AAAAACCGGAAAAGTCATCATCATCGGGTCCGGTGTTAGTGGCCTGGCGGCTGCCCGCCAACT<br>GCAGTCCTTTGGGATGGACGTCACTTTATTGGAAGCTAGAGACCGGGTTGGGGGCAGAGTCGC<br>CACATTTAGGAAAGGGAACTACGTAGCAGATCTGGGAGCTATGGTGGTGACTGGTCTGGGAGGC<br>AATCCTATGGCCGTCGTCTCTAAACAGGTCAACATGGAGCTCGCAAAAATCAAGCAAAAGTGTCC<br>ATTATATGAAGCTAACGGACAAGCGGTCCCGAAGGAAAAAGACGAAATGGTTGAGCAGGAGTTTA<br>ATCGTCTTTTGGAAGCTACCTCGTACCTTTCACATCAACTAGACTTCAACGTGCTAAACAATAAGC<br>CCGTTAGTCTGGGACAGGCGCTTGAGGTAGTTATACAGCTCCAAGAGAAGCACGTAAAAGATGA<br>GCAAATTGAACATTGGAAAAAGATTGTCAAAACCCAGGAAGAACTGAAGGAGCTCCTGAATAAGA<br>TGGTTAACCTGAAGGAAAAGATCAAGGAGTTACATCAGCAATACAAGGAAGCCAGTGAAGTCAAG<br>CCTCCACGGGACATTACCGCAGAGTTTCTTGTGAAAAGCAAGCACCGTGATCTTACAGCCCTGT<br>GCAAAGAATACGACGAGCTAGCAGAGACTCAGGGTAAACTGGAGGAGAAACTTCAGGAACTCGA<br>AGCAAATCCGCCCTCGGACGTGTACTTATCTAGTAGGGATAGACAGATCTTAGACTGGCACTTCG<br>CCAATCTCGAATTTGCGAACGCCACCCCCCTATCGACCTTATCATTGAAGCACTGGGATCAAGAC<br>GATGATTTCGAGTTTACTGGCAGCCATTTGACAGTGCGGAATGGTTATTCCTGCGTGCCCGTTGC<br>CCTGGCTGAGGGACTTGATATTAAACTGAATACTGCTGTAAGGCAGGTGCGTTACACTGCAAGCG<br>GATGCGAAGTGATAGCCGTGAATACTCGAAGTACGAGTCAGACATTCATTTACAAATGCGACGCC<br>GTGCTATGTACTTTACCCTTAGGCGTGCTGAAACAACAGCCGCCCGCTGTGCAGTTCGTGCCAC<br>CGCTGCCTGAGTGGAAGACTTCCGCCGTCCAGCGGATGGGATTTGGCAATCTGAACAAAGTAGT<br>CCTGTGTTTCGACCGGGTGTTTTGGGACCCATCAGTCAACCTTTTCGGCCACGTAGGCTCGACC<br>ACTGCCAGCCGGGGAGAGCTTTTCCTCTTCTGGAATTTGTATAAGGCTCCAATCTTACTGGCATT<br>GGTCGCCGGCGAGGCCGCCGGCATCATGGAAAACATCTCTGATGATGTGATAGTTGGTCGTTGC<br>CTCGCGATCCTCAAAGGTATCTTCGGCAGTTCTGCAGTCCCACAGCCTAAGGAGACTGTAGTGT<br>CCCGATGGAGGGCGGACCCCTGGGCAAGAGGTTCCTATAGCTACGTGGCAGCAGGCTCTTCTG<br>GAAATGATTACGACCTGATGGCGCAGCCTATTACGCCGGGACCCTCCATCCCGGGGGCACCTCA<br>ACCGATTCCTAGACTATTCTTTGCCGGGGAACATACAATCAGGAATTACCCAGCAACAGTCCACG<br>GCGCCTTGCTATCTGGTTTGAGAGAAGCGGGTAGGATTGCCGACCAATTTCTGGGGGCGATGTA<br>TACGCTACCGCGTCAGGCCACTCCAGGAGTTCCTGCTCAGCAGAGTCCCTCTAT |
| **CRBN** | ATGATGGCCGGCGAAGGAGATCAGCAGGACGCTGCGCACAACATGGGCAACCACCTGCCGCTC<br>CTGCCTGCAGAGAGTGAGGAAGAAGATGAAATGGAAGTTGAAGACCAGGATAGTAAAGAAGCCA<br>AAAAACCAAACATCATAAATTTTGACACCAGTCTGCCGACATCACATACATACCTAGGTGCTGATAT<br>GGAAGAATTTCATGGCAGGACTTTGCACGATGACGACAGCTGTCAGGTGATTCCAGTTCTTCCA<br>CAAGTGATGATGATCCTGATTCCCGGACAGACATTACCTCTTCAGCTTTTTCACCCTCAAGAAGT<br>CAGTATGGTGCGGAATTTAATTCAGAAAGATAGAACCTTTGCTGTTCTTGCATACAGCAATGTACA<br>GGAAAGGGAAGCACAGTTTGGAACAACAGCAGAGATATATGCCTATCGAGAAGAACAGGATTTTG<br>GAATTGAGATAGTGAAAGTGAAAGCAATTGGAAGACAAAGGTTCAAAGTCCTTGAGCTAAGAACA<br>CAGTCAGATGGAATCCAGCAAGCTAAAGTGCAAATTCTTCCCGAATGTGTGTTGCCTTCAACCAT<br>GTCTGCAGTTCAATTAGAATCCCTCAATAAGTGCCAGATATTTCCTTCAAAACCTGTCTCAAGAGA<br>AGACCAATGTTCATATAAATGGTGGCAGAAATACCAGAAGAGAAAGTTTCATTGTGCAAATCTAAC<br>TTCATGGCCTCGCTGGCTGTATTCCTTATATGATGCTGAGACCTTAATGGACAGAATCAAGAAACA<br>GCTACGTGAATGGGATGAAAATCTAAAAGATGATTCTCTTCCTTCAAATCCAATAGATTTTTCTTAC<br>AGAGTAGCTGCTTGTCTTCCTATTGATGATGTATTGAGGATTCAGCTCCTTAAAATTGGCAGTGCT<br>ATCCAGCGACTTCGCTGTGAATTAGACATTATGAATAAATGTACTTCCCTTTGCTGTAAACAATGTC<br>AAGAAACAGAAATAACAACCAAAAATGAAATATTCAGTTTATCCTTATGTGGGCCGATGGCAGCTT<br>ATGTGAATCCTCATGGATATGTGCATGAGACACTTACTGTGTATAAGGCTTGCAACTTGAATCTGAT<br>AGGCCGGCCTTCTACAGAACACAGCTGGTTTCCTGGGTATGCCTGGACTGTTGCCCAGTGTAAG<br>ATCTGTGCAAGCCATATTGGATGGAAGTTTACGGCCACCAAAAAAGACATGTCACCTCAAAAATTT<br>TGGGGCTTAACGCGATCTGCTCTGTTGCCCACGATCCCAGACACTGAAGATGAAATAAGTCCAG<br>ACAAAGTAATACTTTGCTTG |

| | |
|---|---|
| **VHL** | ATGCCCCGTCGCGCCGAAAACTGGGACGAGGCCGAGGTGGGCGCCGAGGAGGCTGGAGTGG<br>AGGAATATGGCCCAGAAGAGGATGGGGGCGAAGAGTCCGGTGCGGAGGAGTCGGGTCCCGAA<br>GAGAGCGGACCGGAGGAACTGGGGGCAGAGGAAGAAATGGAGGCTGGGAGGCCGCGGCCTG<br>TGCTTCGTTCAGTAAATTCCCGAGAGCCCAGTCAAGTTATATTTTGTAATCGGTCACCGCGAGTG<br>GTCTTACCTGTTTGGCTCAACTTCGACGGCGAGCCTCAACCTTACCCCACGCTGCCTCCCGGCA<br>CAGGAAGGCGCATCCACAGCTATAGGGGGCATCTATGGCTGTTCCGCGATGCTGGAACCCACGA<br>TGGCCTCCTCGTGAATCAGACTGAGTTATTTGTCCCATCTTTGAATGTAGACGGTCAGCCCATTTT<br>CGCCAACATTACACTCCCAGTGTACACCCTGAAGGAACGCTGCTTGCAGGTGGTGAGGAGTCTG<br>GTCAAGCCAGAGAACTACAGACGGCTTGACATCGTCCGCTCTCTGTATGAAGACCTGGAAGATC<br>ATCCAAACGTTCAGAAAGACCTGGAAAGACTTACTCAGGAAAGAATCGCACACCAGAGAATGGG<br>AGAT |

| **Site directed mutagenesis primers** | **Sequence (5'→3')** | **Sequence (3'→5')** |
|---|---|---|
| **NUDT5-K0** | **lysine to arginine** | |
| **NUDT5-K0_R14K** | CAGAATGGCAAACAGTATATCATTTCAGAGGAG | TACTGTTTGCCATTCTGAGAAGATTCCGTTG |
| **NUDT5-K0_R27K** | CAGAAGGAAAATGGGTCAGACTTGAAAG | GTCTGACCCATTTTCCTTCTGAAATTAACTC |
| **NUDT5-K0_R30K** | GGGTCAAACTTGAAAGAACAACGTAC | TCAAGTTTGACCCATCTTCCTTCTG |
| **NUDT5-K0_R33K** | CTTGAAAAAACAACGTACATGGACC | GTTGTTTTTTCAAGTCTGACCCATC |
| **NUDT5-K0_R42K** | CCTACTGGTAAAACTAGAACTTGGGAATC | GTTCTAGTTTTACCAGTAGGGTCCATGT |
| **NUDT5-K0_R50K** | CAGTGAAACGTACAACCAGGAGAGAG | TGGTTGTACGTTTCACTGATTCCCAAG |
| **NUDT5-K0_R55K** | AACCAGGAAAGAGCAGACTGCG | CTGCTCTTTCCTGGTTGTACGTCTC |
| **NUDT5-K0_R81K** | TGGTCAAACAGTTCCGACCAC | AACTGTTTGACCAGAACGATACACT |
| **NUDT5-K0_R120K** | GCTACAAAGGGGACATTGCCG | TCCCCTTTGTAGCCAGTTTCTTCTT |
| **NUDT5-K0_R159K** | GCCGAAGCCAAGGCCAGG | TTGGCTTCGGCCTTGCGTTT |
| **NUDT5-K0_R161K** | GCCAAAGCCAGGGGATGGAG | CCTGGCTTTGGCCTCGGCCTTGCGT |
| **NUDT5-K0_R175K** | TACCCAAGAATGACCTGCTGCA | CAGGTCATTCTTGGGTAAAGAAATGACTT |
| **NUDT5-K0_R205K** | CACTGAAACACGCAAATGCAAGGC | GCGTGTTTCAGTGCTAGAGCGTAG |
| **NUDT5-K0_R210K** | ATGCAAAGCCATTTGAAGTGCCC | AATGGCTTTGCATTTGCGTGTCTC |
| **NUDT5-K0_R218K** | TTCTTGAAATTTGCGGCCGCA | CCGCAAATTTCAAGAAGGGCACT |

| | | |
|---|---|---|
| **CeTEAM** | | |
| **VHL-CeTEAM_R167Q** | GTGGTGCAGAGTCTGGTCAAGCC | CAGACTCTGCACCACCTGCAAGC |
| **CRBN-CeTEAM_L190F** | GTGTGTTCCCTTCAACCATGTCTG | GAAGGGAACACACATTCGGGAAG |
| **shRNA** | **Sequence (5'→3')** | **Sequence (3'→5')** |
| **shVHL** | CCGGTATCACACTGCCAGTGTATACCTCGAGGTATACACTGGCAGTGTGATATTTTTG | AATTCAAAAATATCACACTGCCAGTGTATACCTCGAGGTATACACTGGCAGTGTGATA |
